# Computationally validated SARS-CoV-2 CTL and HTL Multi-Patch Vaccines designed by reverse epitomics approach, shows potential to cover large ethnically distributed human population worldwide

**DOI:** 10.1101/2020.09.06.284992

**Authors:** Sukrit Srivastava, Sonia Verma, Mohit Kamthania, Deepa Agarwal, Ajay Kumar Saxena, Michael Kolbe, Sarman Singh, Ashwin Kotnis, Brijesh Rathi, Seema. A. Nayar, Ho-Joon Shin, Kapil Vashisht, Kailash C Pandey

## Abstract

**Background:** The SARS-CoV-2 (Severe Acute Respiratory Syndrome Coronavirus 2) is a positive-sense single-stranded RNA coronavirus responsible for the ongoing 2019-2020 COVID-19 outbreak. The highly contagious COVID-19 disease has spread to 216 countries in less than six months. Though several vaccine candidates are being claimed, an effective vaccine is yet to come. In present study we have designed and theoretically validated novel Multi-Patch Vaccines against SARS-CoV-2.

**Methodology:** A novel reverse epitomics approach, “overlapping-epitope-clusters-to-patches” method is utilized to identify multiple antigenic regions from the SARS-CoV-2 proteome. These antigenic regions are here termed as “Ag-Patch or Ag-Patches”, for Antigenic Patch or Patches. The identification of Ag-Patches is based on clusters of overlapping epitopes rising from a particular region of SARS-CoV-2 protein. Further, we have utilized the identified Ag-Patches to design Multi-Patch Vaccines (MPVs), proposing a novel methodology for vaccine design and development. The designed MPVs were analyzed for immunologically crucial parameters, physiochemical properties and cDNA constructs.

**Results:** We identified 73 CTL (Cytotoxic T-Lymphocyte), 49 HTL (Helper T-Lymphocyte) novel Ag-Patches from the proteome of SARS-CoV-2. The identified Ag-Patches utilized to design MPVs cover 768 (518 CTL and 250 HTL) overlapping epitopes targeting different HLA alleles. Such large number of epitope coverage is not possible for multi-epitope vaccines. The large number of epitopes covered implies large number of HLA alleles targeted, and hence large ethnically distributed human population coverage. The MPVs:Toll-Like Receptor ectodomain complex shows stable nature with numerous hydrogen bond formation and acceptable root mean square deviation and fluctuation. Further, the cDNA analysis favors high expression of the MPVs constructs in human cell line.

**Conclusion:** Highly immunogenic novel Ag-Patches are identified from the entire proteome of SARS CoV-2 by a novel reverse epitomics approach. We conclude that the novel Multi-Patch Vaccines could be a highly potential novel approach to combat SARS-CoV-2, with greater effectiveness, high specificity and large human population coverage worldwide.

ABSTRACT FIGURE:

A Multi-Patch Vaccine design to combat SARS-CoV-2 and a method to prepare thereof.
Multi-Patch Vaccine designing to combat SARS-CoV-2 infection by reverse epitomics approach, “Overlapping-epitope-clusters-to-patches” method.

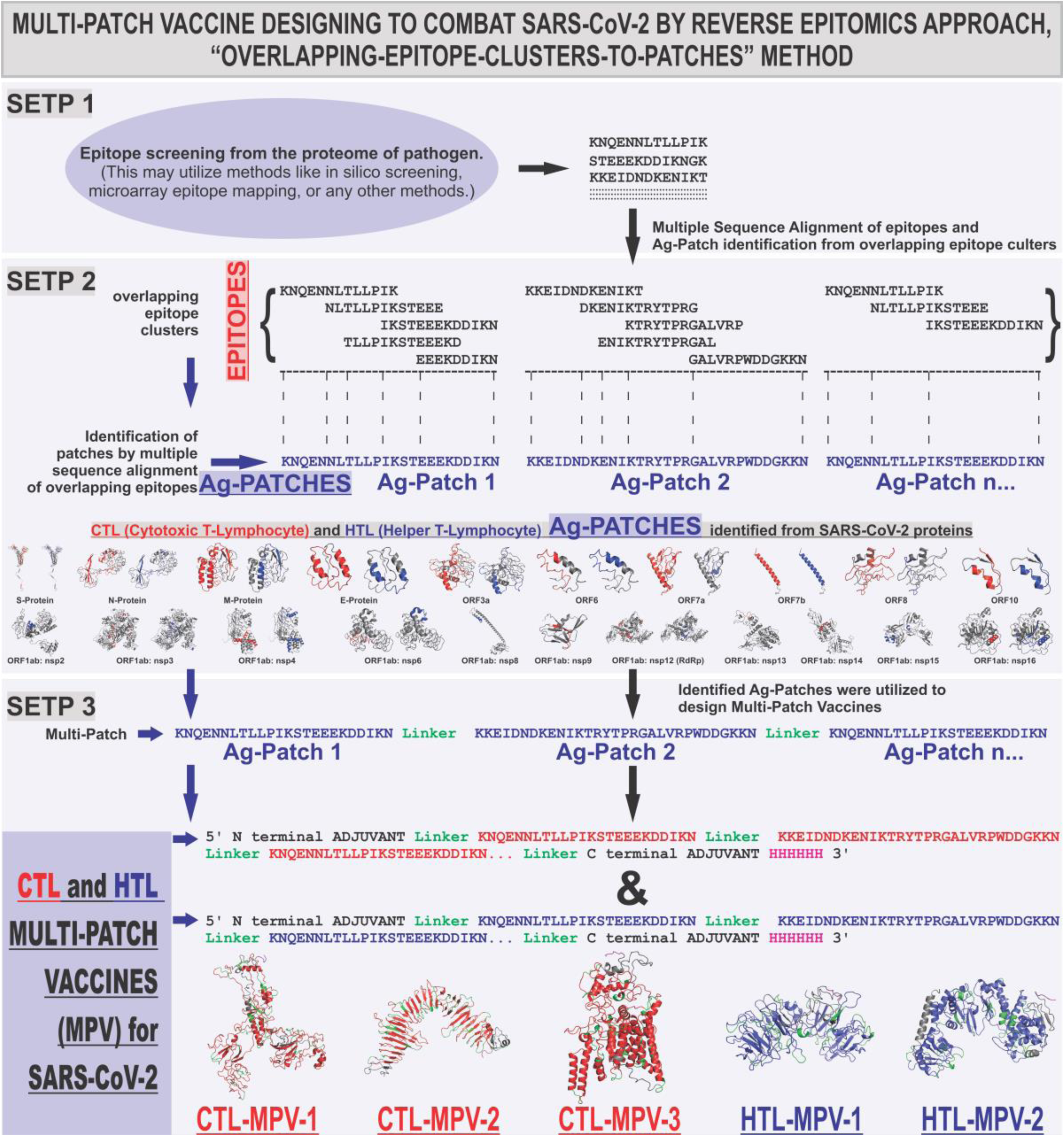

## Introduction

The Severe Acute Respiratory Syndrome Coronavirus 2 (SARS-CoV-2) causes the highly communicable disease COVID-19, the ongoing 2019-2020 disease outbreak which has turned into a pandemic. The disease has already spread to more than 216 countries and territories worldwide. More than 26,415,380 people have already got infected and 870,286 have died worldwide, resulting the ongoing mortality rate to be ∼3.295% (cases/deaths*100) (WHO Coronavirus Disease (COVID-19) Dashboard, 5^th^ September, 2020; WHO Weekly Epidemiological Update, 31^st^ August 2020). The outbreak has raised an urgent need of a specific and efficient vaccine against the SARS-CoV-2 virus.

The long term adaptive immunity crucially involves presentation of antigen as epitope on the surface of Antigen Presenting Cells (APC). The chop down processing of antigen by proteasome and lysosome into small size peptide epitopes of different length like (7,9,13,15… amino acid) pave the way for epitope formation and eventual presentation. The ‘Transporter associated with antigen processing’ (TAP) and further the HLA allele (human leukocyte antigen) molecules facilitate the epitope presentation (Figure 1A). The crucial step in the process of antigen presentation is the cleavage of the antigenic protein molecules to provide small length peptides which acts as epitopes at later stage after their presentation on the surface of APC (Melief et al., 2005; Kahan et al., 2003, Tenzer et al., 2005, Antoniou et al., 2003).

**Figure 1.**
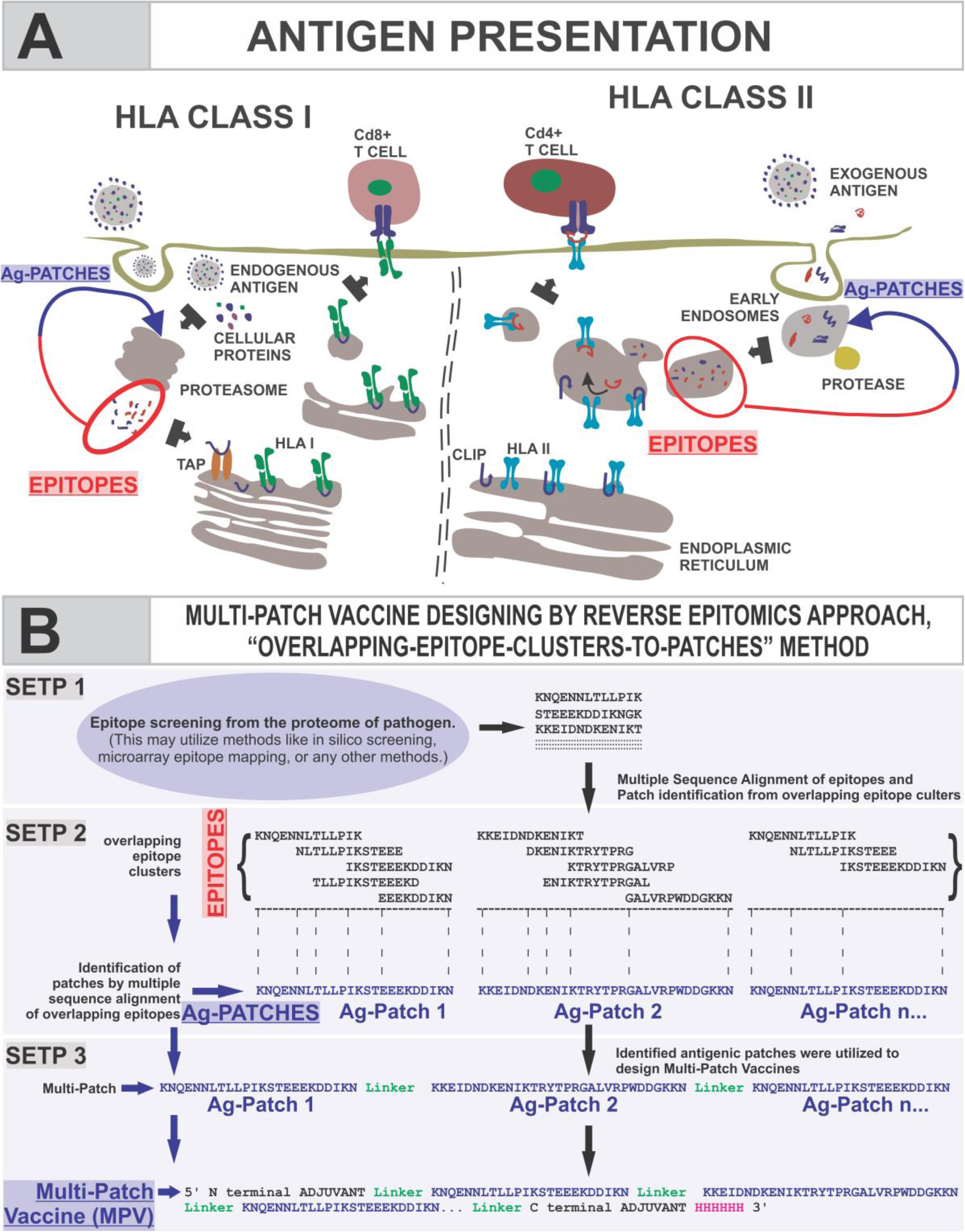
Schematic presentation for revers epitomics approach, “overlapping-epitope-clusters-to-patches” method. **A.** Schematic presentation of the process of epitope presentation by the Antigen Presenting Cells facilitated by TAP transporter and HLA class I, & II alleles. **B.** Presentation of “overlapping-epitope-clusters-to-patches” method (Step 1 & 2) to identify antigenic patches (Ag-Patches) from SARS-CoV-2/pathogen. The cluster of overlapping epitopes leads us to identify antigenic patches from the source protein, a reverse epitomics approach. The identified antigenic patches could be used to design Multi-Patch Vaccine as shown in Step. 3.

In the present study, we have performed the reverse epitomics analysis to identify antigenic patches from SARS-CoV-2 proteins by novel “overlapping-epitope-clusters-to-patches” method. The term “reverse epitomics” would involve search for antigenic patches from the overlapping multiple epitope clusters. This method of identifying antigenic patches (Ag-Patches) from a protein is termed here as “Overlapping-epitope-clusters-to-patches” method. These antigenic patches are termed here as Ag-Patch or Ag-Patches. These Ag-Patches, once administered as vaccine candidates, are expected to be chopped down to produce cluster of overlapping multiple epitope. In our present study we have identified 73 CTL (Cytotoxic T-Lymphocyte) and 49 HTL (Helper T-Lymphocyte) such Ag-Patches from the structural and non-structural proteins of SARS-CoV-2 virus proteome. Further we utilized these identified Ag-Patches from SARS-CoV-2 proteome to design Multi-Patch Vaccine candidates against SARS-CoV-2.

The designed MPVs are supposed to be more effective than both the subunit or single antigenic protein based vaccines and the Multi-Epitope Vaccines (MEVs). MPVs would involve Ag-Patches from entire proteome of SARS-CoV-2 and hence would be more effective than subunit or single protein vaccine. In comparison to the epitope based MEVs, the Ag-Patches based MPVs will provide a more natural antigenic vaccine candidate. The proteasome and lysosomal chop down processing of MPVs by the Antigen Presenting cells would release larger number of epitopes in their intact form for which the MEVs would have a challenge. Moreover the Ag-Patch based MPVs will cover larger number of overlapping epitopes in comparison to the limited number of epitopes covered by epitope based MEVs (Srivastava et al., 2020a; Srivastava et al., 2020b). The larger the number of epitopes covered larger would be the number of HLA alleles targeted and hence larger human population with ethnic distribution of HLA alleles will be covered. Hence, the MPVs would provide more effective and specific solution with larger human population coverage for vaccine development against SARS-CoV-2. The schematic of antigen presentation, the identification of antigenic patches (Ag-Patches) by the reverse epitomics “Overlapping-epitope-clusters-to-patches” method and Multi-Patch Vaccine designing is shown in Figure 1A and 1B.

## Methodology

In the present study, we have identified antigenic patches from all the eleven ORF proteins of SARS-CoV-2 proteome (Supplementary figure S1) (NCBI: SARS-CoV-2 isolate Wuhan-Hu-1, complete genome; gi|1798174254|ref|NC_045512.2|). These Ag-Patches are identified by the overlapping epitope clusters. These identified Ag-Patches are further utilized to design a Multi-Patch Vaccine (MPV) candidate against the SARS-CoV-2 infection. To identify the Ag-Patches from the SARS-CoV-2 proteins we have first screened high scoring CTL and HTL epitopes from all the eleven ORF proteins of the virus. The screening of potential CTL and HTL epitopes was done using various IEDB tools. We further analyzed the immunogenic properties of the screened CTL and HTL epitopes on the basis of their amino acid sequences and epitope databases.

We further performed multiple sequence alignment for all the screened epitopes leading us to obtain clusters of overlapping epitopes from the proteins of SARS-CoV-2. These CTL and HTL epitope clusters further led us to identify the antigenic patches (Ag-Patches) of the SARS-CoV-2 proteins from where the overlapping epitopes clusters were arising. The identified Ag-Patches, in other words, act as the source of multiple CTL and HTL epitopes and hence are the potential stretch of SARS-CoV-2 proteins which can be utilized to elicit T Cell immune response. The pairs of overlapping epitope forming clusters and their HLA allele binders were also utilized to analyze the world population coverage by the screened epitopes on the basis of HLA allele distributed worldwide amongst different ethnic human population.

We further used the identified antigenic patches to design chimeric fusion Multi-Patch Vaccines (MPVs) against SARS-CoV-2, by the GGGGS linker which is flexible in nature and favors peptide chain folding. We also liked human β defensin 2 and 3 (hBD-2 and hBD-3) at its N and C terminal of the Multi-Patch Vaccines candidates respectively as adjuvants, by the rigid linker EAAAK favoring domain formation (Hu et al., 2004; Chen et al., 2013; Hajighahramani et al., 2017; Srivastava et al., 2018; Srivastava et al., 2019). COVID-19 patients usually have lung infection and lung infection is usually coupled with increased expression of hBD-2 and hBD-3. Also, the β-Defensins are involved in the chemotactic activity for memory T cells, monocytes, immature dendritic cells as well as in degranulation of mast cells. The hBDs also enhances the innate and adaptive immunity and therefore are chosen here as adjuvants in the design of the MPVs (Wilson et al., 2013; Duits et al., 2003; Hoover et al., 2000; Yang et al., 2020; Biragyn et al., 2001; Kohlgraf et al., 2010). Further, discontinuous B cell epitopes, linear B cell epitopes and IFN-γ inducing epitopes were also screened from the generated tertiary models of all the MPVs, designed by the identified Ag-Patches of SARS-CoV-2.

The molecular signaling by multiple TLRs is an essential component of the innate immune response against viral infection. Upon lung infection the PBEC (peripheral blood mononuclear cells from infected patients) were found to express TLR3, which is involved in recognition of RNA. Hence we tested all the CTL and HTL MPV models for their molecular interaction with the ectodomain of the TLR3 receptor by molecular docking and molecular dynamics simulation studies (Delneste et al., 2007; Totura et al., 2015; Farina et al., 2005). Further, the codon-optimized cDNA of all the CTL and HTL MPVs were also analyzed and found to favor high expression in human cell line. The Schematic work flow chart and the methodology utilized in the present study are shown in Supplementary figure S2.

### Screening of potential epitopes from SARS-CoV-2 proteome

#### Screening of Cytotoxic T lymphocyte (CTL) Epitope

The screening of Cytotoxic T lymphocyte epitopes was performed by the IEDB (Immune Epitope Database) tools “MHC-I Binding Predictions” (http://tools.iedb.org/mhci/) and “MHC-I Processing Predictions” (Tenzer et al., 2005; Peters et al., 2003; Hoof et al., 2009). The tools use six different methods (viz. Consensus, NN-align, SMM-align, Combinatorial library, Sturniolo and NetMHCIIpan) and generate “Percentile rank” and a “total score” respectively indicating the immunogenic potential of the screened epitopes. The screening methods utilized are based on the total amount of cleavage sites in the submitted protein sequence. The TAP score of a peptide estimates an effective −log of (IC50) values (half maximal inhibitory concentration) for binding to TAP (Transporter associate with Antigen Processing) or its N-terminal prolonged precursors. The MHC (major histocompatibility complex) binding prediction score of a peptide is the –log of (IC50) values for binding to MHC molecule (Calis et al., 2013). The IC(50) (nM) value for each epitope and MHC allele binding pairs were also obtained by the IEDB tool. Epitopes having high, intermediate, and least affinity of binding to their respective HLA allele binders have IC50 values of < 50 nM, < 500 nM and < 5000 nM, respectively.

Immunogenicity of all the screened CTL epitopes was also obtained by using “MHC I Immunogenicity” tool of IEDB with all the parameters set to default analyzing 1st, 2nd, and C-terminus amino acids of the given epitope (Calis et al., 2013). The tool predicts the immunogenicity of a given peptide and MHC (pMHC) complex based on the physicochemical properties of constituting amino acid and their position within the peptide sequence.

#### Screening of Helper T lymphocyte (HTL) Epitopes

To screen out the Helper T lymphocyte epitopes from SARS-CoV-2 proteins, the IEDB tool “MHC-II Binding Predictions” was used. The tool generates “Percentile Rank” for each potential screened peptide. The lower the value of percentile, the higher would be the immunogenic potential of the peptide. This percentile score is generated by the combination of three different methods which includes combinatorial library, SMM_align and Sturniolo method. The generated percentile score is further compared with the other random five million 15-mer peptides of SWISSPROT database (Wang et al., 2010; Sidney et al., 2008; Nielsen et al., 2007; Sturniolo et al., 1999). The percentile rank of the screened peptides is generated from the consensus of all above mentioned methods by the median percentile rank.

#### CTL and HTL epitope Toxicity prediction

The tool ToxinPred was used to analyze the toxicity of screened CTL and HTL epitopes. The tool facilitates to identify the highly toxic or non-toxic short peptides. The toxicity check analysis was done by the “SVM (Swiss-Prot) based” (support vector machine) method utilizing a dataset of 1805 sequences as positive (toxic) and 3593 sequences as negative (non-toxic) from Swissprot along with a comparison to an alternative dataset comprising of the same 1805 positive sequences and 12541 other negative sequences from TrEMBLE (Gupta et al., 2013).

### Overlapping CTL and HTL epitope clusters to Ag-Patches

#### CTL & HTL multi-epitope cluster based Ag-Patch identification

To identify the potentially immunogenic Ag-Patches from SARS-CoV-2 proteome, all the screened high scoring epitopes were aligned against their respective source protein sequence by multiple sequence alignment (MSA) tool Clustal Omega available on the EBI server (Sievers et al., 2011). The Patches of the SARS-CoV-2 protein sequences showing consensus with the clusters of overlapping epitopes were chosen and shortlisted as antigenic patches (Ag-Patches). This approach of search and identification of antigenic patches from source protein in a reverse epitomics manner, i.e. from epitopes to antigenic patches of source protein, is here defined as “Overlapping-epitope-clusters-to-patches” method (Figure 1A, 1B). The here provided reverse epitomics approach, “Overlapping-epitope-clusters-to-patches” method to identify Antigenic Patches (Ag-Patches) from pathogen’s (here SARS-CoV-2) protein is included in filed Patent No: 202011037585.

#### Population Coverage by the overlapping CTL and HTL epitopes

The “Population Coverage” tool of IEDB was used to analyze the world human population coverage for both the CTL & HTL overlapping epitopes and their respective HLA allele binding pairs (Bui et al., 2006). The T cells recognize the complex of a specific MHC molecule with a particular pathogen-derived epitope. The given epitope will elicit an immune response only in an individual that expresses the epitope binding the MHC molecule. This denomination of the MHC restricted T cell responses and the MHC polymorphism amongst human population provides the basis for population coverage analysis. The MHC types are expressed at dramatically different frequencies in the different ethnicities of human population worldwide. In this way, the population coverage by an epitope-MHC pair could be determined (Sturniolo et al., 1999).

#### Conservation analysis of antigenic patches

The shortlisted CTL and HTL epitope cluster based antigenic (Ag-Patches) identified from eleven SARS-CoV-2 ORF proteins were analyzed for their amino acid sequence conservancy by “Epitope Conservancy Analysis” tool of IEDB. The epitope conservancy is the percentage of SARS-CoV-2 ORF protein sequences (retrieved from NCBI) containing the particular epitope cluster based Ag-Patch sequence in 100% amino acid sequence match. The analysis was done for all the identified Ag-Patches against all the source ORF protein sequences of SARS-CoV-2 proteins retrieved from the NCBI protein database (Bui et al., 2007).

### Multi-Patch Vaccines

The identified antigenic patches (Ag-Patches) from the SARS-CoV-2 proteome were further utilized to design three CTL and two HTL Multi-Patch Vaccines as explained in the result section.

#### Physicochemical property analysis of designed MPVs

The physicochemical properties of the amino acid sequences of the designed three CTL and two HTL MPVs were analyzed by the ProtParam tool (Gasteiger et al., 2005). The ProtParam tool performs an empirical investigation for the given query amino acid sequence of a protein. ProtParam computes various physicochemical parameters including amino acid length, molecular weight, theoretical pI, expected half-life (in E.Coli, Yeast & Mammalian cell), Aliphatic index, Grand average of hydropathicity (GRAVY) and instability index score. The aliphatic index and grand average of hydropathicity (GRAVY) indicate the globular and hydrophilic nature of the protein. The instability index score indicates the stable nature of the protein molecule.

#### Interferon-gamma inducing epitope prediction from the MPVs

From the designed amino acid sequence of all the three CTL and two HTL MPVs, the potential interferon-gamma (IFN-γ) epitopes were screened by utilizing the “IFN epitope” tool with “Motif and SVM hybrid” method (MERCI: Motif-EmeRging and with Classes-Identification, and SVM: support vector machine). The tool predicts peptides from the given protein sequences having the potential to induce release of IFN-gamma from CD4+ T cells. This module generates the overlapping, IFN-gamma inducing peptides sequences from the given query sequence. The prediction is based on the IEDB database with 3705 IFN-gamma inducing and 6728 non-inducing MHC class II binding epitopes (Nagpal et al., 2015; Dhanda et al., 2013).

#### MPVs allergenicity and antigenicity prediction

All the designed CTL and HTL MPVs were further analyzed for allergenicity and antigenicity prediction by utilizing the AlgPred and the Vaxigen tools respectively (Saha et al., 2006; Doytchinova and Flower 2007). The AlgPred prediction is based on the similarity with already known allergens & non-allergens database, with any region of the submitted protein. For the screening of allergenicity, the Swiss-prot dataset consisting of 101725 non-allergens and 323 allergens is utilized. The VaxiJen tool utilizes an alignment-free approach, solely based on the physicochemical properties of the given query protein sequence, to predict antigenicity of a give protein sequence. For the prediction of antigenicity, the Bacterial, viral and the tumor protein datasets are utilized by VaxiJen to predict the whole protein antigenicity.

#### Tertiary structure modelling and refinement of MPVs

The tertiary structure of all the designed three CTL and two HTL MPVs were generated by homology modelling utilizing the I-TASSER modelling tool. The I-TASSER is a tool that utilizes the sequence-to-structure-to-function paradigm for protein structure prediction (Roy et al., 2010). The tool generates three-dimensional (3D) atomic models from multiple threading alignments and iterative structural assembly simulations for the submitted amino acid sequence of the protein. It works based on the structure templates identified by LOMETS, a meta-server, from the PDB library. The I-TASSER tool only uses the templates of the highest Z-score which is the difference between the raw and average scores in the unit of standard deviation with reference to the templet protein structure. For each target model, the I-TASSER simulations generate a large ensemble of the structural conformations, known as decoys. To select the final models, I-TASSER uses the SPICKER program to cluster all the decoys based on the pair-wise structure similarity and reports up to five top-scoring models. The Normalized Z-score >1 indicates a good alignment. The Cov score represents the coverage of the threading alignment and is equal to the number of aligned residues divided by the amino acid length of the query protein sequence. Ranking of templet proteins is based on TM-score of the structural alignment between the query structure model and the known templet protein structures. The RMSD (Root Mean Square Deviation) is the deviation between templet residues and query residues that are structurally aligned by TM-align.

The refinement of all the generated three CTL and two HTL MPV models were performed by ModRefiner and GalaxyRefine tools (Dong et al., 2011; Ko et al., 2012). The TM-score generated by ModRefiner indicates the structural similarity of the refined model with the original input protein structure. Closer the TM-Score to 1, higher would be the similarity of original and the refined model. The RMSD value of the refined model shows the conformational deviation from the initial input protein model.

The GalaxyRefine tool refines the input tertiary structure by repeated structure perturbation as well as by subsequent structural relaxation and molecular dynamics simulation. The tool GalaxyRefine generates reliable core structures from multiple templates and then re-builds the loops or termini by using an optimization-based refinement method (Wang et al., 2013; Shin et al., 2014). To avoid any breaks or gaps in the generated 3D model, the GalaxyRefine uses the triaxial loop closure method. The MolProbity score generated for a given refined model indicates the log-weighted combination score of the clash score, the percentage of Ramachandran not favored residues and the percentage of bad side-chain rotamers.

#### Validation of CTL and HTL MPVs refined models

Both the refined three CTL and two HTL MPV tertiary models were further validated by RAMPAGE analysis tool (Lovell et al., 2003; Ramakrishnan et al., 1965). The generated Ramachandran plots for all the MPV models show the sterically allowed and disallowed residues along with their dihedral psi (ψ) and phi (φ) angles.

#### Linear and Discontinuous B-cell epitope prediction from the MPVs

The IEDB tool, Ellipro (ElliPro: Antibody Epitope Prediction tool) available at IEDB server was used to screen the linear and the discontinuous B cell epitopes from all the CTL and HTL MPVs tertiary structure models. The ElliPro method screened B-Cell epitope from the given tertiary structure based on the location of residues in the proteins 3D structure. The farthest residue to be considered was limited to 6 Å. The residues lying outside of an ellipsoid covering 90% of the inner core residues of the protein score highest Protrusion Index (PI) of 0.9; and so on. The discontinuous epitopes predicted by the ElliPro tool are clustered based on the distance “R” in Å between two residues centers of mass lying outside of the largest possible ellipsoid of the protein tertiary structure. The larger the value of R, greater would be the distance between the residues (residue discontinuity) of the screened discontinuous epitopes (Kringelum et al., 2012; Ponomarenko et al., 2008).

#### Molecular interaction analysis of MPVs and immune receptor complexes

Molecular interaction analysis of all the three CTL and two HTL MPVs with Toll-Like receptor 3 (TLR3), was performed by molecular docking followed by a molecular dynamics simulation study. Protein-protein molecular docking was performed by the tool PatchDock (Schneidman-Duhovny et al., 2005). The PatchDock tool utilizes an algorithm for unbound (mimicking real-world environment) docking of molecules for protein-protein complex formation (Duhovny et al., 2002). The algorithm carries out the rigid protein-protein docking with the surface variability/flexibility implicitly addressed through liberal intermolecular penetrations. The algorithm focuses on the (a) initial molecular surface fitting on localized and curvature-based patches of the protein surface (b) use of Geometric Hashing and Pose Clustering for initial transformation detection (c) computation of shape complementarity utilizing the Distance Transformation (d) efficient steric clash detection and geometric fit score based on a multi-resolution shape representation and (d) utilization of the biological information by focusing on the hot-spot rich surface patches (Duhovny et al., 2002; Schneidman-Duhovny et al., 2005). For molecular docking, the X-Ray crystal structure of human TLR3 ectodomain (ECD) was retrieved from PDB databank (PDB ID: 2A0Z) (Bell et al., 2005). The PatchDock molecular docking study provides dynamical properties of the designed system with the complex and also it gives ‘exact’ predictions of bulk properties including the hydrogen bond formation and conformation of the molecules forming the complex.

#### Molecular Dynamics (MD) Simulations study of MPVs and immune receptor complexes

The MPVs-TLR3 complex molecular interactions were further evaluated using Molecular Dynamics (MD) simulations analysis. MD simulation studies were performed for 10 nanoseconds (ns) by using the YASARA tool (Yet Another Scientific Artificial Reality Application) (Krieger and Vriend et al., 2015). The MD simulations studies were carried out in an explicit water environment in a dodecahedron simulation box at a stabilized temperature of 298K, the pressure of 1 atm and pH 7.4, with periodic cell boundary condition. The solvated systems were neutralized with counter ions (NaCl) (concentration 0.9 M). The AMBER14 force field was used on the systems during MD simulation (Maier et al., 2015; Case et al., 2014). The Long-range electrostatic energy and forces were calculated using particle-mesh-based Ewald method (Toukmaji et al., 2000). The solvated structures were energy minimized by the steepest descent method at a temperature of 298K and a stable pressure of 1 atm. Further, the complexes were equilibrated for period of 1 ns. After equilibration, a production MD simulation was run for 10 ns at a stable temperature and pressure and time frames were saved at every 10 ps, for each MD simulations. The RMSD and RMSF values for Cα, Backbone and all the atoms of all the CTL and HTL MPVs in complex with TLR3 were analyzed for each MD simulation study.

#### Analysis of MPVs cDNA for expression in human host cell line

Codon-optimized complementary DNA (cDNA) of all the three CTL and two HTL MPVs were generated for favored expression in Mammalian cell line (Human) by Java Codon Adaptation Tool. The generated cDNA of all the MPVs were further analyzed by GenScript Rare Codon Analysis Tool for its large scale expression potential. The tool analyses the GC content, Codon Adaptation Index (CAI) and the Tandem rare codon frequency in a given query cDNA sequence (Morla et al., 2016; Wu et al., 2010). The CAI indicates the possibility of cDNA expression in the chosen expression system. The tandem rare codon frequency indicates the presence of low-frequency codons in the given query cDNA sequence.

## RESULTS

### Screening of potential epitopes from SARS-CoV-2 proteome

#### Screening of Cytotoxic T lymphocyte (CTL) Epitope

Cytotoxic T lymphocyte (CTL) epitopes were screened by “MHC-I Binding Predictions” and “MHC-I Processing Predictions” tools of IEDB. A total of 1013 CTL epitopes and HLA I alleles pairs were screened (Supplementary table S1, S2). The immunogenicity analysis of all the screened CTL epitopes was also performed by the tool “MHC I Immunogenicity” of IEDB. The higher immunogenicity score of all the screened CTL epitopes suggest the high immunogenic potential of the CTL epitopes (Supplementary table S1, S2; Srivastava et al. 2020a; Srivastava et al. 2020b; and Grifoni et al., 2020a).

#### Screening of Helper T lymphocyte (HTL) Epitopes

The screening of helper T lymphocyte (HTL) epitopes from eleven different ORF proteins of SARS-CoV-2 was performed by the “MHC-II Binding Predictions” tool of IEDB. The tool generates percentile rank for all the screened epitopes. The smaller the value of percentile indicates higher “Percentile Rank” of the epitope. Furthermore, the higher Percentile rank suggests a higher affinity of the peptide to bind its respective HLA allele binders. A total of 314 HTL epitopes-HLA II allele pairs with high percentile rank were screened from the entire proteome of the SARS-CoV-2 virus (Supplementary table S3; Srivastava et al. 2020a; Srivastava et al. 2020b; and Grifoni et al., 2020a).

#### CTL and HTL epitope Toxicity prediction

Toxicity analysis of all the screened CTL and HTL epitopes was also performed by the tool “ToxinPred”. The ToxinPred study of all the screened CTL and HTL epitopes revealed that all the screened epitopes were Non-Toxic (Supplementary table S1, S2, S3).

### Overlapping CTL and HTL epitope clusters to Ag-Patches

#### CTL & HTL multi-epitope cluster based Ag-Patch identification

A total of 73 Ag-Patches from CTL and a total of 49 Ag-Patches from HTL high scoring 768 overlapping epitopes (518 CTL and 250 HTL epitopes) were identified (Table 1 & 2, Figure 2, 3 & 4) by the “Overlapping-epitope-clusters-to-patches” method (Figure 1A, 1B, table S1, S2, S3). The Ag-Patches hence obtained from the CTL and HTL epitope clusters are expected to produce back clusters of upto 768 overlapping epitopes, targeting different HLA allele, while their chop down processing by proteasome and lysosome, in the process of antigen presentation by the Antigen Presenting Cells (APC). This would be crucial step for the Multi-Patch Vaccines and shown strong possibility for the epitopes to be raised in their intact form and presented by APC. On other hand, such large numbers of epitopes are not possible to be accommodated by the Multi-Epitope Vaccines. The MEVs would also face challenge to give raise to the epitopes in their intact form upon chop down processing by proteasome or lysosome, for presentation by APC (Figure 1A, 1B, 2, 3 & 4). The identified and here provided SARS-CoV-2 Ag-Patches are included in filed Patent No: 202011037939.

**Figure 2.**
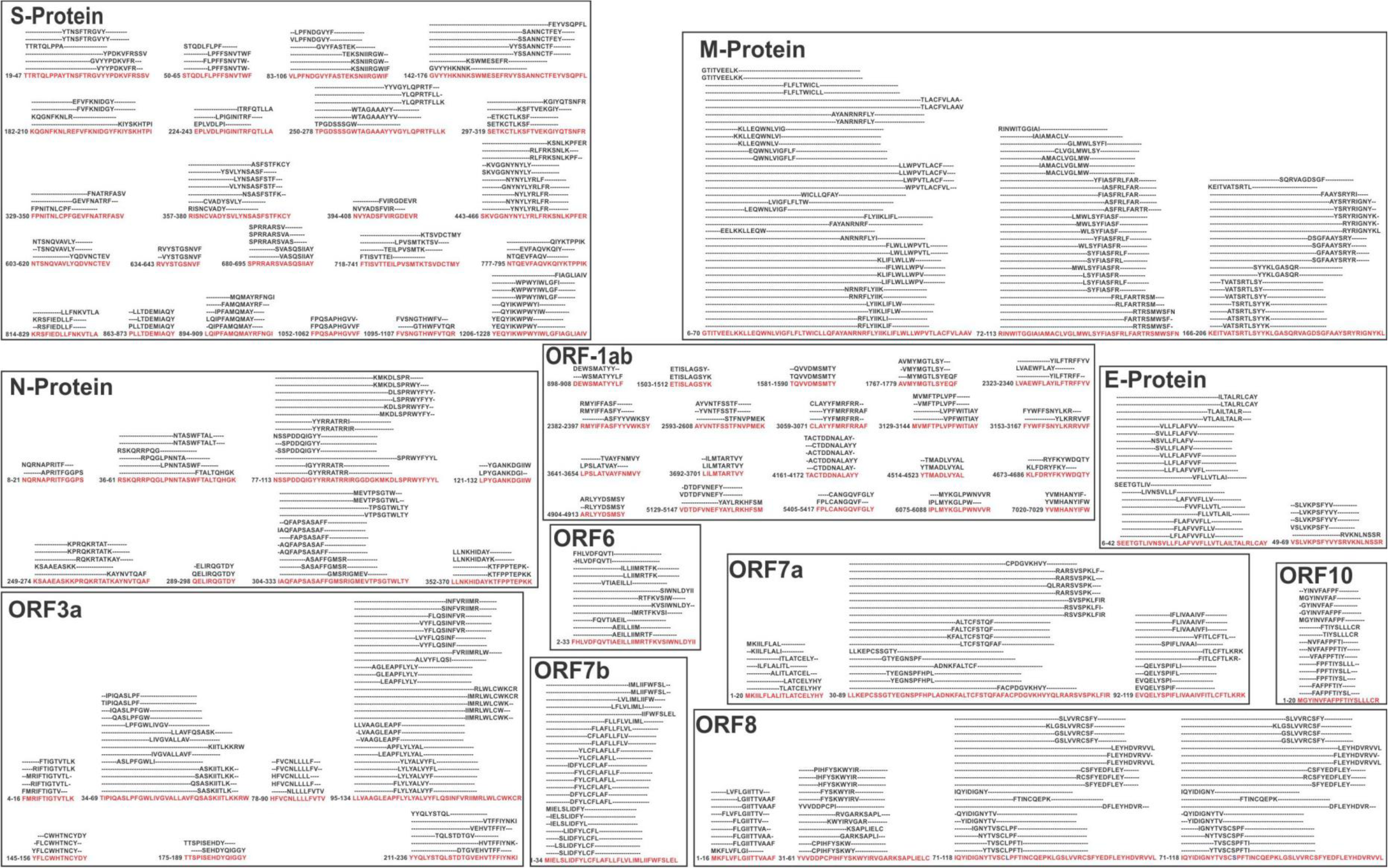
Immunogenic Ag-Patches (antigenic patches) identified from overlapping CTL epitope clusters of SARS-CoV-2 proteins. Immunogenic Ag-Patches are identified (red amino acid sequences) on the basis of the overlapping epitope clusters by the revers epitomics approach, “Overlapping-epitope-clusters-to patches” method.

**Figure 3.**
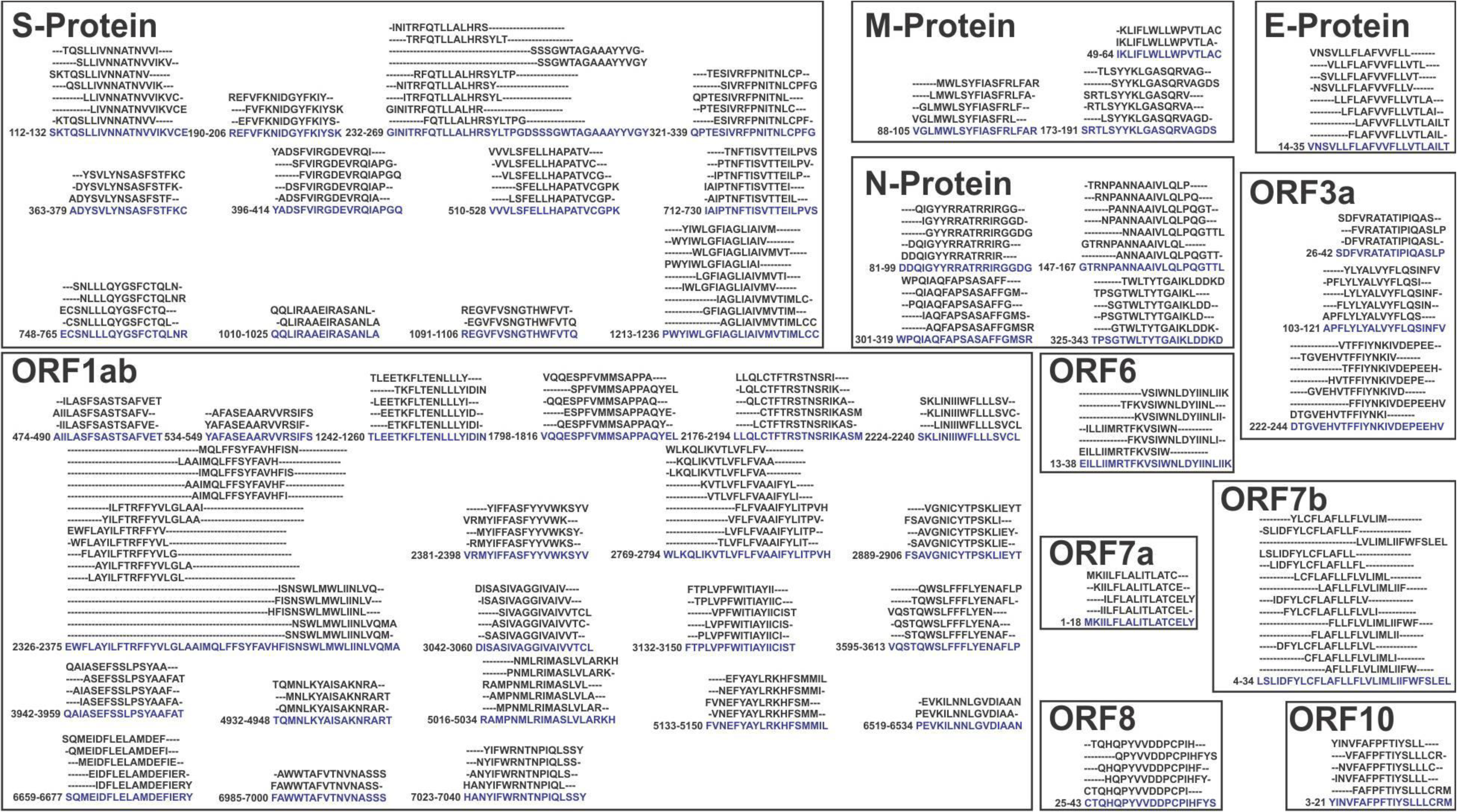
Immunogenic Ag-Patches (antigenic patches) identified from overlapping HTL epitope clusters of SARS-CoV-2 proteins. Immunogenic Ag-Patches identified (blue amino acid sequences) on the basis of the overlapping epitope clusters by the revers epitomics approach, “Overlapping-epitope-clusters-to patches” method.

**Table 1:**
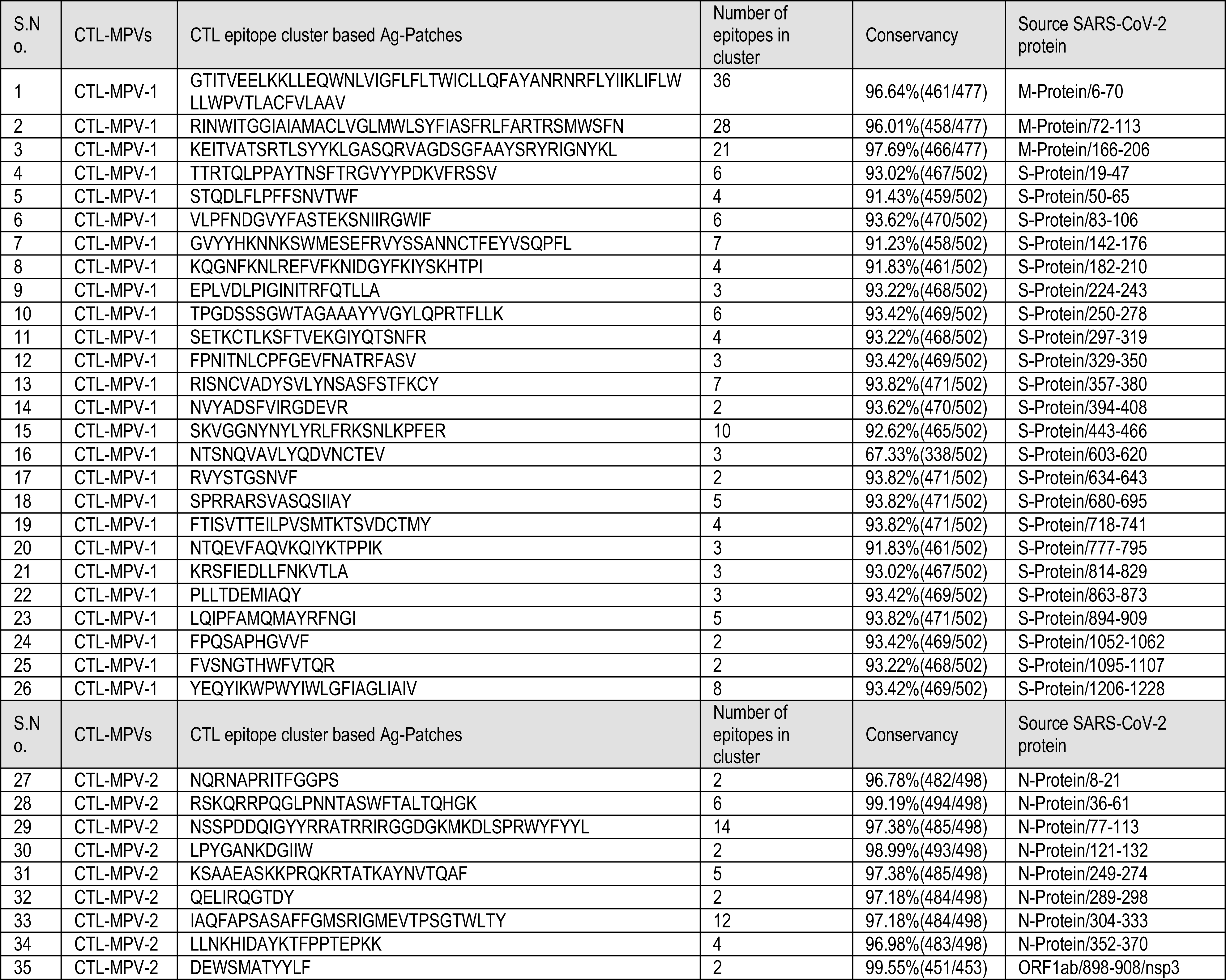
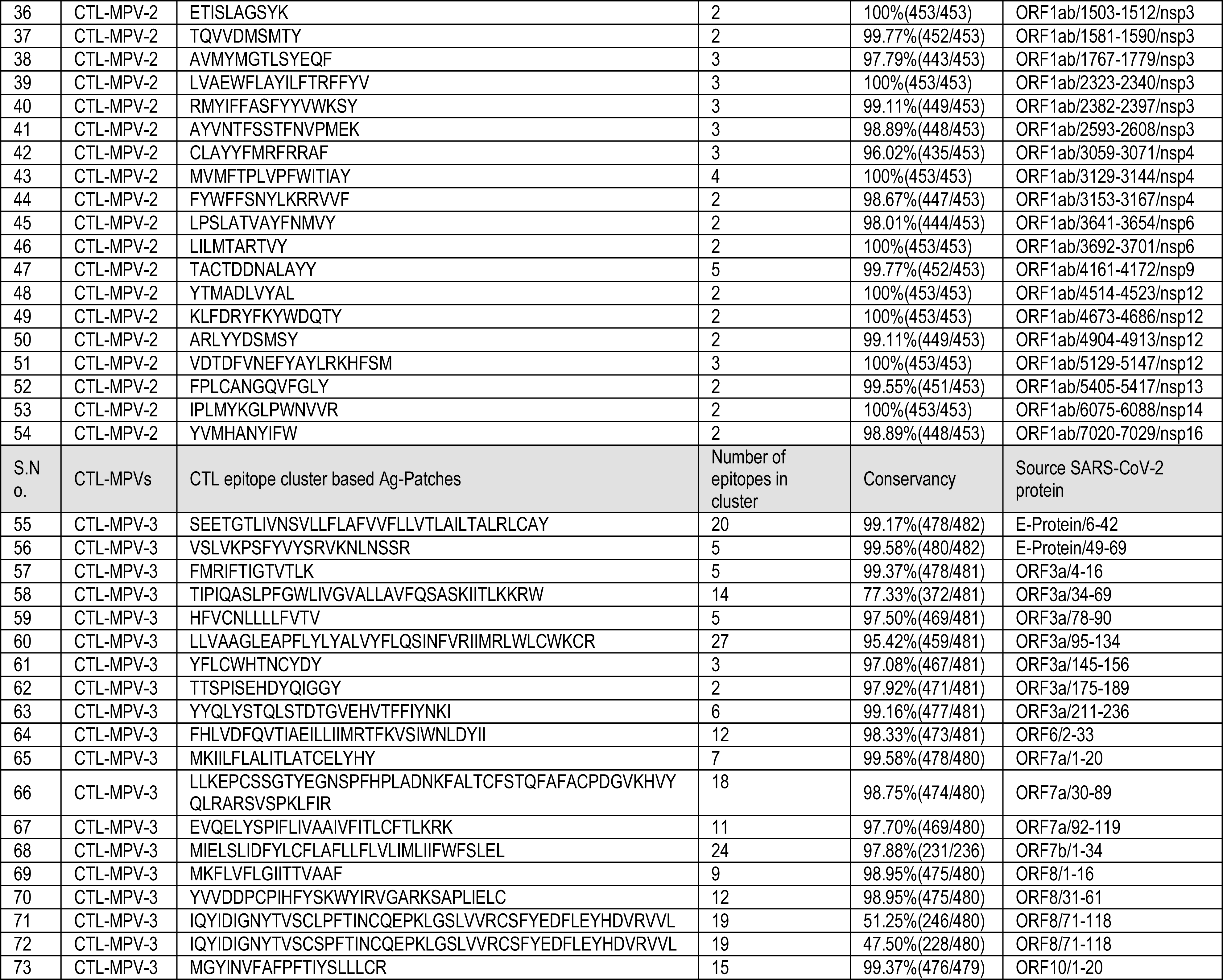
Identified “overlapping-epitope-clusters-to-patches” method based CTL Ag-Patches from the entire proteome of the SARS-CoV-2 virus. The identified highly immunogenic Ag-Patches were utilized to design three CTL (CTL-MPV-1, CTL-MPV-2 & CTL-MPV-3) Multi-Patch Vaccines. The identified Ag-Patches were highly conserved in nature and were identified on the basis of significant number of overlapping epitope forming clusters (Figure 2, 3 & 4).

**Table 2:**
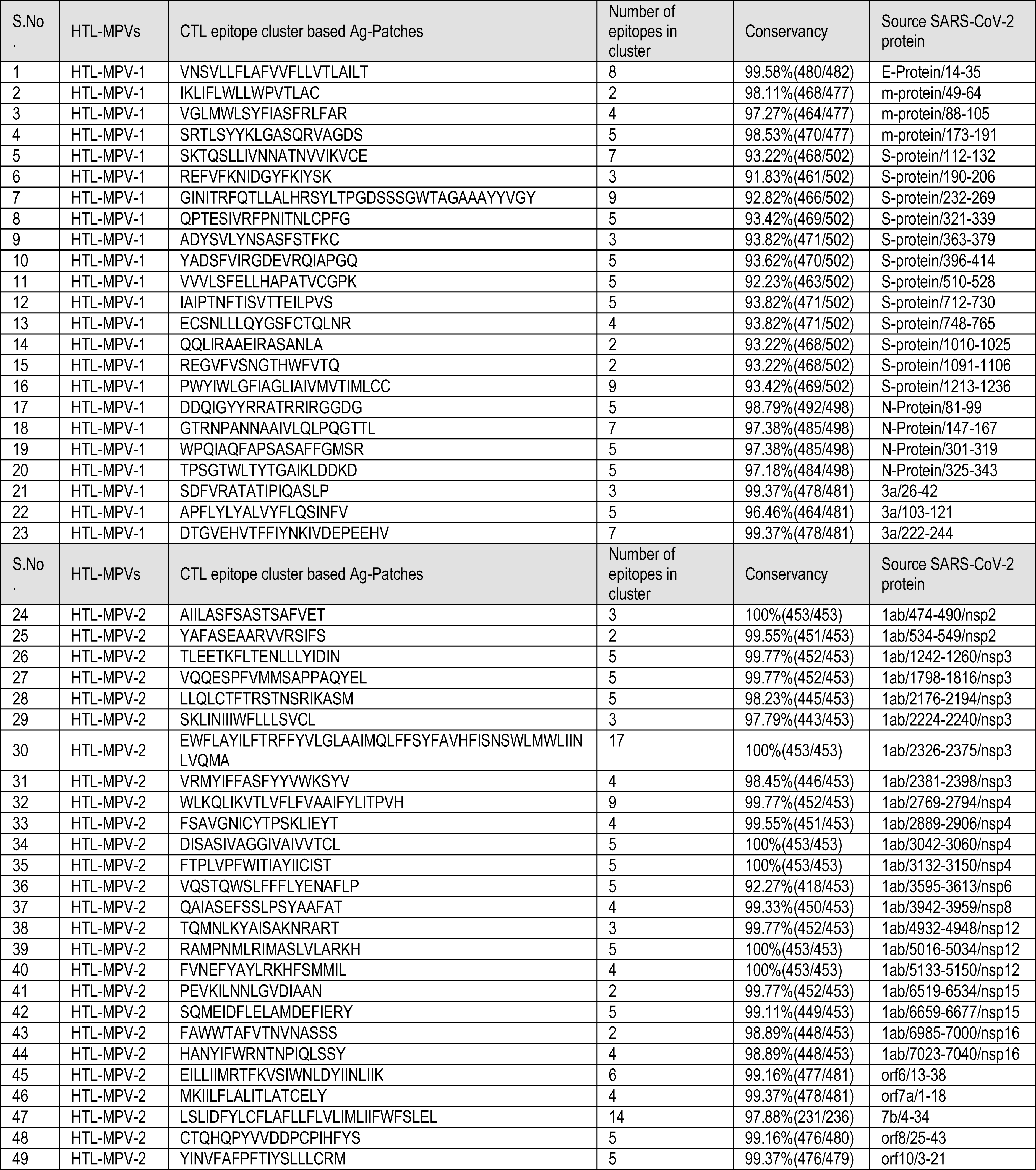
Identified “overlapping-epitope-clusters-to-patches” method based HTL Ag-Patches from the entire proteome of the SARS-CoV-2 virus. The identified highly immunogenic Ag-Patches were utilized to design two HTL (HTL-MPV-1 & HTL-MPV-2) Multi-Patch Vaccines. The identified Ag-Patches were highly conserved in nature and were identified on the basis of significant number of overlapping epitope forming clusters (Figure 2, 3 & 4).

#### Population Coverage by the overlapping CTL and HTL epitopes

The population coverage by all the overlapping CTL & HTL epitopes on the basis of their HLA allele binders was analyzed by the “Population Coverage” tool of IEDB. As all the Ag-Patches identified cover a large number of overlapping epitopes CTL and HTL epitopes, hence large number of HLA alleles are expected to be targeted and hence ethnically distributed large human population worldwide is expected to be covered. The population coverage analysis has shown very convincing results with 99.98% of world population to be covered with Average of 91.11 and standard deviation of 16.97. In our analysis all the human population ethnicity distribution worldwide was included. The countries most affected by COVID-19 have shown particularly significant coverage like France: 100%, Italy: 99.82%, United States: 100%, England: 100%, Germany 100%, Russia: 100%, Brazil: 99.99%; India: 99.94%, China: 99.83% etc. (Supplementary table S4). Epitopes-HLA allele pairs are given in Supplementary 2 file.

#### Conservation analysis of antigenic patches

The identified Ag-Patches were further analyzed for their amino acid sequence conservancy across the source protein sequence library retrieved from NCBI protein database, by the “Epitope Conservancy Analysis” tool of IEDB. We found that all the identified immunogenic Ag-Patches are highly conserved and both the CTL & HTL Ag-Patches are significantly conserved with their 100% amino acid residues largely conserved amongst the protein sequences of SARS-CoV-2 retrieved from NCBI Protein database (CTL epitopes cluster Ag-Patches were 47.50% to 100% conserved (largely above 91.23%) and HTL epitopes cluster Ag-Patches were 91.83% to 100% conserved (Table 1 & 2).

### Multi-Patch Vaccines

#### Design of Multi-Patch Vaccines

The identified overlapping epitope cluster based Ag-Patches from the proteome of the SARS-CoV-2 were utilized to design three CTL and two HTL Multi-Patch vaccines (Figure 4). Short peptide linkers EAAAK and GGGGS were utilized as rigid and flexible linkers respectively. The short peptide linker EAAAK facilitates the domain formation and provides a rigid link between two domains facilitating the protein to fold in a stable tertiary conformation. The short peptide linker GGGGS, with its flexible nature, provides conformational flexibility and hence facilitates stable conformation to the final folded protein structure. The rigid linker was utilized to fuse the human β defensin 2 and 3 (hBD-2 and hBD-3) at N and C terminal of the MPVs, respectively. The human β defensin 2 and 3 have been utilized here as an adjuvant to enhance immunogenic response. The GGGGS was utilized as a flexible linker to fuse the Ag-Patches together, and to facilitate folding of the protein into its tertiary conformation (Hu et al., 2004; Wilson et al., 2013; Duits et al., 2003; Chen et al., 2013; Hajighahramani et al., 2017; Srivastava et al., 2019; Srivastava et al., 2018; Srivastava et al., 2020a; Srivastava et al., 2020b) (Figure 1A, 1B, 2, 3, 4 & 5). The here provided MPV designs against SARS-CoV-2 and the designing method are included in the filed Patent No: 202011037939.

**Figure 4.**
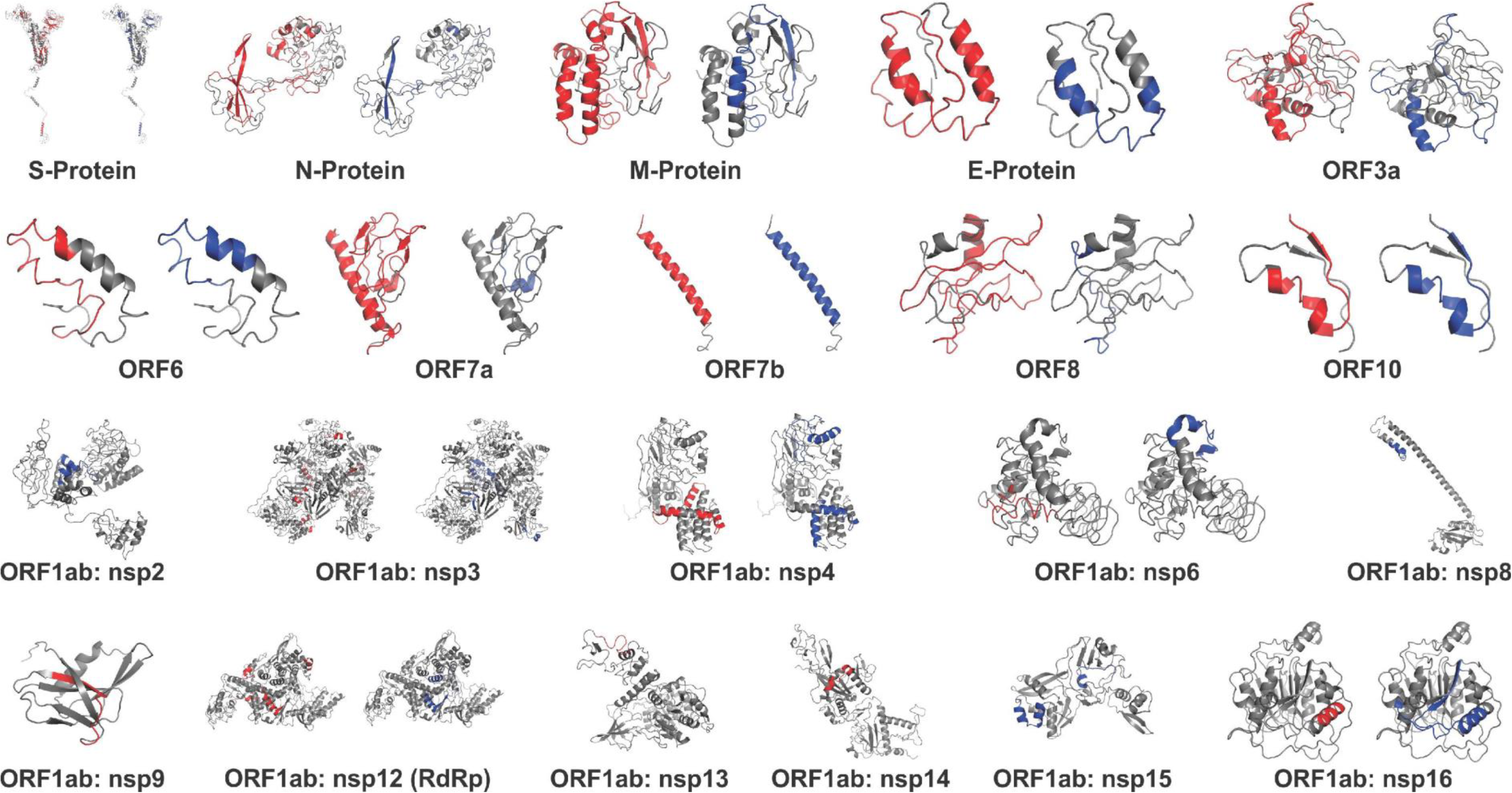
The identified immunogenic Ag-Patches (antigenic patches) shown in the tertiary structure models of the SARS-CoV-2 proteins. The tertiary structure models of all the SARS-CoV-2 proteins have been retrieved from I-TASSER homology modeling server (https://zhanglab.ccmb.med.umich.edu/COVID-19/). The CTL Ag-Patches are shown in red and the HTL Ag-Patches are shown in blue color. The identification of novel Ag-Patches has been done by the reverse epitomics approach, “overlapping-epitope-clusters-to-patches” method. Most of the Ag-Patches identified in the SARS-CoV-2 proteins are observed to be on the exposed surface of the proteins.

**Figure 5.**
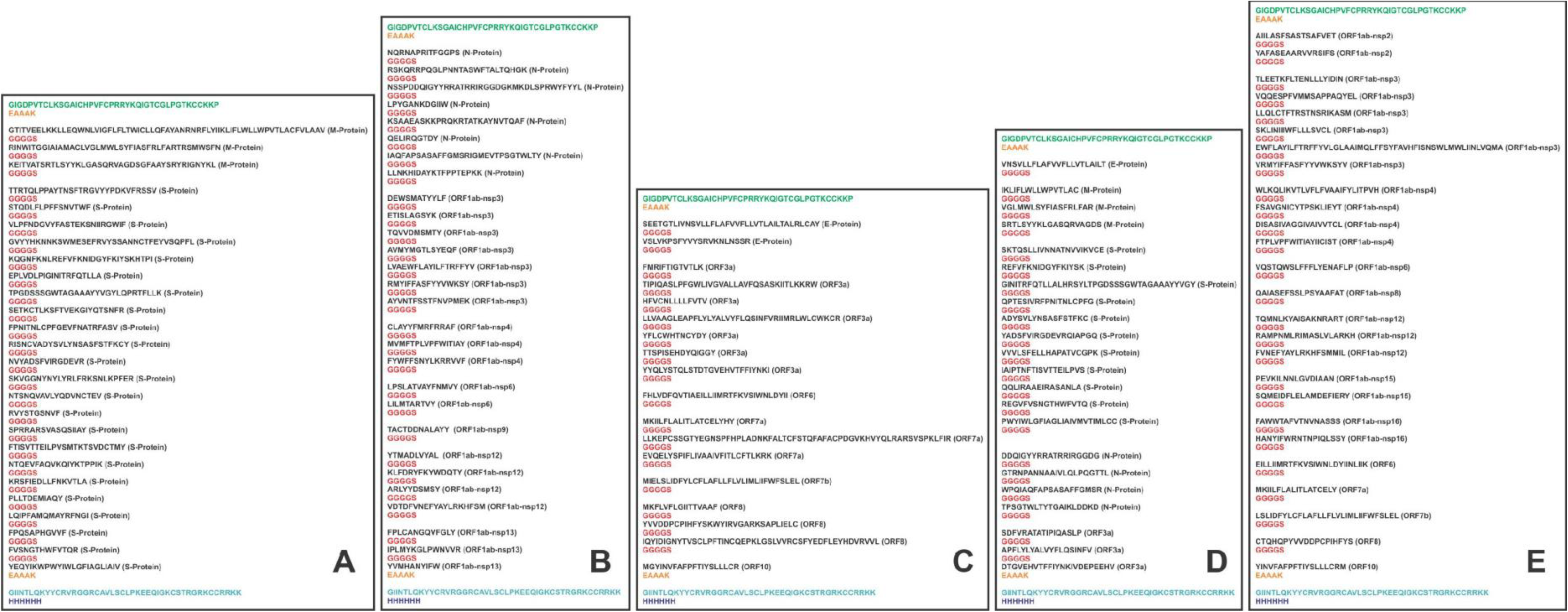
The CTL and HTL Ag-Patches were utilized to design Multi-Patch Vaccine. Short peptide linkers GGGGS and EAAK were used to fuse the Ag-Patches (antigenic patches) and the adjuvants, respectively. The CTL MPV constructs includes (A) CTL-MPV-1, (B) CTL-MPV-2, and (C) CTL-MPV-3. The HTL MPV construct includes (D) HTL-MPV-1 and (E) HTL-MPV-2 (Supplementary table S5).

#### Physicochemical property analysis of designed MPVs

ProtParam analysis for all the designed three CTL and two HTL Multi-Patch Vaccines (MPVs) was performed to analyze their physiochemical properties. The empirical physiochemical properties of the CTL and HTL MPVs are given in Table 3. The molecular weight of all the MPVs range from 66.36 to 89.96 kDa. The expected half-life of all the MPVs to ∼30 Hours in mammalian cells is very favorable for the vaccine protein expression and purification in vitro. The aliphatic index (53.51 to 100.86) and grand average of hydropathicity (GRAVY) (−0.274 to 0.445) of all the MPVs indicate their globular and hydrophilic nature. The instability index score of all the MPVs (39.68 to 53.37) indicates the stable nature of the protein molecules on expression in vitro. Overall, all the physiochemical parameters of all the MPVs suggest a favored expression of MPVs in vitro (Table 3).

**Table 3.**
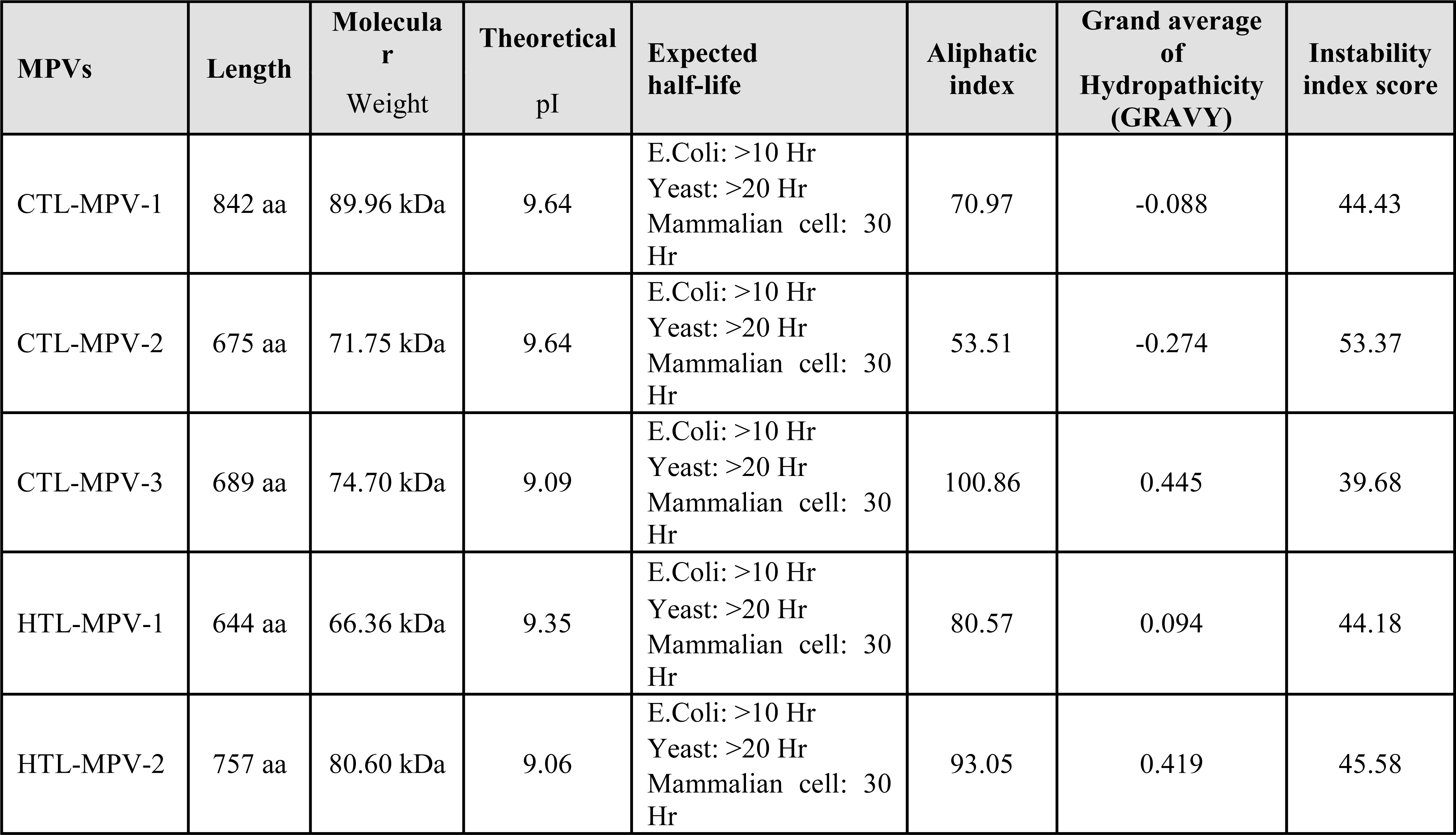
Physicochemical property analysis based on the amino acid sequences of all the designed three CTL and two HTL Multi-Patch Vaccines.

#### Interferon-gamma inducing epitope prediction from the MPVs

The Interferon-gamma (IFN-γ) inducing epitopes are involved in both innate as well as adaptive immune response. The IFN-γ inducing 15 mer peptide epitopes were screened from the amino acid sequence of all the three CTL and two HTL MPVs by utilizing the IFNepitope tool. A total of 257 IFN-γ inducing epitopes were screened from three CTL MPVs (108 from CTL-MPV-1; 66 from CTL-MPV-2 and 83 from CTL-MPV-3); likewise, a total of 181 IFN-γ inducing epitopes were screened from two HTL MPVs (83 from HTL-MPV-1 and 98 from HTL-MPV-2). In the screening, only the INF-γ inducing POSITIVE epitopes with a score of 1 or more than 1 were included (Supplementary table S6 & S7).

#### MPVs allergenicity and antigenicity prediction

We used the AlgPred and Vaxijen tools to analyse the allergic and antigenic nature respectively of all the MPVs on the basis of their amino acid composition. All the three CTL and two HTL MPVs were suggested as NON-ALLERGEN (scoring: −0.46602545 for CTL-MPV-1, −0.71579187 for CTL-MPV-2, −0.90796056 for CTL-MPV-3, −0.56197384 for HTL-MPV-1 and −0.72617142 for HTL-MPV-2; the threshold cutoff being −0.4). Likewise, all the three CTL and two HTL MPVs were found to be potential ANTIGENS as suggested by Vaxijen analysis (scoring: 0.5241 for CTL-MPV-1, 0.4811 for CTL-MPV-2, 0.6534 for CTL-MPV-3, 0.5016 for HTL-MPV-1 and 0.5096 for HTL-MPV-2; default threshold being 0.4). Overall the AlgPred and VaxiJen analysis of all the three CTL MPVs and two HTL MPVs suggests that all the MPVs are non-allergic in nature but they are antigenic. These both properties are essential and favorable for a potential vaccine candidate.

#### Tertiary structure modelling and refinement of MPVs

The tertiary structure models of all the designed three CTL and two HTL MPVs were generated by utilizing the I-TASSER modeling tool (Figure 6). The parameters (C-score, TM-Score and RMSD) of homology modeling have shown acceptable values for all the MPV models (Supplementary table S8). The C-score is a confidence score for estimating the quality of predicted models by I-TASSER. It indicates the significance of threading template alignments and the convergence parameters of the structure assembly simulations. C-score typically rages from −5 to 2, where higher value signifies a model with high confidence and vice-versa. The TM-score indicates the structural alignment between the query structure and the templet structures. The RMSD (Root Mean Square Deviation) is the deviation between the residues that are structurally aligned (by TM-align) to the templet structure.

**Figure 6.**
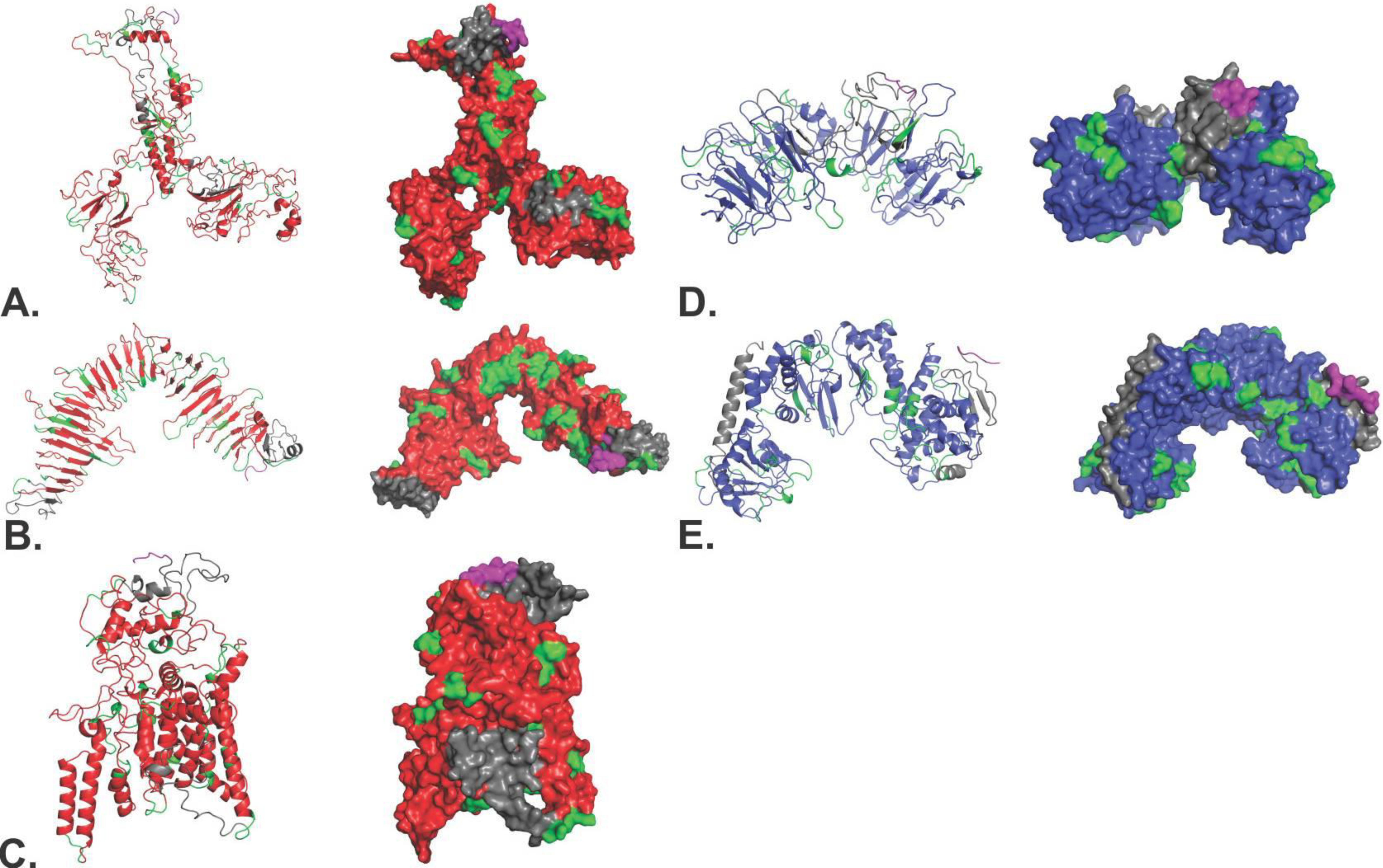
Tertiary structure modelling of CTL and HTL Multi-Patch Vaccines. Tertiary structural models of CTL and HTL MPVs have been shown in both cartoon and surface presentation (A) CTL-MPV-1, (B) CTL-MPV-2, (C) CTL-MPV-3, (D) HTL-MPV-1 and (E) HTL-MPV-2. The Ag-Patches are shown in Red (CTL Ag-Patches), and blue (HTL Ag-Patches). Linkers are in green, adjuvants are in gray and 6xHis tag is shown in magenta.

All the generated three CTL and two HTL MPV tertiary models were further refined using ModRefiner and GalaxyRefine refinement tools to fix any gaps and improve stoichiometric conformational parameters of the models. The refinement by ModRefiner shows the TM-score of 0.9729, 0.9324, 0.9849, 0.9847 and 0.9854 for CTL-MPV-1, CTL-MPV-2, CTL-MPV-3, HTL-MPV-1 and HTL-MPV-1 models respectively. The TM-score being close to 1 suggests that the initial and the refined models were structurally similar. After refinement, the RMSD for CTL-MPV-1, CTL-MPV-2, CTL-MPV-3, HTL-MPV-1 and HTL-MPV-1 models with respect to the initial model was 2.183 Å, 2.634 Å, 1.185 Å, 1.120 Å and 1.162 Å respectively, indicating not much deviation from the initial structure.

Further, all the three CTL and two HTL MPV models were refined by GalaxyRefine and the top model was chosen based on best scorings parameters. All the three CTL and two HTL MPV refinement output models have acceptable Rama favoured, GDT-HA, RMSD value and MolProbity scores (Supplementary table S9). The Rama favored is the percentage of residues which come in the favoured region of the Ramachandran plot, the GDT-HA (global distance test-High Accuracy) indicates the accuracy of the backbone structure, the RMSD (Root Mean Square Deviation) value indicated deviation from the initial model, and the MolProbity score indicates the log-weighted combination score of the clash score, the percentage of Ramachandran not favored residues and the percentage of bad side-chain rotamers.

Overall, after refinement, all the CTL and HTL MPV models had all their stoichiometric parameters in acceptable range and hence all these models were carried forward for further analysis (Supplementary table S8 & S9, Figure 6).

#### Validation of CTL and HTL MPVs refined models

All the three CTL and two HTL MPV models were further analyzed again by the RAMPAGE analysis tool after refinement to their stoichiometric acceptable conformation. All the three refined CTL MPV models (CTL-MPV-1, CTL-MPV-2 & CTL-MPV-3) were found to have 87.0%, 89.20%, & 88.8% favored region residues; 10.2%, 8%, & 7.6% allowed region residues, and 2.7%, 2.8%, & 3.6% outlier region residues, respectively. Likewise, all the two refined HTL MPV models (HTL-MPV-1 & HTL-MPV-2) were found to have 86.9% & 93.0% favored region residues, 9.3% & 5.9% allowed region residues, and only 3.7% & 1.1% outlier region residues respectively (Supplementary figure S3). Hence, all the three CTL and two HTL MPV models were found to have acceptable stoichiometric conformational parameters.

#### Linear and Discontinuous B-cell epitope prediction from the MPVs

The generated tertiary models of CTL and HTL MPVs were further screened for linear and discontinuous B-Cell epitopes. The B-Cell epitope screening was performed by the ElliPro tool available of IEDB. The screening revealed that the CTL MPV carries a total of 41 linear [CTL-MPV-1 (12 epitopes), CTL-MPV-2 (14 epitopes), and CTL-MPV-3 (15 epitopes)] and 19 discontinuous B-cell epitopes [CTL-MPV-1 (7 epitopes), CTL-MPV-2 (5 epitopes), and CTL-MPV-3 (7 epitopes)]. Likewise, the HTL MPVs carries a total of 30 linear [HTL-MPV-1 (16 epitopes) and HTL-MPV-2 (14 epitopes)] and 13 potential discontinuous epitopes [HTL-MPV-1 (8 epitopes) and HTL-MPV-2 (5 epitopes)]. The ElliPro associates each predicted epitope with a score, defined as a PI (Protrusion Index) value. The high PI score of the linear and discontinuous epitopes originating from the CTL and HTL MPVs suggest all the MPVs to have high potential to cause humoral immune response (Supplementary table S10, S11, S12, and S13).

#### Molecular interaction analysis of MPVs and immune receptor complexes

The immune receptor Toll-Like Receptor 3 (TLR3) acts as sentinel for our immune system. The molecular interaction of the designed models of CTL and HTL MPVs with the TLR3 receptor is crucial for the MPVs to act as a vaccine candidate. Hence all the CTL and HTL MPVs were further studied for their molecular interaction with the ectodomain (ECD) of human TLR3 receptor. Molecular docking study for the CTL and HTL MPVs models with the TLR3-ECD crystal structure model (PDB ID: 2A0Z) was performed by utilizing the PatchDock tool. The generated docking complex conformations with the highest docking score were chosen for further study [CTL-MPV-1:TLR3 (docking score: 17696), CTL-MPV-2:TLR3 (docking score: 17118), CTL-MPV-3:TLR3 (docking score: 16562), HTL-MPV-1:TLR3 (docking score: 21432) and HTL-MPV-2:TLR3 (docking score: 17620)]. The highest docking score indicates the MPV and TLR3-ECD complexes to have best geometric shape complementarity fitting conformation as predicted by the PatchDock tool. All the three CTL and two HTL MPVs were fitting into the ectodomain region of TLR3 receptor (Figure 7).

**Figure 7.**
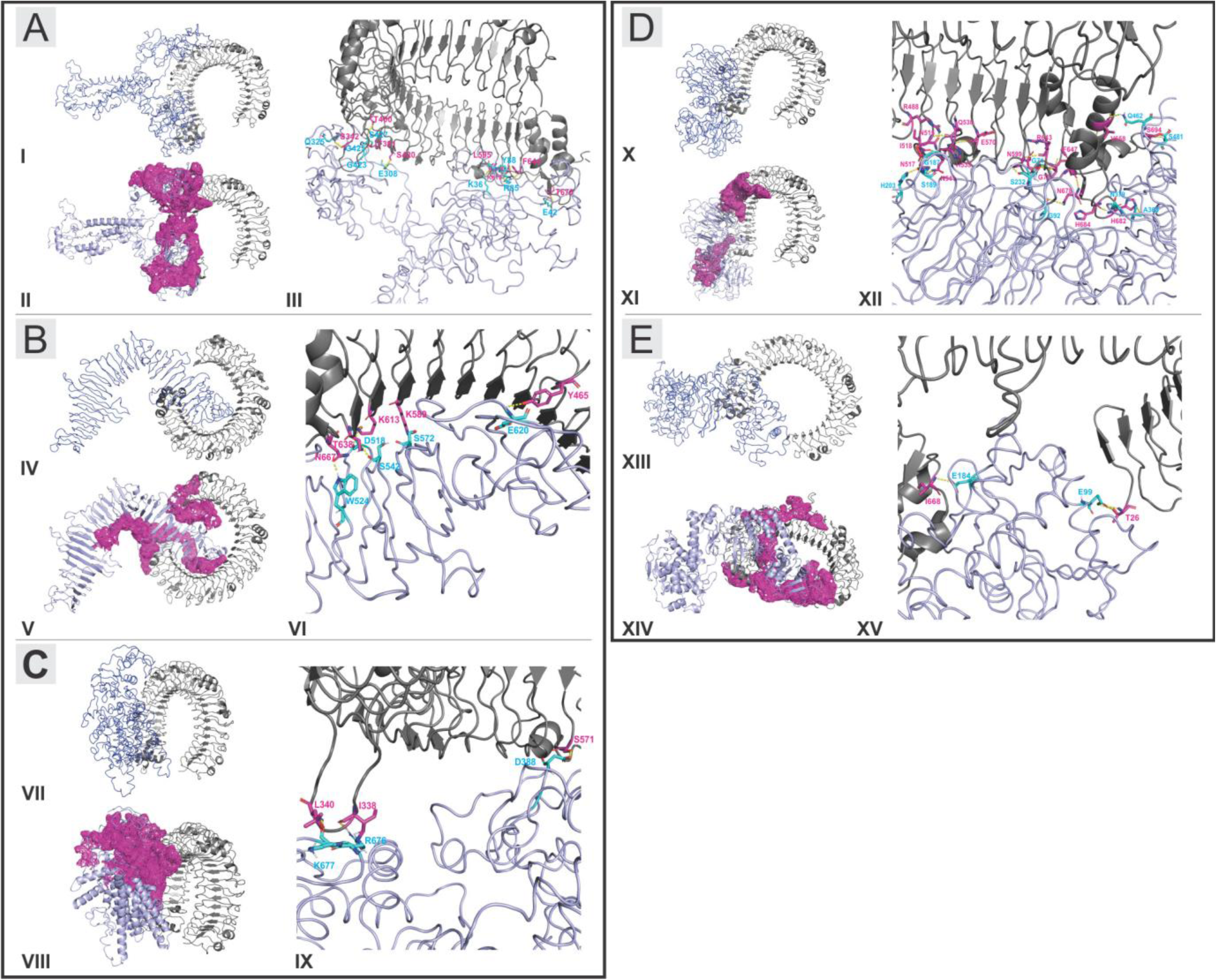
Molecular Docking Study of CTL and HTL MPVs with TLR3-ECD complexes. (A, B, C, D, E): Complex formation by the MPVs (CTL-MPV-1, CTL-MPV-2, CTL-MPV-3, HTL-MPV-1, HTL-MPV-2) and TLR3-ECD, respectively. (A-I, B-IV, C-VII, D-X, E-XIII): B-factor analysis presentation for the docked MPVs to the TLR3-ECD. The presentation is in VIBGYOR color, with blue showing low B-factor and red show high B-factor. Here most of the MPV regions are in blue showing low B-factor and hence suggesting a stable complex formation with TLR3-ECD. (A-II, B-V, C-VIII, D-XI, E-XIV): Binding site formed by the MPVs and TLR3-ECD at the molecular interaction interface, represented in magenta binding patches. All the CTL and HTL MPVs have shown to form prominent and close binding site region with TLR3-ECD. (A-III, B-VI, C-IX, D-XII, E-XV): Representation of hydrogen bond formation is shown. The residues involved in hydrogen bond formation from MPVs are shown in cyan, and the residues from TLR3-ECD are shown in magenta. The hydrogen bonds are shown by yellow dots.

The B-factor analysis of all the MPV & TLR3-ECD complexes was also performed. The B-factor of a tertiary structure indicates the displacement of the atomic positions from an average (mean) value of the structure i.e. the more flexible an atom is the larger the displacement from the mean position will be (mean-squares displacement) (Figure 7: A-I, B-IV, C-VII, D-X, E-XIII). The B-factor analysis of the CTL and HTL MPVs in complex with TLR3-ECD shows that most of the regions of MPVs bound to TLR3-ECD are stable nature in nature. The B-Factor analysis has been represented by the VIBGYOR color presentation with blue representing low B-factor and red representing high B-factor. Since the entire regions of the MPVs in complex with TLR3-ECD are in blue, hence suggest stable complex formation tendency for all the CTL and HTL MPVs with the ectodomain of the human TLR3 receptor.

All the CTL and HTL MPVs have shown to form prominent and close binding site region with TLR3-ECD as represented by binding patches shown in magenta (Figure 7: A-II, B-V, C-VIII, D-XI, E-XIV). The complex forms multiple hydrogen bonds in the interaction interface of MPVs and the ectodomain cavity region of TLR3 receptor (Figure 7: A-III, B-VI, C-IX, D-XII, E-XV). The residues involved in hydrogen bond formation for CTL-MPV-1:TLR3-ECD are GLN325:SER342, GLY423:THR391, GLY425:THR391, SER427:THR400, GLU308:SER420, LYS36:LEY595, GLU42:THR679, TYR88:LYS619, ARG85:LYS619, ARG85:PHE644. The residues involved in hydrogen bond formation for CTL-MPV-2:TLR3-ECD are TRP524:ASN667, SER542:THR638, ASP518:LYS613, SER572:LYS589, GLU620:TYR465. The residues involved in hydrogen bond formation for CTL-MPV-3:TLR3-ECD are LYS677:LEU340, ARG676:ILE338, ASP388:SER571. The residues involved in hydrogen bond formation for HTL-MPV-1:TLR3-ECD are HIS203:ASN517, SER189:ASN541, GLY187:HIS539, SER232:LYS619, GLY92:ASN678, GLY71:THR623, ASN379:HIS684, ASN379:HIS682, ALA363:HIS682, GLN462:VAL658, SER481:SER694. The residues involved in hydrogen bond formation for HTL-MPV-2:TLR3-ECD are GLU184:ILE668, and GLU99:THR26.

#### Molecular Dynamics (MD) Simulations study of MPVs and immune receptor complexes

All the three CTL and two HTL MPVs in complex with TLR3-ECD were further subjected to molecular dynamics (MD) simulation analysis to investigate the stability of the molecular interaction between the MPVs and the TLR3-ECD. All the complexes have shown a very convincing and reasonably stable Root Mean Square Deviation (RMSD) values for Cα, Backbone, and all-atom of the complexes which further stabilizes towards the end MD simulation [(CTL-MPV-1:TLR3-ECD, RMSD: ∼4 to ∼12 Å) (CTL-MPV-2:TLR3-ECD, RMSD: ∼4 to ∼9 Å) (CTL-MPV-3:TLR3-ECD, RMSD: ∼2.5 to ∼5.5 Å) (HTL-MPV-1:TLR3-ECD, RMSD: ∼3 to ∼6 Å) (HTL-MPV-2:TLR3-ECD, RMSD: ∼2.5 to ∼6 Å)] (Figure 8: A-E). The RMSD values for all the MPVs:TLR3-ECD complexes were stable to the above-mentioned RMSD for a given time window of 10 nanoseconds at stable temperature (∼278 K) and pressure (∼1 atm). Molecular docking and molecular dynamics simulation study of all the MPVs & TLR3-ECD complexes suggests a stable complex formation tendency for all the MPVs with TLR3-ECD.

**Figure 8.**
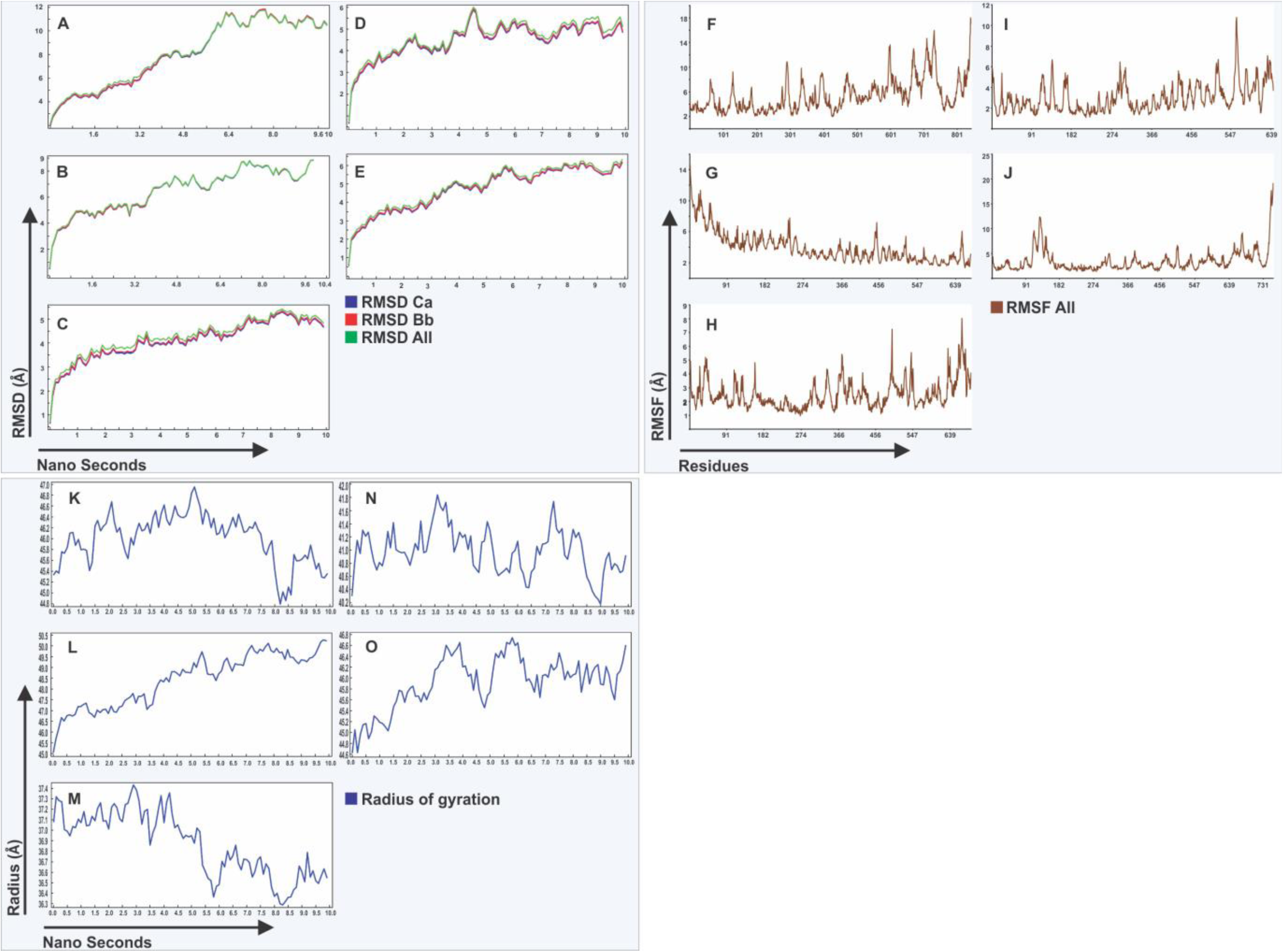
Molecular Dynamics simulation study of MPVs and TLR3-ECD complexes. (A-E): Root Mean Square Deviation (RMSD) for Cα, Backbone and all atoms (RMSD Ca, RMSD Bb, & RMSD All) respectively for (A) CTL-MPV-1:TLR3-ECD, (B) CTL-MPV-2:TLR3-ECD, (C) CTL-MPV-3:TLR3-ECD, (D) HTL-MPV-1:TLR3-ECD, (E) HTL-MPV-2:TLR3-ECD complexes. (F-J) Root Mean Square Fluctuation (RMSF) in the conformation of residues of the MPVs in complex with TLR3-ECD. RMSF: (F) CTL-MPV-1:TLR3-ECD, (G) CTL-MPV-2:TLR3-ECD, (H) CTL-MPV-3:TLR3-ECD, (I) HTL-MPV-1:TLR3-ECD, (J) HTL-MPV-2:TLR3-ECD. (K-O) Radius of gyration for all the MPVs and TLR3-ECD complexes are shown. Rg: (K) CTL-MPV-1:TLR3-ECD, (L) CTL-MPV-2:TLR3-ECD, (M) CTL-MPV-3:TLR3-ECD, (N) HTL-MPV-1:TLR3-ECD, (O) HTL-MPV-2:TLR3-ECD.

Further, most of the amino acid residues of the CTL and HTL MPVs in complexed with TLR3-ECD have shown to have Root Mean Square Fluctuation (RMSF) from their initial confirmation, for all the amino acid residues of the MPVs, in acceptable range [(CTL-MPV-1:TLR3-ECD, RMSF: ∼2 to ∼14 Å) (CTL-MPV-2:TLR3-ECD, RMSF: ∼2 to ∼10.5 Å) (CTL-MPV-3:TLR3-ECD, RMSF: ∼2 to ∼8 Å) (HTL-MPV-1:TLR3-ECD, RMSF: ∼2 to ∼10 Å) (HTL-MPV-2:TLR3-ECD, RMSF: ∼2 to ∼10 Å)] (Figure 8: F-J). These results suggest that all the MPVs and TLR3-CED complexes to have a stable nature.

Furthermore, the radius of gyration analysis for all the MPVs and TLR3-ECD complexes was performed. The Radius of gyration (Rg) indicates the compactness of the complex conformation. Rg is concern with how regular secondary structures are compactly packed in tertiary structure of protein complex throughout the MD simulation. The variation in Rg for all the MPVs and TLR3-ECD complexes remain in acceptable range throughout the MD simulation [(CTL-MPV-1:TLR3-ECD, Rg: ∼44.8 to ∼47.0 Å) (CTL-MPV-2:TLR3-ECD, Rg: ∼46.5 to ∼50.0 Å) (CTL-MPV-3:TLR3-ECD, Rg: ∼36.3 to ∼37.4 Å) (HTL-MPV-1:TLR3-ECD, Rg: ∼40.2 to ∼41.8 Å) (HTL-MPV-2:TLR3-ECD, Rg: ∼44.6 to ∼46.6 Å)] (Figure 8: K-O). This suggests that the MPVs:TLR3-ECD complexes remain structurally compact throughout the 10 ns MD simulation analysis.

#### Analysis of MPVs cDNA for expression in human host cell line

All the three CTL and two HTL MPV construct were further analyzed for feasibility for expression in vitro. Codon-optimized cDNA (complementary DNA) for all the CTL and HTL MPVs were generated for expression in the mammalian host cell line (Human) by the Java Codon Adaptation Tool. The generated optimized cDNA for all the MPVs were further analyzed for their expression feasibility in vitro, by the GenScript Rare Codon Analysis Tool. The analysis revealed that the codon-optimized cDNA of all the CTL and HTL MPVs satisfy all the crucial parameters like GC content, CAI (Codon Adaptation Index) score and tandem rare codons for high-level expression in a mammalian cell line (Human) (Supplementary table S14). The GC content of a cDNA, the CAI score that indicates the possibility of cDNA expression in a chosen expression system (here, human cell line), and the tandem rare codon frequency that indicates the presence of low-frequency codons in cDNA, all these parameters were in acceptable range for all the cDNA of the MPVs. The tandem rare codons which may hinder the proper expression of the cDNA or even interrupt the translational machinery of the chosen expression system was observed to be 0% in all the MPVs. Hence, as per the GenScript Rare Codon analysis, the cDNA of all the MPVs have a high potential for large-scale expression in the human cell line. The workflow concept, from Ag-Patch identification to codon optimization of cDNA and in vivo trial for the here proposed MPVs is shown in Supplementary figure S4.

## DISCUSSION

Majority of the vaccine design against SARS-CoV-2 have put their focus on the single protein, protein subunits or the epitopes from SARS-CoV-2 proteins, mostly S, E, M, N and ORF1ab proteins. The recent strategies to design and develop vaccine to combat SARS-CoV-2 involve subunit vaccines or multi-epitope vaccines. The subunit vaccine involves use of single protein or a multiple subunits of SARS-CoV-2 proteins. On the other hand the multi-epitope vaccine involves fusion of multiple epitopes identified from the proteome of the SARS-CoV-2 by short peptide linkers. Many of the subunit based and multi-epitope based designs have been published claiming potential to activate CD4 and CD8 T cell immune response driving long-term robust adaptive immunity in the vast majority of the population (Abdelmageed et al., 2020; Ahmed et al., 2020; Akhand et al., 2020; An et al., 2000; Banerjee et al., 2020; Baruah et al., 2020; Bhattacharya et al., 2020; Fast et al., 2020; Gragert et al., 2013; Gupta et al., 2020; Herst et al., 2020; Ismail et al., 2020; Khan et al., 2020; Lee et al., 2020; Liu et al., 2020; Lu et al., 2014; Mitra et al., 2020; Nerli et al., 2020; Poran et al., 2020; Ramaiah et al., 2020; Saha et al., 2020; Sheikhshahrokh et al., 2020; Singh et al., 2020; Srivastava et al., 2020a; Srivastava et al., 2020b; ul Qamar et al., 2020; Vashi et al., 2020; Yarmarkovich and Farrel et al., 2020; Yazdani et al., 2020). Numerous highly antigenic regions have also been reported from SARS-CoV-2 proteins which have been recognized on the basis of large population coverage by favorable binding with large number of HLA allele distributed amongst different ethnic human population worldwide (Grifoni et al., 2020b; Yarmarkovich and Warrington et al., 2020).

Although the major focus is on epitope or subunit based vaccine design, the antigen protein subunit and the specific epitope used to design multi-epitope vaccines have some key drawbacks. The Major drawback with single protein or multiple subunit based vaccine is the limited efficiency of the vaccines. On the other hand the epitope based vaccines have major drawback of frequent mutations in the proteome of the SARS-CoV-2 virus. The multi-epitope based vaccines have one more major challenge for presentation of the chosen epitopes by the Antigen Presenting Cells (APC). The chop down processing by proteasome and lysosome would leave a very narrow chance for the epitopes of the multi-epitope vaccines, to remain intact and successfully presented by APC.

In the present study, we have reported a novel method to design a vaccine against SARS-CoV-2 by utilizing multiple antigenic patches from the viral proteins. The Ag-Patches used have been identified by the clusters of overlapping epitopes. The identification of these Ag-Patches was performed by reverse epitomics analysis, of the high scoring CTL and HTL epitopes screened from all the ORF proteins of the SARS-CoV-2 virus. The method is here termed as “Overlapping-epitope-clusters-to-patches” method. All the screened epitopes were well characterized for their conservancy, immunogenicity, non-toxicity and large population coverage. The clusters of the overlapping epitopes lead us to identify the Ag-Patches. These Ag-Patches from all the ORF proteins of the SARS-CoV-2 proteome were utilized further to design Multi-Patch Vaccine (MPV) candidate against the SARS-CoV-2 infection.

The designed MPVs from the antigenic patches of SARS-CoV-2 proteins have several advantages over to the subunit and multi-epitope based vaccines. The Ag-Patches utilized were identified and collected from the entire proteome of the SARS-CoV-2. This would enhance the efficiency of the vaccines and lead the vaccine to be more effective. The MPVs consists of the identified Ag-Patches will have potential to raise multiple epitopes in clusters upon the chop down processing by proteasome and lysosome in the Antigen Presenting Cell (APC). The identified Ag-Patches will also provide larger chance of the epitopes raised after proteasome and lysosomal processing to get presentation by the APC and elicit immune response. Since the Ag-Patches were identified by the large number of epitopes forming clusters, the MPVs designed would have potential to raise larger number of epitopes upon proteasome and lysosomal processing, hence larger number of HLA alleles could be targeted and hence larger ethnic human population could be covered by the MPVs, in comparison to the limited number of epitope used in multi-epitope vaccines. For instance, the three MPVs designed in the present study used the Ag-Patches which were identified by 768 (518 CTL and 250 HTL epitopes) overlapping epitope targeting different HLA alleles. Such inclusion of large number of epitopes and targeting large number of HLA alleles is not possible for multi-epitope based vaccines prepared with limited epitopes.

All the identified Ag-Patches utilized to design MPVs have shown to be highly conserved amongst the protein sequences of SARS-CoV-2 available at NCBI protein database. All the physicochemical properties of SARS-CoV-2, favors their overexpression in vitro. This is further supported by the cDNA analysis of the codon-optimized constructs of all the MPVs for high expression in a mammalian cell line (human). Furthermore, the designed MPVs also show stable binding tendency with the TLR3-ECD which is an essential criteria for an antigen to be recognized and processed by the human immune system.

## CONCLUSION

In the present study we have identified highly immunogenic novel antigenic patches (Ag-Patches) (73 CTL and 49 HTL) from the entire proteome of SARS CoV-2. The Ag-Patches were identified by a novel reverse epitomics approach, “overlapping-epitope-clusters-to-patches” method. These Ag-Patches are highly conserved in nature and found in most of the SARS-CoV-2 protein sequences available at NCBI protein database. These Ag-Patches are identified on the basis of high scoring, immunogenic, overlapping epitopes thoroughly screened from entire proteome of SARS-CoV-2. Further, for the first time we have utilized the identified multiple immunogenic Ag-Patches to design novel Multi-Patch Vaccine (MPV) which proposes a new methodology for vaccine design. The MPVs designed against SARS-CoV-2 in our study have potential to give raise up to a total of 768 epitopes (518 CTL and 250 HTL epitopes) targeting large number of different HLA alleles. Such large number of epitopes is not possible for multi-epitope based vaccine design. The large number of epitopes covered implies large number of HLA alleles targeted. The large number of HLA allele targeted further implies large ethnic human population coverage worldwide. The MPVs, with multiple epitope cluster based Ag-Patches from entire proteome of SARS-CoV-2, are in better position to raise multiple epitopes in comparison to the MEVs upon proteasome or lysosomal chop down processing by the APC. The designed MPVs against SARS-CoV-2 are validated for stable complex formation with ectodomain of TLR3. The physiochemical properties and the codon-optimized cDNA analysis of all the MPVs designed suggest favored large scale expression potential. We conclude that the novel Multi-Patch Vaccine designed by the novel antigenic patches (Ag-Patches) could be a highly potential novel method to combat SARS-CoV-2, with greater effectiveness, high specificity and large human population coverage worldwide.

## AUTHOR CONTRIBUTION

Idea conceived, Methodology designed and performed, Antigenic Patches (Ag-Patches) identified, MPVs designed, Scientific writing: SS; MD Simulation performed: S.V.; Scientific reporting and revising the manuscript: S.S., S.V., M.K., D.A., A.K.S., M.Kolbe, S.Singh., A.K., B.R., S.A.N., H-J.S., K.V., K.C.P.

## DECLARATION OF INTEREST

Two patents have been filed from the report.

## ACKNOWLEDGEMENTS

We are thankful to the Indian Council for Medical Research (ICMR) for providing SRF fellowship to Sonia Verma (ISRM/11(98)/2019). We also thank Rajni Kant Dixit, Scientist D, of ICMR-National Institute of Malaria Research (NIMR) for providing computational facility for experimentations. We also thank Indian Foundation for Fundamental Research for providing the computational and other infrastructural facility for experimentations.

---

**Supplementary table S1:** High Percentile Ranking CTL epitopes-HLA allele pairs screened from entire proteome of SARS-CoV-2 by the “MHC-I Processing Predictions” tool of IEDB. These epitopes were further utilized to identify the potentially immunogenic multiple epitope cluster based CTL Ag-Patches from the entire proteome of the SRAS-CoV-2. The epitopes shown in **<u>RED</u>** are the epitope which form Overlapping epitope clusters. The screened epitopes are in consensus with the previous studies [Srivastava et al. 2020a; Srivastava et al. 2020b; and Grifoni et al., 2020a].

**Supplementary table S2:** High Scoring CTL epitopes-HLA allele pairs screened from entire proteome of SARS-CoV-2 by the “MHC-I Binding Predictions” tool of IEDB. These epitopes were further utilized to identify the potentially immunogenic multiple epitope cluster based CTL Ag-Patches from the entire proteome of the SRAS-CoV-2. The epitopes shown in **<u>RED</u>** are the epitope which form Overlapping epitope clusters. The screened epitopes are in consensus with the previous studies [Srivastava et al. 2020a; Srivastava et al. 2020b; and Grifoni et al., 2020a].

**Supplementary table S3:** High Percentile Ranking HTL epitopes-HLA allele pairs screened from entire proteome of SARS-CoV-2 by the “MHC-II Binding Predictions” tool of IEDB. These epitopes were further utilized to identify the potentially immunogenic multiple epitope cluster based HTL Ag-Patches from the entire proteome of the SRAS-CoV-2. The epitopes shown in **<u>BLUE</u>** are the epitope which form Overlapping epitope clusters. The screened epitopes are in consensus with the previous studies [Srivastava et al. 2020a; Srivastava et al. 2020b; and Grifoni et al., 2020a].

**Supplementary table S4:** Population coverage by all the overlapping CTL and HTL epitopes forming epitope clusters.

**Supplementary table S5:** Construct of CTL-MPV-1, CTL-MPV-2, CTL-MPV-3, HTL-MPV-1, and HTL-MPV-2. Physicochemical property analysis based on the amino acid sequences of all the designed three CTL and two HTL Multi-Patch Vaccines.

**Supplementary table S6:** INF-γ inducing POSITIVE epitopes with a score of 1 or more than 1, screened from the CTL MPVs.

**Supplementary table S7:** INF-γ inducing POSITIVE epitopes with a score of 1 or more than 1, screened from the HTL MPVs.

**Supplementary table S8:** Parameters for the tertiary structure homology modeling of all the CTL and HTL MPVs by the I-TASSER tool.

**Supplementary table S9:** Refinement parameter values for CTL and HTL MPV models after refinement by GalaxyRefine tool.

**Supplementary table S10:** B cell linear epitopes screened from CTL MPVs.

**Supplementary table S11:** B cell Discontinuous epitopes screened from CTL MPVs.

**Supplementary table S12:** B cell linear epitopes screened from HTL MPVs.

**Supplementary table S13:** B cell Discontinuous epitopes screened from HTL MPVs.

**Supplementary table S14:** Analysis of codon-optimized cDNA of all the MPVs.

**Supplementary figure S1.**
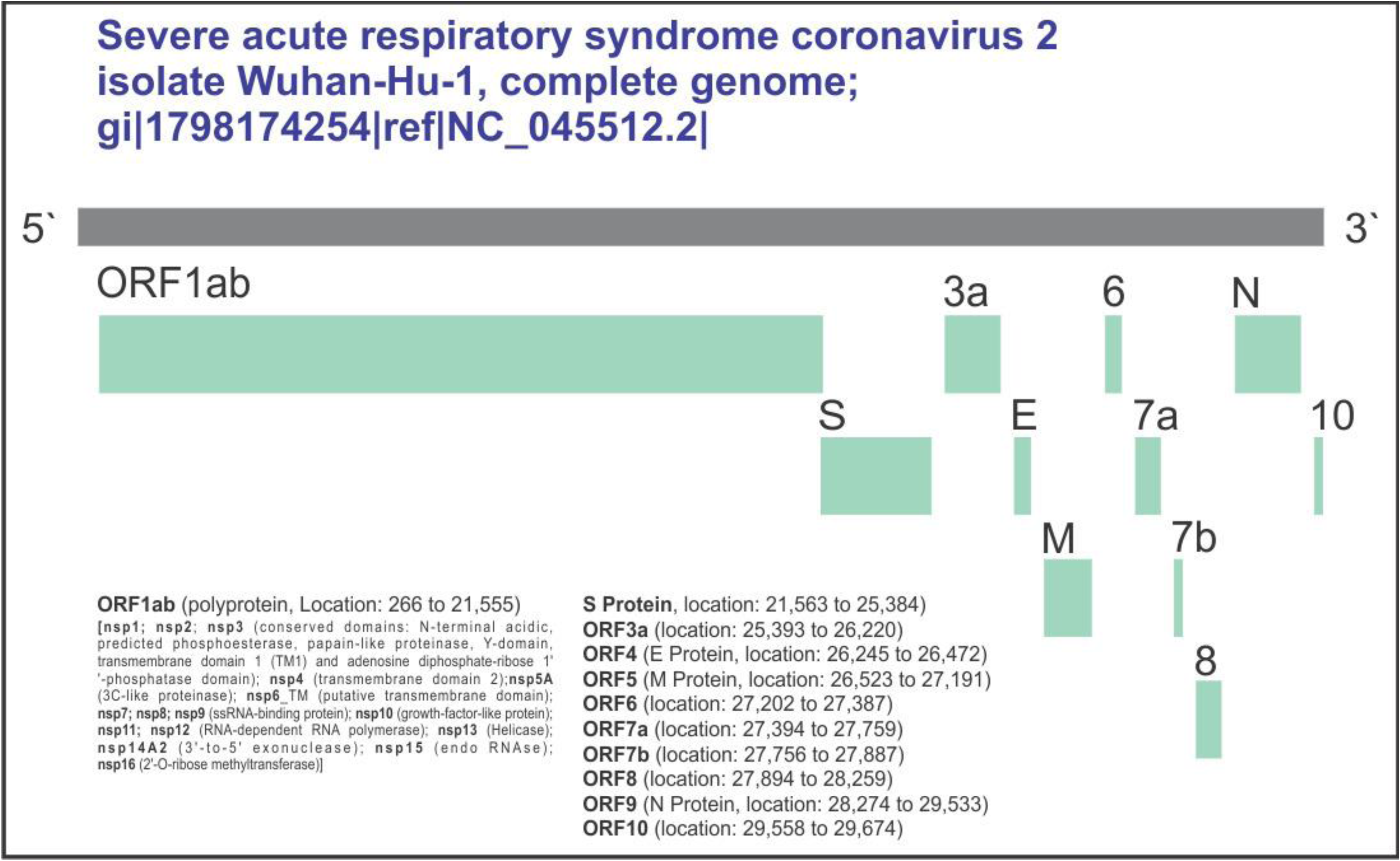
Schematic presentation of all the ORF protein expressed by SARS-CoV-2 genome.

**Supplementary figure S2.**
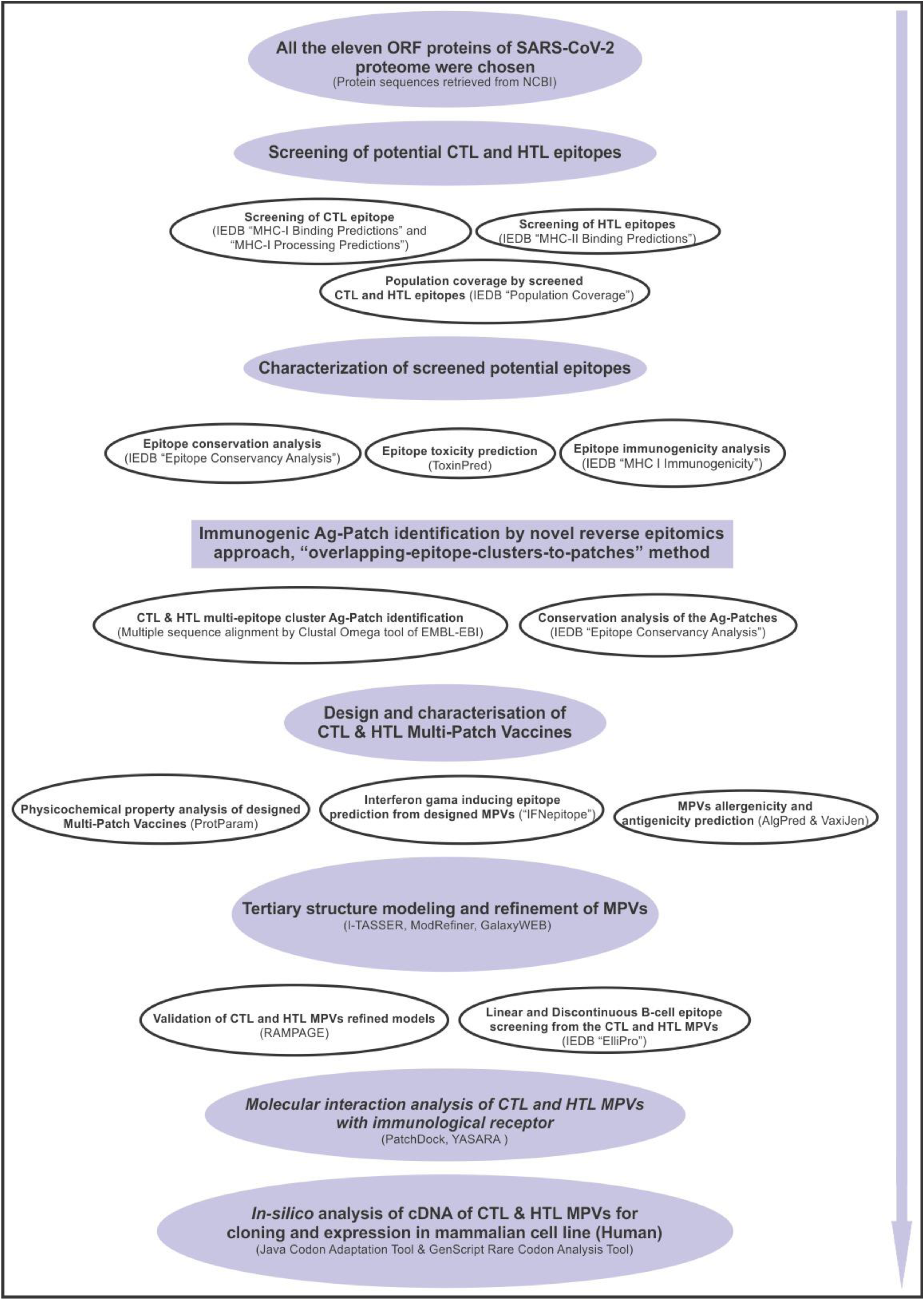
Schematic representation of workflow and methodology.

**Supplementary figure S3.**
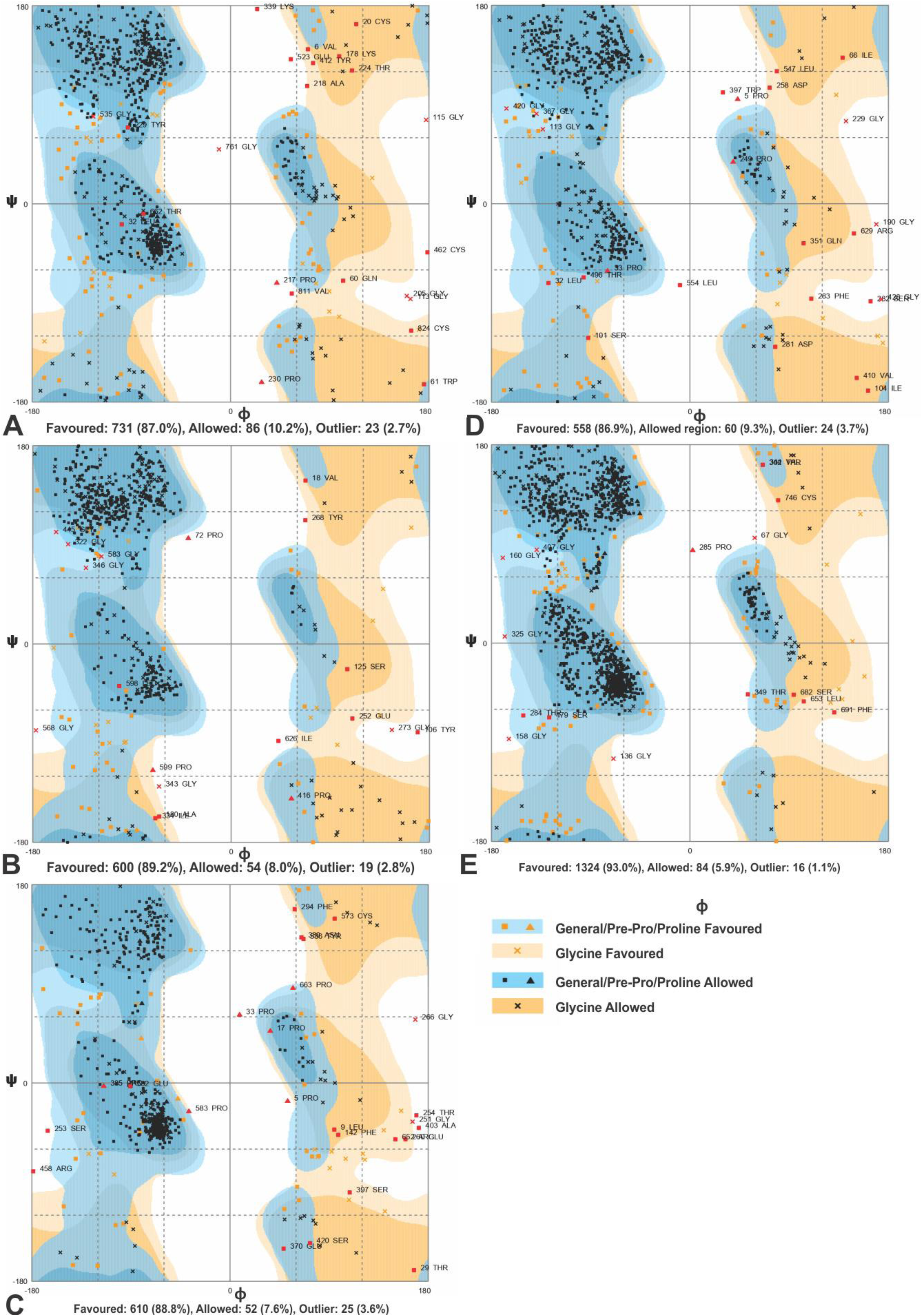
RAMPAGE analysis for all the MPVs **(A)** CTL-MPV-1, **(B)** CTL-MPV-2, **(C)** CTL-MPV-3, **(D)** HTL-MPV-1, **(E)** HTL-MPV-2.

**Supplementary figure S4.**
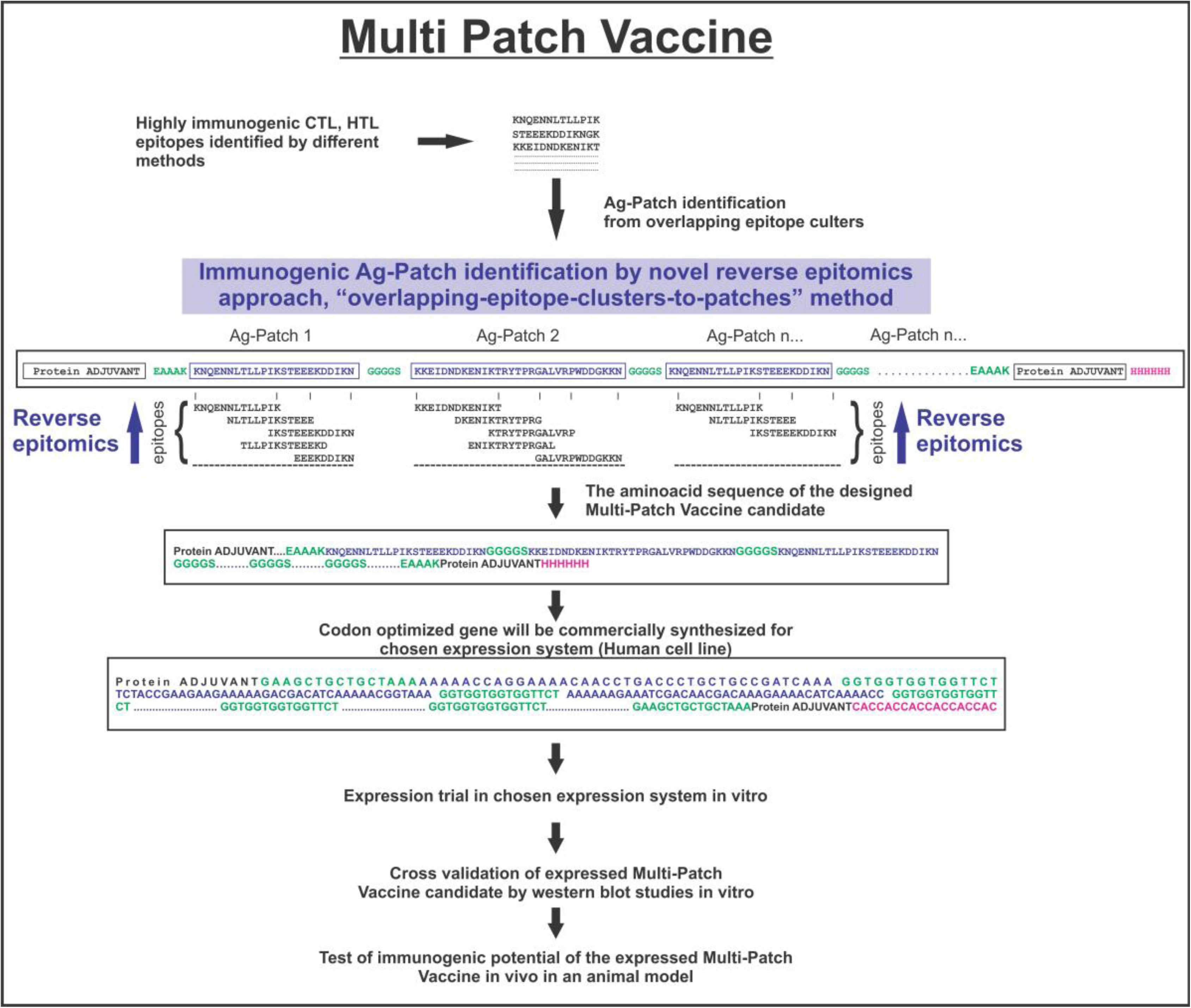
Work flow concept chart from Ag-Patch (antigenic patch) identification to in vivo trial for the proposed MPVs against SARS-CoV-2.

## Notes

### Competing Interest Statement

The authors have declared no competing interest.

## REFERENCES

Abdelmageed, M.I., Abdelmoneim, A.H., Mustafa, M.I., Elfadol, N.M., Murshed, N.S., Shantier, S.W. and Makhawi, A.M., 2020. Design of multi epitope-based peptide vaccine against E protein of human 2019-nCoV: An immunoinformatics approach. bioRxiv 2020. *Preprint*, pp.1–21.

Ahmed, S.F., Quadeer, A.A. and McKay, M.R., 2020. Preliminary identification of potential vaccine targets for the COVID-19 coronavirus (SARS-CoV-2) based on SARS-CoV immunological studies. Viruses, 12(3), p.254.

Akhand, M.R.N., Azim, K.F., Hoque, S.F., Moli, M.A., Joy, B.D., Akter, H., Afif, I.K., Ahmed, N. and Hasan, M., 2020. Genome based Evolutionary study of SARS-CoV-2 towards the Prediction of Epitope Based Chimeric Vaccine. bioRxiv.

An, L.L., Rodriguez, F., Harkins, S., Zhang, J. and Whitton, J.L., 2000. Quantitative and qualitative analyses of the immune responses induced by a multivalent minigene DNA vaccine. Vaccine, 18(20), pp.2132–2141.

Antoniou, A.N., Powis, S.J. and Elliott, T., 2003. Assembly and export of MHC class I peptide ligands. Current opinion in immunology, 15(1), pp.75–81.

Banerjee, A., Santra, D. and Maiti, S., 2020. Energetics based epitope screening in SARS CoV-2 (COVID 19) spike glycoprotein by Immuno-informatic analysis aiming to a suitable vaccine development. BioRxiv.

Baruah, V. and Bose, S., 2020. Immunoinformatics-aided identification of T cell and B cell epitopes in the surface glycoprotein of 2019-nCoV. Journal of medical virology, 92(5), pp.495–500.

Bell, J.K., Botos, I., Hall, P.R., Askins, J., Shiloach, J., Segal, D.M. and Davies, D.R., 2005. The molecular structure of the Toll-like receptor 3 ligand-binding domain. Proceedings of the National Academy of Sciences of the United States of America, 102(31), pp.10976–10980.

Bhattacharya, M., Sharma, A.R., Patra, P., Ghosh, P., Sharma, G., Patra, B.C., Lee, S.S. and Chakraborty, C., 2020. Development of epitope-based peptide vaccine against novel coronavirus 2019 (SARS-COV-2): Immunoinformatics approach. Journal of medical virology, 92(6), pp.618–631.

Biragyn, A., Surenhu, M., Yang, D., Ruffini, P.A., Haines, B.A., Klyushnenkova, E., Oppenheim, J.J. and Kwak, L.W., 2001. Mediators of innate immunity that target immature, but not mature, dendritic cells induce antitumor immunity when genetically fused with nonimmunogenic tumor antigens. The Journal of Immunology, 167(11), pp.6644–6653.

Bui H. H, Sidney J, Dinh K, Southwood S, Newman M. J, Sette A. 2006. Predicting population coverage of T-cell epitope-based diagnostics and vaccines. BMC Bioinformatics 17:153. (http://tools.iedb.org/population/)

Bui HH, Sidney J, Li W, Fusseder N, Sette A. 2007. Development of an epitope conservancy analysis tool to facilitate the design of epitope-based diagnostics and vaccines. BMC Bioinformatics 8:361. (http://tools.iedb.org/conservancy/)

Calis JJA, Maybeno M, Greenbaum JA, Weiskopf D, De Silva AD, Sette A, Kesmir C, Peters B. 2013. Properties of MHC class I presented peptides that enhance immunogenicity. PloS Comp. Biol. 8(1):361. (http://tools.iedb.org/immunogenicity/)

Case, D.A., Babin, V., Berryman, J.T., Betz, R.M., Cai, Q., Cerutti, D.S., Cheatham III, T.E., Darden, T.A., Duke, R.E., Gohlke, H. and Goetz, A.W., 2014. The FF14SB force field. Amber, 14, pp.29–31.

Chen, X., Zaro, J.L. and Shen, W.C., 2013. Fusion protein linkers: property, design and functionality. Advanced drug delivery reviews, 65(10), pp.1357–1369.

Delneste, Y., Beauvillain, C. and Jeannin, P., 2007. Innate immunity: structure and function of TLRs. Medecine sciences: M/S, 23(1), pp.67–73.

Dhanda, S. K., Vir, P. & Raghava, G. P. Designing of interferon-gamma inducing MHC class-II binders. Biol. Direct. 8, 30 (2013).

Dong Xu and Yang Zhang. Improving the Physical Realism and Structural Accuracy of Protein Models by a Two-step Atomic-level Energy Minimization. Biophysical Journal, vol 101, 2525–2534 (2011). (https://zhanglab.ccmb.med.umich.edu/ModRefiner/)

Doytchinova I.A., and Flower D.R., VaxiJen: a server for prediction of protective antigens, tumour antigens and subunit vaccines. BMC Bioinformatics. 2007 8:4. (http://www.ddg-pharmfac.net/vaxijen/VaxiJen/VaxiJen.html)

Duhovny D, Nussinov R, Wolfson HJ. Efficient Unbound Docking of Rigid Molecules. In Gusfield et al., Ed. Proceedings of the 2’nd Workshop on Algorithms in Bioinformatics(WABI) Rome, Italy, Lecture Notes in Computer Science 2452, pp. 185–200, Springer Verlag, 2002

Duits, L.A., Nibbering, P.H., van Strijen, E., Vos, J.B., Mannesse-Lazeroms, S.P., van Sterkenburg, M.A. and Hiemstra, P.S., 2003. Rhinovirus increases human β-defensin-2 and-3 mRNA expression in cultured bronchial epithelial cells. FEMS Immunology & Medical Microbiology, 38(1), pp.59–64.

Farina, C., Krumbholz, M., Giese, T., Hartmann, G., Aloisi, F. and Meinl, E., 2005. Preferential expression and function of Toll-like receptor 3 in human astrocytes. Journal of neuroimmunology, 159(1-2), pp.12–19.

Fast, E. and Chen, B., 2020. Potential T-cell and B-cell Epitopes of 2019-nCoV. bioRxiv.

Gasteiger, E., Hoogland, C., Gattiker, A., Duvaud, S.E., Wilkins, M.R., Appel, R.D. and Bairoch, A., 2005. Protein identification and analysis tools on the ExPASy server (pp. 571–607). Humana Press. (https://web.expasy.org/protparam/)

Gragert, L., Madbouly, A., Freeman, J. and Maiers, M., 2013. Six-locus high resolution HLA haplotype frequencies derived from mixed-resolution DNA typing for the entire US donor registry. Human immunology, 74(10), pp.1313–1320.

Grifoni, A., Sidney, J., Zhang, Y., Scheuermann, R.H., Peters, B. and Sette, A., 2020a. Candidate targets for immune responses to 2019-Novel Coronavirus (nCoV): sequence homology- and bioinformatic-based predictions. CELL-HOST-MICROBE-D-20-00119.

Grifoni, A., Sidney, J., Zhang, Y., Scheuermann, R.H., Peters, B. and Sette, A., 2020b. A sequence homology and bioinformatic approach can predict candidate targets for immune responses to SARS-CoV-2. Cell host & microbe.

Gupta, E., Mishra, R.K. and Niraj, R.R.K., 2020. Identification of potential vaccine candidates against SARS-CoV-2, A step forward to fight novel coronavirus 2019-nCoV: A Reverse Vaccinology Approach. bioRxiv.

Gupta, S., Kapoor, P., Chaudhary, K., Gautam, A., Kumar, R., Raghava, G.P. and Open Source Drug Discovery Consortium, 2013. In silico approach for predicting toxicity of peptides and proteins. PLoS One, 8(9), p.e73957. (http://crdd.osdd.net/raghava/toxinpred/multi_submit.php)

Hajighahramani, N., Nezafat, N., Eslami, M., Negahdaripour, M., Rahmatabadi, S.S. and Ghasemi, Y., 2017. Immunoinformatics analysis and in silico designing of a novel multi-epitope peptide vaccine against Staphylococcus aureus. Infection, Genetics and Evolution, 48, pp.83–94.

Herst, C.V., Burkholz, S., Sidney, J., Sette, A., Harris, P.E., Massey, S., Brasel, T., Cunha-Neto, E., Rosa, D.S., Chao, W.C.H. and Carback, R., 2020. An effective CTL peptide vaccine for ebola zaire based on survivors’ CD8+ targeting of a particular nucleocapsid protein epitope with potential implications for COVID-19 vaccine design. Vaccine.

Hoof, I., Peters, B., Sidney, J., Pedersen, L.E., Sette, A., Lund, O., Buus, S. and Nielsen, M., 2009. NetMHCpan, a method for MHC class I binding prediction beyond humans. Immunogenetics, 61(1), p.1.

Hoover, D.M., Rajashankar, K.R., Blumenthal, R., Puri, A., Oppenheim, J.J., Chertov, O. and Lubkowski, J., 2000. The structure of human β-defensin-2 shows evidence of higher order oligomerization. Journal of Biological Chemistry, 275(42), pp.32911–32918. PDB ID: 1FD3.

Hu W, Li F, Yang X, Li Z, Xia H, Li G, Wang Y, Zhang Z. A flexible peptide linker enhances the immunoreactivity of two copies HBsAg preS1 (21-47) fusion protein. J Biotechnol. 2004;107:83–90.

Ismail, S., Ahmad, S. and Azam, S.S., 2020. Immuno-informatics Characterization SARS-CoV-2 Spike Glycoprotein for Prioritization of Epitope based Multivalent Peptide Vaccine. bioRxiv.

Kahan, B.D., 2003. Individuality: the barrier to optimal immunosuppression. Nature Reviews Immunology, 3(10), pp.831–838.

Khan, A., Alam, A., Imam, N., Siddiqui, M.F. and Ishrat, R., 2020. Design of an Epitope-Based Peptide Vaccine against the Severe Acute Respiratory Syndrome Coronavirus-2 (SARS-CoV-2): A Vaccine Informatics Approach. bioRxiv.

Ko, J., Park, H., Heo, L., and Seok, C., GalaxyWEB server for protein structure prediction and refinement, Nucleic Acids Res. 40 (W1), W294–W297 (2012). (http://galaxy.seoklab.org/cgi-bin/submit.cgi?type=REFINE)

Kohlgraf, K.G., Pingel, L.C., Dietrich, D.E. and Brogden, K.A., 2010. Defensins as anti-inflammatory compounds and mucosal adjuvants. Future microbiology, 5(1), pp.99–113.

Krieger, E. and Vriend, G., 2015. New ways to boost molecular dynamics simulations. Journal of computational chemistry, 36(13), pp.996–1007.

Kringelum, J.V., Lundegaard, C., Lund, O. and Nielsen, M., 2012. Reliable B cell epitope predictions: impacts of method development and improved benchmarking. PLoS computational biology, 8(12).

Lee, C.H. and Koohy, H., 2020. In silico identification of vaccine targets for 2019-nCoV. F1000Research, 9.

Liu, G., Carter, B., Bricken, T., Jain, S., Viard, M., Carrington, M. and Gifford, D.K., 2020. Computationally Optimized SARS-CoV-2 MHC Class I and II Vaccine Formulations Predicted to Target Human Haplotype Distributions. Cell systems.

Lovell, S.C., Davis, I.W., Arendall, W.B., de Bakker, P.I.W., Word, J.M., Prisant, M.G., Richardson, J.S. and Richardson, D.C., 2003. Structure validation by Calpha geometry: phi, psi and Cbeta deviation. Proteins 50, 437e450. (http://mordred.bioc.cam.ac.uk/~rapper/rampage.php)

Lu, Y.C., Yao, X., Crystal, J.S., Li, Y.F., El-Gamil, M., Gross, C., Davis, L., Dudley, M.E., Yang, J.C., Samuels, Y. and Rosenberg, S.A., 2014. Efficient identification of mutated cancer antigens recognized by T cells associated with durable tumor regressions. Clinical Cancer Research, 20(13), pp.3401–3410.

Maier, J.A., Martinez, C., Kasavajhala, K., Wickstrom, L., Hauser, K.E. and Simmerling, C., 2015. ff14SB: improving the accuracy of protein side chain and backbone parameters from ff99SB. Journal of chemical theory and computation, 11(8), pp.3696–3713.

Melief, C.J., 2005. Escort service for cross-priming. Nature immunology, 6(6), pp.543–544.

Mitra, D., Shekhar, N., Pandey, J., Jain, A. and Swaroop, S., 2020. Multi-epitope based peptide vaccine design against SARS-CoV-2 using its spike protein. bioRxiv.

Morla, S., Makhija, A. & Kumar, S. Synonymous codon usage pattern in glycoprotein gene of rabies virus. Gene. 584, 1–6 (2016). (http://www.jcat.de/).

Nagpal, G., Gupta, S., Chaudhary, K., Dhanda, S.K., Prakash, S. and Raghava, G.P., 2015. VaccineDA: Prediction, design and genome-wide screening of oligodeoxynucleotide-based vaccine adjuvants. Scientific reports, 5, p.12478. (http://crdd.osdd.net/raghava/ifnepitope/scan.php)

NCBI: Severe acute respiratory syndrome coronavirus 2 isolate Wuhan-Hu-1, complete genome; gi|1798174254|ref|NC_045512.2|;https://www.ncbi.nlm.nih.gov/projects/sviewer/?id=NC_045512&tracks=[key:sequence_track,name:Sequence,display_name:Sequence,id:STD649220238,annots:Sequence,ShowLabel:false,ColorGaps:false,shown:true,order:1][key:gene_model_track,name:Genes,display_name:Genes,id:STD3194982005,annots:Unnamed,Options:ShowAllButGenes,CDSProductFeats:true,NtRuler:true,AaRuler:true,HighlightMode:2,ShowLabel:true,shown:true,order:9]&v=1:29903&c=null&select=null&slim=0

Nerli, S. and Sgourakis, N.G., 2020. Structure-based modeling of SARS-CoV-2 peptide/HLA-A02 antigens. bioRxiv.

Nielsen, M., Lundegaard, C. and Lund, O., 2007. Prediction of MHC class II binding affinity using SMM-align, a novel stabilization matrix alignment method. BMC bioinformatics, 8(1), p.238.

Peters B, Bulik S, Tampe R, Van Endert PM, Holzhutter HG. 2003. Identifying MHC class I epitopes by predicting the TAP transport efficiency of epitope precursors. J Immunol171:1741–1749.

Ponomarenko JV, Bui H, Li W, Fusseder N, Bourne PE, Sette A, Peters B. 2008. ElliPro: a new structure-based tool for the prediction of antibody epitopes. BMC Bioinformatics 9:514. (http://tools.iedb.org/ellipro/)

Poran, A., Harjanto, D., Malloy, M., Rooney, M.S., Srinivasan, L. and Gaynor, R.B., 2020. Sequence-based prediction of vaccine targets for inducing T cell responses to SARS-CoV-2 utilizing the bioinformatics predictor RECON. bioRxiv.

Ramaiah, A. and Arumugaswami, V., 2020. Insights into cross-species evolution of novel human coronavirus 2019-nCoV and defining immune determinants for vaccine development. bioRxiv.

Ramakrishnan, C. and Ramachandran, G.N., 1965. Stereochemical criteria for polypeptide and protein chain conformations: II. Allowed conformations for a pair of peptide units. Biophysical journal, 5(6), pp.909–933.

Roy, A., Kucukural, A. and Zhang, Y., 2010. I-TASSER: a unified platform for automated protein structure and function prediction. Nature protocols, 5(4), p.725. (https://zhanglab.ccmb.med.umich.edu/I-TASSER/)

Saha, R. and Prasad, B.V., 2020. In silico approach for designing of a multi-epitope based vaccine against novel Coronavirus (SARS-COV-2). bioRxiv.

Saha, S. & Raghava, G. AlgPred: prediction of allergenic proteins and mapping of IgE epitopes. Nucleic. Acids. Res. 34, W202–W209 (2006). (http://crdd.osdd.net/raghava/algpred/submission.html)

Schneidman-Duhovny D, Inbar Y, Nussinov R, Wolfson HJ. PatchDock and SymmDock: servers for rigid and symmetric docking. Nucl. Acids. Res. 33: W363–367, 2005. (http://bioinfo3d.cs.tau.ac.il/PatchDock/)

Sheikhshahrokh, A., Ranjbar, R., Saeidi, E., Dehkordi, F.S., Heiat, M., Ghasemi-Dehkordi, P. and Goodarzi, H., 2020. Frontier therapeutics and vaccine strategies for sars-cov-2 (COVID-19): A review. Iranian Journal of Public Health, 49, pp.18–29.

Shin, W.H., Lee, G.R., Heo, L., Lee, H. and Seok, C., 2014. Prediction of protein structure and interaction by GALAXY protein modeling programs. Bio Design, 2(1), pp.1–11.

Sidney, J., Assarsson, E., Moore, C., Ngo, S., Pinilla, C., Sette, A. and Peters, B., 2008. Quantitative peptide binding motifs for 19 human and mouse MHC class I molecules derived using positional scanning combinatorial peptide libraries. Immunome research, 4(1), p.2.

Sievers, F., Wilm, A., Dineen, D., Gibson, T.J., Karplus, K., Li, W., Lopez, R., McWilliam, H., Remmert, M., Söding, J. and Thompson, J.D., and Higgins D.G., 2011. Fast, scalable generation of high-quality protein multiple sequence alignments using Clustal Omega. Molecular systems biology, 7(1), p.539. (https://www.ebi.ac.uk/Tools/msa/clustalo/)

Singh, A., Thakur, M., Sharma, L.K. and Chandra, K., 2020. Designing a multi-epitope peptide-based vaccine against SARS-CoV-2. bioRxiv.

Srivastava, S., Kamthania, M., Kumar Pandey, R., Kumar Saxena, A., Saxena, V., Kumar Singh, S., Kumar Sharma, R. and Sharma, N., 2019. Design of novel multi-epitope vaccines against severe acute respiratory syndrome validated through multistage molecular interaction and dynamics. Journal of Biomolecular Structure and Dynamics, 37(16), pp.4345–4360.

Srivastava, S., Kamthania, M., Singh, S., Saxena, A.K. and Sharma, N., 2018. Structural basis of development of multi-epitope vaccine against middle east respiratory syndrome using in silico approach. Infection and drug resistance, 11, p.2377.

Srivastava, S., Verma, S., Kamthania, M., Kaur, R., Badyal, R.K., Saxena, A.K., Shin, H.J., Kolbe, M. and Pandey, K.C., 2020b. Structural Basis for Designing Multiepitope Vaccines Against COVID-19 Infection: In Silico Vaccine Design and Validation. JMIR bioinformatics and biotechnology, 1(1), p.e19371.

Srivastava, S., Verma, S., Kamthania, M., Kaur, R., Badyal, R.K., Saxena, A.K., Shin, H.J., Kolbe, M. and Pandey, K., 2020a. Structural basis to design multi-epitope vaccines against Novel Coronavirus 19 (COVID19) infection, the ongoing pandemic emergency: an in silico approach. bioRxiv.

Sturniolo, T., Bono, E., Ding, J., Raddrizzani, L., Tuereci, O., Sahin, U., Braxenthaler, M., Gallazzi, F., Protti, M.P., Sinigaglia, F. and Hammer, J., 1999. Generation of tissue-specific and promiscuous HLA ligand databases using DNA microarrays and virtual HLA class II matrices. Nature biotechnology, 17(6), p.555.

Tenzer S, Peters B, Bulik S, Schoor O, Lemmel C, Schatz MM, Kloetzel PM, Rammensee HG, Schild H, Holzhutter HG. 2005. Modeling the MHC class I pathway by combining predictions of proteasomal cleavage, TAP transport and MHC class I binding. Cell Mol Life Sci 62:1025–1037. (http://tools.iedb.org/processing/)

Totura, A.L., Whitmore, A., Agnihothram, S., Schäfer, A., Katze, M.G., Heise, M.T. and Baric, R.S., 2015. Toll-like receptor 3 signaling via TRIF contributes to a protective innate immune response to severe acute respiratory syndrome coronavirus infection. MBio, 6(3), pp.e00638–15.

Toukmaji, A., Sagui, C., Board, J. and Darden, T., 2000. Efficient particle-mesh Ewald based approach to fixed and induced dipolar interactions. The Journal of chemical physics, 113(24), pp.10913–10927.

ul Qamar, M.T., Rehman, A., Ashfaq, U.A., Awan, M.Q., Fatima, I., Shahid, F. and Chen, L.L., 2020. Designing of a next generation multiepitope based vaccine (MEV) against SARS-COV-2: Immunoinformatics and in silico approaches. BioRxiv.

Vashi, Y., Jagrit, V. and Kumar, S., 2020. Understanding the B and T cells epitopes of spike protein of severe respiratory syndrome coronavirus-2: A computational way to predict the immunogens. bioRxiv.

Wang, P., Sidney, J., Kim, Y., Sette, A., Lund, O., Nielsen, M. and Peters, B., 2010. Peptide binding predictions for HLA DR, DP and DQ molecules. BMC bioinformatics, 11(1), p.568.

Wang, Z. and Xu, J., 2013. Predicting protein contact map using evolutionary and physical constraints by integer programming. Bioinformatics, 29(13), pp.i266–i273 (http://tools.iedb.org/mhcii/).

WHO Coronavirus Disease (COVID-19) Dashboard; 5^th^ September, 2020; https://covid19.who.int/

WHO Weekly Epidemiological Record (WER); 4^th^ September, 2020; https://www.who.int/wer/2020/wer9536/en/;https://apps.who.int/iris/bitstream/handle/10665/334140/WER9536-eng-fre.pdf?ua=1

Wilson, S.S., Wiens, M.E. and Smith, J.G., 2013. Antiviral mechanisms of human defensins. Journal of molecular biology, 425(24), pp.4965–4980.

Wu, X., Wu, S., Li, D., Zhang, J., Hou, L., Ma, J., Liu, W., Ren, D., Zhu, Y. and He, F., 2010. Computational identification of rare codons of Escherichia coli based on codon pairs preference. Bmc Bioinformatics, 11(1), p.61. (https://www.genscript.com/tools/rare-codon-analysis).

Yang, D., Biragyn, A., Kwak, L.W. and Oppenheim, J.J., 2002. Mammalian defensins in immunity: more than just microbicidal. Trends in immunology, 23(6), pp.291–296.

Yarmarkovich, M., Farrel, A., Sison III, A., Di Marco, M., Raman, P., Parris, J.L., Monos, D., Lee, H., Stevanovic, S. and Maris, J.M., 2020. Immunogenicity and Immune Silence in Human Cancer. Frontiers in immunology, 11, p.69.

Yarmarkovich, M., Warrington, J.M., Farrel, A. and Maris, J.M., 2020. Identification of SARS-CoV-2 Vaccine Epitopes Predicted to Induce Long-term Population-Scale Immunity. Cell Reports Medicine.

Yazdani, Z., Rafiei, A., Yazdani, M. and Valadan, R., 2020. Design an efficient multi-epitope peptide vaccine candidate against SARS-CoV-2: An in silico analysis. bioRxiv.

