## Supplementary 1 for "Computationally validated SARS-CoV-2 CTL and HTL Multi-Patch Vaccines designed by reverse epitomics approach, shows potential to cover large ethnically distributed human population worldwide"

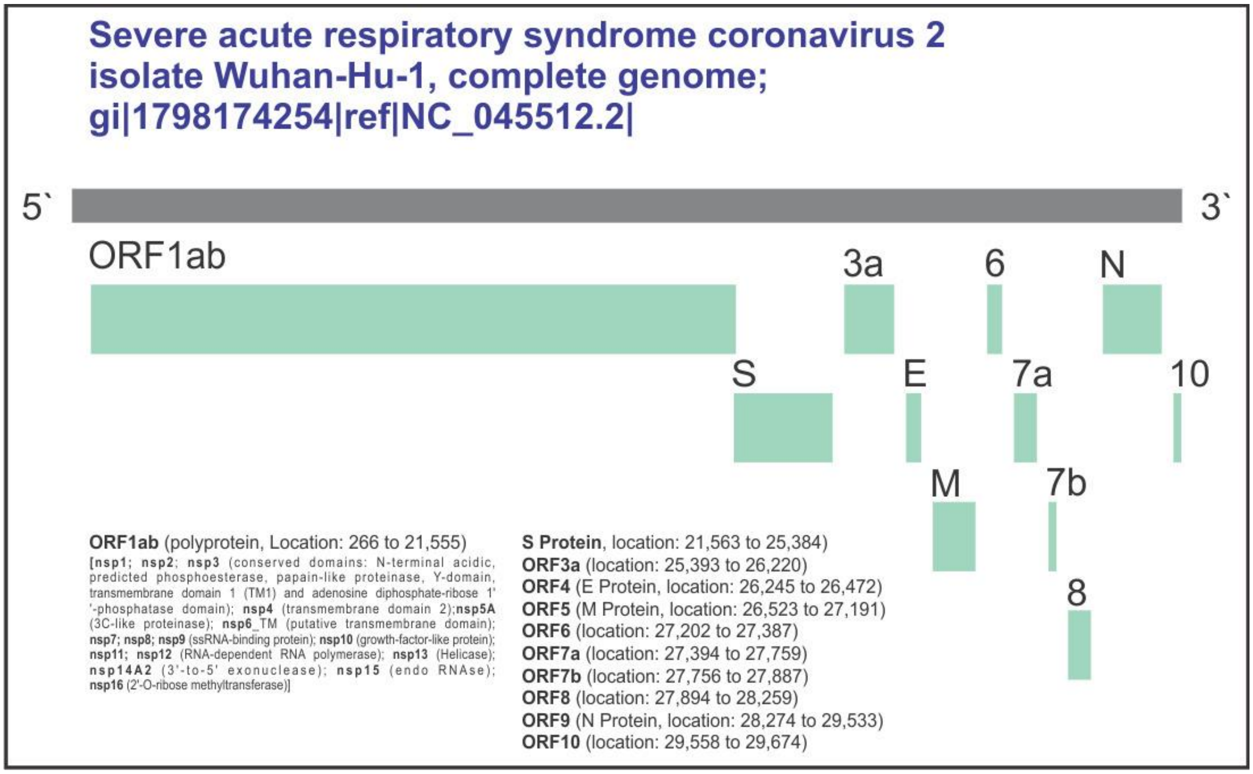

**Supplementary figure S1.** Schematic presentation of all the ORF protein expressed by SARS-CoV-2 genome.

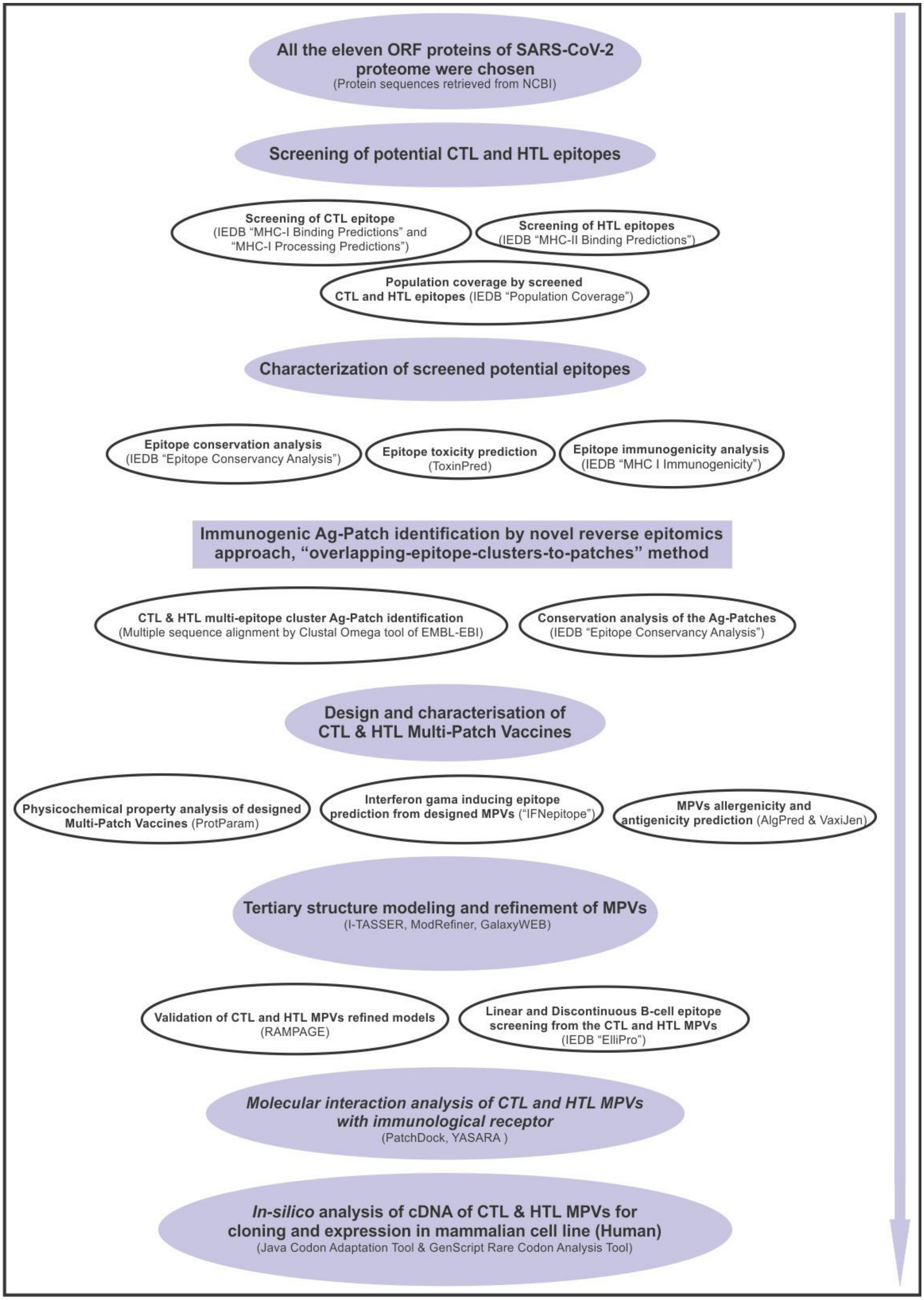

**Supplementary figure S2.** Schematic representation of workflow and methodology.

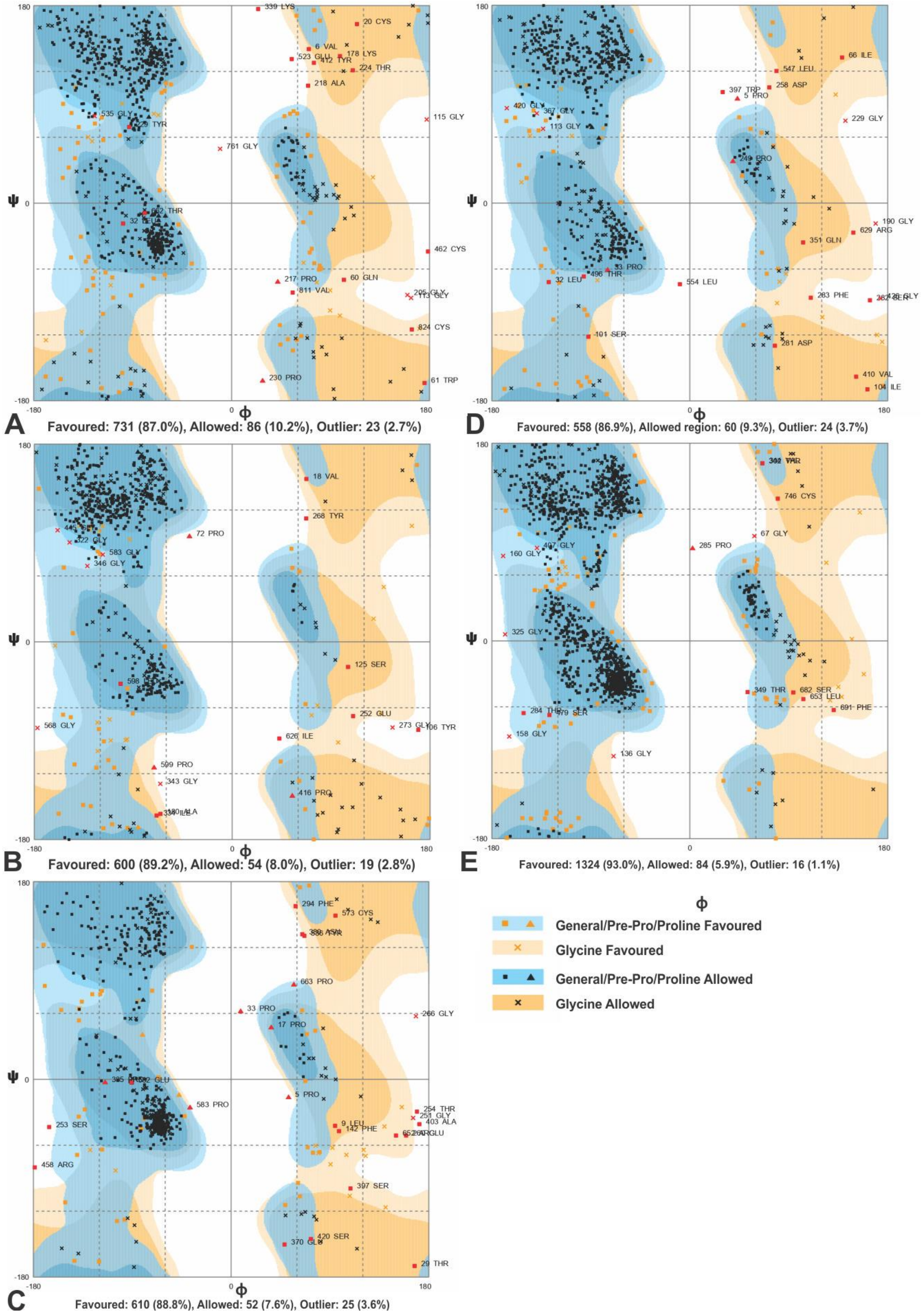

**Supplementary figure S3.** RAMPAGE analysis for all the MPVs **(A)** CTL-MPV-1, **(B)** CTL-MPV-2, **(C)** CTL-MPV-3, **(D)** HTL-MPV-1, **(E)** HTL-MPV-2.

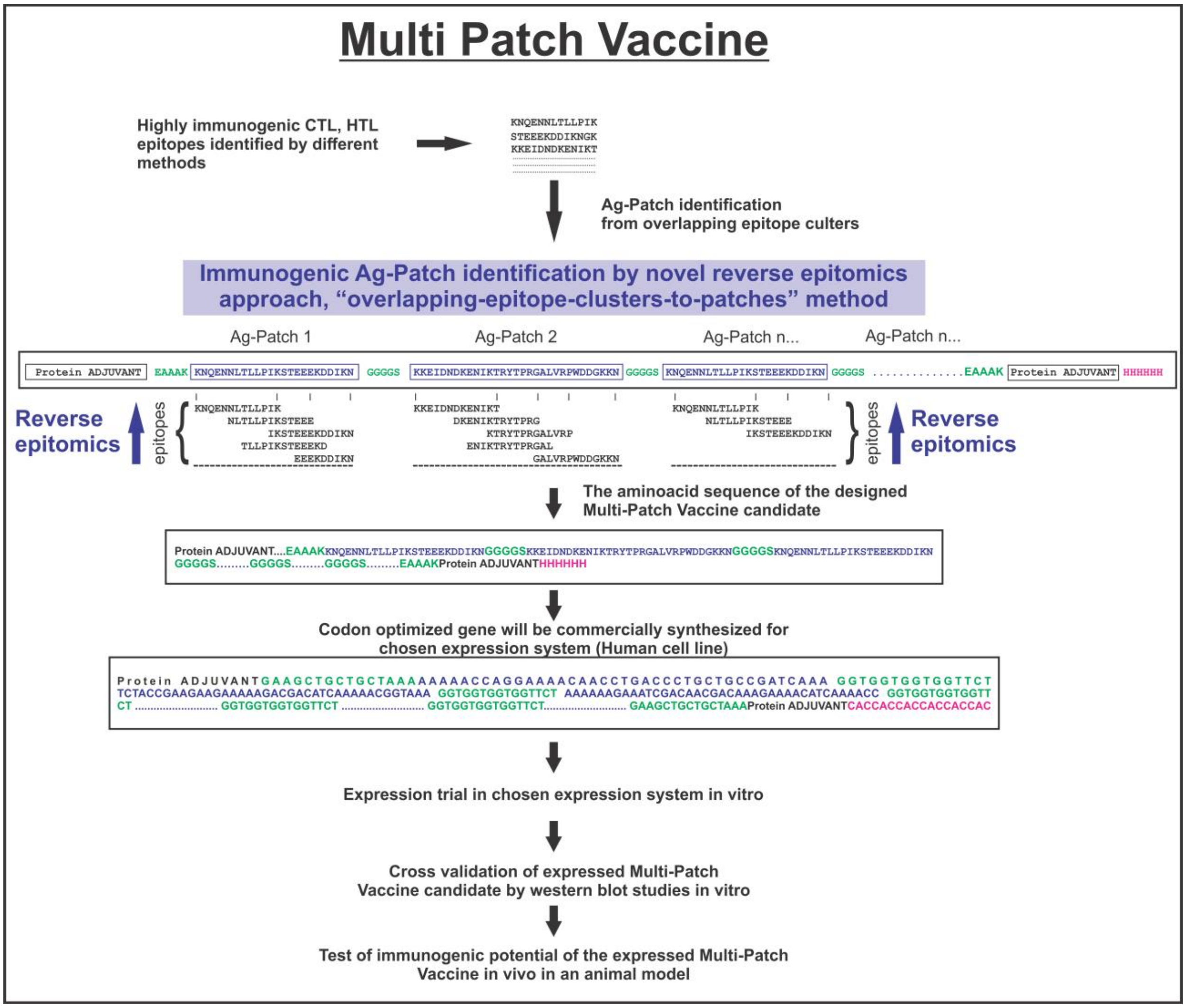

**Supplementary figure S4.** Work flow concept chart from Ag-Patch (antigenic patch) identification to in vivo trial for the proposed MPVs against SARS-CoV-2.

**Supplementary table S1:** High Percentile Ranking CTL epitopes-HLA allele pairs screened from entire proteome of SARS-CoV-2 by the “MHC-I Processing Predictions” tool of IEDB. These epitopes were further utilized to identify the potentially immunogenic multiple epitope cluster based CTL Ag-Patches from the entire proteome of the SRAS-CoV-2. The epitopes shown in **RED** are the epitope which form Overlapping epitope clusters. The screened epitopes are in consensus with the previous studies [Srivastava et al. 2020a; Srivastava et al. 2020b; and Grifoni et al., 2020a].

| S.No. | ORF | Allele | Peptide Length | Peptide | Immunogenicity | Conservancy | Toxicity | Proteasome Score | TAP Score | MHC Score | Processing Score | Total Score | MHC IC50[nM] |
|---|---|---|---|---|---|---|---|---|---|---|---|---|---|
| 1 | E Protein | A*02:01 | 9 | FLAFVVFLL | 0.30188 | 100.00% (35/35) | Non-Toxin | 1.45 | 0.41 | -0.81 | 1.86 | 1.05 | 6.5 |
| 2 | E Protein | A*02:01 | 9 | FLAFVVFLL | 0.30188 | 100.00% (35/35) | Non-Toxin | 1.45 | 0.41 | -0.81 | 1.86 | 1.05 | 6.5 |
| 3 | E Protein | A*02:03 | 9 | FLAFVVFLL | 0.30188 | 100.00% (35/35) | Non-Toxin | 1.45 | 0.41 | -1.17 | 1.86 | 0.69 | 14.9 |
| 4 | E Protein | A*02:06 | 9 | FLAFVVFLL | 0.30188 | 100.00% (35/35) | Non-Toxin | 1.45 | 0.41 | -1.22 | 1.86 | 0.64 | 16.6 |
| 5 | E Protein | A*02:01 | 9 | FLLVTLAIL | 0.17608 | 100.00% (35/35) | Non-Toxin | 1.51 | 0.41 | -1.34 | 1.92 | 0.59 | 21.7 |
| 6 | E Protein | A*02:06 | 9 | FVVFLLVTL | 0.16748 | 100.00% (35/35) | Non-Toxin | 2 | 0.54 | -2.05 | 2.54 | 0.48 | 113.2 |
| 7 | E Protein | B*15:01 | 10 | ILTALRLCAY | 0.05849 | 97.14% (34/35) | Non-Toxin | 1.42 | 1.34 | -1.77 | 2.76 | 0.99 | 58.4 |
| 8 | E Protein | A*30:02 | 10 | ILTALRLCAY | 0.05849 | 97.14% (34/35) | Non-Toxin | 1.42 | 1.34 | -2.24 | 2.76 | 0.52 | 172.9 |
| 9 | E Protein | B*15:01 | 9 | LIVNSVLLF | -0.13119 | 100.00% (35/35) | Non-Toxin | 1.15 | 1.2 | -1.38 | 2.35 | 0.97 | 23.8 |
| 10 | E Protein | B*15:01 | 9 | LLFLAFVVF | 0.2341 | 100.00% (35/35) | Non-Toxin | 1.53 | 1.18 | -2.08 | 2.71 | 0.63 | 120.3 |
| 11 | E Protein | B*15:01 | 9 | LTALRLCAY | 0.01886 | 97.14% (34/35) | Non-Toxin | 1.42 | 1.27 | -1.69 | 2.69 | 1 | 49.5 |
| 12 | E Protein | B*15:01 | 9 | LTALRLCAY | 0.01886 | 97.14% (34/35) | Non-Toxin | 1.42 | 1.27 | -1.69 | 2.69 | 1 | 49.5 |
| 13 | E Protein | A*30:02 | 9 | LTALRLCAY | 0.01886 | 97.14% (34/35) | Non-Toxin | 1.42 | 1.27 | -2.03 | 2.69 | 0.67 | 106 |
| 14 | E Protein | A*01:01 | 9 | LTALRLCAY | 0.01886 | 97.14% (34/35) | Non-Toxin | 1.42 | 1.27 | -2.09 | 2.69 | 0.6 | 123.3 |
| 15 | E Protein | B*15:01 | 9 | LVKPSFYVY | -0.11106 | 100.00% (35/35) | Non-Toxin | 1.51 | 1.35 | -1.42 | 2.86 | 1.44 | 26.3 |
| 16 | E Protein | A*30:02 | 9 | LVKPSFYVY | -0.11106 | 100.00% (35/35) | Non-Toxin | 1.51 | 1.35 | -1.61 | 2.86 | 1.25 | 40.4 |
| 17 | E Protein | B*15:01 | 9 | LVKPSFYVY | -0.11106 | 100.00% (35/35) | Non-Toxin | 1.51 | 1.35 | -1.42 | 2.86 | 1.44 | 26.3 |
| 18 | E Protein | A*30:02 | 9 | LVKPSFYVY | -0.11106 | 100.00% (35/35) | Non-Toxin | 1.51 | 1.35 | -1.61 | 2.86 | 1.25 | 40.4 |
| 19 | E Protein | B*35:01 | 9 | LVKPSFYVY | -0.11106 | 100.00% (35/35) | Non-Toxin | 1.51 | 1.35 | -1.96 | 2.86 | 0.9 | 90.4 |
| 20 | E Protein | A*31:01 | 9 | RVKNLNSSR | -0.32968 | 100.00% (35/35) | Non-Toxin | 0.86 | 0.83 | -0.9 | 1.7 | 0.8 | 7.9 |
| 21 | E Protein | B*15:01 | 10 | SLVKPSFYVY | -0.2443 | 100.00% (35/35) | Non-Toxin | 1.51 | 1.36 | -1.39 | 2.86 | 1.47 | 24.6 |
| 22 | E Protein | B*15:01 | 10 | SLVKPSFYVY | -0.2443 | 100.00% (35/35) | Non-Toxin | 1.51 | 1.36 | -1.39 | 2.86 | 1.47 | 24.6 |
| 23 | E Protein | A*30:02 | 10 | SLVKPSFYVY | -0.2443 | 100.00% (35/35) | Non-Toxin | 1.51 | 1.36 | -1.88 | 2.86 | 0.98 | 75.4 |
| 24 | E Protein | A*11:01 | 10 | SLVKPSFYVY | -0.2443 | 100.00% (35/35) | Non-Toxin | 1.51 | 1.36 | -2.28 | 2.86 | 0.58 | 192.1 |
| 25 | E Protein | A*30:02 | 9 | VSLVKPSFY | -0.25372 | 100.00% (35/35) | Non-Toxin | 1.19 | 1.38 | -1.74 | 2.58 | 0.84 | 54.9 |
| S.No. | ORF | Allele | Peptide Length | Peptide | Immunogenicity | Conservancy | Toxicity | Proteasome Score | TAP Score | MHC Score | Processing Score | Total Score | MHC IC50[nM] |
| 26 | M Protein | A*30:02 | 9 | ATSRTLSYY | -0.11604 | 100.00% (41/41) | Non-Toxin | 1.26 | 1.34 | -1.12 | 2.6 | 1.48 | 13.3 |
| 27 | M Protein | A*11:01 | 9 | ATSRTLSYY | -0.11604 | 100.00% (41/41) | Non-Toxin | 1.26 | 1.34 | -1.52 | 2.6 | 1.08 | 32.9 |
| 28 | M Protein | B*08:01 | 10 | FARTRSMWSF | -0.12986 | 100.00% (41/41) | Non-Toxin | 1.41 | 1.12 | -1.34 | 2.53 | 1.19 | 22.1 |
| 29 | M Protein | A*02:01 | 10 | FLWLLWPVTL | 0.31272 | 100.00% (41/41) | Non-Toxin | 1.85 | 0.46 | -1.16 | 2.31 | 1.16 | 14.3 |
| 30 | M Protein | A*68:01 | 9 | LSYFIASFR | 0.21181 | 100.00% (41/41) | Non-Toxin | 0.84 | 0.72 | -0.46 | 1.56 | 1.1 | 2.9 |
| 31 | M Protein | A*23:01 | 11 | LSYFIASFRLF | 0.2706 | 100.00% (41/41) | Non-Toxin | 1.25 | 1.18 | -1.36 | 2.43 | 1.07 | 23 |
| 32 | M Protein | A*23:01 | 10 | MWLSYFIASF | 0.00197 | 100.00% (41/41) | Non-Toxin | 1.38 | 1.26 | -0.91 | 2.63 | 1.73 | 8.1 |
| 33 | M Protein | A*24:02 | 10 | MWLSYFIASF | 0.00197 | 100.00% (41/41) | Non-Toxin | 1.38 | 1.26 | -1.27 | 2.63 | 1.36 | 18.7 |
| 34 | M Protein | B*15:01 | 10 | MWLSYFIASF | 0.00197 | 100.00% (41/41) | Non-Toxin | 1.38 | 1.26 | -1.33 | 2.63 | 1.3 | 21.6 |
| 35 | M Protein | A*23:01 | 10 | RFLYIIKLIF | 0.11728 | 100.00% (41/41) | Non-Toxin | 1.25 | 1.35 | -1.33 | 2.6 | 1.27 | 21.3 |
| 36 | M Protein | A*30:01 | 8 | RTRSMWSF | -0.25178 | 100.00% (41/41) | Non-Toxin | 1.41 | 1.28 | -1.51 | 2.69 | 1.18 | 32.7 |
| 37 | M Protein | A*30:02 | 9 | SGFAAYSRY | 0.00261 | 100.00% (41/41) | Non-Toxin | 1.53 | 1.17 | -1.4 | 2.69 | 1.3 | 25 |
| 38 | M Protein | B*15:01 | 10 | SQRVAGDSGF | 0.0305 | 100.00% (41/41) | Non-Toxin | 1.33 | 1.24 | -1.35 | 2.58 | 1.23 | 22.4 |
| 39 | M Protein | A*23:01 | 9 | SYFIASFRL | 0.18333 | 100.00% (41/41) | Non-Toxin | 1.45 | 0.62 | -0.99 | 2.08 | 1.09 | 9.8 |
| 40 | M Protein | A*23:01 | 10 | SYFIASFRLF | 0.19632 | 100.00% (41/41) | Non-Toxin | 1.25 | 1.31 | -0.56 | 2.56 | 2 | 3.6 |
| 41 | M Protein | A*24:02 | 10 | SYFIASFRLF | 0.19632 | 100.00% (41/41) | Non-Toxin | 1.25 | 1.31 | -0.76 | 2.56 | 1.8 | 5.7 |
| 42 | M Protein | B*35:01 | 9 | VATSRTLSY | -0.17295 | 100.00% (41/41) | Non-Toxin | 1.34 | 1.31 | -1.38 | 2.65 | 1.27 | 23.9 |

| | | | | | | | | | | | | | |
|---|---|---|---|---|---|---|---|---|---|---|---|---|---|
| 43 | M Protein | B*35:01 | 9 | YANRNRFLY | 0.18472 | 100.00% (41/41) | Non-Toxin | 1.18 | 1.31 | -0.83 | 2.49 | 1.66 | 6.8 |
| 44 | M Protein | A*30:02 | 9 | YANRNRFLY | 0.18472 | 100.00% (41/41) | Non-Toxin | 1.18 | 1.31 | -1.46 | 2.49 | 1.03 | 28.9 |
| 45 | M Protein | A*23:01 | 9 | YFIASFRLF | 0.06887 | 100.00% (41/41) | Non-Toxin | 1.25 | 1.22 | -0.73 | 2.47 | 1.74 | 5.4 |
| 46 | M Protein | A*24:02 | 9 | YFIASFRLF | 0.06887 | 100.00% (41/41) | Non-Toxin | 1.25 | 1.22 | -1 | 2.47 | 1.48 | 9.9 |
| 47 | M Protein | A*30:02 | 9 | YSRYRIGNY | 0.21358 | 100.00% (41/41) | Non-Toxin | 1.36 | 1.37 | -1.61 | 2.73 | 1.12 | 40.6 |
| S.No. | ORF | Allele | Peptide Length | Peptide | Immunogenicity | Conservancy | Toxicity | Proteasome Score | TAP Score | MHC Score | Processing Score | Total Score | MHC IC50[nM] |
| 48 | N Protein | B*15:01 | 10 | AQFAPSASAF | -0.17446 | 100.00% (40/40) | Non-Toxin | 1.23 | 1.25 | -0.7 | 2.48 | 1.78 | 5 |
| 49 | N Protein | B*15:01 | 10 | AQFAPSASAF | -0.17446 | 100.00% (40/40) | Non-Toxin | 1.23 | 1.25 | -0.7 | 2.48 | 1.78 | 5 |
| 50 | N Protein | B*15:01 | 11 | AQFAPSASAFF | -0.11074 | 100.00% (40/40) | Non-Toxin | 1.3 | 1.25 | -1.46 | 2.55 | 1.08 | 29 |
| 51 | N Protein | B*15:01 | 11 | AQFAPSASAFF | -0.11074 | 100.00% (40/40) | Non-Toxin | 1.3 | 1.25 | -1.46 | 2.55 | 1.08 | 29 |
| 52 | N Protein | B*35:01 | 9 | FAPSASAFF | -0.18628 | 100.00% (40/40) | Non-Toxin | 1.3 | 1.05 | -1.54 | 2.35 | 0.81 | 35 |
| 53 | N Protein | A*30:02 | 9 | GTTLPKGFY | -0.11536 | 100.00% (40/40) | Non-Toxin | 1.48 | 1.15 | -1.88 | 2.62 | 0.74 | 76.3 |
| 54 | N Protein | B*15:01 | 11 | IAQFAPSASAF | -0.13353 | 100.00% (40/40) | Non-Toxin | 1.23 | 1.1 | -1.84 | 2.33 | 0.49 | 69.1 |
| 55 | N Protein | A*31:01 | 9 | IGYYRRATR | 0.1499 | 100.00% (40/40) | Non-Toxin | 1.27 | 0.6 | -1.16 | 1.87 | 0.71 | 14.3 |
| 56 | N Protein | A*32:01 | 9 | KAYNVTQAF | -0.00587 | 100.00% (40/40) | Non-Toxin | 1.21 | 1.25 | -0.98 | 2.46 | 1.48 | 9.5 |
| 57 | N Protein | B*15:01 | 9 | KAYNVTQAF | -0.00587 | 100.00% (40/40) | Non-Toxin | 1.21 | 1.25 | -1.34 | 2.46 | 1.12 | 21.9 |
| 58 | N Protein | A*32:01 | 9 | KAYNVTQAF | -0.00587 | 100.00% (40/40) | Non-Toxin | 1.21 | 1.25 | -0.98 | 2.46 | 1.48 | 9.5 |
| 59 | N Protein | B*15:01 | 9 | KAYNVTQAF | -0.00587 | 100.00% (40/40) | Non-Toxin | 1.21 | 1.25 | -1.34 | 2.46 | 1.12 | 21.9 |
| 60 | N Protein | B*58:01 | 9 | KAYNVTQAF | -0.00587 | 100.00% (40/40) | Non-Toxin | 1.21 | 1.25 | -1.48 | 2.46 | 0.98 | 30.1 |
| 61 | N Protein | B*35:01 | 9 | KAYNVTQAF | -0.00587 | 100.00% (40/40) | Non-Toxin | 1.21 | 1.25 | -1.9 | 2.46 | 0.56 | 79.8 |
| 62 | N Protein | A*30:02 | 10 | KDLSPRWYFY | 0.14332 | 100.00% (40/40) | Non-Toxin | 1.58 | 1.21 | -1.68 | 2.79 | 1.1 | 48.3 |
| 63 | N Protein | A*30:02 | 10 | KDLSPRWYFY | 0.14332 | 100.00% (40/40) | Non-Toxin | 1.58 | 1.21 | -1.68 | 2.79 | 1.1 | 48.3 |
| 64 | N Protein | A*03:01 | 10 | KDLSPRWYFY | 0.14332 | 100.00% (40/40) | Non-Toxin | 1.58 | 1.21 | -1.99 | 2.79 | 0.8 | 97.5 |
| 65 | N Protein | A*30:02 | 10 | KMKDLSPRWY | -0.05692 | 100.00% (40/40) | Non-Toxin | 1.31 | 1.35 | -1.56 | 2.66 | 1.1 | 36.2 |
| 66 | N Protein | A*30:02 | 10 | KMKDLSPRWY | -0.05692 | 100.00% (40/40) | Non-Toxin | 1.31 | 1.35 | -1.56 | 2.66 | 1.1 | 36.2 |
| 67 | N Protein | B*15:01 | 10 | KMKDLSPRWY | -0.05692 | 100.00% (40/40) | Non-Toxin | 1.31 | 1.35 | -2.04 | 2.66 | 0.62 | 110.6 |
| 68 | N Protein | A*02:03 | 9 | LLLDRLNQL | -0.01446 | 100.00% (40/40) | Non-Toxin | 1.48 | 0.46 | -0.89 | 1.94 | 1.05 | 7.7 |
| 69 | N Protein | A*02:03 | 9 | LLLDRLNQL | -0.01446 | 100.00% (40/40) | Non-Toxin | 1.48 | 0.46 | -0.89 | 1.94 | 1.05 | 7.7 |
| 70 | N Protein | A*02:01 | 9 | LLLDRLNQL | -0.01446 | 100.00% (40/40) | Non-Toxin | 1.48 | 0.46 | -0.99 | 1.94 | 0.95 | 9.8 |
| 71 | N Protein | A*02:06 | 9 | LLLDRLNQL | -0.01446 | 100.00% (40/40) | Non-Toxin | 1.48 | 0.46 | -1.11 | 1.94 | 0.83 | 12.8 |
| 72 | N Protein | B*15:01 | 9 | LLNKHIDAY | -0.02074 | 100.00% (40/40) | Non-Toxin | 1.31 | 1.24 | -1.58 | 2.55 | 0.98 | 37.6 |
| 73 | N Protein | B*35:01 | 9 | LPAADLDDF | 0.09491 | 100.00% (40/40) | Non-Toxin | 0.94 | 1 | -1.43 | 1.94 | 0.5 | 27.2 |
| 74 | N Protein | B*35:01 | 9 | LPNNTASWF | 0.05582 | 100.00% (40/40) | Non-Toxin | 1.21 | 0.92 | -1.25 | 2.13 | 0.88 | 17.7 |
| 75 | N Protein | B*53:01 | 9 | LPNNTASWF | 0.05582 | 100.00% (40/40) | Non-Toxin | 1.21 | 0.92 | -1.51 | 2.13 | 0.62 | 32.7 |
| 76 | N Protein | A*30:02 | 9 | LSPRWYFYY | 0.35734 | 100.00% (40/40) | Non-Toxin | 1.07 | 1.22 | -1.71 | 2.3 | 0.59 | 50.8 |
| 77 | N Protein | B*44:02 | 9 | MEVTPSGTW | -0.06279 | 100.00% (40/40) | Non-Toxin | 1.44 | 0.37 | -1.26 | 1.8 | 0.54 | 18.2 |
| 78 | N Protein | A*30:02 | 11 | MKDLSPRWYFY | 0.14833 | 100.00% (40/40) | Non-Toxin | 1.58 | 1.28 | -2.31 | 2.86 | 0.55 | 202.5 |
| 79 | N Protein | B*15:01 | 10 | NQRNAPRITF | 0.22131 | 100.00% (40/40) | Non-Toxin | 1.49 | 1.15 | -2.04 | 2.64 | 0.6 | 110.4 |
| 80 | N Protein | A*01:01 | 11 | NSSPDDQIGYY | 0.01726 | 100.00% (40/40) | Non-Toxin | 1.24 | 1.31 | -2.01 | 2.54 | 0.53 | 102.4 |
| 81 | N Protein | A*68:02 | 9 | NTASWFTAL | 0.22775 | 100.00% (40/40) | Non-Toxin | 1.42 | 0.49 | -0.79 | 1.91 | 1.12 | 6.2 |
| 82 | N Protein | A*68:02 | 9 | NTASWFTAL | 0.22775 | 100.00% (40/40) | Non-Toxin | 1.42 | 0.49 | -0.79 | 1.91 | 1.12 | 6.2 |
| 83 | N Protein | A*23:01 | 10 | QFAPSASAFF | -0.17577 | 100.00% (40/40) | Non-Toxin | 1.3 | 1.25 | -1.97 | 2.55 | 0.58 | 92.6 |
| 84 | N Protein | B*15:01 | 10 | RQKRTATKAY | -0.06462 | 100.00% (40/40) | Non-Toxin | 1.51 | 1.34 | -1.55 | 2.85 | 1.3 | 35.5 |
| 85 | N Protein | B*15:01 | 10 | RQKRTATKAY | -0.06462 | 100.00% (40/40) | Non-Toxin | 1.51 | 1.34 | -1.55 | 2.85 | 1.3 | 35.5 |
| 86 | N Protein | A*30:02 | 10 | RQKRTATKAY | -0.06462 | 100.00% (40/40) | Non-Toxin | 1.51 | 1.34 | -2.03 | 2.85 | 0.82 | 107.5 |
| 87 | N Protein | B*35:01 | 9 | SPDDQIGYY | 0.06844 | 100.00% (40/40) | Non-Toxin | 1.24 | 1.11 | -1.8 | 2.35 | 0.55 | 63.2 |
| 88 | N Protein | B*07:02 | 9 | SPRWYFYYL | 0.34101 | 100.00% (40/40) | Non-Toxin | 1.61 | 0.35 | -1.2 | 1.96 | 0.76 | 15.7 |
| 89 | N Protein | B*08:01 | 9 | SPRWYFYYL | 0.34101 | 100.00% (40/40) | Non-Toxin | 1.61 | 0.35 | -1.41 | 1.96 | 0.55 | 25.5 |
| 90 | N Protein | A*01:01 | 10 | SSPDDQIGYY | 0.07924 | 100.00% (40/40) | Non-Toxin | 1.24 | 1.36 | -1.16 | 2.6 | 1.45 | 14.3 |
| 91 | N Protein | A*01:01 | 10 | SSPDDQIGYY | 0.07924 | 100.00% (40/40) | Non-Toxin | 1.24 | 1.36 | -1.16 | 2.6 | 1.45 | 14.3 |
| 92 | N Protein | B*35:01 | 9 | TPSGTWLTY | 0.24003 | 100.00% (40/40) | Non-Toxin | 1.52 | 1.15 | -0.84 | 2.67 | 1.83 | 6.9 |

| | | | | | | | | | | | | | |
|---|---|---|---|---|---|---|---|---|---|---|---|---|---|
| 93 | N Protein | B*35:01 | 9 | TPSGTWLTY | 0.24003 | 100.00% (40/40) | Non-Toxin | 1.52 | 1.15 | -0.84 | 2.67 | 1.83 | 6.9 | |
| 94 | N Protein | B*53:01 | 9 | TPSGTWLTY | 0.24003 | 100.00% (40/40) | Non-Toxin | 1.52 | 1.15 | -2.07 | 2.67 | 0.6 | 117.2 | |
| 95 | N Protein | B*35:01 | 10 | VTPSGTWLTY | 0.12256 | 100.00% (40/40) | Non-Toxin | 1.52 | 1.21 | -1.93 | 2.73 | 0.8 | 84.5 | |
| 96 | N Protein | A*30:02 | 10 | VTPSGTWLTY | 0.12256 | 100.00% (40/40) | Non-Toxin | 1.52 | 1.21 | -2.06 | 2.73 | 0.67 | 115.4 | |
| S.No. | ORF | Allele | Peptide Length | Peptide | Immunogenicity | Conservancy | Toxicity | Proteasome Score | TAP Score | MHC Score | Processing Score | Total Score | MHC IC50[nM] | |
| 97 | ORF10 | A*02:06 | 10 | AQVDVVNFNL | 0.20505 | 100.00% (28/28) | Non-Toxin | 1.4 | 0.55 | -1.35 | 1.95 | 0.6 | 22.6 | |
| 98 | ORF10 | B*35:01 | 8 | FAFPFTIY | 0.2897 | 100.00% (28/28) | Non-Toxin | 1.47 | 1.32 | -1.93 | 2.79 | 0.86 | 85.1 | |
| 99 | ORF10 | A*68:02 | 10 | FAFPFTIYSL | 0.20414 | 100.00% (28/28) | Non-Toxin | 1.38 | 0.47 | -1.39 | 1.85 | 0.46 | 24.5 | |
| 100 | ORF10 | A*68:01 | 10 | FTIYSLLLCR | -0.18372 | 100.00% (28/28) | Non-Toxin | 1.19 | 0.56 | -1.06 | 1.75 | 0.69 | 11.4 | |
| 101 | ORF10 | A*23:01 | 8 | GYINVFAF | 0.22011 | 100.00% (28/28) | Non-Toxin | 1.16 | 1.18 | -1.84 | 2.35 | 0.51 | 68.7 | |
| 102 | ORF10 | A*23:01 | 10 | GYINVFAFPF | 0.32004 | 100.00% (28/28) | Non-Toxin | 1.05 | 1.18 | -0.82 | 2.23 | 1.41 | 6.6 | |
| 103 | ORF10 | A*24:02 | 10 | GYINVFAFPF | 0.32004 | 100.00% (28/28) | Non-Toxin | 1.05 | 1.18 | -1.17 | 2.23 | 1.06 | 14.9 | |
| 104 | ORF10 | A*23:01 | 10 | GYINVFAFPF | 0.32004 | 100.00% (28/28) | Non-Toxin | 1.05 | 1.18 | -0.82 | 2.23 | 1.41 | 6.6 | |
| 105 | ORF10 | A*24:02 | 10 | GYINVFAFPF | 0.32004 | 100.00% (28/28) | Non-Toxin | 1.05 | 1.18 | -1.17 | 2.23 | 1.06 | 14.9 | |
| 106 | ORF10 | B*15:01 | 10 | LLCRMNSRNY | -0.25855 | 100.00% (28/28) | Non-Toxin | 1.23 | 1.31 | -1.93 | 2.53 | 0.6 | 84.8 | |
| 107 | ORF10 | B*35:01 | 9 | MGYINVFAF | 0.28694 | 100.00% (28/28) | Non-Toxin | 1.16 | 1.14 | -1.56 | 2.3 | 0.74 | 36.6 | |
| 108 | ORF10 | A*68:02 | 9 | NVFAFPFTI | 0.30241 | 100.00% (28/28) | Non-Toxin | 1.25 | 0.36 | -0.71 | 1.61 | 0.9 | 5.1 | |
| 109 | ORF10 | A*32:01 | 9 | NVFAFPFTI | 0.30241 | 100.00% (28/28) | Non-Toxin | 1.25 | 0.36 | -1 | 1.61 | 0.61 | 10.1 | |
| 110 | ORF10 | A*02:06 | 9 | NVFAFPFTI | 0.30241 | 100.00% (28/28) | Non-Toxin | 1.25 | 0.36 | -1 | 1.61 | 0.61 | 10.1 | |
| 111 | ORF10 | A*02:01 | 9 | NVFAFPFTI | 0.30241 | 100.00% (28/28) | Non-Toxin | 1.25 | 0.36 | -1.11 | 1.61 | 0.5 | 13 | |
| 112 | ORF10 | A*68:01 | 10 | NVFAFPFTIY | 0.40129 | 100.00% (28/28) | Non-Toxin | 1.47 | 1.4 | -1.84 | 2.86 | 1.02 | 69.7 | |
| 113 | ORF10 | A*68:01 | 10 | NVFAFPFTIY | 0.40129 | 100.00% (28/28) | Non-Toxin | 1.47 | 1.4 | -1.84 | 2.86 | 1.02 | 69.7 | |
| 114 | ORF10 | A*11:01 | 10 | NVFAFPFTIY | 0.40129 | 100.00% (28/28) | Non-Toxin | 1.47 | 1.4 | -2.2 | 2.86 | 0.67 | 157.8 | |
| 115 | ORF10 | A*30:02 | 10 | NVFAFPFTIY | 0.40129 | 100.00% (28/28) | Non-Toxin | 1.47 | 1.4 | -2.29 | 2.86 | 0.57 | 196.8 | |
| 116 | ORF10 | B*15:01 | 10 | NVFAFPFTIY | 0.40129 | 100.00% (28/28) | Non-Toxin | 1.47 | 1.4 | -2.3 | 2.86 | 0.56 | 200.7 | |
| 117 | ORF10 | A*68:01 | 9 | TIYSLLLCR | -0.22977 | 100.00% (28/28) | Non-Toxin | 1.19 | 0.73 | -1.18 | 1.91 | 0.74 | 15.1 | |
| 118 | ORF10 | A*30:02 | 9 | VFAFPFTIY | 0.34042 | 100.00% (28/28) | Non-Toxin | 1.47 | 1.44 | -2.21 | 2.9 | 0.69 | 161.8 | |
| 119 | ORF10 | B*15:01 | 10 | YIAQVDVVNF | 0.02786 | 100.00% (28/28) | Non-Toxin | 1.52 | 1.22 | -1.76 | 2.74 | 0.98 | 57.9 | |
| 120 | ORF10 | B*35:01 | 9 | YINVFAFPF | 0.28259 | 100.00% (28/28) | Non-Toxin | 1.05 | 1.09 | -1.37 | 2.14 | 0.77 | 23.6 | |
| S.No. | ORF | Allele | Peptide Length | Peptide | Immunogenicity | Conservancy | Toxicity | Proteasome Score | TAP Score | MHC Score | Processing Score | Total Score | MHC IC50[nM] | NSP |
| 121 | ORF1ab | A*01:01 | 10 | ACTDDNALAY | 0.10055 | 100.00% (36/36) | Non-Toxin | 1.49 | 1.33 | -0.82 | 2.82 | 2 | 6.6 | nsp9 |
| 122 | ORF1ab | A*01:01 | 11 | ACTDDNALAYY | 0.10874 | 100.00% (36/36) | Non-Toxin | 1.35 | 1.33 | -1.12 | 2.68 | 1.56 | 13.1 | nsp9 |
| 123 | ORF1ab | B*15:01 | 10 | ARLYYDSMSY | -0.38991 | 97.22% (35/36) | Non-Toxin | 1.45 | 1.47 | -1.39 | 2.92 | 1.53 | 24.5 | nsp12 |
| 124 | ORF1ab | B*15:01 | 10 | AVMYMGTLSY | -0.2682 | 100.00% (36/36) | Non-Toxin | 1.3 | 1.35 | -0.95 | 2.64 | 1.7 | 8.9 | nsp3 |
| 125 | ORF1ab | A*23:01 | 9 | AYILFTRFF | 0.29466 | 100.00% (36/36) | Non-Toxin | 1.5 | 1.33 | -1.28 | 2.83 | 1.55 | 19 | nsp3 |
| 126 | ORF1ab | A*23:01 | 10 | AYVNTFSSTF | -0.11398 | 100.00% (36/36) | Non-Toxin | 1.4 | 1.25 | -1.13 | 2.65 | 1.52 | 13.5 | nsp3 |
| 127 | ORF1ab | A*01:01 | 9 | CTDDNALAY | 0.07355 | 100.00% (36/36) | Non-Toxin | 1.49 | 1.23 | -0.45 | 2.72 | 2.27 | 2.8 | nsp9 |
| 128 | ORF1ab | A*01:01 | 10 | CTDDNALAYY | 0.08174 | 100.00% (36/36) | Non-Toxin | 1.35 | 1.23 | -0.57 | 2.58 | 2.01 | 3.7 | nsp9 |
| 129 | ORF1ab | A*01:01 | 9 | DTDFVNEFY | 0.31201 | 97.22% (35/36) | Non-Toxin | 1.55 | 1.07 | -0.91 | 2.62 | 1.71 | 8.1 | nsp12 |
| 130 | ORF1ab | A*26:01 | 9 | ETISLAGSY | -0.1653 | 100.00% (36/36) | Non-Toxin | 1.4 | 1.21 | -0.53 | 2.61 | 2.08 | 3.4 | nsp3 |
| 131 | ORF1ab | B*35:01 | 9 | FAIGLALYY | 0.09181 | 97.22% (35/36) | Non-Toxin | 1.2 | 1.27 | -0.71 | 2.47 | 1.77 | 5.1 | nsp13 |
| 132 | ORF1ab | B*35:01 | 9 | FAVDAAKAY | -0.04849 | 100.00% (36/36) | Non-Toxin | 1.73 | 1.35 | -0.46 | 3.08 | 2.62 | 2.9 | nsp10 |
| 133 | ORF1ab | A*02:03 | 9 | FLNRFTTTL | 0.25596 | 100.00% (36/36) | Non-Toxin | 1.81 | 0.32 | -0.46 | 2.12 | 1.66 | 2.9 | nsp5 |
| 134 | ORF1ab | B*35:01 | 10 | FPLCANGQVF | -0.06779 | 97.22% (35/36) | Non-Toxin | 1.55 | 0.96 | -1 | 2.51 | 1.51 | 10 | nsp13 |
| 135 | ORF1ab | B*35:01 | 9 | FVSLAIDAY | 0.1401 | 97.22% (35/36) | Non-Toxin | 1.26 | 1.24 | -0.96 | 2.5 | 1.53 | 9.2 | nsp12 |
| 136 | ORF1ab | A*01:01 | 9 | GTDLEGNFY | 0.18838 | 100.00% (36/36) | Non-Toxin | 1.42 | 1.17 | -0.99 | 2.59 | 1.61 | 9.7 | nsp5 |
| 137 | ORF1ab | B*15:01 | 9 | ILMTARTVY | 0.12576 | 100.00% (36/36) | Non-Toxin | 1.65 | 1.34 | -1.1 | 2.99 | 1.89 | 12.5 | nsp6 |
| 138 | ORF1ab | B*35:01 | 10 | IPMDSTVKNY | -0.3049 | 97.22% (35/36) | Non-Toxin | 1.53 | 1.16 | -1.18 | 2.7 | 1.52 | 15.1 | nsp15 |
| 139 | ORF1ab | A*01:01 | 9 | ISDYDYYRY | 0.04872 | 97.22% (35/36) | Non-Toxin | 1.49 | 1.29 | -1.06 | 2.77 | 1.71 | 11.5 | nsp12 |
| 140 | ORF1ab | B*58:01 | 9 | KAYKIEELF | 0.08367 | 97.22% (35/36) | Non-Toxin | 1.24 | 1.19 | -0.9 | 2.43 | 1.53 | 7.9 | nsp14 |
| 141 | ORF1ab | A*32:01 | 9 | KLFDRYFKY | 0.08004 | 97.22% (35/36) | Non-Toxin | 1.51 | 1.39 | -1.17 | 2.9 | 1.74 | 14.7 | nsp12 |

| | | | | | | | | | | | | | | |
|---|---|---|---|---|---|---|---|---|---|---|---|---|---|---|
| 142 | ORF1ab | A*30:02 | 9 | KLFDRYFKY | 0.08004 | 97.22% (35/36) | Non-Toxin | 1.51 | 1.39 | -1.27 | 2.9 | 1.64 | 18.5 | nsp12 |
| 143 | ORF1ab | A*03:01 | 9 | KLFDRYFKY | 0.08004 | 97.22% (35/36) | Non-Toxin | 1.51 | 1.39 | -1.31 | 2.9 | 1.6 | 20.2 | nsp12 |
| 144 | ORF1ab | B*15:01 | 10 | LILMTARTVY | 0.0012 | 100.00% (36/36) | Non-Toxin | 1.65 | 1.36 | -1.23 | 3.01 | 1.78 | 16.9 | nsp6 |
| 145 | ORF1ab | B*15:01 | 9 | LMNVLTLVY | 0.07994 | 100.00% (36/36) | Non-Toxin | 1.51 | 1.26 | -1.16 | 2.77 | 1.61 | 14.5 | nsp6 |
| 146 | ORF1ab | B*15:01 | 10 | LMSNLGMPSY | -0.30933 | 100.00% (36/36) | Non-Toxin | 1.55 | 1.34 | -1 | 2.9 | 1.89 | 10.1 | nsp3 |
| 147 | ORF1ab | B*35:01 | 9 | LPGVYSVIY | 0.00581 | 100.00% (36/36) | Non-Toxin | 1.39 | 1.02 | -0.81 | 2.42 | 1.6 | 6.5 | nsp4 |
| 148 | ORF1ab | B*35:01 | 9 | LPSLATVAY | 0.06748 | 100.00% (36/36) | Non-Toxin | 1.47 | 1.15 | -0.34 | 2.62 | 2.28 | 2.2 | nsp6 |
| 149 | ORF1ab | B*35:01 | 9 | LVAEWFLAY | 0.45285 | 100.00% (36/36) | Non-Toxin | 1.21 | 1.33 | -1.01 | 2.54 | 1.52 | 10.3 | nsp3 |
| 150 | ORF1ab | B*35:01 | 10 | LVPFWITIAY | 0.63122 | 100.00% (36/36) | Non-Toxin | 1.42 | 1.26 | -0.91 | 2.68 | 1.77 | 8.1 | nsp4 |
| 151 | ORF1ab | B*15:01 | 9 | MMSAPPAQY | -0.07023 | 100.00% (36/36) | Non-Toxin | 1.27 | 1.36 | -1.11 | 2.63 | 1.52 | 12.9 | nsp3 |
| 152 | ORF1ab | B*35:01 | 9 | MSNLGMPSY | -0.25158 | 100.00% (36/36) | Non-Toxin | 1.55 | 1.29 | -1.28 | 2.84 | 1.56 | 18.9 | nsp3 |
| 153 | ORF1ab | B*15:01 | 10 | MVMFTPLVPF | 0.10572 | 100.00% (36/36) | Non-Toxin | 1.2 | 1.18 | -0.74 | 2.38 | 1.64 | 5.5 | nsp4 |
| 154 | ORF1ab | A*23:01 | 11 | MYMGTLSYEQF | -0.14366 | 100.00% (36/36) | Non-Toxin | 1.36 | 1.28 | -0.96 | 2.65 | 1.68 | 9.2 | nsp3 |
| 155 | ORF1ab | B*44:03 | 10 | QEYADVFHLY | 0.23099 | 97.22% (35/36) | Non-Toxin | 1.22 | 1.36 | -1.01 | 2.57 | 1.57 | 10.2 | nsp12 |
| 156 | ORF1ab | B*35:01 | 9 | QVVDMSMTY | -0.41653 | 100.00% (36/36) | Non-Toxin | 1.49 | 1.37 | -1.03 | 2.87 | 1.84 | 10.6 | nsp3 |
| 157 | ORF1ab | B*15:01 | 9 | RLYYDSMSY | -0.38391 | 97.22% (35/36) | Non-Toxin | 1.45 | 1.53 | -1 | 2.98 | 1.97 | 10.1 | nsp12 |
| 158 | ORF1ab | A*03:01 | 9 | RLYYDSMSY | -0.38391 | 97.22% (35/36) | Non-Toxin | 1.45 | 1.53 | -1.02 | 2.98 | 1.96 | 10.4 | nsp12 |
| 159 | ORF1ab | A*30:02 | 9 | RLYYDSMSY | -0.38391 | 97.22% (35/36) | Non-Toxin | 1.45 | 1.53 | -1.14 | 2.98 | 1.84 | 13.9 | nsp12 |
| 160 | ORF1ab | A*32:01 | 9 | RLYYDSMSY | -0.38391 | 97.22% (35/36) | Non-Toxin | 1.45 | 1.53 | -1.16 | 2.98 | 1.82 | 14.4 | nsp12 |
| 161 | ORF1ab | A*32:01 | 9 | RMYIFFASF | 0.29328 | 100.00% (36/36) | Non-Toxin | 1.32 | 1.36 | -0.82 | 2.68 | 1.86 | 6.6 | nsp3 |
| 162 | ORF1ab | A*30:02 | 10 | RMYIFFASFY | 0.32633 | 100.00% (36/36) | Non-Toxin | 1.28 | 1.53 | -0.97 | 2.81 | 1.83 | 9.4 | nsp3 |
| 163 | ORF1ab | A*03:01 | 10 | RMYIFFASFY | 0.32633 | 100.00% (36/36) | Non-Toxin | 1.28 | 1.53 | -1 | 2.81 | 1.8 | 10.1 | nsp3 |
| 164 | ORF1ab | B*15:01 | 10 | RTIAFGGCVF | 0.24009 | 100.00% (36/36) | Non-Toxin | 1.51 | 1.21 | -1.17 | 2.73 | 1.56 | 14.7 | nsp2 |
| 165 | ORF1ab | A*30:02 | 10 | RYFKYWDQTY | -0.0211 | 97.22% (35/36) | Non-Toxin | 1.49 | 1.48 | -1.42 | 2.97 | 1.54 | 26.6 | nsp12 |
| 166 | ORF1ab | A*30:02 | 10 | RYFRLTLGVY | 0.15936 | 100.00% (36/36) | Non-Toxin | 1.41 | 1.6 | -1.34 | 3.01 | 1.66 | 22.1 | nsp6 |
| 167 | ORF1ab | B*44:03 | 9 | SEFSSLPSY | -0.40603 | 100.00% (36/36) | Non-Toxin | 1.49 | 1.35 | -1.03 | 2.84 | 1.82 | 10.6 | nsp8 |
| 168 | ORF1ab | A*23:01 | 10 | SYFVVKRHTF | 0.00844 | 97.22% (35/36) | Non-Toxin | 1.5 | 1.23 | -1.02 | 2.73 | 1.71 | 10.4 | nsp12 |
| 169 | ORF1ab | A*01:01 | 11 | TACTDDNALAY | 0.10879 | 100.00% (36/36) | Non-Toxin | 1.49 | 1.27 | -1.19 | 2.75 | 1.56 | 15.6 | nsp9 |
| 170 | ORF1ab | A*02:01 | 9 | TMADLVYAL | 0.08282 | 97.22% (35/36) | Non-Toxin | 1.73 | 0.44 | -0.56 | 2.17 | 1.62 | 3.6 | nsp12 |
| 171 | ORF1ab | A*02:03 | 9 | TMADLVYAL | 0.08282 | 97.22% (35/36) | Non-Toxin | 1.73 | 0.44 | -0.64 | 2.17 | 1.53 | 4.4 | nsp12 |
| 172 | ORF1ab | A*02:06 | 9 | TMADLVYAL | 0.08282 | 97.22% (35/36) | Non-Toxin | 1.73 | 0.44 | -0.67 | 2.17 | 1.5 | 4.7 | nsp12 |
| 173 | ORF1ab | B*15:01 | 10 | TQVVDMSMTY | -0.37571 | 100.00% (36/36) | Non-Toxin | 1.49 | 1.32 | -1.25 | 2.81 | 1.56 | 17.6 | nsp3 |
| 174 | ORF1ab | A*01:01 | 10 | TTDPSFLGRY | -0.00266 | 100.00% (36/36) | Non-Toxin | 1.29 | 1.16 | -0.83 | 2.46 | 1.63 | 6.7 | nsp3 |
| 175 | ORF1ab | B*35:01 | 9 | TVAYFNMVY | -0.00719 | 100.00% (36/36) | Non-Toxin | 1.52 | 1.39 | -1.23 | 2.92 | 1.69 | 16.9 | nsp6 |
| 176 | ORF1ab | A*01:01 | 10 | VDTDFVNEFY | 0.33593 | 97.22% (35/36) | Non-Toxin | 1.55 | 1.09 | -0.9 | 2.64 | 1.74 | 7.9 | nsp12 |
| 177 | ORF1ab | B*15:01 | 10 | VMFLARGIVF | 0.28368 | 100.00% (36/36) | Non-Toxin | 1.57 | 1.23 | -1.09 | 2.8 | 1.71 | 12.4 | nsp6 |
| 178 | ORF1ab | B*15:01 | 9 | VMFTPLVPF | 0.08418 | 100.00% (36/36) | Non-Toxin | 1.2 | 1.24 | -0.92 | 2.44 | 1.52 | 8.4 | nsp4 |
| 179 | ORF1ab | B*15:01 | 9 | VMYMGTLSY | -0.21438 | 100.00% (36/36) | Non-Toxin | 1.3 | 1.5 | -0.8 | 2.79 | 1.99 | 6.3 | nsp3 |
| 180 | ORF1ab | A*03:01 | 9 | VMYMGTLSY | -0.21438 | 100.00% (36/36) | Non-Toxin | 1.3 | 1.5 | -0.98 | 2.79 | 1.82 | 9.5 | nsp3 |
| 181 | ORF1ab | A*30:02 | 9 | VMYMGTLSY | -0.21438 | 100.00% (36/36) | Non-Toxin | 1.3 | 1.5 | -1.09 | 2.79 | 1.7 | 12.4 | nsp3 |
| 182 | ORF1ab | B*35:01 | 9 | VPFWITIAY | 0.56221 | 100.00% (36/36) | Non-Toxin | 1.42 | 1.23 | -0.51 | 2.65 | 2.14 | 3.2 | nsp4 |
| 183 | ORF1ab | B*35:01 | 9 | VPWDTIANY | 0.28654 | 100.00% (36/36) | Non-Toxin | 1.37 | 1.26 | -1.06 | 2.63 | 1.57 | 11.4 | nsp3 |
| 184 | ORF1ab | B*15:01 | 9 | VVYRGTTTY | 0.17586 | 97.22% (35/36) | Non-Toxin | 1.54 | 1.51 | -1.41 | 3.05 | 1.64 | 25.5 | nsp13 |
| 185 | ORF1ab | A*30:02 | 9 | VVYRGTTTY | 0.17586 | 97.22% (35/36) | Non-Toxin | 1.54 | 1.51 | -1.43 | 3.05 | 1.62 | 27 | nsp13 |
| 186 | ORF1ab | A*23:01 | 9 | WSMATYYLF | 0.00709 | 97.22% (35/36) | Non-Toxin | 1.17 | 1.06 | -0.73 | 2.23 | 1.5 | 5.4 | nsp3 |
| 187 | ORF1ab | B*35:01 | 8 | YAFEHIVY | 0.33037 | 97.22% (35/36) | Non-Toxin | 1.68 | 1.32 | -1.49 | 3 | 1.51 | 30.9 | nsp15 |
| 188 | ORF1ab | B*08:01 | 8 | YLKRRVVF | 0.10618 | 100.00% (36/36) | Non-Toxin | 1.62 | 1.16 | -1.2 | 2.78 | 1.58 | 15.7 | nsp4 |
| 189 | ORF1ab | B*35:01 | 9 | YPNASFDNF | 0.00131 | 100.00% (36/36) | Non-Toxin | 1.27 | 0.86 | -0.56 | 2.12 | 1.56 | 3.6 | nsp3 |
| 190 | ORF1ab | A*02:01 | 10 | YTMADLVYAL | 0.05177 | 97.22% (35/36) | Non-Toxin | 1.73 | 0.47 | -0.58 | 2.2 | 1.62 | 3.8 | nsp12 |
| 191 | ORF1ab | A*02:06 | 10 | YTMADLVYAL | 0.05177 | 97.22% (35/36) | Non-Toxin | 1.73 | 0.47 | -0.63 | 2.2 | 1.57 | 4.3 | nsp12 |
| 192 | ORF1ab | A*02:03 | 10 | YTMADLVYAL | 0.05177 | 97.22% (35/36) | Non-Toxin | 1.73 | 0.47 | -0.67 | 2.2 | 1.53 | 4.7 | nsp12 |

| | | | | | | | | | | | | | | |
|---|---|---|---|---|---|---|---|---|---|---|---|---|---|---|
| 193 | ORF1ab | B*35:01 | 9 | YVMHANYIF | 0.0822 | 97.22% (35/36) | Non-Toxin | 1.37 | 1.17 | -1.03 | 2.53 | 1.5 | 10.8 | nsp16 |
| 194 | ORF1ab | B*35:01 | 9 | YVNTFSSTF | -0.12171 | 100.00% (36/36) | Non-Toxin | 1.4 | 1.18 | -1.01 | 2.58 | 1.57 | 10.2 | nsp3 |
| 195 | ORF1ab | B*15:01 | 9 | YVNTFSSTF | -0.12171 | 100.00% (36/36) | Non-Toxin | 1.4 | 1.18 | -1.07 | 2.58 | 1.5 | 11.8 | nsp3 |
| 196 | ORF1ab | A*23:01 | 10 | YYFMRFRRAF | 0.14096 | 100.00% (36/36) | Non-Toxin | 1.34 | 1.32 | -0.95 | 2.66 | 1.7 | 9 | nsp4 |
| S.No. | ORF | Allele | Peptide Length | Peptide | Immunogenicity | Conservancy | Toxicity | Proteasome Score | TAP Score | MHC Score | Processing Score | Total Score | MHC IC50[nM] | |
| 197 | ORF3a | B*44:03 | 11 | AGLEAPFLYLY | 0.21841 | 100.00% (28/28) | Non-Toxin | 1.51 | 1.29 | -1.63 | 2.8 | 1.17 | 42.9 | |
| 198 | ORF3a | B*44:03 | 11 | AGLEAPFLYLY | 0.21841 | 100.00% (28/28) | Non-Toxin | 1.51 | 1.29 | -1.63 | 2.8 | 1.17 | 42.9 | |
| 199 | ORF3a | B*44:02 | 11 | AGLEAPFLYLY | 0.21841 | 100.00% (28/28) | Non-Toxin | 1.51 | 1.29 | -1.94 | 2.8 | 0.86 | 86.3 | |
| 200 | ORF3a | A*02:03 | 9 | ALSKGVHFV | -0.10314 | 100.00% (28/28) | Non-Toxin | 1.07 | 0.23 | -0.57 | 1.3 | 0.73 | 3.7 | |
| 201 | ORF3a | A*02:03 | 9 | ALVYFLQSI | -0.08118 | 100.00% (28/28) | Non-Toxin | 1.25 | 0.31 | -0.83 | 1.56 | 0.73 | 6.8 | |
| 202 | ORF3a | A*26:01 | 10 | DTGVEHVTFF | 0.32156 | 100.00% (28/28) | Non-Toxin | 1.24 | 0.89 | -1.64 | 2.13 | 0.49 | 43.5 | |
| 203 | ORF3a | B*15:01 | 9 | FLYLYALVY | 0.03563 | 100.00% (28/28) | Non-Toxin | 1.58 | 1.26 | -2.26 | 2.83 | 0.58 | 181.1 | |
| 204 | ORF3a | A*23:01 | 10 | FLYLYALVYF | 0.04438 | 100.00% (28/28) | Non-Toxin | 1.18 | 1.09 | -1.79 | 2.27 | 0.48 | 61.1 | |
| 205 | ORF3a | B*44:03 | 10 | GLEAPFLYLY | 0.15503 | 100.00% (28/28) | Non-Toxin | 1.51 | 1.2 | -1.26 | 2.71 | 1.46 | 18.1 | |
| 206 | ORF3a | B*44:02 | 10 | GLEAPFLYLY | 0.15503 | 100.00% (28/28) | Non-Toxin | 1.51 | 1.2 | -1.61 | 2.71 | 1.1 | 40.7 | |
| 207 | ORF3a | B*44:03 | 10 | GLEAPFLYLY | 0.15503 | 100.00% (28/28) | Non-Toxin | 1.51 | 1.2 | -1.26 | 2.71 | 1.46 | 18.1 | |
| 208 | ORF3a | B*44:02 | 10 | GLEAPFLYLY | 0.15503 | 100.00% (28/28) | Non-Toxin | 1.51 | 1.2 | -1.61 | 2.71 | 1.1 | 40.7 | |
| 209 | ORF3a | A*23:01 | 10 | HFVCNLLLLF | -0.08423 | 100.00% (28/28) | Non-Toxin | 1.34 | 1.19 | -1.75 | 2.52 | 0.77 | 56.3 | |
| 210 | ORF3a | B*35:01 | 9 | IPIQASLPF | -0.20683 | 100.00% (28/28) | Non-Toxin | 0.98 | 1.01 | -0.43 | 1.99 | 1.56 | 2.7 | |
| 211 | ORF3a | B*35:01 | 9 | IPIQASLPF | -0.20683 | 100.00% (28/28) | Non-Toxin | 0.98 | 1.01 | -0.43 | 1.99 | 1.56 | 2.7 | |
| 212 | ORF3a | B*53:01 | 9 | IPIQASLPF | -0.20683 | 100.00% (28/28) | Non-Toxin | 0.98 | 1.01 | -1.28 | 1.99 | 0.71 | 19.2 | |
| 213 | ORF3a | B*07:02 | 9 | IPIQASLPF | -0.20683 | 100.00% (28/28) | Non-Toxin | 0.98 | 1.01 | -1.39 | 1.99 | 0.6 | 24.7 | |
| 214 | ORF3a | B*15:01 | 10 | IVGVALLAVF | 0.12654 | 100.00% (28/28) | Non-Toxin | 1.3 | 1.12 | -1.76 | 2.42 | 0.66 | 57.1 | |
| 215 | ORF3a | B*44:03 | 9 | LEAPFLYLY | 0.0955 | 100.00% (28/28) | Non-Toxin | 1.51 | 1.21 | -1.3 | 2.72 | 1.41 | 20.1 | |
| 216 | ORF3a | B*44:03 | 9 | LEAPFLYLY | 0.0955 | 100.00% (28/28) | Non-Toxin | 1.51 | 1.21 | -1.3 | 2.72 | 1.41 | 20.1 | |
| 217 | ORF3a | B*44:02 | 9 | LEAPFLYLY | 0.0955 | 100.00% (28/28) | Non-Toxin | 1.51 | 1.21 | -1.73 | 2.72 | 0.99 | 53.3 | |
| 218 | ORF3a | B*15:01 | 11 | LLVAAGLEAPF | 0.23386 | 100.00% (28/28) | Non-Toxin | 0.86 | 1.16 | -1.49 | 2.02 | 0.54 | 30.8 | |
| 219 | ORF3a | B*15:01 | 10 | LVAAGLEAPF | 0.19506 | 100.00% (28/28) | Non-Toxin | 0.86 | 1.19 | -1.1 | 2.05 | 0.95 | 12.5 | |
| 220 | ORF3a | A*23:01 | 10 | LVYFLQSINF | -0.05419 | 100.00% (28/28) | Non-Toxin | 1.3 | 1.31 | -1.39 | 2.61 | 1.23 | 24.3 | |
| 221 | ORF3a | A*23:01 | 10 | LVYFLQSINF | -0.05419 | 100.00% (28/28) | Non-Toxin | 1.3 | 1.31 | -1.39 | 2.61 | 1.23 | 24.3 | |
| 222 | ORF3a | A*24:02 | 10 | LVYFLQSINF | -0.05419 | 100.00% (28/28) | Non-Toxin | 1.3 | 1.31 | -1.84 | 2.61 | 0.77 | 69 | |
| 223 | ORF3a | B*15:01 | 10 | LVYFLQSINF | -0.05419 | 100.00% (28/28) | Non-Toxin | 1.3 | 1.31 | -1.93 | 2.61 | 0.68 | 85.6 | |
| 224 | ORF3a | A*23:01 | 9 | LYLYALVYF | 0.05302 | 100.00% (28/28) | Non-Toxin | 1.18 | 1.18 | -1.29 | 2.36 | 1.07 | 19.6 | |
| 225 | ORF3a | A*23:01 | 9 | LYLYALVYF | 0.05302 | 100.00% (28/28) | Non-Toxin | 1.18 | 1.18 | -1.29 | 2.36 | 1.07 | 19.6 | |
| 226 | ORF3a | A*24:02 | 9 | LYLYALVYF | 0.05302 | 100.00% (28/28) | Non-Toxin | 1.18 | 1.18 | -1.86 | 2.36 | 0.5 | 71.7 | |
| 227 | ORF3a | A*02:01 | 10 | LYLYALVYFL | 0.12412 | 100.00% (28/28) | Non-Toxin | 1.45 | 0.5 | -1 | 1.94 | 0.94 | 10.1 | |
| 228 | ORF3a | A*32:01 | 10 | RIFTIGTVTL | 0.33372 | 100.00% (28/28) | Non-Toxin | 1.77 | 0.64 | -1.59 | 2.41 | 0.82 | 38.6 | |
| 229 | ORF3a | A*31:01 | 9 | RLWLCWKCR | 0.00325 | 100.00% (28/28) | Non-Toxin | 1.27 | 0.85 | -1.25 | 2.11 | 0.87 | 17.7 | |
| 230 | ORF3a | B*15:01 | 10 | TIPIQASLPF | -0.11595 | 100.00% (28/28) | Non-Toxin | 0.98 | 1.21 | -1.31 | 2.18 | 0.87 | 20.6 | |
| 231 | ORF3a | B*35:01 | 10 | TIPIQASLPF | -0.11595 | 100.00% (28/28) | Non-Toxin | 0.98 | 1.21 | -1.39 | 2.18 | 0.79 | 24.8 | |
| 232 | ORF3a | A*02:03 | 9 | TVYSHLLLV | -0.16245 | 100.00% (28/28) | Non-Toxin | 1.17 | 0.29 | -0.79 | 1.45 | 0.66 | 6.2 | |
| 233 | ORF3a | A*02:06 | 9 | TVYSHLLLV | -0.16245 | 100.00% (28/28) | Non-Toxin | 1.17 | 0.29 | -0.91 | 1.45 | 0.54 | 8.2 | |
| 234 | ORF3a | B*35:01 | 9 | VAAGLEAPF | 0.15679 | 100.00% (28/28) | Non-Toxin | 0.86 | 1.18 | -1.2 | 2.04 | 0.84 | 15.8 | |
| 235 | ORF3a | B*44:03 | 9 | VEHVTFFIY | 0.3766 | 100.00% (28/28) | Non-Toxin | 1.28 | 1.24 | -1.9 | 2.52 | 0.62 | 78.7 | |
| 236 | ORF3a | A*30:02 | 10 | VVNPVMEPIY | 0.01859 | 100.00% (28/28) | Non-Toxin | 1.25 | 1.3 | -1.92 | 2.55 | 0.63 | 82.7 | |
| 237 | ORF3a | A*23:01 | 9 | VYFLQSINF | -0.13315 | 100.00% (28/28) | Non-Toxin | 1.3 | 1.31 | -0.96 | 2.61 | 1.65 | 9.2 | |
| 238 | ORF3a | A*24:02 | 9 | VYFLQSINF | -0.13315 | 100.00% (28/28) | Non-Toxin | 1.3 | 1.31 | -1.28 | 2.61 | 1.33 | 19.2 | |
| 239 | ORF3a | A*23:01 | 9 | VYFLQSINF | -0.13315 | 100.00% (28/28) | Non-Toxin | 1.3 | 1.31 | -0.96 | 2.61 | 1.65 | 9.2 | |
| 240 | ORF3a | A*24:02 | 9 | VYFLQSINF | -0.13315 | 100.00% (28/28) | Non-Toxin | 1.3 | 1.31 | -1.28 | 2.61 | 1.33 | 19.2 | |
| 241 | ORF3a | A*33:01 | 10 | YFLQSINFVR | -0.03483 | 100.00% (28/28) | Non-Toxin | 1.09 | 0.74 | -1.27 | 1.82 | 0.56 | 18.5 | |
| 242 | ORF3a | A*02:01 | 9 | YLYALVYFL | 0.13151 | 100.00% (28/28) | Non-Toxin | 1.45 | 0.52 | -0.38 | 1.96 | 1.58 | 2.4 | |

| | | | | | | | | | | | | | |
|---|---|---|---|---|---|---|---|---|---|---|---|---|---|
| 243 | ORF3a | A*02:06 | 9 | YLYALVYFL | 0.13151 | 100.00% (28/28) | Non-Toxin | 1.45 | 0.52 | -0.61 | 1.96 | 1.35 | 4.1 |
| 244 | ORF3a | A*02:03 | 9 | YLYALVYFL | 0.13151 | 100.00% (28/28) | Non-Toxin | 1.45 | 0.52 | -0.76 | 1.96 | 1.21 | 5.7 |
| 245 | ORF3a | A*02:01 | 9 | YLYALVYFL | 0.13151 | 100.00% (28/28) | Non-Toxin | 1.45 | 0.52 | -0.38 | 1.96 | 1.58 | 2.4 |
| 246 | ORF3a | A*02:06 | 9 | YLYALVYFL | 0.13151 | 100.00% (28/28) | Non-Toxin | 1.45 | 0.52 | -0.61 | 1.96 | 1.35 | 4.1 |
| 247 | ORF3a | A*02:03 | 9 | YLYALVYFL | 0.13151 | 100.00% (28/28) | Non-Toxin | 1.45 | 0.52 | -0.76 | 1.96 | 1.21 | 5.7 |
| 248 | ORF3a | A*24:02 | 9 | YYQLYSTQL | -0.24301 | 100.00% (28/28) | Non-Toxin | 1.66 | 0.4 | -1.56 | 2.06 | 0.5 | 36.4 |
| 249 | ORF3a | A*23:01 | 9 | YYQLYSTQL | -0.24301 | 100.00% (28/28) | Non-Toxin | 1.66 | 0.4 | -1.58 | 2.06 | 0.48 | 38.3 |
| S.No. | ORF | Allele | Peptide Length | Peptide | Immunogenicity | Conservancy | Toxicity | Proteasome Score | TAP Score | MHC Score | Processing Score | Total Score | MHC IC50[nM] |
| 250 | ORF6 | A*02:01 | 10 | FHLVDFQVTI | 0.12202 | 100.00% (34/34) | Non-Toxin | 1.57 | 0.22 | -1.16 | 1.78 | 0.62 | 14.5 |
| 251 | ORF6 | A*02:03 | 10 | FHLVDFQVTI | 0.12202 | 100.00% (34/34) | Non-Toxin | 1.57 | 0.22 | -1.17 | 1.78 | 0.61 | 14.9 |
| 252 | ORF6 | A*02:06 | 9 | FQVTIAEIL | 0.38115 | 100.00% (34/34) | Non-Toxin | 1.59 | 0.38 | -1.58 | 1.97 | 0.39 | 38.1 |
| 253 | ORF6 | A*02:03 | 9 | HLVDFQVTI | 0.0982 | 100.00% (34/34) | Non-Toxin | 1.57 | 0.19 | -1.22 | 1.76 | 0.53 | 16.7 |
| 254 | ORF6 | A*02:01 | 9 | HLVDFQVTI | 0.0982 | 100.00% (34/34) | Non-Toxin | 1.57 | 0.19 | -1.25 | 1.76 | 0.51 | 17.7 |
| 255 | ORF6 | A*30:02 | 9 | KVSIWNLDY | 0.29343 | 100.00% (34/34) | | 1.21 | 1.33 | -1.46 | 2.54 | 1.08 | 28.8 |
| 256 | ORF6 | A*30:02 | 9 | KVSIWNLDY | 0.29343 | 100.00% (34/34) | Non-Toxin | 1.21 | 1.33 | -1.46 | 2.54 | 1.08 | 28.8 |
| 257 | ORF6 | A*30:02 | 10 | LSKSLTENKY | -0.24668 | 100.00% (34/34) | Non-Toxin | 1.52 | 1.35 | -2.37 | 2.87 | 0.5 | 234.3 |
| 258 | ORF6 | B*58:01 | 8 | RTFKVSIW | -0.18221 | 100.00% (34/34) | Non-Toxin | 1.42 | 0.54 | -1.34 | 1.97 | 0.63 | 21.7 |
| 259 | ORF6 | B*57:01 | 8 | RTFKVSIW | -0.18221 | 100.00% (34/34) | Non-Toxin | 1.42 | 0.54 | -1.38 | 1.97 | 0.59 | 24 |
| S.No. | ORF | Allele | Peptide Length | Peptide | Immunogenicity | Conservancy | Toxicity | Proteasome Score | TAP Score | MHC Score | Processing Score | Total Score | MHC IC50[nM] |
| 260 | ORF7a | A*02:03 | 10 | ALITLATCEL | 0.1591 | 100.00% (28/28) | Non-Toxin | 1.69 | 0.54 | -1.66 | 2.23 | 0.57 | 45.8 |
| 261 | ORF7a | B*15:01 | 9 | ALTCFSTQF | -0.1183 | 100.00% (28/28) | Non-Toxin | 1.46 | 1.11 | -1.91 | 2.57 | 0.66 | 82.2 |
| 262 | ORF7a | B*35:01 | 9 | CPDGVKHVY | -0.07008 | 100.00% (28/28) | | 1.43 | 1.11 | -1.34 | 2.54 | 1.2 | 22 |
| 263 | ORF7a | B*35:01 | 9 | CPDGVKHVY | -0.07008 | 100.00% (28/28) | Non-Toxin | 1.43 | 1.11 | -1.34 | 2.54 | 1.2 | 22 |
| 264 | ORF7a | B*35:01 | 11 | FACPDGVKHVY | -0.08544 | 100.00% (28/28) | Non-Toxin | 1.43 | 1.32 | -1.97 | 2.76 | 0.78 | 94.4 |
| 265 | ORF7a | B*15:01 | 10 | FALTCFSTQF | -0.09369 | 100.00% (28/28) | Non-Toxin | 1.46 | 1.13 | -1.93 | 2.59 | 0.67 | 84.9 |
| 266 | ORF7a | B*35:01 | 10 | FALTCFSTQF | -0.09369 | 100.00% (28/28) | Non-Toxin | 1.46 | 1.13 | -2.07 | 2.59 | 0.52 | 118.4 |
| 267 | ORF7a | A*68:01 | 10 | FITLCFTLKR | -0.03588 | 100.00% (28/28) | Non-Toxin | 1.16 | 0.66 | -1.25 | 1.83 | 0.58 | 17.8 |
| 268 | ORF7a | B*15:01 | 9 | FLIVAAIVF | 0.29611 | 100.00% (28/28) | Non-Toxin | 1.56 | 1.1 | -2.17 | 2.67 | 0.5 | 146.3 |
| 269 | ORF7a | B*15:01 | 9 | GTYEGNSPF | -0.01964 | 100.00% (28/28) | Non-Toxin | 0.85 | 1.12 | -1.34 | 1.96 | 0.63 | 21.7 |
| 270 | ORF7a | A*23:01 | 10 | IFLIVAAIVF | 0.38189 | 100.00% (28/28) | Non-Toxin | 1.56 | 1.13 | -1.98 | 2.69 | 0.71 | 96.6 |
| 271 | ORF7a | A*02:01 | 9 | ILFLALITL | 0.1895 | 100.00% (28/28) | Non-Toxin | 1.92 | 0.49 | -1.83 | 2.4 | 0.57 | 68.1 |
| 272 | ORF7a | A*30:02 | 9 | ITLATCELY | 0.10084 | 100.00% (28/28) | Non-Toxin | 1.23 | 1.3 | -1.73 | 2.53 | 0.8 | 53.3 |
| 273 | ORF7a | B*58:01 | 9 | ITLATCELY | 0.10084 | 100.00% (28/28) | Non-Toxin | 1.23 | 1.3 | -1.84 | 2.53 | 0.68 | 69.7 |
| 274 | ORF7a | A*23:01 | 11 | KFALTCFSTQF | -0.08981 | 100.00% (28/28) | Non-Toxin | 1.46 | 1.29 | -2.18 | 2.75 | 0.57 | 150.6 |
| 275 | ORF7a | B*35:01 | 9 | LATCELYHY | 0.06119 | 100.00% (28/28) | Non-Toxin | 1.32 | 1.28 | -1.93 | 2.6 | 0.67 | 84.3 |
| 276 | ORF7a | B*15:01 | 11 | LLKEPCSSGTY | -0.2981 | 100.00% (28/28) | Non-Toxin | 1.39 | 1.3 | -1.96 | 2.69 | 0.73 | 91.4 |
| 277 | ORF7a | B*58:01 | 10 | LTCFSTQFAF | -0.01038 | 100.00% (28/28) | Non-Toxin | 1.32 | 1.16 | -1.54 | 2.48 | 0.94 | 34.8 |
| 278 | ORF7a | B*15:01 | 10 | LTCFSTQFAF | -0.01038 | 100.00% (28/28) | Non-Toxin | 1.32 | 1.16 | -1.62 | 2.48 | 0.87 | 41.3 |
| 279 | ORF7a | A*30:01 | 9 | RARSVSPKL | -0.40056 | 100.00% (28/28) | Non-Toxin | 1.57 | 0.59 | -1.23 | 2.15 | 0.93 | 16.9 |
| 280 | ORF7a | A*30:01 | 10 | RARSVSPKLF | -0.46949 | 100.00% (28/28) | Non-Toxin | 1.09 | 1.27 | -1.89 | 2.37 | 0.48 | 77.5 |
| 281 | ORF7a | A*31:01 | 10 | RSVSPKLFIR | -0.20775 | 100.00% (28/28) | Non-Toxin | 0.98 | 0.87 | -1.3 | 1.85 | 0.55 | 20 |
| S.No. | ORF | Allele | Peptide Length | Peptide | Immunogenicity | Conservancy | Toxicity | Proteasome Score | TAP Score | MHC Score | Processing Score | Total Score | MHC IC50[nM] |
| 282 | ORF7b | A*23:01 | 11 | FYLCFLAFLLF | 0.17781 | 100.00% (2/2) | Non-Toxin | 1.24 | 1.12 | -1.75 | 2.36 | 0.61 | 56.8 |
| 283 | ORF7b | B*44:03 | 9 | IELSLIDFY | 0.03153 | 100.00% (2/2) | Non-Toxin | 1.13 | 1.24 | -1.83 | 2.37 | 0.54 | 67.4 |
| 284 | ORF7b | A*02:01 | 10 | IELSLIDFYL | 0.06625 | 100.00% (2/2) | Non-Toxin | 1.42 | 0.39 | -1.28 | 1.81 | 0.53 | 18.9 |
| 285 | ORF7b | A*02:01 | 9 | IIFWFSLEL | 0.2683 | 100.00% (2/2) | Non-Toxin | 1.65 | 0.55 | -1.2 | 2.21 | 1.01 | 15.7 |
| 286 | ORF7b | A*02:01 | 9 | IIFWFSLEL | 0.2683 | 100.00% (2/2) | Non-Toxin | 1.65 | 0.55 | -1.2 | 2.21 | 1.01 | 15.7 |
| 287 | ORF7b | A*02:06 | 9 | IIFWFSLEL | 0.2683 | 100.00% (2/2) | Non-Toxin | 1.65 | 0.55 | -1.55 | 2.21 | 0.66 | 35.5 |
| 288 | ORF7b | A*02:01 | 9 | LIDFYLCFL | 0.13386 | 100.00% (2/2) | Non-Toxin | 1.7 | 0.42 | -1.64 | 2.12 | 0.49 | 43.4 |
| 289 | ORF7b | B*58:01 | 10 | LSLIDFYLCF | 0.22158 | 100.00% (2/2) | Non-Toxin | 1.38 | 1.05 | -1.96 | 2.43 | 0.48 | 90.2 |
| 290 | ORF7b | B*44:03 | 10 | MIELSLIDFY | 0.06184 | 100.00% (2/2) | Non-Toxin | 1.13 | 1.24 | -1.63 | 2.36 | 0.74 | 42.5 |

| | | | | | | | | | | | | | |
|---|---|---|---|---|---|---|---|---|---|---|---|---|---|
| 291 | ORF7b | B*15:01 | 9 | SLIDFYLCF | 0.13518 | 100.00% (2/2) | Non-Toxin | 1.38 | 1.18 | -1.76 | 2.56 | 0.8 | 57.2 |
| 292 | ORF7b | A*02:01 | 10 | SLIDFYLCFL | 0.18838 | 100.00% (2/2) | Non-Toxin | 1.7 | 0.5 | -0.74 | 2.2 | 1.46 | 5.5 |
| 293 | ORF7b | A*02:03 | 10 | SLIDFYLCFL | 0.18838 | 100.00% (2/2) | Non-Toxin | 1.7 | 0.5 | -0.79 | 2.2 | 1.41 | 6.2 |
| 294 | ORF7b | A*02:06 | 10 | SLIDFYLCFL | 0.18838 | 100.00% (2/2) | Non-Toxin | 1.7 | 0.5 | -0.93 | 2.2 | 1.27 | 8.6 |
| 295 | ORF7b | A*02:01 | 10 | SLIDFYLCFL | 0.18838 | 100.00% (2/2) | Non-Toxin | 1.7 | 0.5 | -0.74 | 2.2 | 1.46 | 5.5 |
| 296 | ORF7b | A*02:03 | 10 | SLIDFYLCFL | 0.18838 | 100.00% (2/2) | Non-Toxin | 1.7 | 0.5 | -0.79 | 2.2 | 1.41 | 6.2 |
| 297 | ORF7b | A*02:06 | 10 | SLIDFYLCFL | 0.18838 | 100.00% (2/2) | Non-Toxin | 1.7 | 0.5 | -0.93 | 2.2 | 1.27 | 8.6 |
| 298 | ORF7b | A*02:01 | 9 | YLCFLAFLL | 0.21865 | 100.00% (2/2) | Non-Toxin | 1.36 | 0.37 | -0.9 | 1.73 | 0.83 | 7.9 |
| S.No. | ORF | Allele | Peptide Length | Peptide | Immunogenicity | Conservancy | Toxicity | Proteasome Score | TAP Score | MHC Score | Processing Score | Total Score | MHC IC50[nM] |
| 299 | ORF8 | A*30:02 | 10 | CSFYEDFLEY | 0.31272 | 100.00% (34/34) | Non-Toxin | 1.47 | 1.38 | -1.96 | 2.85 | 0.89 | 91.9 |
| 300 | ORF8 | A*01:01 | 10 | CSFYEDFLEY | 0.31272 | 100.00% (34/34) | Non-Toxin | 1.47 | 1.38 | -2.03 | 2.85 | 0.82 | 108.2 |
| 301 | ORF8 | B*58:01 | 10 | CSFYEDFLEY | 0.31272 | 100.00% (34/34) | Non-Toxin | 1.47 | 1.38 | -2.12 | 2.85 | 0.73 | 132.5 |
| 302 | ORF8 | A*11:01 | 10 | CSFYEDFLEY | 0.31272 | 100.00% (34/34) | Non-Toxin | 1.47 | 1.38 | -2.29 | 2.85 | 0.56 | 195 |
| 303 | ORF8 | B*15:01 | 10 | CSFYEDFLEY | 0.31272 | 100.00% (34/34) | Non-Toxin | 1.47 | 1.38 | -2.37 | 2.85 | 0.48 | 236.6 |
| 304 | ORF8 | B*15:01 | 9 | GIITTVAAF | 0.2148 | 100.00% (34/34) | Non-Toxin | 1.27 | 1.13 | -1.26 | 2.4 | 1.14 | 18.1 |
| 305 | ORF8 | B*15:01 | 9 | GIITTVAAF | 0.2148 | 100.00% (34/34) | Non-Toxin | 1.27 | 1.13 | -1.26 | 2.4 | 1.14 | 18.1 |
| 306 | ORF8 | A*30:02 | 10 | GSLVVRCSFY | 0.00657 | 100.00% (34/34) | Non-Toxin | 1.38 | 1.24 | -1.82 | 2.61 | 0.79 | 66.4 |
| 307 | ORF8 | A*33:01 | 9 | HFYSKWYIR | -0.09452 | 100.00% (34/34) | Non-Toxin | 1.08 | 0.76 | -0.68 | 1.84 | 1.16 | 4.8 |
| 308 | ORF8 | A*31:01 | 9 | HFYSKWYIR | -0.09452 | 100.00% (34/34) | Non-Toxin | 1.08 | 0.76 | -0.79 | 1.84 | 1.05 | 6.2 |
| 309 | ORF8 | A*33:01 | 9 | HFYSKWYIR | -0.09452 | 100.00% (34/34) | Non-Toxin | 1.08 | 0.76 | -0.68 | 1.84 | 1.16 | 4.8 |
| 310 | ORF8 | A*31:01 | 9 | HFYSKWYIR | -0.09452 | 100.00% (34/34) | Non-Toxin | 1.08 | 0.76 | -0.79 | 1.84 | 1.05 | 6.2 |
| 311 | ORF8 | A*31:01 | 10 | IHFYSKWYIR | -0.05367 | 100.00% (34/34) | Non-Toxin | 1.08 | 0.64 | -1.2 | 1.72 | 0.53 | 15.7 |
| 312 | ORF8 | A*30:02 | 9 | IQYIDIGNY | 0.30442 | 100.00% (34/34) | Non-Toxin | 1.2 | 1.33 | -1.66 | 2.53 | 0.87 | 45.4 |
| 313 | ORF8 | B*15:01 | 9 | IQYIDIGNY | 0.30442 | 100.00% (34/34) | Non-Toxin | 1.2 | 1.33 | -1.66 | 2.53 | 0.87 | 45.4 |
| 314 | ORF8 | B*40:01 | 10 | LEYHDVRVVL | 0.20083 | 100.00% (34/34) | Non-Toxin | 1.75 | 0.4 | -1.64 | 2.15 | 0.51 | 43.5 |
| 315 | ORF8 | B*15:01 | 10 | LGIITTVAAF | 0.34746 | 100.00% (34/34) | Non-Toxin | 1.27 | 0.97 | -1.1 | 2.24 | 1.14 | 12.7 |
| 316 | ORF8 | B*15:01 | 10 | LGIITTVAAF | 0.34746 | 100.00% (34/34) | Non-Toxin | 1.27 | 0.97 | -1.1 | 2.24 | 1.14 | 12.7 |
| 317 | ORF8 | B*15:01 | 10 | LQSCTQHQPY | -0.25674 | 100.00% (34/34) | Non-Toxin | 1 | 1.33 | -1.57 | 2.32 | 0.75 | 37 |
| 318 | ORF8 | A*30:02 | 9 | SFYEDFLEY | 0.28049 | 100.00% (34/34) | Non-Toxin | 1.47 | 1.44 | -2.04 | 2.92 | 0.88 | 110.3 |
| 319 | ORF8 | B*35:01 | 9 | SFYEDFLEY | 0.28049 | 100.00% (34/34) | Non-Toxin | 1.47 | 1.44 | -2.28 | 2.92 | 0.63 | 192.4 |
| S.No. | ORF | Allele | Peptide Length | Peptide | Immunogenicity | Conservancy | Toxicity | Proteasome Score | TAP Score | MHC Score | Processing Score | Total Score | MHC IC50[nM] |
| 320 | S protein | A*30:02 | 9 | ASFSTFKCY | -0.19397 | 100.00% (41/41) | Non-Toxin | 1.47 | 1.42 | -1.6 | 2.89 | 1.29 | 39.6 |
| 321 | S protein | B*35:01 | 9 | CVADYSVLY | -0.09595 | 100.00% (41/41) | Non-Toxin | 1.51 | 1.38 | -1.39 | 2.89 | 1.5 | 24.4 |
| 322 | S protein | A*26:01 | 9 | CVADYSVLY | -0.09595 | 100.00% (41/41) | Non-Toxin | 1.51 | 1.38 | -1.56 | 2.89 | 1.34 | 36 |
| 323 | S protein | A*01:01 | 9 | CVADYSVLY | -0.09595 | 100.00% (41/41) | Non-Toxin | 1.51 | 1.38 | -1.74 | 2.89 | 1.16 | 54.5 |
| 324 | S protein | B*35:01 | 9 | FAMQMAYRF | -0.28061 | 100.00% (41/41) | Non-Toxin | 1.45 | 1.05 | -0.8 | 2.5 | 1.7 | 6.3 |
| 325 | S protein | B*53:01 | 9 | FAMQMAYRF | -0.28061 | 100.00% (41/41) | Non-Toxin | 1.45 | 1.05 | -1.15 | 2.5 | 1.35 | 14.1 |
| 326 | S protein | B*58:01 | 9 | FAMQMAYRF | -0.28061 | 100.00% (41/41) | Non-Toxin | 1.45 | 1.05 | -1.42 | 2.5 | 1.07 | 26.4 |
| 327 | S protein | A*23:01 | 9 | FAMQMAYRF | -0.28061 | 100.00% (41/41) | Non-Toxin | 1.45 | 1.05 | -1.43 | 2.5 | 1.07 | 26.7 |
| 328 | S protein | B*40:01 | 9 | FEYVSQPFL | -0.17076 | 100.00% (41/41) | Non-Toxin | 1.84 | 0.4 | -1.14 | 2.24 | 1.09 | 13.9 |
| 329 | S protein | B*53:01 | 10 | FLPFFSNVTW | 0.11853 | 100.00% (41/41) | Non-Toxin | 1.69 | 0.37 | -0.88 | 2.06 | 1.18 | 7.6 |
| 330 | S protein | B*35:01 | 10 | FPNITNLCPF | 0.1009 | 100.00% (41/41) | Non-Toxin | 1.02 | 0.94 | -0.59 | 1.97 | 1.38 | 3.9 |
| 331 | S protein | B*35:01 | 11 | FPQSAPHGVVF | -0.05441 | 100.00% (41/41) | Non-Toxin | 1.56 | 0.94 | -1.47 | 2.5 | 1.02 | 29.7 |
| 332 | S protein | B*35:01 | 9 | FVFKNIDGY | -0.0215 | 100.00% (41/41) | Non-Toxin | 1.1 | 1.26 | -1.18 | 2.36 | 1.19 | 15.1 |
| 333 | S protein | A*26:01 | 9 | FVFKNIDGY | -0.0215 | 100.00% (41/41) | Non-Toxin | 1.1 | 1.26 | -1.36 | 2.36 | 1.01 | 22.8 |
| 334 | S protein | B*58:01 | 9 | GTITSGWTF | 0.16268 | 100.00% (41/41) | Non-Toxin | 1.58 | 1.11 | -1.65 | 2.69 | 1.04 | 44.6 |
| 335 | S protein | B*35:01 | 9 | IPFAMQMAY | -0.32801 | 100.00% (41/41) | Non-Toxin | 1.42 | 1.17 | -0.36 | 2.6 | 2.24 | 2.3 |
| 336 | S protein | B*53:01 | 9 | IPFAMQMAY | -0.32801 | 100.00% (41/41) | Non-Toxin | 1.42 | 1.17 | -1.23 | 2.6 | 1.37 | 16.9 |
| 337 | S protein | B*58:01 | 10 | KRSFIEDLLF | 0.29624 | 100.00% (41/41) | Non-Toxin | 1.23 | 1.26 | -1.11 | 2.49 | 1.37 | 12.9 |
| 338 | S protein | A*30:02 | 10 | KSFTVEKGIY | 0.11812 | 100.00% (41/41) | Non-Toxin | 1.25 | 1.4 | -1.37 | 2.65 | 1.28 | 23.4 |
| 339 | S protein | B*58:01 | 10 | KSNIIRGWIF | 0.60842 | 100.00% (41/41) | Non-Toxin | 1.45 | 1.05 | -1.43 | 2.5 | 1.07 | 27 |

| | | | | | | | | | | | | | |
|---|---|---|---|---|---|---|---|---|---|---|---|---|---|
| 340 | S protein | A*30:02 | 10 | KVGGNYNYLY | 0.01951 | 100.00% (41/41) | Non-Toxin | 1.4 | 1.33 | -1.43 | 2.73 | 1.29 | 27.1 |
| 341 | S protein | A*23:01 | 10 | KWPWYIWLGF | 0.56424 | 100.00% (41/41) | Non-Toxin | 1.26 | 1.25 | -1.04 | 2.51 | 1.47 | 11 |
| 342 | S protein | A*24:02 | 10 | KWPWYIWLGF | 0.56424 | 100.00% (41/41) | Non-Toxin | 1.26 | 1.25 | -1.14 | 2.51 | 1.37 | 13.9 |
| 343 | S protein | A*01:01 | 10 | LLTDEMIAQY | 0.05204 | 100.00% (41/41) | Non-Toxin | 1.21 | 1.2 | -1.08 | 2.42 | 1.33 | 12.1 |
| 344 | S protein | B*53:01 | 9 | LPFFSNVTW | 0.04613 | 100.00% (41/41) | Non-Toxin | 1.69 | 0.26 | -0.77 | 1.95 | 1.18 | 5.9 |
| 345 | S protein | B*35:01 | 9 | LPFNDGVYF | 0.11767 | 100.00% (41/41) | Non-Toxin | 1.41 | 1.04 | -0.54 | 2.45 | 1.9 | 3.5 |
| 346 | S protein | B*53:01 | 9 | LPFNDGVYF | 0.11767 | 100.00% (41/41) | Non-Toxin | 1.41 | 1.04 | -0.98 | 2.45 | 1.47 | 9.6 |
| 347 | S protein | B*35:01 | 11 | LQIPFAMQMAY | -0.22124 | 100.00% (41/41) | Non-Toxin | 1.42 | 1.4 | -1.25 | 2.82 | 1.57 | 17.8 |
| 348 | S protein | B*15:01 | 11 | LQIPFAMQMAY | -0.22124 | 100.00% (41/41) | Non-Toxin | 1.42 | 1.4 | -1.64 | 2.82 | 1.19 | 43.2 |
| 349 | S protein | A*01:01 | 9 | LTDEMIAQY | 0.02757 | 100.00% (41/41) | Non-Toxin | 1.21 | 1.21 | -0.72 | 2.42 | 1.71 | 5.2 |
| 350 | S protein | A*23:01 | 10 | LYNSASFSTF | -0.29831 | 100.00% (41/41) | Non-Toxin | 1.33 | 1.2 | -0.95 | 2.52 | 1.57 | 9 |
| 351 | S protein | A*24:02 | 10 | LYNSASFSTF | -0.29831 | 100.00% (41/41) | Non-Toxin | 1.33 | 1.2 | -0.99 | 2.52 | 1.53 | 9.8 |
| 352 | S protein | A*68:01 | 10 | NVYADSFVIR | 0.12147 | 100.00% (41/41) | Non-Toxin | 1.47 | 0.81 | -1.03 | 2.28 | 1.24 | 10.8 |
| 353 | S protein | A*24:02 | 9 | NYNYLYRLF | 0.0171 | 100.00% (41/41) | Non-Toxin | 1.2 | 1.18 | -1.37 | 2.38 | 1.01 | 23.4 |
| 354 | S protein | A*33:01 | 10 | NYNYLYRLFR | 0.08754 | 100.00% (41/41) | Non-Toxin | 1.16 | 0.72 | -0.86 | 1.88 | 1.02 | 7.3 |
| 355 | S protein | B*35:01 | 10 | QIPFAMQMAY | -0.25308 | 100.00% (41/41) | Non-Toxin | 1.42 | 1.35 | -1.16 | 2.77 | 1.61 | 14.5 |
| 356 | S protein | A*30:02 | 9 | RISNCVADY | -0.02787 | 100.00% (41/41) | Non-Toxin | 1.16 | 1.47 | -1.26 | 2.63 | 1.37 | 18.2 |
| 357 | S protein | B*58:01 | 9 | RSFIEDLLF | 0.27446 | 100.00% (41/41) | Non-Toxin | 1.23 | 1.32 | -0.72 | 2.54 | 1.82 | 5.3 |
| 358 | S protein | B*57:01 | 9 | RSFIEDLLF | 0.27446 | 100.00% (41/41) | Non-Toxin | 1.23 | 1.32 | -1.18 | 2.54 | 1.36 | 15.2 |
| 359 | S protein | B*15:01 | 10 | RVYSTGSNVF | -0.23394 | 100.00% (41/41) | Non-Toxin | 1.51 | 1.32 | -1.02 | 2.83 | 1.81 | 10.5 |
| 360 | S protein | A*32:01 | 10 | RVYSTGSNVF | -0.23394 | 100.00% (41/41) | Non-Toxin | 1.51 | 1.32 | -1.69 | 2.83 | 1.14 | 48.9 |
| 361 | S protein | B*35:01 | 9 | SANNCTFEY | 0.13273 | 100.00% (41/41) | Non-Toxin | 1.18 | 1.3 | -1.11 | 2.48 | 1.37 | 12.8 |
| 362 | S protein | B*15:01 | 10 | SVASQSIIAY | -0.16721 | 100.00% (41/41) | Non-Toxin | 1.32 | 1.42 | -1.74 | 2.74 | 1 | 55.2 |
| 363 | S protein | B*35:01 | 9 | VASQSIIAY | -0.0709 | 100.00% (41/41) | Non-Toxin | 1.32 | 1.34 | -0.86 | 2.66 | 1.8 | 7.2 |
| 364 | S protein | A*24:02 | 10 | VYSSANNCTF | -0.21728 | 100.00% (41/41) | Non-Toxin | 1.42 | 1.3 | -1.62 | 2.71 | 1.09 | 41.7 |
| 365 | S protein | A*23:01 | 10 | VYSSANNCTF | -0.21728 | 100.00% (41/41) | Non-Toxin | 1.42 | 1.3 | -1.71 | 2.71 | 1.01 | 50.8 |
| 366 | S protein | A*26:01 | 9 | WTAGAAAYY | 0.15259 | 100.00% (41/41) | Non-Toxin | 1.24 | 1.24 | -1.06 | 2.48 | 1.41 | 11.6 |
| 367 | S protein | A*68:01 | 9 | WTAGAAAYY | 0.15259 | 100.00% (41/41) | Non-Toxin | 1.24 | 1.24 | -1.37 | 2.48 | 1.11 | 23.5 |
| 368 | S protein | A*02:01 | 9 | YLQPRTFLL | 0.1305 | 100.00% (41/41) | Non-Toxin | 1.46 | 0.39 | -0.66 | 1.85 | 1.18 | 4.6 |
| 369 | S protein | B*35:01 | 9 | YSSANNCTF | -0.04954 | 100.00% (41/41) | Non-Toxin | 1.42 | 1.13 | -1.29 | 2.54 | 1.26 | 19.4 |
| 370 | S protein | B*58:01 | 9 | YSSANNCTF | -0.04954 | 100.00% (41/41) | Non-Toxin | 1.42 | 1.13 | -1.49 | 2.54 | 1.05 | 31.1 |
| 371 | S protein | B*15:01 | 10 | YSVLYNSASF | -0.22703 | 100.00% (41/41) | Non-Toxin | 1.39 | 1.08 | -1.36 | 2.47 | 1.11 | 23 |
| 372 | S protein | A*30:02 | 10 | YTNSFTRGVY | 0.08467 | 95.12% (39/41) | Non-Toxin | 1.36 | 1.28 | -1.26 | 2.64 | 1.38 | 18.2 |
| 373 | S protein | A*01:01 | 10 | YTNSFTRGVY | 0.08467 | 95.12% (39/41) | Non-Toxin | 1.36 | 1.28 | -1.45 | 2.64 | 1.19 | 28 |
| 374 | S protein | B*15:01 | 10 | YTNSFTRGVY | 0.08467 | 95.12% (39/41) | Non-Toxin | 1.36 | 1.28 | -1.6 | 2.64 | 1.04 | 40.2 |
| 375 | S protein | A*01:01 | 11 | YTNSFTRGVYY | 0.09821 | 95.12% (39/41) | Non-Toxin | 1.45 | 1.28 | -1.51 | 2.72 | 1.22 | 32.1 |
| 376 | S protein | A*23:01 | 11 | YYVGYLQPRTF | -0.02378 | 100.00% (41/41) | Non-Toxin | 1.36 | 1.31 | -1.38 | 2.67 | 1.29 | 24 |
| 377 | S protein | A*24:02 | 11 | YYVGYLQPRTF | -0.02378 | 100.00% (41/41) | Non-Toxin | 1.36 | 1.31 | -1.67 | 2.67 | 1 | 46.5 |

**Supplementary table S2:** High Scoring CTL epitopes-HLA allele pairs screened from entire proteome of SARS-CoV-2 by the "MHC-I Binding Predictions" tool of IEDB. These epitopes were further utilized to identify the potentially immunogenic multiple epitope cluster based CTL Ag-Patches from the entire proteome of the SRAS-CoV-2. The epitopes shown in **RED** are the epitope which form Overlapping epitope clusters. The screened epitopes are in consensus with the previous studies [Srivastava et al. 2020a; Srivastava et al. 2020b; and Grifoni et al., 2020a].

| S.No. | ORF | Allele | Length | Peptide | Immunogenicity | Method used | Percentile Rank | Conservancy | Toxicity |
|---|---|---|---|---|---|---|---|---|---|
| 378 | E Protein | A*02:01 | 9 | FLAFVVFLL | 0.30188 | Consensus (ann/comblib_sidney2008/smm) | 0.2 | 99.59% (480/482) | Non-Toxin |
| 379 | E Protein | A*02:03 | 9 | FLAFVVFLL | 0.30188 | Consensus (ann/smm) | 0.25 | 99.59% (480/482) | Non-Toxin |
| 380 | E Protein | A*02:06 | 9 | FLAFVVFLL | 0.30188 | Consensus (ann/smm) | 0.53 | 99.59% (480/482) | Non-Toxin |
| 381 | E Protein | A*02:01 | 10 | FLAFVVFLLV | 0.30526 | Consensus (ann/smm) | 0.15 | 99.59% (480/482) | Non-Toxin |
| 382 | E Protein | A*02:03 | 10 | FLAFVVFLLV | 0.30526 | Consensus (ann/smm) | 0.23 | 99.59% (480/482) | Non-Toxin |
| 383 | E Protein | A*02:06 | 10 | FLAFVVFLLV | 0.30526 | Consensus (ann/smm) | 0.47 | 99.59% (480/482) | Non-Toxin |
| 384 | E Protein | A*02:01 | 9 | FLLVTLAIL | 0.17608 | Consensus (ann/comblib_sidney2008/smm) | 0.43 | 99.59% (480/482) | Non-Toxin |
| 385 | E Protein | A*02:06 | 9 | FVVFLLVTL | 0.16748 | Consensus (ann/smm) | 0.46 | 99.59% (480/482) | Non-Toxin |
| 386 | E Protein | B*15:01 | 10 | ILTALRLCAY | 0.05849 | Consensus (ann/smm) | 0.41 | 99.17% (478/482) | Non-Toxin |
| 387 | E Protein | B*51:01 | 9 | LAFVVFLLV | 0.2141 | Consensus (ann/comblib_sidney2008/smm) | 0.2 | 99.59% (480/482) | Non-Toxin |
| 388 | E Protein | A*02:01 | 11 | LFLAFVVFLLV | 0.32453 | Consensus (ann/smm) | 0.39 | 99.59% (480/482) | Non-Toxin |
| 389 | E Protein | B*15:01 | 9 | LLFLAFVVF | 0.2341 | Consensus (ann/comblib_sidney2008/smm) | 0.1 | 99.59% (480/482) | Non-Toxin |
| 390 | E Protein | A*32:01 | 9 | LLFLAFVVF | 0.2341 | Consensus (ann/comblib_sidney2008/smm) | 0.3 | 99.59% (480/482) | Non-Toxin |
| 391 | E Protein | A*02:01 | 10 | LLFLAFVVFL | 0.32104 | Consensus (ann/smm) | 0.48 | 99.59% (480/482) | Non-Toxin |
| 392 | E Protein | A*01:01 | 9 | LTALRLCAY | 0.01886 | Consensus (ann/smm) | 0.12 | 99.17% (478/482) | Non-Toxin |
| 393 | E Protein | A*26:01 | 9 | LTALRLCAY | 0.01886 | Consensus (ann/smm) | 0.41 | 99.17% (478/482) | Non-Toxin |
| 394 | E Protein | B*15:01 | 9 | LVKPSFYVY | -0.11106 | Consensus (ann/comblib_sidney2008/smm) | 0.2 | 99.59% (480/482) | Non-Toxin |
| 395 | E Protein | A*30:02 | 9 | LVKPSFYVY | -0.11106 | Consensus (ann/smm) | 0.46 | 99.59% (480/482) | Non-Toxin |
| 396 | E Protein | A*68:02 | 10 | NSVLLFLAFV | 0.19642 | Consensus (ann/smm) | 0.38 | 99.59% (480/482) | Non-Toxin |
| 397 | E Protein | A*31:01 | 9 | RVKNLNSSR | -0.32968 | Consensus (ann/smm) | 0.16 | 99.59% (480/482) | Non-Toxin |
| 398 | E Protein | B*40:01 | 9 | SEETGTLIV | 0.2095 | Consensus (ann/smm) | 0.45 | 99.59% (480/482) | Non-Toxin |
| 399 | E Protein | A*02:06 | 9 | SLVKPSFYV | -0.27349 | Consensus (ann/smm) | 0.4 | 99.59% (480/482) | Non-Toxin |
| 400 | E Protein | A*02:01 | 9 | SLVKPSFYV | -0.27349 | Consensus (ann/comblib_sidney2008/smm) | 0.5 | 99.59% (480/482) | Non-Toxin |
| 401 | E Protein | B*15:01 | 10 | SLVKPSFYVY | -0.2443 | Consensus (ann/smm) | 0.28 | 99.59% (480/482) | Non-Toxin |
| 402 | E Protein | A*30:02 | 10 | SLVKPSFYVY | -0.2443 | Consensus (ann/smm) | 0.58 | 99.59% (480/482) | Non-Toxin |
| 403 | E Protein | A*02:06 | 9 | SVLLFLAFV | 0.19022 | Consensus (ann/smm) | 0.33 | 99.59% (480/482) | Non-Toxin |
| 404 | E Protein | A*02:06 | 10 | SVLLFLAFVV | 0.24819 | Consensus (ann/smm) | 0.48 | 99.59% (480/482) | Non-Toxin |
| 405 | E Protein | A*68:01 | 9 | TLAILTALR | 0.1989 | Consensus (ann/smm) | 0.21 | 99.17% (478/482) | Non-Toxin |
| 406 | E Protein | A*23:01 | 9 | VFLLVTLAI | 0.07548 | Consensus (ann/smm) | 0.5 | 99.59% (480/482) | Non-Toxin |
| 407 | E Protein | A*02:01 | 9 | VLLFLAFVV | 0.26315 | Consensus (ann/comblib_sidney2008/smm) | 0.3 | 99.59% (480/482) | Non-Toxin |
| 408 | E Protein | B*15:01 | 10 | VLLFLAFVVF | 0.31066 | Consensus (ann/smm) | 0.46 | 99.59% (480/482) | Non-Toxin |
| 409 | E Protein | A*30:02 | 9 | VSLVKPSFY | -0.25372 | Consensus (ann/smm) | 0.33 | 99.59% (480/482) | Non-Toxin |
| 410 | E Protein | A*68:01 | 10 | VTLAILTALR | 0.21765 | Consensus (ann/smm) | 0.52 | 99.17% (478/482) | Non-Toxin |
| S.No. | ORF | Allele | Length | Peptide | Immunogenicity | Method used | Percentile Rank | Conservancy | Toxicity |
| 411 | M Protein | B*58:01 | 10 | AMACLVGLMW | -0.09221 | Consensus (ann/smm) | 0.41 | 98.53% (470/477) | Non-Toxin |
| 412 | M Protein | B*44:02 | 10 | AMACLVGLMW | -0.09221 | Consensus (ann/smm) | 0.41 | 98.53% (470/477) | Non-Toxin |
| 413 | M Protein | A*30:01 | 9 | ANRNRFLYI | 0.15937 | Consensus (ann/comblib_sidney2008/smm) | 0.5 | 99.16% (473/477) | Non-Toxin |
| 414 | M Protein | A*31:01 | 8 | ASFRLFAR | 0.2225 | ann | 0.27 | 99.79% (476/477) | Non-Toxin |
| 415 | M Protein | A*31:01 | 10 | ASFRLFARTR | 0.29647 | Consensus (ann/smm) | 0.27 | 99.79% (476/477) | Non-Toxin |
| 416 | M Protein | A*30:02 | 9 | ATSRTLSYY | -0.11604 | Consensus (ann/smm) | 0.17 | 98.95% (472/477) | Non-Toxin |
| 417 | M Protein | A*01:01 | 9 | ATSRTLSYY | -0.11604 | Consensus (ann/smm) | 0.17 | 98.95% (472/477) | Non-Toxin |
| 418 | M Protein | A*26:01 | 9 | ATSRTLSYY | -0.11604 | Consensus (ann/smm) | 0.26 | 98.95% (472/477) | Non-Toxin |
| 419 | M Protein | A*11:01 | 10 | ATSRTLSYYK | -0.13563 | Consensus (ann/smm) | 0.06 | 98.95% (472/477) | Non-Toxin |

| | | | | | | | | | |
|---|---|---|---|---|---|---|---|---|---|
| 420 | M Protein | A*03:01 | 10 | ATSRTLSYYK | -0.13563 | Consensus (ann/smm) | 0.14 | 98.95% (472/477) | Non-Toxin |
| 421 | M Protein | A*30:01 | 10 | ATSRTLSYYK | -0.13563 | Consensus (ann/smm) | 0.17 | 98.95% (472/477) | Non-Toxin |
| 422 | M Protein | A*31:01 | 9 | AVILRGHLR | 0.13516 | Consensus (ann/smm) | 0.43 | 99.79% (476/477) | Non-Toxin |
| 423 | M Protein | A*30:02 | 10 | AYANRNRFLY | 0.19133 | Consensus (ann/smm) | 0.24 | 98.95% (472/477) | Non-Toxin |
| 424 | M Protein | A*30:02 | 10 | AYSRYRIGNY | 0.19528 | Consensus (ann/smm) | 0.27 | 98.32% (469/477) | Non-Toxin |
| 425 | M Protein | B*15:01 | 10 | CLVGLMWLSY | -0.03181 | Consensus (ann/smm) | 0.49 | 97.48% (465/477) | Non-Toxin |
| 426 | M Protein | A*30:02 | 10 | DSGFAAYSRY | 0.09214 | Consensus (ann/smm) | 0.38 | 98.32% (469/477) | Non-Toxin |
| 427 | M Protein | B*44:03 | 10 | EELKKLLEQW | -0.43502 | Consensus (ann/smm) | 0.2 | 98.74% (471/477) | Non-Toxin |
| 428 | M Protein | B*44:02 | 10 | EELKKLLEQW | -0.43502 | Consensus (ann/smm) | 0.24 | 98.74% (471/477) | Non-Toxin |
| 429 | M Protein | A*23:01 | 11 | EQWNLVIGFLF | 0.34861 | Consensus (ann/smm) | 0.38 | 99.16% (473/477) | Non-Toxin |
| 430 | M Protein | B*51:01 | 9 | FAAYSRYRI | -0.07628 | Consensus (ann/comblib_sidney2008/smm) | 0.46 | 98.32% (469/477) | Non-Toxin |
| 431 | M Protein | A*68:02 | 9 | FAAYSRYRI | -0.07628 | Consensus (ann/comblib_sidney2008/smm) | 0.5 | 98.32% (469/477) | Non-Toxin |
| 432 | M Protein | B*35:01 | 9 | FAYANRNRF | 0.10537 | Consensus (ann/comblib_sidney2008/smm) | 0.3 | 98.95% (472/477) | Non-Toxin |
| 433 | M Protein | A*68:01 | 10 | FIASFRLFAR | 0.12185 | Consensus (ann/smm) | 0.12 | 99.79% (476/477) | Non-Toxin |
| 434 | M Protein | A*33:01 | 10 | FIASFRLFAR | 0.12185 | Consensus (ann/smm) | 0.14 | 99.79% (476/477) | Non-Toxin |
| 435 | M Protein | A*31:01 | 10 | FIASFRLFAR | 0.12185 | Consensus (ann/smm) | 0.3 | 99.79% (476/477) | Non-Toxin |
| 436 | M Protein | A*02:01 | 9 | FLFLTWICL | 0.35397 | Consensus (ann/comblib_sidney2008/smm) | 0.4 | 98.95% (472/477) | Non-Toxin |
| 437 | M Protein | A*02:01 | 10 | FLFLTWICLL | 0.35364 | Consensus (ann/smm) | 0.15 | 98.95% (472/477) | Non-Toxin |
| 438 | M Protein | A*02:01 | 10 | FLWLLWPVTL | 0.31272 | Consensus (ann/smm) | 0.28 | 98.53% (470/477) | Non-Toxin |
| 439 | M Protein | A*02:01 | 10 | FLYIIKLIFL | 0.2226 | Consensus (ann/smm) | 0.29 | 98.53% (470/477) | Non-Toxin |
| 440 | M Protein | B*08:01 | 10 | FRLFARTRSM | 0.18626 | Consensus (ann/smm) | 0.43 | 99.79% (476/477) | Non-Toxin |
| 441 | M Protein | A*31:01 | 9 | GFAAYSRYR | -0.06574 | Consensus (ann/smm) | 0.41 | 98.32% (469/477) | Non-Toxin |
| 442 | M Protein | A*02:01 | 9 | GLMWLSYFI | 0.06464 | Consensus (ann/comblib_sidney2008/smm) | 0.2 | 97.48% (465/477) | Non-Toxin |
| 443 | M Protein | A*32:01 | 9 | GLMWLSYFI | 0.06464 | Consensus (ann/comblib_sidney2008/smm) | 0.4 | 97.48% (465/477) | Non-Toxin |
| 444 | M Protein | A*11:01 | 9 | GTITVEELK | 0.29473 | Consensus (ann/smm) | 0.36 | 98.74% (471/477) | Non-Toxin |
| 445 | M Protein | A*68:01 | 9 | GTITVEELK | 0.29473 | Consensus (ann/smm) | 0.47 | 98.74% (471/477) | Non-Toxin |
| 446 | M Protein | A*11:01 | 10 | GTITVEELKK | 0.17885 | Consensus (ann/smm) | 0.27 | 98.53% (470/477) | Non-Toxin |
| 447 | M Protein | B*51:01 | 9 | IAIAMACLV | -0.10358 | Consensus (ann/comblib_sidney2008/smm) | 0.5 | 99.58% (475/477) | Non-Toxin |
| 448 | M Protein | B*58:01 | 11 | IAMACLVGLMW | -0.12079 | Consensus (ann/smm) | 0.42 | 98.53% (470/477) | Non-Toxin |
| 449 | M Protein | A*33:01 | 9 | IASFRLFAR | 0.22572 | Consensus (ann/smm) | 0.23 | 99.79% (476/477) | Non-Toxin |
| 450 | M Protein | A*68:01 | 9 | IASFRLFAR | 0.22572 | Consensus (ann/smm) | 0.26 | 99.79% (476/477) | Non-Toxin |
| 451 | M Protein | A*31:01 | 9 | IASFRLFAR | 0.22572 | Consensus (ann/smm) | 0.29 | 99.79% (476/477) | Non-Toxin |
| 452 | M Protein | A*02:06 | 9 | IFLWLLWPV | 0.37851 | Consensus (ann/smm) | 0.28 | 98.53% (470/477) | Non-Toxin |
| 453 | M Protein | B*40:01 | 11 | KEITVATSRTL | 0.10899 | ann | 0.42 | 98.74% (471/477) | Non-Toxin |
| 454 | M Protein | A*32:01 | 11 | KKLLEQWNLVI | 0.19431 | ann | 0.39 | 98.74% (471/477) | Non-Toxin |
| 455 | M Protein | B*58:01 | 9 | KLIFLWLLW | 0.34287 | Consensus (ann/comblib_sidney2008/smm) | 0.2 | 98.32% (469/477) | Non-Toxin |
| 456 | M Protein | A*32:01 | 9 | KLIFLWLLW | 0.34287 | Consensus (ann/comblib_sidney2008/smm) | 0.3 | 98.32% (469/477) | Non-Toxin |
| 457 | M Protein | B*57:01 | 9 | KLIFLWLLW | 0.34287 | Consensus (ann/smm) | 0.42 | 98.32% (469/477) | Non-Toxin |
| 458 | M Protein | A*02:01 | 11 | KLIFLWLLWPV | 0.52512 | Consensus (ann/smm) | 0.14 | 98.11% (468/477) | Non-Toxin |
| 459 | M Protein | A*32:01 | 11 | KLIFLWLLWPV | 0.52512 | ann | 0.2 | 98.11% (468/477) | Non-Toxin |
| 460 | M Protein | A*02:06 | 9 | KLLEQWNLV | 0.18092 | Consensus (ann/smm) | 0.17 | 98.95% (472/477) | Non-Toxin |
| 461 | M Protein | A*02:01 | 9 | KLLEQWNLV | 0.18092 | Consensus (ann/comblib_sidney2008/smm) | 0.4 | 98.95% (472/477) | Non-Toxin |
| 462 | M Protein | A*32:01 | 11 | KLLEQWNLVIG | 0.29591 | ann | 0.38 | 98.95% (472/477) | Non-Toxin |
| 463 | M Protein | B*44:02 | 10 | LEQWNLVIGF | 0.33917 | Consensus (ann/smm) | 0.29 | 99.16% (473/477) | Non-Toxin |
| 464 | M Protein | B*44:03 | 10 | LEQWNLVIGF | 0.33917 | Consensus (ann/smm) | 0.48 | 99.16% (473/477) | Non-Toxin |
| 465 | M Protein | A*02:06 | 10 | LIFLWLLWPV | 0.40176 | Consensus (ann/smm) | 0.08 | 98.32% (469/477) | Non-Toxin |
| 466 | M Protein | A*02:01 | 10 | LIFLWLLWPV | 0.40176 | Consensus (ann/smm) | 0.15 | 98.32% (469/477) | Non-Toxin |
| 467 | M Protein | A*02:03 | 10 | LIFLWLLWPV | 0.40176 | Consensus (ann/smm) | 0.3 | 98.32% (469/477) | Non-Toxin |
| 468 | M Protein | B*15:01 | 10 | LLWPVTLACF | 0.12982 | Consensus (ann/smm) | 0.13 | 98.53% (470/477) | Non-Toxin |
| 469 | M Protein | A*02:01 | 11 | LLWPVTLACFV | 0.18767 | Consensus (ann/smm) | 0.35 | 98.53% (470/477) | Non-Toxin |
| 470 | M Protein | B*15:01 | 11 | LMWLSYFIASF | 0.07168 | ann | 0.42 | 97.69% (466/477) | Non-Toxin |

| | | | | | | | | | |
|---|---|---|---|---|---|---|---|---|---|
| 471 | M Protein | A*23:01 | 11 | LMWLSYFIASF | 0.07168 | Consensus (ann/smm) | 0.47 | 97.69% (466/477) | Non-Toxin |
| 472 | M Protein | A*68:01 | 9 | LSYFIASFR | 0.21181 | Consensus (ann/smm) | 0.11 | 97.90% (467/477) | Non-Toxin |
| 473 | M Protein | A*31:01 | 9 | LSYFIASFR | 0.21181 | Consensus (ann/smm) | 0.17 | 97.90% (467/477) | Non-Toxin |
| 474 | M Protein | A*33:01 | 9 | LSYFIASFR | 0.21181 | Consensus (ann/smm) | 0.41 | 97.90% (467/477) | Non-Toxin |
| 475 | M Protein | A*03:01 | 9 | LSYFIASFR | 0.21181 | Consensus (ann/smm) | 0.48 | 97.90% (467/477) | Non-Toxin |
| 476 | M Protein | A*23:01 | 11 | LSYFIASFRLF | 0.2706 | Consensus (ann/smm) | 0.34 | 97.90% (467/477) | Non-Toxin |
| 477 | M Protein | B*53:01 | 10 | LVIGFLFLTW | 0.30402 | Consensus (ann/smm) | 0.15 | 98.95% (472/477) | Non-Toxin |
| 478 | M Protein | A*23:01 | 9 | LWLLWPVTL | 0.24802 | Consensus (ann/smm) | 0.43 | 98.53% (470/477) | Non-Toxin |
| 479 | M Protein | A*24:02 | 9 | LWLLWPVTL | 0.24802 | Consensus (ann/smm) | 0.43 | 98.53% (470/477) | Non-Toxin |
| 480 | M Protein | A*24:02 | 9 | LWPVTLACF | 0.06682 | Consensus (ann/smm) | 0.2 | 98.74% (471/477) | Non-Toxin |
| 481 | M Protein | A*23:01 | 10 | LYIIKLIFLW | 0.17392 | Consensus (ann/smm) | 0.17 | 98.53% (470/477) | Non-Toxin |
| 482 | M Protein | A*24:02 | 10 | LYIIKLIFLW | 0.17392 | Consensus (ann/smm) | 0.23 | 98.53% (470/477) | Non-Toxin |
| 483 | M Protein | B*57:01 | 9 | MACLVGLMW | -0.06852 | Consensus (ann/smm) | 0.15 | 98.53% (470/477) | Non-Toxin |
| 484 | M Protein | B*58:01 | 9 | MACLVGLMW | -0.06852 | Consensus (ann/comblib_sidney2008/smm) | 0.2 | 98.53% (470/477) | Non-Toxin |
| 485 | M Protein | B*53:01 | 9 | MACLVGLMW | -0.06852 | Consensus (ann/comblib_sidney2008/smm) | 0.2 | 98.53% (470/477) | Non-Toxin |
| 486 | M Protein | A*23:01 | 10 | MWLSYFIASF | 0.00197 | Consensus (ann/smm) | 0.11 | 97.69% (466/477) | Non-Toxin |
| 487 | M Protein | A*24:02 | 10 | MWLSYFIASF | 0.00197 | Consensus (ann/smm) | 0.13 | 97.69% (466/477) | Non-Toxin |
| 488 | M Protein | A*32:01 | 10 | MWLSYFIASF | 0.00197 | Consensus (ann/smm) | 0.46 | 97.69% (466/477) | Non-Toxin |
| 489 | M Protein | A*68:01 | 11 | MWLSYFIASFR | 0.03554 | ann | 0.09 | 97.69% (466/477) | Non-Toxin |
| 490 | M Protein | A*33:01 | 11 | MWLSYFIASFR | 0.03554 | ann | 0.1 | 97.69% (466/477) | Non-Toxin |
| 491 | M Protein | A*30:01 | 10 | NRNRFLYIIK | 0.33978 | Consensus (ann/smm) | 0.45 | 98.95% (472/477) | Non-Toxin |
| 492 | M Protein | A*23:01 | 10 | QWNLVIGFLF | 0.28076 | Consensus (ann/smm) | 0.12 | 99.16% (473/477) | Non-Toxin |
| 493 | M Protein | A*24:02 | 10 | QWNLVIGFLF | 0.28076 | Consensus (ann/smm) | 0.14 | 99.16% (473/477) | Non-Toxin |
| 494 | M Protein | A*23:01 | 9 | RFLYIIKLI | 0.05908 | Consensus (ann/smm) | 0.47 | 98.74% (471/477) | Non-Toxin |
| 495 | M Protein | A*23:01 | 10 | RFLYIIKLIF | 0.11728 | Consensus (ann/smm) | 0.12 | 98.74% (471/477) | Non-Toxin |
| 496 | M Protein | A*24:02 | 10 | RFLYIIKLIF | 0.11728 | Consensus (ann/smm) | 0.45 | 98.74% (471/477) | Non-Toxin |
| 497 | M Protein | A*31:01 | 9 | RIAGHHLGR | 0.11919 | Consensus (ann/smm) | 0.46 | 99.58% (475/477) | Non-Toxin |
| 498 | M Protein | A*32:01 | 11 | RINWITGGIAI | 0.58827 | ann | 0.33 | 99.37% (474/477) | Non-Toxin |
| 499 | M Protein | A*32:01 | 11 | RINWITGGIAI | 0.58827 | ann | 0.33 | 99.37% (474/477) | Non-Toxin |
| 500 | M Protein | B*08:01 | 9 | RLFARTRSM | 0.11133 | Consensus (ann/comblib_sidney2008/smm) | 0.3 | 99.79% (476/477) | Non-Toxin |
| 501 | M Protein | A*32:01 | 9 | RLFARTRSM | 0.11133 | Consensus (ann/comblib_sidney2008/smm) | 0.4 | 99.79% (476/477) | Non-Toxin |
| 502 | M Protein | B*15:01 | 9 | RLFARTRSM | 0.11133 | Consensus (ann/comblib_sidney2008/smm) | 0.5 | 99.79% (476/477) | Non-Toxin |
| 503 | M Protein | A*30:01 | 9 | RNRFLYIIK | 0.3104 | Consensus (ann/comblib_sidney2008/smm) | 0.2 | 98.95% (472/477) | Non-Toxin |
| 504 | M Protein | A*32:01 | 8 | RTRSMWSF | -0.25178 | ann | 0.1 | 99.58% (475/477) | Non-Toxin |
| 505 | M Protein | A*30:01 | 8 | RTRSMWSF | -0.25178 | ann | 0.29 | 99.58% (475/477) | Non-Toxin |
| 506 | M Protein | B*57:01 | 8 | RTRSMWSF | -0.25178 | ann | 0.39 | 99.58% (475/477) | Non-Toxin |
| 507 | M Protein | A*30:01 | 9 | RTRSMWSFN | -0.18338 | Consensus (ann/comblib_sidney2008/smm) | 0.3 | 99.37% (474/477) | Non-Toxin |
| 508 | M Protein | A*30:01 | 8 | RYRIGNYK | 0.17451 | ann | 0.19 | 98.53% (470/477) | Non-Toxin |
| 509 | M Protein | A*23:01 | 9 | RYRIGNYKL | 0.04851 | Consensus (ann/smm) | 0.4 | 98.53% (470/477) | Non-Toxin |
| 510 | M Protein | A*30:02 | 9 | SGFAAYSRY | 0.00261 | Consensus (ann/smm) | 0.41 | 98.32% (469/477) | Non-Toxin |
| 511 | M Protein | A*31:01 | 10 | SGFAAYSRYR | -0.00234 | Consensus (ann/smm) | 0.35 | 98.32% (469/477) | Non-Toxin |
| 512 | M Protein | A*01:01 | 9 | SSDNIALLV | 0.15128 | Consensus (ann/smm) | 0.2 | 99.16% (473/477) | Non-Toxin |
| 513 | M Protein | A*33:01 | 8 | SYFIASFR | 0.15309 | ann | 0.42 | 98.74% (471/477) | Non-Toxin |
| 514 | M Protein | A*24:02 | 9 | SYFIASFRL | 0.18333 | Consensus (ann/smm) | 0.22 | 98.74% (471/477) | Non-Toxin |
| 515 | M Protein | A*23:01 | 9 | SYFIASFRL | 0.18333 | Consensus (ann/smm) | 0.36 | 98.74% (471/477) | Non-Toxin |
| 516 | M Protein | A*23:01 | 10 | SYFIASFRLF | 0.19632 | Consensus (ann/smm) | 0.11 | 98.74% (471/477) | Non-Toxin |
| 517 | M Protein | A*24:02 | 10 | SYFIASFRLF | 0.19632 | Consensus (ann/smm) | 0.11 | 98.74% (471/477) | Non-Toxin |
| 518 | M Protein | A*33:01 | 10 | SYYKLGASQR | -0.36647 | Consensus (ann/smm) | 0.27 | 99.37% (474/477) | Non-Toxin |
| 519 | M Protein | A*02:03 | 9 | TLACFVLAA | 0.12481 | Consensus (ann/smm) | 0.42 | 99.16% (473/477) | Non-Toxin |
| 520 | M Protein | A*02:03 | 10 | TLACFVLAAV | 0.15809 | Consensus (ann/smm) | 0.12 | 98.32% (469/477) | Non-Toxin |
| 521 | M Protein | A*02:01 | 10 | TLACFVLAAV | 0.15809 | Consensus (ann/smm) | 0.21 | 98.32% (469/477) | Non-Toxin |

| | | | | | | | | | |
|---|---|---|---|---|---|---|---|---|---|
| 522 | M Protein | A*30:01 | 9 | TSRTLSYYK | -0.11595 | Consensus (ann/comblib_sidney2008/smm) | 0.3 | 99.16% (473/477) | Non-Toxin |
| 523 | M Protein | A*11:01 | 9 | TSRTLSYYK | -0.11595 | Consensus (ann/smm) | 0.35 | 99.16% (473/477) | Non-Toxin |
| 524 | M Protein | B*15:01 | 10 | TVATSRTLSY | -0.12842 | Consensus (ann/smm) | 0.21 | 98.74% (471/477) | Non-Toxin |
| 525 | M Protein | A*26:01 | 10 | TVATSRTLSY | -0.12842 | Consensus (ann/smm) | 0.22 | 98.74% (471/477) | Non-Toxin |
| 526 | M Protein | A*01:01 | 10 | VATSRTLSYY | -0.21789 | Consensus (ann/smm) | 0.28 | 98.95% (472/477) | Non-Toxin |
| 527 | M Protein | A*30:02 | 10 | VATSRTLSYY | -0.21789 | Consensus (ann/smm) | 0.41 | 98.95% (472/477) | Non-Toxin |
| 528 | M Protein | A*11:01 | 11 | VATSRTLSYYK | -0.23748 | Consensus (ann/smm) | 0.17 | 98.95% (472/477) | Non-Toxin |
| 529 | M Protein | A*01:01 | 9 | WICLLQFAY | -0.02684 | Consensus (ann/smm) | 0.37 | 98.95% (472/477) | Non-Toxin |
| 530 | M Protein | B*15:01 | 9 | WLSYFIASF | 0.11822 | Consensus (ann/comblib_sidney2008/smm) | 0.3 | 97.69% (466/477) | Non-Toxin |
| 531 | M Protein | A*32:01 | 9 | WLSYFIASF | 0.11822 | Consensus (ann/comblib_sidney2008/smm) | 0.5 | 97.69% (466/477) | Non-Toxin |
| 532 | M Protein | A*68:01 | 10 | WLSYFIASFR | 0.15179 | Consensus (ann/smm) | 0.21 | 97.69% (466/477) | Non-Toxin |
| 533 | M Protein | A*33:01 | 10 | WLSYFIASFR | 0.15179 | Consensus (ann/smm) | 0.23 | 97.69% (466/477) | Non-Toxin |
| 534 | M Protein | A*31:01 | 10 | WLSYFIASFR | 0.15179 | Consensus (ann/smm) | 0.4 | 97.69% (466/477) | Non-Toxin |
| 535 | M Protein | B*51:01 | 10 | WPVTLACFVL | 0.15193 | Consensus (ann/smm) | 0.34 | 98.95% (472/477) | Non-Toxin |
| 536 | M Protein | A*01:01 | 9 | YANRNRFLY | 0.18472 | Consensus (ann/smm) | 0.28 | 99.16% (473/477) | Non-Toxin |
| 537 | M Protein | B*35:01 | 9 | YANRNRFLY | 0.18472 | Consensus (ann/comblib_sidney2008/smm) | 0.4 | 99.16% (473/477) | Non-Toxin |
| 538 | M Protein | A*23:01 | 9 | YFIASFRLF | 0.06887 | Consensus (ann/smm) | 0.11 | 99.37% (474/477) | Non-Toxin |
| 539 | M Protein | A*24:02 | 9 | YFIASFRLF | 0.06887 | Consensus (ann/smm) | 0.18 | 99.37% (474/477) | Non-Toxin |
| 540 | M Protein | A*33:01 | 11 | YFIASFRLFAR | 0.19709 | ann | 0.03 | 99.37% (474/477) | Non-Toxin |
| 541 | M Protein | B*58:01 | 9 | YIIKLIFLW | 0.033 | Consensus (ann/comblib_sidney2008/smm) | 0.3 | 98.53% (470/477) | Non-Toxin |
| 542 | M Protein | B*53:01 | 9 | YIIKLIFLW | 0.033 | Consensus (ann/comblib_sidney2008/smm) | 0.3 | 98.53% (470/477) | Non-Toxin |
| 543 | M Protein | B*57:01 | 9 | YIIKLIFLW | 0.033 | Consensus (ann/smm) | 0.48 | 98.53% (470/477) | Non-Toxin |
| 544 | M Protein | A*26:01 | 9 | YSRYRIGNY | 0.21358 | Consensus (ann/smm) | 0.4 | 98.53% (470/477) | Non-Toxin |
| 545 | M Protein | A*30:02 | 9 | YSRYRIGNY | 0.21358 | Consensus (ann/smm) | 0.49 | 98.53% (470/477) | Non-Toxin |
| 546 | M Protein | A*03:01 | 10 | YSRYRIGNYK | 0.21736 | Consensus (ann/smm) | 0.47 | 98.53% (470/477) | Non-Toxin |
| 547 | M Protein | A*33:01 | 9 | YYKLGASQR | -0.21863 | Consensus (ann/smm) | 0.27 | 99.37% (474/477) | Non-Toxin |
| S.No. | ORF | Allele | Length | Peptide | Immunogenicity | Method used | Percentile Rank | Conservancy | Toxicity |
| 548 | N Protein | B*07:02 | 10 | APRITFGGPS | 0.35654 | Consensus (ann/smm) | 0.39 | 96.79% (482/498) | Non-Toxin |
| 549 | N Protein | B*15:01 | 10 | AQFAPSASAF | -0.17446 | Consensus (ann/smm) | 0.16 | 97.59% (486/498) | Non-Toxin |
| 550 | N Protein | B*15:01 | 11 | AQFAPSASAFF | -0.11074 | ann | 0.12 | 97.59% (486/498) | Non-Toxin |
| 551 | N Protein | A*11:01 | 9 | ASAFFGMSR | 0.03154 | Consensus (ann/smm) | 0.38 | 97.59% (486/498) | Non-Toxin |
| 552 | N Protein | A*26:01 | 10 | DLSPRWYFYY | 0.2944 | Consensus (ann/smm) | 0.31 | 99.60% (496/498) | Non-Toxin |
| 553 | N Protein | A*01:01 | 10 | DLSPRWYFYY | 0.2944 | Consensus (ann/smm) | 0.35 | 99.60% (496/498) | Non-Toxin |
| 554 | N Protein | A*26:01 | 9 | ELIRQGTDY | 0.0601 | Consensus (ann/smm) | 0.39 | 97.39% (485/498) | Non-Toxin |
| 555 | N Protein | B*35:01 | 9 | FAPSASAFF | -0.18628 | Consensus (ann/comblib_sidney2008/smm) | 0.4 | 97.59% (486/498) | Non-Toxin |
| 556 | N Protein | A*68:01 | 9 | FTALTQHGK | -0.0226 | Consensus (ann/smm) | 0.18 | 99.40% (495/498) | Non-Toxin |
| 557 | N Protein | A*02:03 | 9 | GMSRIGMEV | 0.07018 | Consensus (ann/smm) | 0.46 | 97.59% (486/498) | Non-Toxin |
| 558 | N Protein | A*31:01 | 9 | GYYRRATRR | 0.20111 | Consensus (ann/smm) | 0.45 | 99.40% (495/498) | Non-Toxin |
| 559 | N Protein | B*15:01 | 11 | IAQFAPSASAF | -0.13353 | ann | 0.4 | 97.39% (485/498) | Non-Toxin |
| 560 | N Protein | A*31:01 | 9 | IGYYRRATR | 0.1499 | Consensus (ann/smm) | 0.38 | 99.40% (495/498) | Non-Toxin |
| 561 | N Protein | A*32:01 | 9 | KAYNVTQAF | -0.00587 | Consensus (ann/comblib_sidney2008/smm) | 0.2 | 97.59% (486/498) | Non-Toxin |
| 562 | N Protein | B*58:01 | 9 | KAYNVTQAF | -0.00587 | Consensus (ann/comblib_sidney2008/smm) | 0.4 | 97.59% (486/498) | Non-Toxin |
| 563 | N Protein | A*30:02 | 10 | KDLSPRWYFY | 0.14332 | Consensus (ann/smm) | 0.47 | 99.60% (496/498) | Non-Toxin |
| 564 | N Protein | A*31:01 | 8 | KMKDLSPR | -0.22357 | ann | 0.15 | 99.60% (496/498) | Non-Toxin |
| 565 | N Protein | A*30:02 | 10 | KMKDLSPRWY | -0.05692 | Consensus (ann/smm) | 0.33 | 99.60% (496/498) | Non-Toxin |
| 566 | N Protein | B*07:02 | 9 | KPRQKRTAT | -0.20542 | Consensus (ann/comblib_sidney2008/smm) | 0.1 | 97.79% (487/498) | Non-Toxin |
| 567 | N Protein | B*07:02 | 10 | KPRQKRTATK | -0.16712 | Consensus (ann/smm) | 0.2 | 97.79% (487/498) | Non-Toxin |
| 568 | N Protein | A*03:01 | 9 | KSAAEASKK | -0.07922 | Consensus (ann/smm) | 0.45 | 97.79% (487/498) | Non-Toxin |
| 569 | N Protein | A*11:01 | 9 | KTFPPTEPK | 0.1306 | Consensus (ann/smm) | 0.11 | 96.99% (483/498) | Non-Toxin |
| 570 | N Protein | A*30:01 | 9 | KTFPPTEPK | 0.1306 | Consensus (ann/comblib_sidney2008/smm) | 0.2 | 96.99% (483/498) | Non-Toxin |
| 571 | N Protein | A*03:01 | 9 | KTFPPTEPK | 0.1306 | Consensus (ann/smm) | 0.23 | 96.99% (483/498) | Non-Toxin |

| | | | | | | | | | |
|---|---|---|---|---|---|---|---|---|---|
| 572 | N Protein | A*11:01 | 10 | KTFPPTEPKK | 0.01273 | Consensus (ann/smm) | 0.28 | 96.99% (483/498) | Non-Toxin |
| 573 | N Protein | B*08:01 | 9 | LLLDRLNQL | -0.01446 | Consensus (ann/comblib_sidney2008/smm) | 0.4 | 97.59% (486/498) | Non-Toxin |
| 574 | N Protein | B*15:01 | 9 | LLNKHIDAY | -0.02074 | Consensus (ann/comblib_sidney2008/smm) | 0.3 | 97.59% (486/498) | Non-Toxin |
| 575 | N Protein | A*03:01 | 10 | LLNKHIDAYK | -0.00626 | Consensus (ann/smm) | 0.43 | 97.59% (486/498) | Non-Toxin |
| 576 | N Protein | B*35:01 | 9 | LPAADLDDF | 0.09491 | Consensus (ann/comblib_sidney2008/smm) | 0.5 | 97.59% (486/498) | Non-Toxin |
| 577 | N Protein | B*53:01 | 9 | LPNNTASWF | 0.05582 | Consensus (ann/comblib_sidney2008/smm) | 0.3 | 99.40% (495/498) | Non-Toxin |
| 578 | N Protein | B*51:01 | 10 | LPYGANKDGI | -0.09978 | Consensus (ann/smm) | 0.23 | 99.20% (494/498) | Non-Toxin |
| 579 | N Protein | A*01:01 | 9 | LSPRWYFYY | 0.35734 | Consensus (ann/smm) | 0.22 | 99.60% (496/498) | Non-Toxin |
| 580 | N Protein | B*44:02 | 9 | MEVTPSGTW | -0.06279 | Consensus (ann/smm) | 0.06 | 97.39% (485/498) | Non-Toxin |
| 581 | N Protein | B*44:03 | 9 | MEVTPSGTW | -0.06279 | Consensus (ann/smm) | 0.13 | 97.39% (485/498) | Non-Toxin |
| 582 | N Protein | B*40:01 | 10 | MEVTPSGTWL | 0.07464 | Consensus (ann/smm) | 0.2 | 97.39% (485/498) | Non-Toxin |
| 583 | N Protein | A*01:01 | 11 | NSSPDDQIGYY | 0.01726 | Consensus (ann/smm) | 0.49 | 99.40% (495/498) | Non-Toxin |
| 584 | N Protein | A*68:02 | 9 | NTASWFTAL | 0.22775 | Consensus (ann/comblib_sidney2008/smm) | 0.4 | 99.60% (496/498) | Non-Toxin |
| 585 | N Protein | A*68:02 | 10 | NTASWFTALT | 0.23901 | Consensus (ann/smm) | 0.48 | 99.40% (495/498) | Non-Toxin |
| 586 | N Protein | B*44:03 | 10 | QELIRQGTDY | 0.14554 | Consensus (ann/smm) | 0.25 | 97.19% (484/498) | Non-Toxin |
| 587 | N Protein | B*44:02 | 10 | QELIRQGTDY | 0.14554 | Consensus (ann/smm) | 0.28 | 97.19% (484/498) | Non-Toxin |
| 588 | N Protein | B*07:02 | 10 | RPQGLPNNTA | -0.01397 | Consensus (ann/smm) | 0.46 | 99.40% (495/498) | Non-Toxin |
| 589 | N Protein | B*15:01 | 10 | RQKRTATKAY | -0.06462 | Consensus (ann/smm) | 0.2 | 97.79% (487/498) | Non-Toxin |
| 590 | N Protein | A*30:02 | 10 | RQKRTATKAY | -0.06462 | Consensus (ann/smm) | 0.28 | 97.79% (487/498) | Non-Toxin |
| 591 | N Protein | A*30:01 | 9 | RSKQRRPQG | -0.16448 | Consensus (ann/comblib_sidney2008/smm) | 0.4 | 99.60% (496/498) | Non-Toxin |
| 592 | N Protein | A*30:01 | 9 | RSRNSSRNS | -0.26664 | Consensus (ann/comblib_sidney2008/smm) | 0.4 | 93.37% (465/498) | Non-Toxin |
| 593 | N Protein | A*68:01 | 10 | SASAFFGMSR | 0.00071 | Consensus (ann/smm) | 0.31 | 97.59% (486/498) | Non-Toxin |
| 594 | N Protein | A*11:01 | 10 | SASAFFGMSR | 0.00071 | Consensus (ann/smm) | 0.47 | 97.59% (486/498) | Non-Toxin |
| 595 | N Protein | B*07:02 | 9 | SPRWYFYYL | 0.34101 | Consensus (ann/comblib_sidney2008/smm) | 0.2 | 99.60% (496/498) | Non-Toxin |
| 596 | N Protein | B*08:01 | 9 | SPRWYFYYL | 0.34101 | Consensus (ann/comblib_sidney2008/smm) | 0.2 | 99.60% (496/498) | Non-Toxin |
| 597 | N Protein | A*01:01 | 10 | SSPDDQIGYY | 0.07924 | Consensus (ann/smm) | 0.2 | 99.40% (495/498) | Non-Toxin |
| 598 | N Protein | A*30:02 | 10 | SSPDDQIGYY | 0.07924 | Consensus (ann/smm) | 0.42 | 99.40% (495/498) | Non-Toxin |
| 599 | N Protein | B*35:01 | 9 | TPSGTWLTY | 0.24003 | Consensus (ann/comblib_sidney2008/smm) | 0.2 | 97.39% (485/498) | Non-Toxin |
| 600 | N Protein | B*53:01 | 9 | TPSGTWLTY | 0.24003 | Consensus (ann/comblib_sidney2008/smm) | 0.3 | 97.39% (485/498) | Non-Toxin |
| 601 | N Protein | B*53:01 | 10 | YGANKDGIIW | 0.03977 | Consensus (ann/smm) | 0.24 | 99.40% (495/498) | Non-Toxin |
| 602 | N Protein | A*24:02 | 9 | YYRRATRRI | 0.21744 | Consensus (ann/smm) | 0.26 | 99.40% (495/498) | Non-Toxin |
| 603 | N Protein | A*33:01 | 10 | YYRRATRRIR | 0.31494 | Consensus (ann/smm) | 0.32 | 99.40% (495/498) | Non-Toxin |
| S.No. | ORF | Allele | Length | Peptide | Immunogenicity | Method used | Percentile Rank | Conservancy | Toxicity |
| 604 | ORF10 | A*02:06 | 10 | FAFPFTIYSL | 0.20414 | Consensus (ann/smm) | 0.48 | 99.79% (478/479) | Non-Toxin |
| 605 | ORF10 | B*51:01 | 8 | FPFTIYSL | 0.06356 | Consensus (ann/smm) | 0.39 | 100.00% (479/479) | Non-Toxin |
| 606 | ORF10 | B*51:01 | 9 | FPFTIYSLL | 0.05708 | Consensus (ann/comblib_sidney2008/smm) | 0.3 | 100.00% (479/479) | Non-Toxin |
| 607 | ORF10 | B*53:01 | 9 | FPFTIYSLL | 0.05708 | Consensus (ann/comblib_sidney2008/smm) | 0.5 | 100.00% (479/479) | Non-Toxin |
| 608 | ORF10 | B*53:01 | 10 | FPFTIYSLLL | 0.03149 | Consensus (ann/smm) | 0.13 | 100.00% (479/479) | Non-Toxin |
| 609 | ORF10 | B*51:01 | 10 | FPFTIYSLLL | 0.03149 | Consensus (ann/smm) | 0.14 | 100.00% (479/479) | Non-Toxin |
| 610 | ORF10 | A*68:01 | 10 | FTIYSLLLCR | -0.18372 | Consensus (ann/smm) | 0.12 | 100.00% (479/479) | Non-Toxin |
| 611 | ORF10 | A*33:01 | 10 | FTIYSLLLCR | -0.18372 | Consensus (ann/smm) | 0.35 | 100.00% (479/479) | Non-Toxin |
| 612 | ORF10 | A*23:01 | 10 | GYINVFAFPF | 0.32004 | Consensus (ann/smm) | 0.11 | 99.37% (476/479) | Non-Toxin |
| 613 | ORF10 | A*24:02 | 10 | GYINVFAFPF | 0.32004 | Consensus (ann/smm) | 0.13 | 99.37% (476/479) | Non-Toxin |
| 614 | ORF10 | B*35:01 | 9 | MGYINVFAF | 0.28694 | Consensus (ann/comblib_sidney2008/smm) | 0.1 | 99.37% (476/479) | Non-Toxin |
| 615 | ORF10 | A*23:01 | 9 | MGYINVFAF | 0.28694 | Consensus (ann/smm) | 0.5 | 99.37% (476/479) | Non-Toxin |
| 616 | ORF10 | A*23:01 | 11 | MGYINVFAFPF | 0.40977 | Consensus (ann/smm) | 0.41 | 99.37% (476/479) | Non-Toxin |
| 617 | ORF10 | A*68:02 | 9 | NVFAFPFTI | 0.30241 | Consensus (ann/comblib_sidney2008/smm) | 0.4 | 99.58% (477/479) | Non-Toxin |
| 618 | ORF10 | A*32:01 | 9 | NVFAFPFTI | 0.30241 | Consensus (ann/comblib_sidney2008/smm) | 0.5 | 99.58% (477/479) | Non-Toxin |
| 619 | ORF10 | A*26:01 | 10 | NVFAFPFTIY | 0.40129 | Consensus (ann/smm) | 0.48 | 99.58% (477/479) | Non-Toxin |
| 620 | ORF10 | A*68:01 | 9 | TIYSLLLCR | -0.22977 | Consensus (ann/smm) | 0.47 | 100.00% (479/479) | Non-Toxin |
| 621 | ORF10 | B*53:01 | 9 | YINVFAFPF | 0.28259 | Consensus (ann/comblib_sidney2008/smm) | 0.35 | 99.37% (476/479) | Non-Toxin |

| | | | | | | | | | | |
|---|---|---|---|---|---|---|---|---|---|---|
| 622 | ORF10 | A*32:01 | 9 | YINVFAFPF | 0.28259 | Consensus (ann/comblib_sidney2008/smm) | 0.4 | 99.37% (476/479) | Non-Toxin | |
| S.No. | ORF | Allele | Length | Peptide | Immunogenicity | Method used | Percentile Rank | Conservancy | Toxicity | NSP |
| 623 | ORF-1ab | A*30:02 | 10 | ASFYYVWKSY | 0.00073 | Consensus (ann/smm) | 0.07 | 99.77%(452/453) | Non-Toxin | nsp3 |
| 624 | ORF-1ab | A*30:02 | 10 | CANGQVFGLY | 0.09172 | Consensus (ann/smm) | 0.1 | 99.55%(451/453) | Non-Toxin | nsp13 |
| 625 | ORF-1ab | A*30:02 | 10 | CANGQVFGLY | 0.09172 | Consensus (ann/smm) | 0.1 | 99.55%(451/453) | Non-Toxin | nsp13 |
| 626 | ORF-1ab | A*33:01 | 11 | CLAYYFMRFRR | 0.12614 | ann | 0.07 | 100%(453/453) | Non-Toxin | nsp4 |
| 627 | ORF-1ab | A*01:01 | 9 | CTDDNALAY | 0.07355 | Consensus (ann/smm) | 0.06 | 99.77%(452/453) | Non-Toxin | nsp9 |
| 628 | ORF-1ab | A*01:01 | 10 | CTDDNALAYY | 0.08174 | Consensus (ann/smm) | 0.06 | 99.77%(452/453) | Non-Toxin | nsp9 |
| 629 | ORF-1ab | B*44:03 | 9 | DEWSMATYY | -0.19814 | Consensus (ann/smm) | 0.07 | 99.55%(451/453) | Non-Toxin | nsp3 |
| 630 | ORF-1ab | A*26:01 | 9 | ETISLAGSY | -0.1653 | Consensus (ann/smm) | 0.1 | 100%(453/453) | Non-Toxin | nsp3 |
| 631 | ORF-1ab | A*68:01 | 10 | ETISLAGSYK | -0.20585 | Consensus (ann/smm) | 0.06 | 100%(453/453) | Non-Toxin | nsp3 |
| 632 | ORF-1ab | B*35:01 | 9 | FAVDAAKAY | -0.04849 | Consensus (ann/comblib_sidney2008/smm) | 0.1 | 98.89%(448/453) | Non-Toxin | nsp10 |
| 633 | ORF-1ab | A*02:01 | 10 | FLFVAAIFYL | 0.37766 | Consensus (ann/smm) | 0.06 | 99.77%(452/453) | Non-Toxin | nsp4 |
| 634 | ORF-1ab | A*02:03 | 9 | FLNGSCGSV | -0.24791 | Consensus (ann/smm) | 0.06 | 99.55%(451/453) | Non-Toxin | nsp5 |
| 635 | ORF-1ab | B*53:01 | 10 | FPLCANGQVF | -0.06779 | Consensus (ann/smm) | 0.1 | 99.55%(451/453) | Non-Toxin | nsp13 |
| 636 | ORF-1ab | B*53:01 | 10 | FPLCANGQVF | -0.06779 | Consensus (ann/smm) | 0.1 | 99.55%(451/453) | Non-Toxin | nsp13 |
| 637 | ORF-1ab | A*33:01 | 11 | FYWFFSNYLKR | -0.00254 | ann | 0.04 | 98.67%(447/453) | Non-Toxin | nsp4 |
| 638 | ORF-1ab | B*53:01 | 10 | IPLMYKGLPW | -0.37784 | Consensus (ann/smm) | 0.07 | 100%(453/453) | Non-Toxin | nsp14 |
| 639 | ORF-1ab | B*53:01 | 10 | IPLMYKGLPW | -0.37784 | Consensus (ann/smm) | 0.07 | 100%(453/453) | Non-Toxin | nsp14 |
| 640 | ORF-1ab | B*07:02 | 9 | IPRRNVATL | 0.15714 | Consensus (ann/comblib_sidney2008/smm) | 0.1 | 100%(453/453) | Non-Toxin | nsp13 |
| 641 | ORF-1ab | B*07:02 | 9 | IPRRNVATL | 0.15714 | Consensus (ann/comblib_sidney2008/smm) | 0.1 | 100%(453/453) | Non-Toxin | nsp13 |
| 642 | ORF-1ab | B*58:01 | 8 | ISNSWLMW | -0.11151 | ann | 0.05 | 100%(453/453) | Non-Toxin | nsp3 |
| 643 | ORF-1ab | A*30:02 | 9 | KMNYQVNGY | -0.06542 | Consensus (ann/smm) | 0.07 | 100%(453/453) | Non-Toxin | nsp14 |
| 644 | ORF-1ab | A*30:02 | 9 | KMNYQVNGY | -0.06542 | Consensus (ann/smm) | 0.07 | 100%(453/453) | Non-Toxin | nsp14 |
| 645 | ORF-1ab | A*30:01 | 9 | KVKYLYFIK | 0.08856 | Consensus (ann/comblib_sidney2008/smm) | 0.1 | 100%(453/453) | Non-Toxin | nsp9 |
| 646 | ORF-1ab | B*15:01 | 9 | LMNVLTLVY | 0.07994 | Consensus (ann/comblib_sidney2008/smm) | 0.1 | 98.01%(444/453) | Non-Toxin | nsp6 |
| 647 | ORF-1ab | B*35:01 | 9 | LPSLATVAY | 0.06748 | Consensus (ann/comblib_sidney2008/smm) | 0.1 | 98.01%(444/453) | Non-Toxin | nsp6 |
| 648 | ORF-1ab | B*53:01 | 10 | LPVNVAFELW | 0.27341 | Consensus (ann/smm) | 0.06 | 100%(453/453) | Non-Toxin | nsp15 |
| 649 | ORF-1ab | A*33:01 | 11 | MYKGLPWNVVR | 0.21107 | ann | 0.06 | 100%(453/453) | Non-Toxin | nsp14 |
| 650 | ORF-1ab | A*33:01 | 11 | MYKGLPWNVVR | 0.21107 | ann | 0.06 | 100%(453/453) | Non-Toxin | nsp14 |
| 651 | ORF-1ab | B*44:02 | 9 | QEILGTVSW | 0.03976 | Consensus (ann/smm) | 0.06 | 100%(453/453) | Non-Toxin | nsp3 |
| 652 | ORF-1ab | A*32:01 | 9 | RMYIFFASF | 0.29328 | Consensus (ann/comblib_sidney2008/smm) | 0.1 | 99.11%(449/453) | Non-Toxin | nsp3 |
| 653 | ORF-1ab | B*15:01 | 9 | RMYIFFASF | 0.29328 | Consensus (ann/comblib_sidney2008/smm) | 0.1 | 99.11%(449/453) | Non-Toxin | nsp3 |
| 654 | ORF-1ab | A*30:02 | 10 | RMYIFFASFY | 0.32633 | Consensus (ann/smm) | 0.06 | 99.11%(449/453) | Non-Toxin | nsp3 |
| 655 | ORF-1ab | A*03:01 | 10 | RMYIFFASFY | 0.32633 | Consensus (ann/smm) | 0.1 | 99.11%(449/453) | Non-Toxin | nsp3 |
| 656 | ORF-1ab | A*30:02 | 10 | RYFRLTLGVY | 0.15936 | Consensus (ann/smm) | 0.06 | 100%(453/453) | Non-Toxin | nsp6 |
| 657 | ORF-1ab | B*44:02 | 11 | SEMVMCGGSLY | -0.32016 | ann | 0.03 | 100%(453/453) | Non-Toxin | nsp12 |
| 658 | ORF-1ab | A*11:01 | 10 | SIINNTVYTK | 0.12661 | Consensus (ann/smm) | 0.06 | 100%(453/453) | Non-Toxin | nsp15 |
| 659 | ORF-1ab | A*11:01 | 9 | STFNVPMEK | -0.02845 | Consensus (ann/smm) | 0.06 | 99.77%(452/453) | Non-Toxin | nsp3 |
| 660 | ORF-1ab | A*30:02 | 9 | STNVTIATY | 0.25822 | Consensus (ann/smm) | 0.09 | 99.33%(450/453) | Non-Toxin | nsp3 |
| 661 | ORF-1ab | A*32:01 | 8 | TYNLWNTF | 0.22911 | ann | 0.1 | 98.67%(447/453) | Non-Toxin | nsp14 |
| 662 | ORF-1ab | B*35:01 | 9 | VPFWITIAY | 0.56221 | Consensus (ann/comblib_sidney2008/smm) | 0.1 | 100%(453/453) | Non-Toxin | nsp4 |
| 663 | ORF-1ab | B*08:01 | 10 | YAYLRKHFSM | -0.13937 | Consensus (ann/smm) | 0.07 | 100%(453/453) | Non-Toxin | nsp12 |
| 664 | ORF-1ab | A*02:06 | 10 | YIFFASFYYV | 0.13772 | Consensus (ann/smm) | 0.07 | 99.55%(451/453) | Non-Toxin | nsp3 |
| 665 | ORF-1ab | A*02:06 | 10 | YILFTRFFYV | 0.40924 | Consensus (ann/smm) | 0.07 | 100%(453/453) | Non-Toxin | nsp3 |
| 666 | ORF-1ab | B*53:01 | 10 | YVMHANYIFW | 0.18459 | Consensus (ann/smm) | 0.1 | 98.89%(448/453) | Non-Toxin | nsp16 |
| 667 | ORF-1ab | A*33:01 | 8 | YYFMRFRR | 0.06558 | ann | 0.1 | 100%(453/453) | Non-Toxin | nsp4 |
| S.No. | ORF | Allele | Length | Peptide | Immunogenicity | Method used | Percentile Rank | Conservancy | Toxicity | |
| 668 | ORF3a | B*44:02 | 11 | AGLEAPFLYLY | 0.21841 | ann | 0.41 | 96.67%(465/481) | Non-Toxin | |
| 669 | ORF3a | A*02:03 | 9 | ALSKGVHFV | -0.10314 | Consensus (ann/smm) | 0.12 | 99.67%(477/481) | Non-Toxin | |
| 670 | ORF3a | A*02:01 | 9 | ALSKGVHFV | -0.10314 | Consensus (ann/comblib_sidney2008/smm) | 0.4 | 99.67%(477/481) | Non-Toxin | |

| | | | | | | | | | |
|---|---|---|---|---|---|---|---|---|---|
| 671 | ORF3a | B*07:02 | 9 | APFLYLYAL | 0.03254 | Consensus (ann/comblib_sidney2008/smm) | 0.5 | 96.88%(466/481) | Non-Toxin |
| 672 | ORF3a | A*30:01 | 9 | ASKIITLKK | 0.0947 | Consensus (ann/comblib_sidney2008/smm) | 0.4 | 99.37%(478/481) | Non-Toxin |
| 673 | ORF3a | A*11:01 | 9 | ASKIITLKK | 0.0947 | Consensus (ann/smm) | 0.46 | 99.37%(478/481) | Non-Toxin |
| 674 | ORF3a | B*58:01 | 9 | ASLPFGWLI | 0.3116 | Consensus (ann/comblib_sidney2008/smm) | 0.3 | 98.96%(476/481) | Non-Toxin |
| 675 | ORF3a | A*30:02 | 9 | CWHTNCYDY | 0.00235 | Consensus (ann/smm) | 0.38 | 97.29%(468/481) | Non-Toxin |
| 676 | ORF3a | A*68:01 | 9 | DATPSDFVR | -0.01586 | Consensus (ann/smm) | 0.21 | 99.58%(479/481) | Non-Toxin |
| 677 | ORF3a | A*26:01 | 10 | DTGVEHVTFF | 0.32156 | Consensus (ann/smm) | 0.41 | 99.58%(479/481) | Non-Toxin |
| 678 | ORF3a | A*26:01 | 10 | EIKDATPSDF | -0.10888 | Consensus (ann/smm) | 0.28 | 99.58%(479/481) | Non-Toxin |
| 679 | ORF3a | A*01:01 | 9 | FLCWHTNCY | 0.23647 | Consensus (ann/smm) | 0.42 | 97.29%(468/481) | Non-Toxin |
| 680 | ORF3a | A*68:01 | 9 | FLQSINFVR | 0.04236 | Consensus (ann/smm) | 0.23 | 97.08%(467/481) | Non-Toxin |
| 681 | ORF3a | A*33:01 | 9 | FLQSINFVR | 0.04236 | Consensus (ann/smm) | 0.39 | 97.08%(467/481) | Non-Toxin |
| 682 | ORF3a | B*15:01 | 9 | FLYLYALVY | 0.03563 | Consensus (ann/comblib_sidney2008/smm) | 0.42 | 96.88%(466/481) | Non-Toxin |
| 683 | ORF3a | A*01:01 | 9 | FLYLYALVY | 0.03563 | Consensus (ann/smm) | 0.45 | 96.88%(466/481) | Non-Toxin |
| 684 | ORF3a | A*23:01 | 10 | FLYLYALVYF | 0.04438 | Consensus (ann/smm) | 0.27 | 96.88%(466/481) | Non-Toxin |
| 685 | ORF3a | A*02:03 | 10 | FMRIFTIGTV | 0.47908 | Consensus (ann/smm) | 0.17 | 99.37%(478/481) | Non-Toxin |
| 686 | ORF3a | A*68:01 | 9 | FTIGTVTLK | 0.18024 | Consensus (ann/smm) | 0.16 | 99.37%(478/481) | Non-Toxin |
| 687 | ORF3a | A*11:01 | 9 | FTIGTVTLK | 0.18024 | Consensus (ann/smm) | 0.37 | 99.37%(478/481) | Non-Toxin |
| 688 | ORF3a | A*26:01 | 9 | FVCNLLLLF | -0.06109 | Consensus (ann/smm) | 0.36 | 97.92%(471/481) | Non-Toxin |
| 689 | ORF3a | A*02:06 | 10 | FVCNLLLLFV | 0.00299 | Consensus (ann/smm) | 0.21 | 97.50%(469/481) | Non-Toxin |
| 690 | ORF3a | A*02:01 | 10 | FVCNLLLLFV | 0.00299 | Consensus (ann/smm) | 0.42 | 97.50%(469/481) | Non-Toxin |
| 691 | ORF3a | B*57:01 | 9 | FVRIIMRLW | 0.15222 | Consensus (ann/smm) | 0.27 | 96.67%(465/481) | Non-Toxin |
| 692 | ORF3a | A*01:01 | 10 | GLEAPFLYLY | 0.15503 | Consensus (ann/smm) | 0.43 | 97.08%(467/481) | Non-Toxin |
| 693 | ORF3a | A*23:01 | 9 | HFVCNLLLL | -0.07343 | Consensus (ann/smm) | 0.48 | 97.92%(471/481) | Non-Toxin |
| 694 | ORF3a | A*23:01 | 10 | HFVCNLLLLF | -0.08423 | Consensus (ann/smm) | 0.28 | 97.92%(471/481) | Non-Toxin |
| 695 | ORF3a | A*26:01 | 9 | HTIDGSSGV | -0.17703 | Consensus (ann/smm) | 0.23 | 95.42%(459/481) | Non-Toxin |
| 696 | ORF3a | A*68:02 | 9 | HTIDGSSGV | -0.17703 | Consensus (ann/comblib_sidney2008/smm) | 0.3 | 95.42%(459/481) | Non-Toxin |
| 697 | ORF3a | A*68:01 | 9 | HVTFFIYNK | 0.36278 | Consensus (ann/smm) | 0.16 | 99.79%(480/481) | Non-Toxin |
| 698 | ORF3a | A*11:01 | 9 | HVTFFIYNK | 0.36278 | Consensus (ann/smm) | 0.24 | 99.79%(480/481) | Non-Toxin |
| 699 | ORF3a | A*30:01 | 9 | HVTFFIYNK | 0.36278 | Consensus (ann/comblib_sidney2008/smm) | 0.34 | 99.79%(480/481) | Non-Toxin |
| 700 | ORF3a | B*58:01 | 9 | IIMRLWLCW | 0.15193 | Consensus (ann/comblib_sidney2008/smm) | 0.3 | 96.67%(465/481) | Non-Toxin |
| 701 | ORF3a | B*57:01 | 9 | IIMRLWLCW | 0.15193 | Consensus (ann/smm) | 0.49 | 96.67%(465/481) | Non-Toxin |
| 702 | ORF3a | A*03:01 | 10 | IIMRLWLCWK | 0.27346 | Consensus (ann/smm) | 0.13 | 96.67%(465/481) | Non-Toxin |
| 703 | ORF3a | A*11:01 | 10 | IIMRLWLCWK | 0.27346 | Consensus (ann/smm) | 0.36 | 96.67%(465/481) | Non-Toxin |
| 704 | ORF3a | A*03:01 | 9 | IMRLWLCWK | 0.29482 | Consensus (ann/smm) | 0.34 | 96.67%(465/481) | Non-Toxin |
| 705 | ORF3a | A*31:01 | 11 | IMRLWLCWKCR | 0.15305 | Consensus (ann/smm) | 0.41 | 96.67%(465/481) | Non-Toxin |
| 706 | ORF3a | A*33:01 | 11 | IMRLWLCWKCR | 0.15305 | ann | 0.46 | 96.67%(465/481) | Non-Toxin |
| 707 | ORF3a | A*33:01 | 9 | INFVRIIMR | 0.26494 | Consensus (ann/smm) | 0.41 | 96.88%(466/481) | Non-Toxin |
| 708 | ORF3a | B*53:01 | 9 | IPIQASLPF | -0.20683 | Consensus (ann/comblib_sidney2008/smm) | 0.2 | 99.67%(477/481) | Non-Toxin |
| 709 | ORF3a | B*35:01 | 9 | IPIQASLPF | -0.20683 | Consensus (ann/comblib_sidney2008/smm) | 0.2 | 99.67%(477/481) | Non-Toxin |
| 710 | ORF3a | B*07:02 | 9 | IPIQASLPF | -0.20683 | Consensus (ann/comblib_sidney2008/smm) | 0.3 | 99.67%(477/481) | Non-Toxin |
| 711 | ORF3a | B*51:01 | 10 | IPYNSVTSSI | -0.32835 | Consensus (ann/smm) | 0.11 | 97.92%(471/481) | Non-Toxin |
| 712 | ORF3a | B*07:02 | 10 | IPYNSVTSSI | -0.32835 | Consensus (ann/smm) | 0.46 | 97.92%(471/481) | Non-Toxin |
| 713 | ORF3a | B*53:01 | 9 | IQASLPFGW | -0.05641 | Consensus (ann/comblib_sidney2008/smm) | 0.4 | 98.96%(476/481) | Non-Toxin |
| 714 | ORF3a | B*58:01 | 9 | IQASLPFGW | -0.05641 | Consensus (ann/comblib_sidney2008/smm) | 0.5 | 98.96%(476/481) | Non-Toxin |
| 715 | ORF3a | B*15:01 | 10 | IVGVALLAVF | 0.12654 | Consensus (ann/smm) | 0.32 | 99.67%(477/481) | Non-Toxin |
| 716 | ORF3a | B*57:01 | 9 | KIITLKKRW | -0.2833 | Consensus (ann/smm) | 0.23 | 99.67%(477/481) | Non-Toxin |
| 717 | ORF3a | B*58:01 | 9 | KIITLKKRW | -0.2833 | Consensus (ann/comblib_sidney2008/smm) | 0.4 | 99.67%(477/481) | Non-Toxin |
| 718 | ORF3a | B*44:03 | 9 | LEAPFLYLY | 0.0955 | Consensus (ann/smm) | 0.24 | 97.29%(468/481) | Non-Toxin |
| 719 | ORF3a | B*44:02 | 9 | LEAPFLYLY | 0.0955 | Consensus (ann/smm) | 0.4 | 97.29%(468/481) | Non-Toxin |
| 720 | ORF3a | B*40:01 | 11 | LEAPFLYLYAL | 0.11044 | ann | 0.49 | 96.88%(466/481) | Non-Toxin |
| 721 | ORF3a | A*02:06 | 10 | LIVGVALLAV | 0.12886 | Consensus (ann/smm) | 0.2 | 99.37%(478/481) | Non-Toxin |

| | | | | | | | | | |
|---|---|---|---|---|---|---|---|---|---|
| 722 | ORF3a | A*03:01 | 10 | LLAVFQSASK | -0.16393 | Consensus (ann/smm) | 0.28 | 77.33%(372/481) | Non-Toxin |
| 723 | ORF3a | B*15:01 | 11 | LLVAAGLEAPF | 0.23386 | ann | 0.28 | 97.08%(467/481) | Non-Toxin |
| 724 | ORF3a | B*51:01 | 10 | LPFGWLIVGV | 0.45692 | Consensus (ann/smm) | 0.22 | 98.96%(476/481) | Non-Toxin |
| 725 | ORF3a | B*15:01 | 10 | LVAAGLEAPF | 0.19506 | Consensus (ann/smm) | 0.27 | 97.08%(467/481) | Non-Toxin |
| 726 | ORF3a | A*23:01 | 9 | LYLYALVYF | 0.05302 | Consensus (ann/smm) | 0.11 | 96.88%(466/481) | Non-Toxin |
| 727 | ORF3a | A*24:02 | 9 | LYLYALVYF | 0.05302 | Consensus (ann/smm) | 0.14 | 96.88%(466/481) | Non-Toxin |
| 728 | ORF3a | A*23:01 | 10 | LYLYALVYFL | 0.12412 | Consensus (ann/smm) | 0.18 | 96.88%(466/481) | Non-Toxin |
| 729 | ORF3a | A*24:02 | 10 | LYLYALVYFL | 0.12412 | Consensus (ann/smm) | 0.35 | 96.88%(466/481) | Non-Toxin |
| 730 | ORF3a | A*32:01 | 11 | MRIFTIGTVTL | 0.45546 | ann | 0.35 | 99.37%(478/481) | Non-Toxin |
| 731 | ORF3a | A*02:06 | 9 | NLLLLFVTV | 0.14216 | Consensus (ann/smm) | 0.49 | 97.50%(469/481) | Non-Toxin |
| 732 | ORF3a | B*58:01 | 8 | QASLPFGW | 0.06314 | ann | 0.35 | 98.96%(476/481) | Non-Toxin |
| 733 | ORF3a | A*68:01 | 10 | QSASKIITLK | -0.08261 | Consensus (ann/smm) | 0.4 | 77.33%(372/481) | Non-Toxin |
| 734 | ORF3a | A*32:01 | 11 | RIFTIGTVTLK | 0.3426 | ann | 0.36 | 99.37%(478/481) | Non-Toxin |
| 735 | ORF3a | A*31:01 | 9 | RLWLCWKCR | 0.00325 | Consensus (ann/smm) | 0.3 | 96.88%(466/481) | Non-Toxin |
| 736 | ORF3a | A*11:01 | 9 | SASKIITLK | 0.01046 | Consensus (ann/smm) | 0.3 | 99.37%(478/481) | Non-Toxin |
| 737 | ORF3a | A*68:01 | 9 | SASKIITLK | 0.01046 | Consensus (ann/smm) | 0.4 | 99.37%(478/481) | Non-Toxin |
| 738 | ORF3a | A*11:01 | 10 | SASKIITLKK | -0.11032 | Consensus (ann/smm) | 0.35 | 99.37%(478/481) | Non-Toxin |
| 739 | ORF3a | B*44:03 | 10 | SEHDYQIGGY | 0.0901 | Consensus (ann/smm) | 0.19 | 99.58%(479/481) | Non-Toxin |
| 740 | ORF3a | B*44:02 | 10 | SEHDYQIGGY | 0.0901 | Consensus (ann/smm) | 0.38 | 99.58%(479/481) | Non-Toxin |
| 741 | ORF3a | A*68:01 | 10 | SINFVRIIMR | 0.3413 | Consensus (ann/smm) | 0.39 | 96.88%(466/481) | Non-Toxin |
| 742 | ORF3a | A*02:06 | 9 | TQLSTDTGV | -0.05883 | Consensus (ann/smm) | 0.39 | 99.58%(479/481) | Non-Toxin |
| 743 | ORF3a | A*01:01 | 10 | TTSPISEHDY | 0.03815 | Consensus (ann/smm) | 0.22 | 97.92%(471/481) | Non-Toxin |
| 744 | ORF3a | A*02:06 | 9 | TVYSHLLLV | -0.16245 | Consensus (ann/smm) | 0.44 | 97.92%(471/481) | Non-Toxin |
| 745 | ORF3a | B*44:03 | 9 | VEHVTFFIY | 0.3766 | Consensus (ann/smm) | 0.18 | 99.58%(479/481) | Non-Toxin |
| 746 | ORF3a | A*32:01 | 9 | VTFFIYNKI | 0.15046 | Consensus (ann/comblib_sidney2008/smm) | 0.4 | 99.58%(479/481) | Non-Toxin |
| 747 | ORF3a | A*24:02 | 9 | VYFLQSINF | -0.13315 | Consensus (ann/smm) | 0.23 | 96.88%(466/481) | Non-Toxin |
| 748 | ORF3a | A*23:01 | 9 | VYFLQSINF | -0.13315 | Consensus (ann/smm) | 0.24 | 96.88%(466/481) | Non-Toxin |
| 749 | ORF3a | A*33:01 | 11 | VYFLQSINFVR | -0.00063 | ann | 0.43 | 96.88%(466/481) | Non-Toxin |
| 750 | ORF3a | B*40:01 | 11 | WESGVKDCVVL | -0.15959 | ann | 0.21 | 96.25%(463/481) | Non-Toxin |
| 751 | ORF3a | A*01:01 | 10 | YFLCWHTNCY | 0.18893 | Consensus (ann/smm) | 0.48 | 97.08%(467/481) | Non-Toxin |
| 752 | ORF3a | A*33:01 | 10 | YFLQSINFVR | -0.03483 | Consensus (ann/smm) | 0.2 | 96.88%(466/481) | Non-Toxin |
| 753 | ORF3a | A*02:01 | 9 | YLYALVYFL | 0.13151 | Consensus (ann/comblib_sidney2008/smm) | 0.1 | 96.88%(466/481) | Non-Toxin |
| 754 | ORF3a | A*02:03 | 9 | YLYALVYFL | 0.13151 | Consensus (ann/smm) | 0.18 | 96.88%(466/481) | Non-Toxin |
| 755 | ORF3a | A*02:06 | 9 | YLYALVYFL | 0.13151 | Consensus (ann/smm) | 0.37 | 96.88%(466/481) | Non-Toxin |
| 756 | ORF3a | A*24:02 | 9 | YYQLYSTQL | -0.24301 | Consensus (ann/smm) | 0.29 | 99.58%(479/481) | Non-Toxin |
| S.No. | ORF | Allele | Length | Peptide | Immunogenicity | Method used | Percentile Rank | Conservancy | Toxicity |
| 757 | ORF6 | B*40:01 | 8 | AEILLIIM | 0.25884 | Consensus (ann/smm) | 0.41 | 99.58% (479/481) | Non-Toxin |
| 758 | ORF6 | B*44:02 | 11 | AEILLIIMRTF | 0.1815 | ann | 0.06 | 99.58% (479/481) | Non-Toxin |
| 759 | ORF6 | A*02:06 | 9 | FQVTIAEIL | 0.38115 | Consensus (ann/smm) | 0.52 | 99.17% (477/481) | Non-Toxin |
| 760 | ORF6 | A*03:01 | 10 | ILLIIMRTFK | 0.2388 | Consensus (ann/smm) | 0.17 | 99.58% (479/481) | Non-Toxin |
| 761 | ORF6 | A*32:01 | 9 | IMRTFKVSI | -0.09496 | Consensus (ann/comblib_sidney2008/smm) | 0.44 | 99.79% (480/481) | Non-Toxin |
| 762 | ORF6 | A*30:02 | 9 | KVSIWNLDY | 0.29343 | Consensus (ann/smm) | 0.41 | 99.58% (479/481) | Non-Toxin |
| 763 | ORF6 | A*03:01 | 9 | LLIIMRTFK | 0.156 | Consensus (ann/smm) | 0.36 | 99.58% (479/481) | Non-Toxin |
| 764 | ORF6 | A*30:02 | 10 | LSKSLTENKY | -0.24668 | Consensus (ann/smm) | 0.48 | 98.75% (475/481) | Non-Toxin |
| 765 | ORF6 | B*57:01 | 8 | RTFKVSIW | -0.18221 | ann | 0.05 | 99.79% (480/481) | Non-Toxin |
| 766 | ORF6 | B*58:01 | 8 | RTFKVSIW | -0.18221 | ann | 0.07 | 99.79% (480/481) | Non-Toxin |
| 767 | ORF6 | A*32:01 | 8 | RTFKVSIW | -0.18221 | ann | 0.44 | 99.79% (480/481) | Non-Toxin |
| 768 | ORF6 | A*32:01 | 9 | SIWNLDYII | 0.15011 | Consensus (ann/comblib_sidney2008/smm) | 0.2 | 99.58% (479/481) | Non-Toxin |
| 769 | ORF6 | B*58:01 | 9 | VTIAEILLI | 0.28951 | Consensus (ann/comblib_sidney2008/smm) | 0.34 | 99.17% (477/481) | Non-Toxin |
| S.No. | ORF | Allele | Length | Peptide | Immunogenicity | Method used | Percentile Rank | Conservancy | Toxicity |
| 770 | ORF7a | B*44:02 | 10 | ADNKFALTCF | -0.07618 | Consensus (ann/smm) | 0.36 | 98.75% (474/480) | Non-Toxin |

| | | | | | | | | | |
|---|---|---|---|---|---|---|---|---|---|
| 771 | ORF7a | A*03:01 | 10 | CVRGTTVLLK | 0.14952 | Consensus (ann/smm) | 0.17 | 99.58% (478/480) | Non-Toxin |
| 772 | ORF7a | A*26:01 | 9 | EVQELYSPI | -0.09723 | Consensus (ann/smm) | 0.21 | 98.12% (471/480) | Non-Toxin |
| 773 | ORF7a | A*68:02 | 9 | EVQELYSPI | -0.09723 | Consensus (ann/comblib_sidney2008/smm) | 0.5 | 98.12% (471/480) | Non-Toxin |
| 774 | ORF7a | A*26:01 | 10 | EVQELYSPIF | -0.03858 | Consensus (ann/smm) | 0.22 | 98.12% (471/480) | Non-Toxin |
| 775 | ORF7a | B*53:01 | 10 | FALTCFSTQF | -0.09369 | Consensus (ann/smm) | 0.22 | 98.75% (474/480) | Non-Toxin |
| 776 | ORF7a | A*33:01 | 10 | FITLCFTLKR | -0.03588 | Consensus (ann/smm) | 0.32 | 97.92% (470/480) | Non-Toxin |
| 777 | ORF7a | A*68:01 | 10 | FITLCFTLKR | -0.03588 | Consensus (ann/smm) | 0.48 | 97.92% (470/480) | Non-Toxin |
| 778 | ORF7a | B*15:01 | 9 | FLIVAAIVF | 0.29611 | Consensus (ann/comblib_sidney2008/smm) | 0.2 | 98.96% (475/480) | Non-Toxin |
| 779 | ORF7a | A*02:01 | 10 | FLIVAAIVFI | 0.38946 | Consensus (ann/smm) | 0.21 | 98.33% (472/480) | Non-Toxin |
| 780 | ORF7a | A*32:01 | 9 | GTYEGNSPF | -0.01964 | Consensus (ann/comblib_sidney2008/smm) | 0.4 | 98.96% (475/480) | Non-Toxin |
| 781 | ORF7a | A*23:01 | 10 | IFLIVAAIVF | 0.38189 | Consensus (ann/smm) | 0.34 | 98.33% (472/480) | Non-Toxin |
| 782 | ORF7a | B*15:01 | 10 | IFLIVAAIVF | 0.38189 | Consensus (ann/smm) | 0.47 | 98.33% (472/480) | Non-Toxin |
| 783 | ORF7a | A*03:01 | 10 | ITLCFTLKRK | -0.06825 | Consensus (ann/smm) | 0.17 | 97.92% (470/480) | Non-Toxin |
| 784 | ORF7a | A*11:01 | 10 | ITLCFTLKRK | -0.06825 | Consensus (ann/smm) | 0.33 | 97.92% (470/480) | Non-Toxin |
| 785 | ORF7a | A*32:01 | 9 | KIILFLALI | 0.16214 | Consensus (ann/comblib_sidney2008/smm) | 0.4 | 99.58% (478/480) | Non-Toxin |
| 786 | ORF7a | B*08:01 | 9 | MKIILFLAL | 0.29002 | Consensus (ann/comblib_sidney2008/smm) | 0.5 | 99.58% (478/480) | Non-Toxin |
| 787 | ORF7a | B*40:01 | 9 | QELYSPIFL | 0.00186 | Consensus (ann/smm) | 0.19 | 98.33% (472/480) | Non-Toxin |
| 788 | ORF7a | B*44:02 | 10 | QELYSPIFLI | 0.03838 | Consensus (ann/smm) | 0.22 | 98.33% (472/480) | Non-Toxin |
| 789 | ORF7a | B*44:03 | 10 | QELYSPIFLI | 0.03838 | Consensus (ann/smm) | 0.25 | 98.33% (472/480) | Non-Toxin |
| 790 | ORF7a | A*03:01 | 10 | QLRARSVSPK | -0.16177 | Consensus (ann/smm) | 0.16 | 98.33% (472/480) | Non-Toxin |
| 791 | ORF7a | A*30:01 | 8 | RARSVSPK | -0.27456 | ann | 0.11 | 98.33% (472/480) | Non-Toxin |
| 792 | ORF7a | A*30:01 | 9 | RARSVSPKL | -0.40056 | Consensus (ann/comblib_sidney2008/smm) | 0.2 | 98.33% (472/480) | Non-Toxin |
| 793 | ORF7a | B*58:01 | 9 | RSVSPKLFI | -0.30783 | Consensus (ann/comblib_sidney2008/smm) | 0.2 | 98.33% (472/480) | Non-Toxin |
| 794 | ORF7a | A*31:01 | 10 | RSVSPKLFIR | -0.20775 | Consensus (ann/smm) | 0.31 | 98.33% (472/480) | Non-Toxin |
| 795 | ORF7a | B*07:02 | 10 | SPIFLIVAAI | 0.37454 | Consensus (ann/smm) | 0.47 | 98.33% (472/480) | Non-Toxin |
| 796 | ORF7a | A*31:01 | 9 | SVSPKLFIR | -0.10874 | Consensus (ann/smm) | 0.45 | 98.33% (472/480) | Non-Toxin |
| 797 | ORF7a | A*11:01 | 9 | SVSPKLFIR | -0.10874 | Consensus (ann/smm) | 0.46 | 98.33% (472/480) | Non-Toxin |
| 798 | ORF7a | A*01:01 | 10 | TLATCELYHY | 0.1021 | Consensus (ann/smm) | 0.35 | 99.79% (479/480) | Non-Toxin |
| 799 | ORF7a | B*40:01 | 11 | TYEGNSPFHPL | 0.01942 | ann | 0.46 | 98.96% (475/480) | Non-Toxin |
| 800 | ORF7a | A*23:01 | 9 | VFITLCFTL | 0.14219 | Consensus (ann/smm) | 0.3 | 98.33% (472/480) | Non-Toxin |
| 801 | ORF7a | A*24:02 | 9 | VFITLCFTL | 0.14219 | Consensus (ann/smm) | 0.44 | 98.33% (472/480) | Non-Toxin |
| 802 | ORF7a | B*40:01 | 10 | YEGNSPFHPL | -0.03639 | Consensus (ann/smm) | 0.28 | 98.96% (475/480) | Non-Toxin |
| S.No. | ORF | Allele | Length | Peptide | Immunogenicity | Method used | Percentile Rank | Conservancy | Toxicity |
| 803 | ORF7b | A*02:01 | 10 | CFLAFLLFLV | 0.22085 | Consensus (ann/smm) | 0.47 | 98.31% (232/236) | Non-Toxin |
| 804 | ORF7b | A*02:06 | 10 | CFLAFLLFLV | 0.22085 | Consensus (ann/smm) | 0.48 | 98.31% (232/236) | Non-Toxin |
| 805 | ORF7b | A*23:01 | 9 | DFYLCFLAF | 0.05884 | Consensus (ann/smm) | 0.29 | 98.31% (232/236) | Non-Toxin |
| 806 | ORF7b | A*33:01 | 10 | DFYLCFLAFL | 0.14012 | Consensus (ann/smm) | 0.4 | 98.31% (232/236) | Non-Toxin |
| 807 | ORF7b | A*02:03 | 9 | FLAFLLFLV | 0.20158 | Consensus (ann/smm) | 0.07 | 98.31% (232/236) | Non-Toxin |
| 808 | ORF7b | A*02:06 | 9 | FLAFLLFLV | 0.20158 | Consensus (ann/smm) | 0.08 | 98.31% (232/236) | Non-Toxin |
| 809 | ORF7b | A*02:01 | 9 | FLAFLLFLV | 0.20158 | Consensus (ann/comblib_sidney2008/smm) | 0.1 | 98.31% (232/236) | Non-Toxin |
| 810 | ORF7b | A*68:02 | 9 | FLAFLLFLV | 0.20158 | Consensus (ann/comblib_sidney2008/smm) | 0.33 | 98.31% (232/236) | Non-Toxin |
| 811 | ORF7b | A*02:01 | 10 | FLAFLLFLVL | 0.23386 | Consensus (ann/smm) | 0.26 | 98.31% (232/236) | Non-Toxin |
| 812 | ORF7b | A*02:01 | 10 | FLLFLVLIML | 0.14288 | Consensus (ann/smm) | 0.3 | 99.58% (235/236) | Non-Toxin |
| 813 | ORF7b | A*23:01 | 9 | FYLCFLAFL | 0.14713 | Consensus (ann/smm) | 0.36 | 98.31% (232/236) | Non-Toxin |
| 814 | ORF7b | A*24:02 | 9 | FYLCFLAFL | 0.14713 | Consensus (ann/smm) | 0.54 | 98.31% (232/236) | Non-Toxin |
| 815 | ORF7b | A*23:01 | 10 | FYLCFLAFLL | 0.1745 | Consensus (ann/smm) | 0.13 | 98.31% (232/236) | Non-Toxin |
| 816 | ORF7b | A*24:02 | 10 | FYLCFLAFLL | 0.1745 | Consensus (ann/smm) | 0.28 | 98.31% (232/236) | Non-Toxin |
| 817 | ORF7b | A*23:01 | 10 | IDFYLCFLAF | 0.09468 | Consensus (ann/smm) | 0.36 | 98.31% (232/236) | Non-Toxin |
| 818 | ORF7b | B*44:03 | 9 | IELSLIDFY | 0.03153 | Consensus (ann/smm) | 0.19 | 99.58% (235/236) | Non-Toxin |
| 819 | ORF7b | B*44:02 | 9 | IELSLIDFY | 0.03153 | Consensus (ann/smm) | 0.42 | 99.58% (235/236) | Non-Toxin |
| 820 | ORF7b | A*32:01 | 10 | IMLIIFWFSL | 0.58457 | Consensus (ann/smm) | 0.4 | 99.58% (235/236) | Non-Toxin |

| | | | | | | | | | |
|---|---|---|---|---|---|---|---|---|---|
| 821 | ORF7b | B*08:01 | 10 | IMLIIFWFSL | 0.58457 | Consensus (ann/smm) | 0.41 | 99.58% (235/236) | Non-Toxin |
| 822 | ORF7b | A*02:01 | 10 | IMLIIFWFSL | 0.58457 | Consensus (ann/smm) | 0.43 | 99.58% (235/236) | Non-Toxin |
| 823 | ORF7b | A*23:01 | 9 | LFLVLIMLI | -0.00226 | Consensus (ann/smm) | 0.44 | 99.58% (235/236) | Non-Toxin |
| 824 | ORF7b | B*53:01 | 10 | LVLIMLIIFW | 0.25452 | Consensus (ann/smm) | 0.32 | 99.58% (235/236) | Non-Toxin |
| 825 | ORF7b | A*01:01 | 10 | MIELSLIDFY | 0.06184 | Consensus (ann/smm) | 0.35 | 99.58% (235/236) | Non-Toxin |
| 826 | ORF7b | A*02:01 | 9 | MLIIFWFSL | 0.50177 | Consensus (ann/comblib_sidney2008/smm) | 0.2 | 99.58% (235/236) | Non-Toxin |
| 827 | ORF7b | A*02:06 | 9 | MLIIFWFSL | 0.50177 | Consensus (ann/smm) | 0.41 | 99.58% (235/236) | Non-Toxin |
| 828 | ORF7b | A*02:01 | 10 | SLIDFYLCFL | 0.18838 | Consensus (ann/smm) | 0.14 | 98.31% (232/236) | Non-Toxin |
| 829 | ORF7b | A*02:03 | 10 | SLIDFYLCFL | 0.18838 | Consensus (ann/smm) | 0.49 | 98.31% (232/236) | Non-Toxin |
| 830 | ORF7b | A*02:01 | 9 | YLCFLAFLL | 0.21865 | Consensus (ann/comblib_sidney2008/smm) | 0.3 | 98.31% (232/236) | Non-Toxin |
| 831 | ORF7b | A*23:01 | 10 | YLCFLAFLLF | 0.22196 | Consensus (ann/smm) | 0.42 | 98.31% (232/236) | Non-Toxin |
| S.No. | ORF | Allele | Length | Peptide | Immunogenicity | Method used | Percentile Rank | Conservancy | Toxicity |
| 832 | ORF8 | B*53:01 | 9 | CPIHFYSKW | -0.07935 | Consensus (ann/comblib_sidney2008/smm) | 0.2 | 99.38% (477/480) | Non-Toxin |
| 833 | ORF8 | B*51:01 | 9 | CPIHFYSKW | -0.07935 | Consensus (ann/comblib_sidney2008/smm) | 0.8 | 99.38% (477/480) | Non-Toxin |
| 834 | ORF8 | B*53:01 | 10 | CPIHFYSKWY | -0.02216 | Consensus (ann/smm) | 0.2 | 99.38% (477/480) | Non-Toxin |
| 835 | ORF8 | A*01:01 | 10 | CSFYEDFLEY | 0.31272 | Consensus (ann/smm) | 0.24 | 99.79% (479/480) | Non-Toxin |
| 836 | ORF8 | A*30:02 | 10 | CSFYEDFLEY | 0.31272 | Consensus (ann/smm) | 0.8 | 99.79% (479/480) | Non-Toxin |
| 837 | ORF8 | A*33:01 | 9 | DFLEYHDVR | 0.16684 | Consensus (ann/smm) | 0.13 | 99.58% (478/480) | Non-Toxin |
| 838 | ORF8 | A*02:03 | 10 | FLEYHDVRVV | 0.18854 | Consensus (ann/smm) | 0.7 | 99.79% (479/480) | Non-Toxin |
| 839 | ORF8 | B*40:01 | 11 | FLEYHDVRVVL | 0.22976 | ann | 0.93 | 99.79% (479/480) | Non-Toxin |
| 840 | ORF8 | A*02:03 | 8 | FLGIITTV | 0.34382 | Consensus (ann/smm) | 0.51 | 99.17% (476/480) | Non-Toxin |
| 841 | ORF8 | A*02:01 | 8 | FLGIITTV | 0.34382 | Consensus (ann/smm) | 0.72 | 99.17% (476/480) | Non-Toxin |
| 842 | ORF8 | A*02:03 | 9 | FLGIITTVA | 0.36794 | Consensus (ann/smm) | 0.72 | 99.17% (476/480) | Non-Toxin |
| 843 | ORF8 | A*02:03 | 10 | FLGIITTVAA | 0.40656 | Consensus (ann/smm) | 0.85 | 99.17% (476/480) | Non-Toxin |
| 844 | ORF8 | B*15:01 | 11 | FLGIITTVAAF | 0.44486 | ann | 0.7 | 99.17% (476/480) | Non-Toxin |
| 845 | ORF8 | A*02:01 | 11 | FLVFLGIITTV | 0.46372 | Consensus (ann/smm) | 0.46 | 99.17% (476/480) | Non-Toxin |
| 846 | ORF8 | A*68:01 | 9 | FTINCQEPK | -0.04683 | Consensus (ann/smm) | 0.18 | 100.00% (480/480) | Non-Toxin |
| 847 | ORF8 | A*11:01 | 9 | FTINCQEPK | -0.04683 | Consensus (ann/smm) | 0.54 | 100.00% (480/480) | Non-Toxin |
| 848 | ORF8 | A*31:01 | 8 | FYSKWYIR | 0.05384 | ann | 0.38 | 99.38% (477/480) | Non-Toxin |
| 849 | ORF8 | A*33:01 | 8 | FYSKWYIR | 0.05384 | ann | 0.47 | 99.38% (477/480) | Non-Toxin |
| 850 | ORF8 | A*24:02 | 9 | FYSKWYIRV | 0.08408 | Consensus (ann/smm) | 0.8 | 99.38% (477/480) | Non-Toxin |
| 851 | ORF8 | A*30:01 | 9 | GARKSAPLI | -0.34031 | Consensus (ann/comblib_sidney2008/smm) | 0.3 | 99.17% (476/480) | Non-Toxin |
| 852 | ORF8 | B*15:01 | 9 | GIITTVAAF | 0.2148 | Consensus (ann/comblib_sidney2008/smm) | 0.3 | 99.17% (476/480) | Non-Toxin |
| 853 | ORF8 | A*26:01 | 9 | GIITTVAAF | 0.2148 | Consensus (ann/smm) | 0.45 | 99.17% (476/480) | Non-Toxin |
| 854 | ORF8 | B*57:01 | 9 | GSLVVRCSF | -0.0153 | Consensus (ann/smm) | 0.88 | 100.00% (480/480) | Non-Toxin |
| 855 | ORF8 | A*30:02 | 10 | GSLVVRCSFY | 0.00657 | Consensus (ann/smm) | 0.24 | 100.00% (480/480) | Non-Toxin |
| 856 | ORF8 | A*31:01 | 9 | HFYSKWYIR | -0.09452 | Consensus (ann/smm) | 0.11 | 99.38% (477/480) | Non-Toxin |
| 857 | ORF8 | A*33:01 | 9 | HFYSKWYIR | -0.09452 | Consensus (ann/smm) | 0.12 | 99.38% (477/480) | Non-Toxin |
| 858 | ORF8 | A*23:01 | 11 | IGNYTVSCLPF | -0.15551 | Consensus (ann/smm) | 1 | 51.46% (247/480) | Non-Toxin |
| 859 | ORF8 | A*31:01 | 10 | IHFYSKWYIR | -0.05367 | Consensus (ann/smm) | 0.53 | 99.38% (477/480) | Non-Toxin |
| 860 | ORF8 | A*30:02 | 9 | IQYIDIGNY | 0.30442 | Consensus (ann/smm) | 0.12 | 99.38% (477/480) | Non-Toxin |
| 861 | ORF8 | B*15:01 | 9 | IQYIDIGNY | 0.30442 | Consensus (ann/comblib_sidney2008/smm) | 0.34 | 99.38% (477/480) | Non-Toxin |
| 862 | ORF8 | A*32:01 | 11 | KLGSLVVRCSF | -0.17931 | ann | 0.83 | 100.00% (480/480) | Non-Toxin |
| 863 | ORF8 | B*58:01 | 9 | KSAPLIELC | 0.19404 | Consensus (ann/comblib_sidney2008/smm) | 0.4 | 99.17% (476/480) | Non-Toxin |
| 864 | ORF8 | A*31:01 | 9 | KWYIRVGAR | 0.27344 | Consensus (ann/smm) | 0.46 | 99.38% (477/480) | Non-Toxin |
| 865 | ORF8 | B*40:01 | 10 | LEYHDVRVVL | 0.20083 | Consensus (ann/smm) | 0.36 | 99.79% (479/480) | Non-Toxin |
| 866 | ORF8 | B*15:01 | 10 | LGIITTVAAF | 0.34746 | Consensus (ann/smm) | 0.31 | 99.17% (476/480) | Non-Toxin |
| 867 | ORF8 | B*15:01 | 10 | LQSCTQHQPY | -0.25674 | Consensus (ann/smm) | 0.17 | 96.04% (461/480) | Non-Toxin |
| 868 | ORF8 | A*02:03 | 10 | LVFLGIITTV | 0.37016 | Consensus (ann/smm) | 0.48 | 99.17% (476/480) | Non-Toxin |
| 869 | ORF8 | A*02:06 | 10 | LVFLGIITTV | 0.37016 | Consensus (ann/smm) | 0.79 | 99.17% (476/480) | Non-Toxin |
| 870 | ORF8 | B*51:01 | 9 | MKFLVFLGI | 0.18768 | Consensus (ann/comblib_sidney2008/smm) | 0.98 | 99.17% (476/480) | Non-Toxin |

| | | | | | | | | | |
|---|---|---|---|---|---|---|---|---|---|
| 871 | ORF8 | A*23:01 | 9 | NYTVSCLPF | -0.17355 | Consensus (ann/smm) | 0.23 | 51.46% (247/480) | Non-Toxin |
| 872 | ORF8 | A*24:02 | 9 | NYTVSCLPF | -0.17355 | Consensus (ann/smm) | 0.34 | 51.46% (247/480) | Non-Toxin |
| 873 | ORF8 | A*33:01 | 11 | PIHFYSKWYIR | 0.03675 | ann | 0.94 | 99.38% (477/480) | Non-Toxin |
| 874 | ORF8 | A*01:01 | 9 | QSCTQHQPY | -0.16503 | Consensus (ann/smm) | 0.28 | 96.04% (461/480) | Non-Toxin |
| 875 | ORF8 | A*23:01 | 10 | QYIDIGNYTV | 0.24159 | Consensus (ann/smm) | 0.58 | 99.79% (479/480) | Non-Toxin |
| 876 | ORF8 | A*01:01 | 11 | RCSFYEDFLEY | 0.33894 | Consensus (ann/smm) | 0.56 | 99.79% (479/480) | Non-Toxin |
| 877 | ORF8 | B*08:01 | 10 | RVGARKSAPL | -0.23842 | Consensus (ann/smm) | 0.44 | 99.17% (476/480) | Non-Toxin |
| 878 | ORF8 | A*30:02 | 9 | SLVVRCSFY | -0.01663 | Consensus (ann/smm) | 0.57 | 100.00% (480/480) | Non-Toxin |
| 879 | ORF8 | A*68:02 | 9 | TVSCLPFTI | -0.00771 | Consensus (ann/comblib_sidney2008/smm) | 0.9 | 51.46% (247/480) | Non-Toxin |
| 880 | ORF8 | A*02:01 | 9 | YIDIGNYTV | 0.18759 | Consensus (ann/comblib_sidney2008/smm) | 0.5 | 99.79% (479/480) | Non-Toxin |
| 881 | ORF8 | A*02:06 | 9 | YIDIGNYTV | 0.18759 | Consensus (ann/smm) | 0.73 | 99.79% (479/480) | Non-Toxin |
| 882 | ORF8 | A*02:06 | 10 | YTVSCLPFTI | -0.10533 | Consensus (ann/smm) | 0.53 | 51.46% (247/480) | Non-Toxin |
| 883 | ORF8 | A*02:06 | 9 | YVVDDPCPI | -0.0051 | Consensus (ann/smm) | 0.68 | 99.17% (476/480) | Non-Toxin |
| S.No. | ORF | Allele | Length | Peptide | Immunogenicity | Method used | Percentile Rank | Conservancy | Toxicity |
| 884 | S Protein | B*40:01 | 9 | AEIRASANL | 0.00689 | Consensus (ann/smm) | 0.12 | 93.23% (468/502) | Non-Toxin |
| 885 | S Protein | B*44:03 | 9 | AEIRASANL | 0.00689 | Consensus (ann/smm) | 0.2 | 93.23% (468/502) | Non-Toxin |
| 886 | S Protein | B*44:02 | 9 | AEIRASANL | 0.00689 | Consensus (ann/smm) | 0.23 | 93.23% (468/502) | Non-Toxin |
| 887 | S Protein | B*44:02 | 9 | AENSVAYSN | -0.19132 | Consensus (ann/smm) | 0.21 | 93.82% (471/502) | Non-Toxin |
| 888 | S Protein | B*44:03 | 9 | AENSVAYSN | -0.19132 | Consensus (ann/smm) | 0.27 | 93.82% (471/502) | Non-Toxin |
| 889 | S Protein | B*44:02 | 9 | AEVQIDRLI | 0.08452 | Consensus (ann/smm) | 0.18 | 92.63% (465/502) | Non-Toxin |
| 890 | S Protein | B*44:03 | 9 | AEVQIDRLI | 0.08452 | Consensus (ann/smm) | 0.2 | 92.63% (465/502) | Non-Toxin |
| 891 | S Protein | A*26:01 | 9 | CVADYSVLY | -0.09595 | Consensus (ann/smm) | 0.17 | 93.82% (471/502) | Non-Toxin |
| 892 | S Protein | A*26:01 | 10 | EFVFKNIDGY | 0.0787 | Consensus (ann/smm) | 0.22 | 92.03% (462/502) | Non-Toxin |
| 893 | S Protein | B*51:01 | 8 | EPLVDLPI | 0.03974 | Consensus (ann/smm) | 0.18 | 93.63% (470/502) | Non-Toxin |
| 894 | S Protein | A*26:01 | 9 | ETKCTLKSF | -0.37555 | Consensus (ann/smm) | 0.17 | 93.43% (469/502) | Non-Toxin |
| 895 | S Protein | A*26:01 | 10 | EVFAQVKQIY | -0.21823 | Consensus (ann/smm) | 0.13 | 92.43% (464/502) | Non-Toxin |
| 896 | S Protein | B*35:01 | 9 | FAMQMAYRF | -0.28061 | Consensus (ann/comblib_sidney2008/smm) | 0.2 | 93.82% (471/502) | Non-Toxin |
| 897 | S Protein | B*40:01 | 9 | FEYVSQPFL | -0.17076 | Consensus (ann/smm) | 0.28 | 92.83% (466/502) | Non-Toxin |
| 898 | S Protein | A*02:03 | 9 | FIAGLIAIV | 0.27206 | Consensus (ann/smm) | 0.16 | 93.82% (471/502) | Non-Toxin |
| 899 | S Protein | B*53:01 | 10 | FLPFFSNVTW | 0.11853 | Consensus (ann/smm) | 0.21 | 93.43% (469/502) | Non-Toxin |
| 900 | S Protein | B*08:01 | 9 | FNATRFASV | 0.14872 | Consensus (ann/comblib_sidney2008/smm) | 0.2 | 93.43% (469/502) | Non-Toxin |
| 901 | S Protein | B*53:01 | 10 | FPNITNLCPF | 0.1009 | Consensus (ann/smm) | 0.06 | 93.82% (471/502) | Non-Toxin |
| 902 | S Protein | B*35:01 | 10 | FPNITNLCPF | 0.1009 | Consensus (ann/smm) | 0.08 | 93.82% (471/502) | Non-Toxin |
| 903 | S Protein | B*51:01 | 10 | FPNITNLCPF | 0.1009 | Consensus (ann/smm) | 0.17 | 93.82% (471/502) | Non-Toxin |
| 904 | S Protein | B*51:01 | 10 | FPQSAPHGVV | -0.0936 | Consensus (ann/smm) | 0.27 | 93.43% (469/502) | Non-Toxin |
| 905 | S Protein | A*68:02 | 9 | FTISVTTEI | 0.04473 | Consensus (ann/comblib_sidney2008/smm) | 0.2 | 93.82% (471/502) | Non-Toxin |
| 906 | S Protein | A*26:01 | 9 | FVFKNIDGY | -0.0215 | Consensus (ann/smm) | 0.11 | 92.03% (462/502) | Non-Toxin |
| 907 | S Protein | A*02:06 | 10 | FVFLVLLPLV | 0.02996 | Consensus (ann/smm) | 0.1 | 92.83% (466/502) | Non-Toxin |
| 908 | S Protein | A*02:01 | 10 | FVFLVLLPLV | 0.02996 | Consensus (ann/smm) | 0.28 | 92.83% (466/502) | Non-Toxin |
| 909 | S Protein | A*68:01 | 9 | FVIRGDEVR | 0.25778 | Consensus (ann/smm) | 0.17 | 93.63% (470/502) | Non-Toxin |
| 910 | S Protein | A*02:06 | 10 | FVSNGTHWFV | 0.29638 | Consensus (ann/smm) | 0.18 | 93.43% (469/502) | Non-Toxin |
| 911 | S Protein | A*02:01 | 10 | FVSNGTHWFV | 0.29638 | Consensus (ann/smm) | 0.21 | 93.43% (469/502) | Non-Toxin |
| 912 | S Protein | A*68:02 | 10 | FVSNGTHWFV | 0.29638 | Consensus (ann/smm) | 0.27 | 93.43% (469/502) | Non-Toxin |
| 913 | S Protein | B*44:03 | 9 | GEVFNATRF | 0.22473 | Consensus (ann/smm) | 0.2 | 93.63% (470/502) | Non-Toxin |
| 914 | S Protein | A*33:01 | 11 | GNYNYLYRLFR | 0.08205 | ann | 0.11 | 92.63% (465/502) | Non-Toxin |
| 915 | S Protein | A*31:01 | 9 | GTHWFVTQR | 0.35133 | Consensus (ann/smm) | 0.24 | 93.23% (468/502) | Non-Toxin |
| 916 | S Protein | A*03:01 | 9 | GVYFASTEK | 0.09023 | Consensus (ann/smm) | 0.2 | 93.63% (470/502) | Non-Toxin |
| 917 | S Protein | A*11:01 | 9 | GVYFASTEK | 0.09023 | Consensus (ann/smm) | 0.23 | 93.63% (470/502) | Non-Toxin |
| 918 | S Protein | A*03:01 | 9 | GVYYHKNNK | -0.18566 | Consensus (ann/smm) | 0.2 | 92.03% (462/502) | Non-Toxin |
| 919 | S Protein | A*68:01 | 10 | GVYYPDKVFR | -0.09388 | Consensus (ann/smm) | 0.27 | 93.82% (471/502) | Non-Toxin |
| 920 | S Protein | B*51:01 | 9 | IAIPTNFTI | 0.18523 | Consensus (ann/comblib_sidney2008/smm) | 0.2 | 93.82% (471/502) | Non-Toxin |

| | | | | | | | | | |
|---|---|---|---|---|---|---|---|---|---|
| 921 | S Protein | A*30:02 | 9 | IGAGICASY | 0.06201 | Consensus (ann/smm) | 0.21 | 93.82% (471/502) | Non-Toxin |
| 922 | S Protein | B*35:01 | 9 | IPFAMQMAY | -0.32801 | Consensus (ann/comblib_sidney2008/smm) | 0.1 | 93.82% (471/502) | Non-Toxin |
| 923 | S Protein | A*30:01 | 9 | ITRFQTLLA | 0.0425 | Consensus (ann/comblib_sidney2008/smm) | 0.2 | 93.43% (469/502) | Non-Toxin |
| 924 | S Protein | A*31:01 | 10 | KGIYQTSNFR | -0.12831 | Consensus (ann/smm) | 0.28 | 93.43% (469/502) | Non-Toxin |
| 925 | S Protein | A*32:01 | 9 | KIYSKHTPI | -0.32094 | Consensus (ann/comblib_sidney2008/smm) | 0.2 | 93.03% (467/502) | Non-Toxin |
| 926 | S Protein | A*31:01 | 9 | KQGNFKNLR | -0.09645 | Consensus (ann/smm) | 0.19 | 92.63% (465/502) | Non-Toxin |
| 927 | S Protein | A*30:02 | 10 | KSFTVEKGIY | 0.11812 | Consensus (ann/smm) | 0.11 | 93.23% (468/502) | Non-Toxin |
| 928 | S Protein | B*58:01 | 8 | KSNIIRGW | 0.33874 | ann | 0.25 | 93.82% (471/502) | Non-Toxin |
| 929 | S Protein | B*57:01 | 8 | KSNIIRGW | 0.33874 | ann | 0.29 | 93.82% (471/502) | Non-Toxin |
| 930 | S Protein | A*31:01 | 9 | KSNLKPFER | -0.0764 | Consensus (ann/smm) | 0.14 | 92.63% (465/502) | Non-Toxin |
| 931 | S Protein | A*31:01 | 9 | KSWMESEFR | -0.01013 | Consensus (ann/smm) | 0.26 | 92.43% (464/502) | Non-Toxin |
| 932 | S Protein | A*30:02 | 9 | KTSVDCTMY | -0.11115 | Consensus (ann/smm) | 0.12 | 93.82% (471/502) | Non-Toxin |
| 933 | S Protein | A*30:02 | 10 | KVGGNYNYLY | 0.01951 | Consensus (ann/smm) | 0.21 | 92.63% (465/502) | Non-Toxin |
| 934 | S Protein | A*24:02 | 10 | KWPWYIWLGF | 0.56424 | Consensus (ann/smm) | 0.11 | 93.43% (469/502) | Non-Toxin |
| 935 | S Protein | A*23:01 | 10 | KWPWYIWLGF | 0.56424 | Consensus (ann/smm) | 0.12 | 93.43% (469/502) | Non-Toxin |
| 936 | S Protein | A*32:01 | 10 | KWPWYIWLGF | 0.56424 | Consensus (ann/smm) | 0.18 | 93.43% (469/502) | Non-Toxin |
| 937 | S Protein | A*02:03 | 9 | LLFNKVTLA | -0.11337 | Consensus (ann/smm) | 0.23 | 93.23% (468/502) | Non-Toxin |
| 938 | S Protein | A*01:01 | 10 | LLTDEMIAQY | 0.05204 | Consensus (ann/smm) | 0.28 | 93.43% (469/502) | Non-Toxin |
| 939 | S Protein | B*51:01 | 10 | LPDDFTGCVI | 0.19184 | Consensus (ann/smm) | 0.14 | 93.63% (470/502) | Non-Toxin |
| 940 | S Protein | B*53:01 | 10 | LPDDFTGCVI | 0.19184 | Consensus (ann/smm) | 0.28 | 93.63% (470/502) | Non-Toxin |
| 941 | S Protein | B*53:01 | 9 | LPFFSNVTW | 0.04613 | Consensus (ann/comblib_sidney2008/smm) | 0.2 | 93.43% (469/502) | Non-Toxin |
| 942 | S Protein | B*53:01 | 10 | LPFFSNVTWF | 0.18944 | Consensus (ann/smm) | 0.13 | 91.83% (461/502) | Non-Toxin |
| 943 | S Protein | B*51:01 | 10 | LPFFSNVTWF | 0.18944 | Consensus (ann/smm) | 0.2 | 91.83% (461/502) | Non-Toxin |
| 944 | S Protein | B*53:01 | 10 | LPIGINITRF | 0.38888 | Consensus (ann/smm) | 0.12 | 93.43% (469/502) | Non-Toxin |
| 945 | S Protein | B*35:01 | 10 | LPIGINITRF | 0.38888 | Consensus (ann/smm) | 0.17 | 93.43% (469/502) | Non-Toxin |
| 946 | S Protein | B*51:01 | 9 | LPLVSSQCV | -0.40815 | Consensus (ann/comblib_sidney2008/smm) | 0.2 | 93.82% (471/502) | Non-Toxin |
| 947 | S Protein | B*51:01 | 10 | LPVSMTKTSV | -0.55317 | Consensus (ann/smm) | 0.2 | 93.82% (471/502) | Non-Toxin |
| 948 | S Protein | B*15:01 | 11 | LQIPFAMQMAY | -0.22124 | ann | 0.28 | 93.82% (471/502) | Non-Toxin |
| 949 | S Protein | A*01:01 | 9 | LTDEMIAQY | 0.02757 | Consensus (ann/smm) | 0.11 | 93.43% (469/502) | Non-Toxin |
| 950 | S Protein | A*24:02 | 10 | LYNSASFSTF | -0.29831 | Consensus (ann/smm) | 0.11 | 93.82% (471/502) | Non-Toxin |
| 951 | S Protein | A*23:01 | 10 | LYNSASFSTF | -0.29831 | Consensus (ann/smm) | 0.14 | 93.82% (471/502) | Non-Toxin |
| 952 | S Protein | A*02:06 | 10 | MQMAYRFNGI | 0.15371 | Consensus (ann/smm) | 0.18 | 93.82% (471/502) | Non-Toxin |
| 953 | S Protein | A*68:01 | 9 | NSASFSTFK | -0.09434 | Consensus (ann/smm) | 0.16 | 93.82% (471/502) | Non-Toxin |
| 954 | S Protein | A*68:02 | 9 | NTQEVFAQV | 0.17889 | Consensus (ann/comblib_sidney2008/smm) | 0.2 | 92.83% (466/502) | Non-Toxin |
| 955 | S Protein | A*01:01 | 10 | NTSNQVAVLY | -0.06762 | Consensus (ann/smm) | 0.27 | 93.82% (471/502) | Non-Toxin |
| 956 | S Protein | A*68:01 | 10 | NVYADSFVIR | 0.12147 | Consensus (ann/smm) | 0.14 | 93.82% (471/502) | Non-Toxin |
| 957 | S Protein | A*33:01 | 8 | NYLYRLFR | 0.13144 | ann | 0.07 | 92.63% (465/502) | Non-Toxin |
| 958 | S Protein | A*24:02 | 9 | NYNYLYRLF | 0.0171 | Consensus (ann/smm) | 0.17 | 92.63% (465/502) | Non-Toxin |
| 959 | S Protein | A*33:01 | 10 | NYNYLYRLFR | 0.08754 | Consensus (ann/smm) | 0.07 | 92.63% (465/502) | Non-Toxin |
| 960 | S Protein | A*01:01 | 11 | PLLTDEMIAQY | 0.07418 | Consensus (ann/smm) | 0.17 | 93.43% (469/502) | Non-Toxin |
| 961 | S Protein | A*23:01 | 9 | PYRVVVLSF | 0.03138 | Consensus (ann/smm) | 0.2 | 92.63% (465/502) | Non-Toxin |
| 962 | S Protein | A*03:01 | 9 | QIYKTPPIK | -0.12244 | Consensus (ann/smm) | 0.27 | 92.23% (463/502) | Non-Toxin |
| 963 | S Protein | A*23:01 | 9 | QYIKWPWYI | 0.21624 | Consensus (ann/smm) | 0.11 | 93.43% (469/502) | Non-Toxin |
| 964 | S Protein | A*24:02 | 9 | QYIKWPWYI | 0.21624 | Consensus (ann/smm) | 0.11 | 93.43% (469/502) | Non-Toxin |
| 965 | S Protein | A*23:01 | 10 | QYIKWPWYIW | 0.31425 | Consensus (ann/smm) | 0.16 | 93.43% (469/502) | Non-Toxin |
| 966 | S Protein | A*24:02 | 10 | QYIKWPWYIW | 0.31425 | Consensus (ann/smm) | 0.21 | 93.43% (469/502) | Non-Toxin |
| 967 | S Protein | A*30:02 | 9 | RISNCVADY | -0.02787 | Consensus (ann/smm) | 0.12 | 93.82% (471/502) | Non-Toxin |
| 968 | S Protein | A*03:01 | 9 | RLFRKSNLK | -0.28759 | Consensus (ann/smm) | 0.1 | 92.63% (465/502) | Non-Toxin |
| 969 | S Protein | A*32:01 | 11 | RLFRKSNLKPF | -0.48624 | ann | 0.26 | 92.63% (465/502) | Non-Toxin |
| 970 | S Protein | B*57:01 | 9 | RSFIEDLLF | 0.27446 | Consensus (ann/smm) | 0.23 | 93.43% (469/502) | Non-Toxin |
| 971 | S Protein | B*15:01 | 10 | RVYSTGSNVF | -0.23394 | Consensus (ann/smm) | 0.21 | 93.82% (471/502) | Non-Toxin |

| | | | | | | | | | |
|---|---|---|---|---|---|---|---|---|---|
| 972 | S Protein | A*30:02 | 9 | SANNCTFEY | 0.13273 | Consensus (ann/smm) | 0.26 | 92.83% (466/502) | Non-Toxin |
| 973 | S Protein | B*44:03 | 10 | SETKCTLKSF | -0.5082 | Consensus (ann/smm) | 0.19 | 93.43% (469/502) | Non-Toxin |
| 974 | S Protein | A*30:02 | 11 | SKVGGNYNYLY | 0.05491 | Consensus (ann/smm) | 0.18 | 92.63% (465/502) | Non-Toxin |
| 975 | S Protein | B*07:02 | 8 | SPRRARSV | 0.01608 | Consensus (ann/smm) | 0.17 | 93.82% (471/502) | Non-Toxin |
| 976 | S Protein | B*07:02 | 9 | SPRRARSVA | 0.0402 | Consensus (ann/comblib_sidney2008/smm) | 0.1 | 93.82% (471/502) | Non-Toxin |
| 977 | S Protein | B*07:02 | 10 | SPRRARSVAS | 0.05935 | Consensus (ann/smm) | 0.14 | 93.82% (471/502) | Non-Toxin |
| 978 | S Protein | A*30:02 | 10 | SSANNCTFEY | 0.14123 | Consensus (ann/smm) | 0.23 | 92.83% (466/502) | Non-Toxin |
| 979 | S Protein | A*01:01 | 10 | SSANNCTFEY | 0.14123 | Consensus (ann/smm) | 0.25 | 92.83% (466/502) | Non-Toxin |
| 980 | S Protein | A*32:01 | 9 | STQDLFLPF | 0.06828 | Consensus (ann/comblib_sidney2008/smm) | 0.2 | 93.03% (467/502) | Non-Toxin |
| 981 | S Protein | A*26:01 | 10 | SVASQSIIAY | -0.16721 | Consensus (ann/smm) | 0.19 | 93.82% (471/502) | Non-Toxin |
| 982 | S Protein | B*44:03 | 10 | TECSNLLLQY | -0.28855 | Consensus (ann/smm) | 0.28 | 93.82% (471/502) | Non-Toxin |
| 983 | S Protein | A*11:01 | 10 | TEILPVSMTK | -0.21981 | Consensus (ann/smm) | 0.28 | 93.82% (471/502) | Non-Toxin |
| 984 | S Protein | B*44:02 | 10 | TEKSNIIRGW | 0.07559 | Consensus (ann/smm) | 0.24 | 93.63% (470/502) | Non-Toxin |
| 985 | S Protein | B*53:01 | 9 | TPGDSSSGW | -0.40333 | Consensus (ann/comblib_sidney2008/smm) | 0.2 | 93.43% (469/502) | Non-Toxin |
| 986 | S Protein | A*01:01 | 9 | TSNQVAVLY | -0.01327 | Consensus (ann/smm) | 0.28 | 93.82% (471/502) | Non-Toxin |
| 987 | S Protein | A*30:01 | 9 | TTRTQLPPA | -0.08322 | Consensus (ann/comblib_sidney2008/smm) | 0.2 | 93.43% (469/502) | Non-Toxin |
| 988 | S Protein | A*30:02 | 9 | VLPFNDGVY | 0.1815 | Consensus (ann/smm) | 0.28 | 93.82% (471/502) | Non-Toxin |
| 989 | S Protein | A*24:02 | 11 | VLYNSASFSTF | -0.29855 | Consensus (ann/smm) | 0.2 | 93.82% (471/502) | Non-Toxin |
| 990 | S Protein | A*11:01 | 10 | VTLADAGFIK | 0.30393 | Consensus (ann/smm) | 0.22 | 93.23% (468/502) | Non-Toxin |
| 991 | S Protein | A*03:01 | 9 | VTYVPAQEK | 0.02711 | Consensus (ann/smm) | 0.27 | 93.63% (470/502) | Non-Toxin |
| 992 | S Protein | A*23:01 | 10 | VYSSANNCTF | -0.21728 | Consensus (ann/smm) | 0.12 | 92.63% (465/502) | Non-Toxin |
| 993 | S Protein | A*24:02 | 10 | VYSSANNCTF | -0.21728 | Consensus (ann/smm) | 0.12 | 92.63% (465/502) | Non-Toxin |
| 994 | S Protein | A*24:02 | 9 | VYSTGSNVF | -0.11871 | Consensus (ann/smm) | 0.17 | 93.82% (471/502) | Non-Toxin |
| 995 | S Protein | A*31:01 | 9 | VYYPDKVFR | -0.09052 | Consensus (ann/smm) | 0.18 | 93.82% (471/502) | Non-Toxin |
| 996 | S Protein | B*53:01 | 9 | WPWYIWLGF | 0.41673 | Consensus (ann/comblib_sidney2008/smm) | 0.2 | 93.43% (469/502) | Non-Toxin |
| 997 | S Protein | B*51:01 | 10 | WPWYIWLGFI | 0.50004 | Consensus (ann/smm) | 0.14 | 93.43% (469/502) | Non-Toxin |
| 998 | S Protein | A*26:01 | 9 | WTAGAAAYY | 0.15259 | Consensus (ann/smm) | 0.11 | 93.43% (469/502) | Non-Toxin |
| 999 | S Protein | A*30:02 | 9 | WTAGAAAYY | 0.15259 | Consensus (ann/smm) | 0.12 | 93.43% (469/502) | Non-Toxin |
| 1000 | S Protein | A*01:01 | 9 | WTAGAAAYY | 0.15259 | Consensus (ann/smm) | 0.17 | 93.43% (469/502) | Non-Toxin |
| 1001 | S Protein | A*68:02 | 10 | WTAGAAAYYV | 0.15455 | Consensus (ann/smm) | 0.06 | 93.43% (469/502) | Non-Toxin |
| 1002 | S Protein | B*44:02 | 9 | YEQYIKWPW | 0.06574 | Consensus (ann/smm) | 0.11 | 93.63% (470/502) | Non-Toxin |
| 1003 | S Protein | B*44:03 | 9 | YEQYIKWPW | 0.06574 | Consensus (ann/smm) | 0.12 | 93.63% (470/502) | Non-Toxin |
| 1004 | S Protein | B*44:03 | 10 | YEQYIKWPWY | 0.20685 | Consensus (ann/smm) | 0.14 | 93.43% (469/502) | Non-Toxin |
| 1005 | S Protein | B*44:02 | 10 | YEQYIKWPWY | 0.20685 | Consensus (ann/smm) | 0.24 | 93.43% (469/502) | Non-Toxin |
| 1006 | S Protein | A*02:01 | 9 | YLQPRTFLL | 0.1305 | Consensus (ann/comblib_sidney2008/smm) | 0.3 | 93.43% (469/502) | Non-Toxin |
| 1007 | S Protein | A*03:01 | 10 | YLQPRTFLLK | 0.1338 | Consensus (ann/smm) | 0.18 | 93.43% (469/502) | Non-Toxin |
| 1008 | S Protein | A*33:01 | 9 | YNYLYRLFR | 0.0918 | Consensus (ann/smm) | 0.12 | 92.63% (465/502) | Non-Toxin |
| 1009 | S Protein | B*07:02 | 10 | YPDKVFRSSV | -0.24316 | Consensus (ann/smm) | 0.26 | 93.82% (471/502) | Non-Toxin |
| 1010 | S Protein | A*02:06 | 9 | YQDVNCTEV | 0.08295 | Consensus (ann/smm) | 0.19 | 67.33% (338/502) | Non-Toxin |
| 1011 | S Protein | A*01:01 | 10 | YTNSFTRGVY | 0.08467 | Consensus (ann/smm) | 0.17 | 93.43% (469/502) | Non-Toxin |
| 1012 | S Protein | A*30:02 | 10 | YTNSFTRGVY | 0.08467 | Consensus (ann/smm) | 0.18 | 93.43% (469/502) | Non-Toxin |
| 1013 | S Protein | A*24:02 | 11 | YYVGYLQPRTF | -0.02378 | Consensus (ann/smm) | 0.15 | 93.43% (469/502) | Non-Toxin |

**Supplementary table S3:** High Percentile Ranking HTL epitopes-HLA allele pairs screened from entire proteome of SARS-CoV-2 by the "MHC-II Binding Predictions" tool of IEDB. These epitopes were further utilized to identify the potentially immunogenic multiple epitope cluster based HTL Ag-Patches from the entire proteome of the SRAS-CoV-2. The epitopes shown in **BLUE** are the epitope which form Overlapping epitope clusters. The screened epitopes are in consensus with the previous studies [Srivastava et al. 2020a; Srivastava et al. 2020b; and Grifoni et al., 2020a].

| S.No. | ORF | Allele | Length | Method used | Peptide | Percentile Rank | Adjusted rank | Conservancy | Toxicity |
|---|---|---|---|---|---|---|---|---|---|
| 1 | E Protein | DPA1*03:01/DPB1*04:02 | 15 | Consensus (comb.lib./smm/nn) | FLAFVVFLLVTLAIL | 0.06 | 0.06 | 99.59% (480/482) | Non-Toxin |
| 2 | E Protein | DPA1*03:01/DPB1*04:02 | 15 | Consensus (comb.lib./smm/nn) | LAFVVFLLVTLAILT | 0.06 | 0.06 | 99.59% (480/482) | Non-Toxin |
| 3 | E Protein | DPA1*03:01/DPB1*04:02 | 15 | Consensus (comb.lib./smm/nn) | LFLAFVVFLLVTLAI | 0.03 | 0.03 | 99.59% (480/482) | Non-Toxin |
| 4 | E Protein | DPA1*01:03/DPB1*02:01 | 15 | Consensus (comb.lib./smm/nn) | LFLAFVVFLLVTLAI | 0.04 | 0.04 | 99.59% (480/482) | Non-Toxin |
| 5 | E Protein | DPA1*01/DPB1*04:01 | 15 | Consensus (comb.lib./smm) | LFLAFVVFLLVTLAI | 0.08 | 0.08 | 99.59% (480/482) | Non-Toxin |
| 6 | E Protein | DPA1*02:01/DPB1*01:01 | 15 | Consensus (comb.lib./smm/nn) | LFLAFVVFLLVTLAI | 0.1 | 0.1 | 99.59% (480/482) | Non-Toxin |
| 7 | E Protein | DPA1*03:01/DPB1*04:02 | 15 | Consensus (comb.lib./smm/nn) | LLFLAFVVFLLVTLA | 0.02 | 0.02 | 99.59% (480/482) | Non-Toxin |
| 8 | E Protein | DPA1*01:03/DPB1*02:01 | 15 | Consensus (comb.lib./smm/nn) | LLFLAFVVFLLVTLA | 0.03 | 0.03 | 99.59% (480/482) | Non-Toxin |
| 9 | E Protein | DPA1*01/DPB1*04:01 | 15 | Consensus (comb.lib./smm) | LLFLAFVVFLLVTLA | 0.06 | 0.06 | 99.59% (480/482) | Non-Toxin |
| 10 | E Protein | DPA1*02:01/DPB1*01:01 | 15 | Consensus (comb.lib./smm/nn) | LLFLAFVVFLLVTLA | 0.1 | 0.1 | 99.59% (480/482) | Non-Toxin |
| 11 | E Protein | DPA1*01:03/DPB1*02:01 | 15 | Consensus (comb.lib./smm/nn) | NSVLLFLAFVVFLLV | 0.04 | 0.04 | 99.59% (480/482) | Non-Toxin |
| 12 | E Protein | DPA1*03:01/DPB1*04:02 | 15 | Consensus (comb.lib./smm/nn) | NSVLLFLAFVVFLLV | 0.06 | 0.06 | 99.59% (480/482) | Non-Toxin |
| 13 | E Protein | DPA1*01/DPB1*04:01 | 15 | Consensus (comb.lib./smm) | NSVLLFLAFVVFLLV | 0.07 | 0.07 | 99.59% (480/482) | Non-Toxin |
| 14 | E Protein | DPA1*02:01/DPB1*01:01 | 15 | Consensus (comb.lib./smm/nn) | NSVLLFLAFVVFLLV | 0.1 | 0.1 | 99.59% (480/482) | Non-Toxin |
| 15 | E Protein | DPA1*01:03/DPB1*02:01 | 15 | Consensus (comb.lib./smm/nn) | SVLLFLAFVVFLLVT | 0.04 | 0.04 | 99.59% (480/482) | Non-Toxin |
| 16 | E Protein | DPA1*03:01/DPB1*04:02 | 15 | Consensus (comb.lib./smm/nn) | SVLLFLAFVVFLLVT | 0.06 | 0.06 | 99.59% (480/482) | Non-Toxin |
| 17 | E Protein | DPA1*01/DPB1*04:01 | 15 | Consensus (comb.lib./smm) | SVLLFLAFVVFLLVT | 0.07 | 0.07 | 99.59% (480/482) | Non-Toxin |
| 18 | E Protein | DPA1*02:01/DPB1*01:01 | 15 | Consensus (comb.lib./smm/nn) | SVLLFLAFVVFLLVT | 0.1 | 0.1 | 99.59% (480/482) | Non-Toxin |
| 19 | E Protein | DPA1*03:01/DPB1*04:02 | 15 | Consensus (comb.lib./smm/nn) | VLLFLAFVVFLLVTL | 0.02 | 0.02 | 99.59% (480/482) | Non-Toxin |
| 20 | E Protein | DPA1*01:03/DPB1*02:01 | 15 | Consensus (comb.lib./smm/nn) | VLLFLAFVVFLLVTL | 0.04 | 0.04 | 99.59% (480/482) | Non-Toxin |
| 21 | E Protein | DPA1*01/DPB1*04:01 | 15 | Consensus (comb.lib./smm) | VLLFLAFVVFLLVTL | 0.06 | 0.06 | 99.59% (480/482) | Non-Toxin |
| 22 | E Protein | DPA1*02:01/DPB1*01:01 | 15 | Consensus (comb.lib./smm/nn) | VLLFLAFVVFLLVTL | 0.1 | 0.1 | 99.59% (480/482) | Non-Toxin |
| 23 | E Protein | DPA1*02:01/DPB1*01:01 | 15 | Consensus (comb.lib./smm/nn) | VNSVLLFLAFVVFLL | 0.1 | 0.1 | 99.59% (480/482) | Non-Toxin |
| S.No. | ORF | Allele | Length | Method used | Peptide | Percentile Rank | Adjusted rank | Conservancy | Toxicity |
| 24 | M Protein | DPA1*01:03/DPB1*02:01 | 15 | Consensus (comb.lib./smm/nn) | GLMWLSYFIASFRLF | 0.05 | 0.05 | 97.48% (465/477) | Non-Toxin |
| 25 | M Protein | DQA1*01:01/DQB1*05:01 | 15 | Consensus (comb.lib./smm/nn) | IKLIFLWLLWPVTLA | 0.07 | 0.07 | 97.48% (465/477) | Non-Toxin |
| 26 | M Protein | DQA1*01:01/DQB1*05:01 | 15 | Consensus (comb.lib./smm/nn) | KLIFLWLLWPVTLAC | 0.11 | 0.11 | 97.48% (465/477) | Non-Toxin |
| 27 | M Protein | DPA1*01:03/DPB1*02:01 | 15 | Consensus (comb.lib./smm/nn) | LMWLSYFIASFRLFA | 0.05 | 0.05 | 97.48% (465/477) | Non-Toxin |
| 28 | M Protein | DRB1*09:01 | 15 | Consensus (comb.lib./smm/nn) | LSYYKLGASQRVAGD | 0.06 | 0.06 | 97.48% (465/477) | Non-Toxin |
| 29 | M Protein | DRB1*01:01 | 15 | Consensus (comb.lib./smm/nn) | LSYYKLGASQRVAGD | 0.67 | 0.67 | 97.48% (465/477) | Non-Toxin |
| 30 | M Protein | DPA1*01:03/DPB1*02:01 | 15 | Consensus (comb.lib./smm/nn) | MWLSYFIASFRLFAR | 0.08 | 0.08 | 97.48% (465/477) | Non-Toxin |
| 31 | M Protein | DRB1*09:01 | 15 | Consensus (comb.lib./smm/nn) | RTLSYYKLGASQRVA | 0.06 | 0.06 | 97.48% (465/477) | Non-Toxin |
| 32 | M Protein | DRB1*01:01 | 15 | Consensus (comb.lib./smm/nn) | RTLSYYKLGASQRVA | 0.67 | 0.67 | 97.48% (465/477) | Non-Toxin |
| 33 | M Protein | DRB1*09:01 | 15 | Consensus (comb.lib./smm/nn) | SRTLSYYKLGASQRV | 0.07 | 0.07 | 97.48% (465/477) | Non-Toxin |
| 34 | M Protein | DRB1*09:01 | 15 | Consensus (comb.lib./smm/nn) | SYYKLGASQRVAGDS | 0.48 | 0.48 | 97.48% (465/477) | Non-Toxin |
| 35 | M Protein | DRB1*09:01 | 15 | Consensus (comb.lib./smm/nn) | TLSYYKLGASQRVAG | 0.07 | 0.07 | 97.48% (465/477) | Non-Toxin |
| 36 | M Protein | DRB1*01:01 | 15 | Consensus (comb.lib./smm/nn) | TLSYYKLGASQRVAG | 0.67 | 0.67 | 97.48% (465/477) | Non-Toxin |
| 37 | M Protein | DPA1*01:03/DPB1*02:01 | 15 | Consensus (comb.lib./smm/nn) | VGLMWLSYFIASFRL | 0.07 | 0.07 | 97.48% (465/477) | Non-Toxin |
| S.No. | ORF | Allele | Length | Method used | Peptide | Percentile Rank | Adjusted rank | Conservancy | Toxicity |
| 38 | N Protein | DQA1*01:02/DQB1*06:02 | 15 | Consensus (comb.lib./smm/nn) | ANNAAIVLQLPQGTT | 0.2 | 0.2 | 97.59% (486/498) | Non-Toxin |
| 39 | N Protein | DRB1*09:01 | 15 | Consensus (comb.lib./smm/nn) | AQFAPSASAFFGMSR | 0.01 | 0.01 | 97.59% (486/498) | Non-Toxin |
| 40 | N Protein | DRB1*11:01 | 15 | Consensus (smm/nn/sturniolo) | DDQIGYYRRATRRIR | 0.42 | 0.42 | 97.59% (486/498) | Non-Toxin |
| 41 | N Protein | DRB1*11:01 | 15 | Consensus (smm/nn/sturniolo) | DQIGYYRRATRRIRG | 0.42 | 0.42 | 97.59% (486/498) | Non-Toxin |

| | | | | | | | | | | |
|---|---|---|---|---|---|---|---|---|---|---|
| 42 | N Protein | DRB5*01:01 | 15 | Consensus (smm/nn/sturniolo) | DQIGYYRRATRRIRG | 0.58 | 0.58 | 97.59% (486/498) | Non-Toxin | |
| 43 | N Protein | DQA1*01:02/DQB1*06:02 | 15 | Consensus (comb.lib./smm/nn) | GTRNPANNAAIVLQL | 0.05 | 0.05 | 97.59% (486/498) | Non-Toxin | |
| 44 | N Protein | DRB1*07:01 | 15 | Consensus (comb.lib./smm/nn) | GTWLTYTGAIKLDDK | 0.58 | 0.58 | 97.59% (486/498) | Non-Toxin | |
| 45 | N Protein | DRB1*11:01 | 15 | Consensus (smm/nn/sturniolo) | GYYRRATRRIRGGDG | 0.42 | 0.42 | 97.59% (486/498) | Non-Toxin | |
| 46 | N Protein | DRB1*09:01 | 15 | Consensus (comb.lib./smm/nn) | IAQFAPSASAFFGMS | 0.01 | 0.01 | 97.59% (486/498) | Non-Toxin | |
| 47 | N Protein | DRB1*11:01 | 15 | Consensus (smm/nn/sturniolo) | IGYYRRATRRIRGGD | 0.42 | 0.42 | 97.59% (486/498) | Non-Toxin | |
| 48 | N Protein | DQA1*01:02/DQB1*06:02 | 15 | Consensus (comb.lib./smm/nn) | NNAAIVLQLPQGTTL | 0.42 | 0.42 | 97.59% (486/498) | Non-Toxin | |
| 49 | N Protein | DQA1*01:02/DQB1*06:02 | 15 | Consensus (comb.lib./smm/nn) | NPANNAAIVLQLPQG | 0.03 | 0.03 | 97.59% (486/498) | Non-Toxin | |
| 50 | N Protein | DQA1*01:02/DQB1*06:02 | 15 | Consensus (comb.lib./smm/nn) | PANNAAIVLQLPQGT | 0.04 | 0.04 | 97.59% (486/498) | Non-Toxin | |
| 51 | N Protein | DRB1*09:01 | 15 | Consensus (comb.lib./smm/nn) | PQIAQFAPSASAFFG | 0.01 | 0.01 | 97.59% (486/498) | Non-Toxin | |
| 52 | N Protein | DRB1*07:01 | 15 | Consensus (comb.lib./smm/nn) | PSGTWLTYTGAIKLD | 0.58 | 0.58 | 97.59% (486/498) | Non-Toxin | |
| 53 | N Protein | DRB1*09:01 | 15 | Consensus (comb.lib./smm/nn) | QIAQFAPSASAFFGM | 0.01 | 0.01 | 97.59% (486/498) | Non-Toxin | |
| 54 | N Protein | DRB1*11:01 | 15 | Consensus (smm/nn/sturniolo) | QIGYYRRATRRIRGG | 0.39 | 0.39 | 97.59% (486/498) | Non-Toxin | |
| 55 | N Protein | DRB5*01:01 | 15 | Consensus (smm/nn/sturniolo) | QIGYYRRATRRIRGG | 0.58 | 0.58 | 97.59% (486/498) | Non-Toxin | |
| 56 | N Protein | DQA1*01:02/DQB1*06:02 | 15 | Consensus (comb.lib./smm/nn) | RNPANNAAIVLQLPQ | 0.04 | 0.04 | 97.59% (486/498) | Non-Toxin | |
| 57 | N Protein | DRB1*07:01 | 15 | Consensus (comb.lib./smm/nn) | SGTWLTYTGAIKLDD | 0.58 | 0.58 | 97.59% (486/498) | Non-Toxin | |
| 58 | N Protein | DRB1*07:01 | 15 | Consensus (comb.lib./smm/nn) | TPSGTWLTYTGAIKL | 0.58 | 0.58 | 97.59% (486/498) | Non-Toxin | |
| 59 | N Protein | DQA1*01:02/DQB1*06:02 | 15 | Consensus (comb.lib./smm/nn) | TRNPANNAAIVLQLP | 0.04 | 0.04 | 97.59% (486/498) | Non-Toxin | |
| 60 | N Protein | DRB1*07:01 | 15 | Consensus (comb.lib./smm/nn) | TWLTYTGAIKLDDKD | 0.58 | 0.58 | 97.59% (486/498) | Non-Toxin | |
| 61 | N Protein | DRB1*09:01 | 15 | Consensus (comb.lib./smm/nn) | WPQIAQFAPSASAFF | 0.01 | 0.01 | 97.59% (486/498) | Non-Toxin | |
| S.No. | ORF | Allele | Length | Method used | Peptide | Percentile Rank | Adjusted rank | Conservancy | Toxicity | |
| 62 | ORF10 | DPA1*01/DPB1*04:01 | 15 | Consensus (comb.lib./smm) | FAFPFTIYSLLLCRM | 0.56 | 0.56 | 99.79% (478/479) | Non-Toxin | |
| 63 | ORF10 | DPA1*02:01/DPB1*01:01 | 15 | Consensus (comb.lib./smm/nn) | FAFPFTIYSLLLCRM | 1.1 | 1.1 | 99.79% (478/479) | Non-Toxin | |
| 64 | ORF10 | DPA1*03:01/DPB1*04:02 | 15 | Consensus (comb.lib./smm/nn) | FAFPFTIYSLLLCRM | 1.1 | 1.1 | 99.79% (478/479) | Non-Toxin | |
| 65 | ORF10 | DPA1*01:03/DPB1*02:01 | 15 | Consensus (comb.lib./smm/nn) | INVFAFPFTIYSLLL | 0.29 | 0.29 | 99.37% (476/479) | Non-Toxin | |
| 66 | ORF10 | DPA1*01/DPB1*04:01 | 15 | Consensus (comb.lib./smm) | INVFAFPFTIYSLLL | 0.46 | 0.46 | 99.37% (476/479) | Non-Toxin | |
| 67 | ORF10 | DPA1*02:01/DPB1*01:01 | 15 | Consensus (comb.lib./smm/nn) | INVFAFPFTIYSLLL | 0.72 | 0.72 | 99.37% (476/479) | Non-Toxin | |
| 68 | ORF10 | DPA1*01:03/DPB1*02:01 | 15 | Consensus (comb.lib./smm/nn) | NVFAFPFTIYSLLLC | 0.4 | 0.4 | 99.58% (477/479) | Non-Toxin | |
| 69 | ORF10 | DPA1*01/DPB1*04:01 | 15 | Consensus (comb.lib./smm) | NVFAFPFTIYSLLLC | 0.56 | 0.56 | 99.58% (477/479) | Non-Toxin | |
| 70 | ORF10 | DPA1*02:01/DPB1*01:01 | 15 | Consensus (comb.lib./smm/nn) | NVFAFPFTIYSLLLC | 0.69 | 0.69 | 99.58% (477/479) | Non-Toxin | |
| 71 | ORF10 | DPA1*01/DPB1*04:01 | 15 | Consensus (comb.lib./smm) | VFAFPFTIYSLLLCR | 0.56 | 0.56 | 99.58% (477/479) | Non-Toxin | |
| 72 | ORF10 | DPA1*02:01/DPB1*01:01 | 15 | Consensus (comb.lib./smm/nn) | VFAFPFTIYSLLLCR | 0.66 | 0.66 | 99.58% (477/479) | Non-Toxin | |
| 73 | ORF10 | DPA1*01:03/DPB1*02:01 | 15 | Consensus (comb.lib./smm/nn) | VFAFPFTIYSLLLCR | 0.71 | 0.71 | 99.58% (477/479) | Non-Toxin | |
| 74 | ORF10 | DPA1*03:01/DPB1*04:02 | 15 | Consensus (comb.lib./smm/nn) | VFAFPFTIYSLLLCR | 1.1 | 1.1 | 99.58% (477/479) | Non-Toxin | |
| 75 | ORF10 | DPA1*01:03/DPB1*02:01 | 15 | Consensus (comb.lib./smm/nn) | YINVFAFPFTIYSLL | 0.29 | 0.29 | 99.37% (476/479) | Non-Toxin | |
| S.No. | ORF | Allele | Length | Method used | Peptide | Percentile Rank | Adjusted rank | Conservancy | Toxicity | NSP |
| 76 | ORF1ab | DPA1*01:03/DPB1*02:01 | 15 | Consensus (comb.lib./smm/nn) | AAIMQLFFSYFAVHF | 0.05 | 0.05 | 100%(453/453) | Non-Toxin | nsp3 |
| 77 | ORF1ab | DQA1*01:02/DQB1*06:02 | 15 | Consensus (comb.lib./smm/nn) | AFASEAARVVRSIFS | 0.08 | 0.08 | 99.55%(451/453) | Non-Toxin | nsp2 |
| 78 | ORF1ab | DRB1*04:01 | 15 | Consensus (smm/nn/sturniolo) | AIASEFSSLPSYAAF | 0.13 | 0.13 | 99.33%(450/453) | Non-Toxin | nsp8 |
| 79 | ORF1ab | DRB1*09:01 | 15 | Consensus (comb.lib./smm/nn) | AIILASFSASTSAFV | 0.01 | 0.01 | 100%(453/453) | Non-Toxin | nsp2 |
| 80 | ORF1ab | DPA1*01:03/DPB1*02:01 | 15 | Consensus (comb.lib./smm/nn) | AIMQLFFSYFAVHFI | 0.05 | 0.05 | 100%(453/453) | Non-Toxin | nsp3 |
| 81 | ORF1ab | DRB1*15:01 | 15 | Consensus (smm/nn/sturniolo) | AMPNMLRIMASLVLA | 0.01 | 0.01 | 100%(453/453) | Non-Toxin | nsp12 |
| 82 | ORF1ab | DRB3*02:02 | 15 | NetMHCIIpan | ANYIFWRNTNPIQLS | 0.05 | 0.05 | 98.89%(448/453) | Non-Toxin | nsp16 |
| 83 | ORF1ab | DRB1*04:01 | 15 | Consensus (smm/nn/sturniolo) | ASEFSSLPSYAAFAT | 0.12 | 0.12 | 99.33%(450/453) | Non-Toxin | nsp8 |
| 84 | ORF1ab | DQA1*05:01/DQB1*03:01 | 15 | Consensus (comb.lib./smm/nn) | ASIVAGGIVAIVVTC | 0.03 | 0.03 | 100%(453/453) | Non-Toxin | nsp4 |
| 85 | ORF1ab | DRB1*07:01 | 15 | Consensus (comb.lib./smm/nn) | AVGNICYTPSKLIEY | 0.09 | 0.09 | 99.55%(451/453) | Non-Toxin | nsp4 |
| 86 | ORF1ab | DRB3*02:02 | 15 | NetMHCIIpan | AWWTAFVTNVNASSS | 0.01 | 0.01 | 98.89%(448/453) | Non-Toxin | nsp16 |
| 87 | ORF1ab | DPA1*01:03/DPB1*02:01 | 15 | Consensus (comb.lib./smm/nn) | AYILFTRFFYVLGLA | 0.01 | 0.01 | 100%(453/453) | Non-Toxin | nsp3 |
| 88 | ORF1ab | DPA1*01/DPB1*04:01 | 15 | Consensus (comb.lib./smm) | AYILFTRFFYVLGLA | 0.08 | 0.08 | 100%(453/453) | Non-Toxin | nsp3 |
| 89 | ORF1ab | DRB1*07:01 | 15 | Consensus (comb.lib./smm/nn) | CTFTRSTNSRIKASM | 0.12 | 0.12 | 98.67%(447/453) | Non-Toxin | nsp3 |
| 90 | ORF1ab | DQA1*05:01/DQB1*03:01 | 15 | Consensus (comb.lib./smm/nn) | DISASIVAGGIVAIV | 0.03 | 0.03 | 100%(453/453) | Non-Toxin | nsp4 |

| | | | | | | | | | | |
|---|---|---|---|---|---|---|---|---|---|---|
| 91 | ORF1ab | DPA1*03:01/DPB1*04:02 | 15 | Consensus (comb.lib./smm/nn) | EETKFLTENLLLYID | 0.03 | 0.03 | 99.77%(452/453) | Non-Toxin | nsp3 |
| 92 | ORF1ab | DRB1*11:01 | 15 | Consensus (smm/nn/sturniolo) | EFYAYLRKHFSMMIL | 0.05 | 0.05 | 100%(453/453) | Non-Toxin | nsp12 |
| 93 | ORF1ab | DQA1*05:01/DQB1*02:01 | 15 | Consensus (comb.lib./smm/nn) | EIDFLELAMDEFIER | 0.04 | 0.04 | 99.11%(449/453) | Non-Toxin | nsp15 |
| 94 | ORF1ab | DRB1*01:01 | 15 | Consensus (comb.lib./smm/nn) | ESPFVMMSAPPAQYE | 0.01 | 0.01 | 100%(453/453) | Non-Toxin | nsp3 |
| 95 | ORF1ab | DPA1*03:01/DPB1*04:02 | 15 | Consensus (comb.lib./smm/nn) | ETKFLTENLLLYIDI | 0.03 | 0.03 | 99.77%(452/453) | Non-Toxin | nsp3 |
| 96 | ORF1ab | DRB1*13:02 | 15 | Consensus (smm/nn/sturniolo) | EVKILNNLGVDIAAN | 0.12 | 0.12 | 99.77%(452/453) | Non-Toxin | nsp15 |
| 97 | ORF1ab | DPA1*01:03/DPB1*02:01 | 15 | Consensus (comb.lib./smm/nn) | EWFLAYILFTRFFYV | 0.03 | 0.03 | 100%(453/453) | Non-Toxin | nsp3 |
| 98 | ORF1ab | DRB3*02:02 | 15 | NetMHCIIpan | FAWWTAFVTNVNASS | 0.01 | 0.01 | 98.89%(448/453) | Non-Toxin | nsp16 |
| 99 | ORF1ab | DQA1*01:01/DQB1*05:01 | 15 | Consensus (comb.lib./smm/nn) | FISNSWLMWLIINLV | 0.05 | 0.05 | 100%(453/453) | Non-Toxin | nsp3 |
| 100 | ORF1ab | DPA1*01:03/DPB1*02:01 | 15 | Consensus (comb.lib./smm/nn) | FLAYILFTRFFYVLG | 0.01 | 0.01 | 100%(453/453) | Non-Toxin | nsp3 |
| 101 | ORF1ab | DPA1*02:01/DPB1*01:01 | 15 | Consensus (comb.lib./smm/nn) | FLAYILFTRFFYVLG | 0.07 | 0.07 | 100%(453/453) | Non-Toxin | nsp3 |
| 102 | ORF1ab | DPA1*01/DPB1*04:01 | 15 | Consensus (comb.lib./smm) | FLAYILFTRFFYVLG | 0.09 | 0.09 | 100%(453/453) | Non-Toxin | nsp3 |
| 103 | ORF1ab | DPA1*01/DPB1*04:01 | 15 | Consensus (comb.lib./smm) | FLFVAAIFYLITPVH | 0.11 | 0.11 | 99.77%(452/453) | Non-Toxin | nsp4 |
| 104 | ORF1ab | DRB1*07:01 | 15 | Consensus (comb.lib./smm/nn) | FSAVGNICYTPSKLI | 0.06 | 0.06 | 99.55%(451/453) | Non-Toxin | nsp4 |
| 105 | ORF1ab | DRB1*07:01 | 15 | Consensus (comb.lib./smm/nn) | FTPLVPFWITIAYII | 0.03 | 0.03 | 100%(453/453) | Non-Toxin | nsp4 |
| 106 | ORF1ab | DRB1*11:01 | 15 | Consensus (smm/nn/sturniolo) | FVNEFYAYLRKHFSM | 0.11 | 0.11 | 100%(453/453) | Non-Toxin | nsp12 |
| 107 | ORF1ab | DRB3*02:02 | 15 | NetMHCIIpan | HANYIFWRNTNPIQL | 0.11 | 0.11 | 98.89%(448/453) | Non-Toxin | nsp16 |
| 108 | ORF1ab | DQA1*01:01/DQB1*05:01 | 15 | Consensus (comb.lib./smm/nn) | HFISNSWLMWLIINL | 0.05 | 0.05 | 100%(453/453) | Non-Toxin | nsp3 |
| 109 | ORF1ab | DRB1*04:01 | 15 | Consensus (smm/nn/sturniolo) | IASEFSSLPSYAAFA | 0.13 | 0.13 | 99.33%(450/453) | Non-Toxin | nsp8 |
| 110 | ORF1ab | DQA1*05:01/DQB1*02:01 | 15 | Consensus (comb.lib./smm/nn) | IDFLELAMDEFIERY | 0.05 | 0.05 | 99.33%(450/453) | Non-Toxin | nsp15 |
| 111 | ORF1ab | DRB1*09:01 | 15 | Consensus (comb.lib./smm/nn) | IILASFSASTSAFVE | 0.01 | 0.01 | 100%(453/453) | Non-Toxin | nsp2 |
| 112 | ORF1ab | DRB1*09:01 | 15 | Consensus (comb.lib./smm/nn) | ILASFSASTSAFVET | 0.02 | 0.02 | 100%(453/453) | Non-Toxin | nsp2 |
| 113 | ORF1ab | DPA1*01:03/DPB1*02:01 | 15 | Consensus (comb.lib./smm/nn) | ILFTRFFYVLGLAAI | 0.11 | 0.11 | 100%(453/453) | Non-Toxin | nsp3 |
| 114 | ORF1ab | DPA1*01:03/DPB1*02:01 | 15 | Consensus (comb.lib./smm/nn) | IMQLFFSYFAVHFIS | 0.05 | 0.05 | 100%(453/453) | Non-Toxin | nsp3 |
| 115 | ORF1ab | DQA1*05:01/DQB1*03:01 | 15 | Consensus (comb.lib./smm/nn) | ISASIVAGGIVAIVV | 0.03 | 0.03 | 100%(453/453) | Non-Toxin | nsp4 |
| 116 | ORF1ab | DQA1*01:01/DQB1*05:01 | 15 | Consensus (comb.lib./smm/nn) | ISNSWLMWLIINLVQ | 0.07 | 0.07 | 100%(453/453) | Non-Toxin | nsp3 |
| 117 | ORF1ab | DPA1*03:01/DPB1*04:02 | 15 | Consensus (comb.lib./smm/nn) | KLINIIIWFLLLSVC | 0.09 | 0.09 | 97.79%(443/453) | Non-Toxin | nsp3 |
| 118 | ORF1ab | DPA1*02:01/DPB1*05:01 | 15 | Consensus (comb.lib./smm/nn) | KQLIKVTLVFLFVAA | 0.13 | 0.13 | 99.77%(452/453) | Non-Toxin | nsp4 |
| 119 | ORF1ab | DPA1*02:01/DPB1*05:01 | 15 | Consensus (comb.lib./smm/nn) | KVTLVFLFVAAIFYL | 0.04 | 0.04 | 99.77%(452/453) | Non-Toxin | nsp4 |
| 120 | ORF1ab | DPA1*01:03/DPB1*02:01 | 15 | Consensus (comb.lib./smm/nn) | LAAIMQLFFSYFAVH | 0.07 | 0.07 | 100%(453/453) | Non-Toxin | nsp3 |
| 121 | ORF1ab | DPA1*01:03/DPB1*02:01 | 15 | Consensus (comb.lib./smm/nn) | LAYILFTRFFYVLGL | 0.01 | 0.01 | 100%(453/453) | Non-Toxin | nsp3 |
| 122 | ORF1ab | DPA1*01/DPB1*04:01 | 15 | Consensus (comb.lib./smm) | LAYILFTRFFYVLGL | 0.08 | 0.08 | 100%(453/453) | Non-Toxin | nsp3 |
| 123 | ORF1ab | DRB1*07:01 | 15 | Consensus (comb.lib./smm/nn) | LCTFTRSTNSRIKAS | 0.12 | 0.12 | 99.11%(449/453) | Non-Toxin | nsp3 |
| 124 | ORF1ab | DPA1*03:01/DPB1*04:02 | 15 | Consensus (comb.lib./smm/nn) | LEETKFLTENLLLYI | 0.03 | 0.03 | 99.77%(452/453) | Non-Toxin | nsp3 |
| 125 | ORF1ab | DPA1*03:01/DPB1*04:02 | 15 | Consensus (comb.lib./smm/nn) | LINIIIWFLLLSVCL | 0.09 | 0.09 | 97.79%(443/453) | Non-Toxin | nsp3 |
| 126 | ORF1ab | DPA1*02:01/DPB1*05:01 | 15 | Consensus (comb.lib./smm/nn) | LKQLIKVTLVFLFVA | 0.13 | 0.13 | 99.77%(452/453) | Non-Toxin | nsp4 |
| 127 | ORF1ab | DRB1*07:01 | 15 | Consensus (comb.lib./smm/nn) | LLQLCTFTRSTNSRI | 0.12 | 0.12 | 98.89%(448/453) | Non-Toxin | nsp3 |
| 128 | ORF1ab | DRB1*07:01 | 15 | Consensus (comb.lib./smm/nn) | LQLCTFTRSTNSRIK | 0.12 | 0.12 | 98.89%(448/453) | Non-Toxin | nsp3 |
| 129 | ORF1ab | DPA1*01/DPB1*04:01 | 15 | Consensus (comb.lib./smm) | LVFLFVAAIFYLITP | 0.11 | 0.11 | 99.77%(452/453) | Non-Toxin | nsp4 |
| 130 | ORF1ab | DRB1*07:01 | 15 | Consensus (comb.lib./smm/nn) | LVPFWITIAYIICIS | 0.03 | 0.03 | 100%(453/453) | Non-Toxin | nsp4 |
| 131 | ORF1ab | DQA1*05:01/DQB1*02:01 | 15 | Consensus (comb.lib./smm/nn) | MEIDFLELAMDEFIE | 0.04 | 0.04 | 99.11%(449/453) | Non-Toxin | nsp15 |
| 132 | ORF1ab | DPA1*02:01/DPB1*14:01 | 15 | NetMHCIIpan | MNLKYAISAKNRART | 0.08 | 0.08 | 99.77%(452/453) | Non-Toxin | nsp12 |
| 133 | ORF1ab | DRB1*15:01 | 15 | Consensus (smm/nn/sturniolo) | MPNMLRIMASLVLAR | 0.01 | 0.01 | 100%(453/453) | Non-Toxin | nsp12 |
| 134 | ORF1ab | DPA1*01:03/DPB1*02:01 | 15 | Consensus (comb.lib./smm/nn) | MQLFFSYFAVHFISN | 0.05 | 0.05 | 100%(453/453) | Non-Toxin | nsp3 |
| 135 | ORF1ab | DPA1*01:03/DPB1*02:01 | 15 | Consensus (comb.lib./smm/nn) | MYIFFASFYYVWKSY | 0.04 | 0.04 | 99.11%(449/453) | Non-Toxin | nsp3 |
| 136 | ORF1ab | DRB1*11:01 | 15 | Consensus (smm/nn/sturniolo) | NEFYAYLRKHFSMMI | 0.02 | 0.02 | 100%(453/453) | Non-Toxin | nsp12 |
| 137 | ORF1ab | DRB1*15:01 | 15 | Consensus (smm/nn/sturniolo) | NMLRIMASLVLARKH | 0.01 | 0.01 | 100%(453/453) | Non-Toxin | nsp12 |
| 138 | ORF1ab | DQA1*01:01/DQB1*05:01 | 15 | Consensus (comb.lib./smm/nn) | NSWLMWLIINLVQMA | 0.1 | 0.1 | 100%(453/453) | Non-Toxin | nsp3 |
| 139 | ORF1ab | DRB3*02:02 | 15 | NetMHCIIpan | NYIFWRNTNPIQLSS | 0.04 | 0.04 | 98.89%(448/453) | Non-Toxin | nsp16 |
| 140 | ORF1ab | DRB1*13:02 | 15 | Consensus (smm/nn/sturniolo) | PEVKILNNLGVDIAA | 0.12 | 0.12 | 99.77%(452/453) | Non-Toxin | nsp15 |
| 141 | ORF1ab | DRB1*07:01 | 15 | Consensus (comb.lib./smm/nn) | PLVPFWITIAYIICI | 0.02 | 0.02 | 100%(453/453) | Non-Toxin | nsp4 |

| | | | | | | | | | | |
|---|---|---|---|---|---|---|---|---|---|---|
| 142 | ORF1ab | DRB1*15:01 | 15 | Consensus (smm/nn/sturniolo) | PNMLRIMASLVLARK | 0.01 | 0.01 | 100%(453/453) | Non-Toxin | nsp12 |
| 143 | ORF1ab | DRB1*04:01 | 15 | Consensus (smm/nn/sturniolo) | QAIASEFSSLPSYAA | 0.13 | 0.13 | 99.33%(450/453) | Non-Toxin | nsp8 |
| 144 | ORF1ab | DRB1*01:01 | 15 | Consensus (comb.lib./smm/nn) | QESPFVMMSAPPAQY | 0.01 | 0.01 | 100%(453/453) | Non-Toxin | nsp3 |
| 145 | ORF1ab | DRB1*07:01 | 15 | Consensus (comb.lib./smm/nn) | QLCTFTRSTNSRIKA | 0.12 | 0.12 | 98.67%(447/453) | Non-Toxin | nsp3 |
| 146 | ORF1ab | DQA1*05:01/DQB1*02:01 | 15 | Consensus (comb.lib./smm/nn) | QMEIDFLELAMDEFI | 0.03 | 0.03 | 99.11%(449/453) | Non-Toxin | nsp15 |
| 147 | ORF1ab | DPA1*02:01/DPB1*14:01 | 15 | NetMHCIIpan | QMNLKYAISAKNRAR | 0.07 | 0.07 | 99.77%(452/453) | Non-Toxin | nsp12 |
| 148 | ORF1ab | DRB1*01:01 | 15 | Consensus (comb.lib./smm/nn) | QQESPFVMMSAPPAQ | 0.1 | 0.1 | 99.77%(452/453) | Non-Toxin | nsp3 |
| 149 | ORF1ab | DQA1*01:01/DQB1*05:01 | 15 | Consensus (comb.lib./smm/nn) | QSTQWSLFFFLYENA | 0.1 | 0.1 | 92.27%(418/453) | Non-Toxin | nsp6 |
| 150 | ORF1ab | DPA1*01:03/DPB1*02:01 | 15 | Consensus (comb.lib./smm/nn) | QWSLFFFLYENAFLP | 0.04 | 0.04 | 92.27%(418/453) | Non-Toxin | nsp6 |
| 151 | ORF1ab | DRB1*15:01 | 15 | Consensus (smm/nn/sturniolo) | RAMPNMLRIMASLVL | 0.01 | 0.01 | 100%(453/453) | Non-Toxin | nsp12 |
| 152 | ORF1ab | DPA1*01:03/DPB1*02:01 | 15 | Consensus (comb.lib./smm/nn) | RMYIFFASFYYVWKS | 0.04 | 0.04 | 99.11%(449/453) | Non-Toxin | nsp3 |
| 153 | ORF1ab | DQA1*05:01/DQB1*03:01 | 15 | Consensus (comb.lib./smm/nn) | SASIVAGGIVAIVVT | 0.03 | 0.03 | 100%(453/453) | Non-Toxin | nsp4 |
| 154 | ORF1ab | DRB1*07:01 | 15 | Consensus (comb.lib./smm/nn) | SAVGNICYTPSKLIE | 0.09 | 0.09 | 99.55%(451/453) | Non-Toxin | nsp4 |
| 155 | ORF1ab | DQA1*05:01/DQB1*03:01 | 15 | Consensus (comb.lib./smm/nn) | SIVAGGIVAIVVTCL | 0.03 | 0.03 | 100%(453/453) | Non-Toxin | nsp4 |
| 156 | ORF1ab | DPA1*03:01/DPB1*04:02 | 15 | Consensus (comb.lib./smm/nn) | SKLINIIIWFLLLSV | 0.09 | 0.09 | 97.79%(443/453) | Non-Toxin | nsp3 |
| 157 | ORF1ab | DQA1*01:01/DQB1*05:01 | 15 | Consensus (comb.lib./smm/nn) | SNSWLMWLIINLVQM | 0.07 | 0.07 | 100%(453/453) | Non-Toxin | nsp3 |
| 158 | ORF1ab | DRB1*01:01 | 15 | Consensus (comb.lib./smm/nn) | SPFVMMSAPPAQYEL | 0.01 | 0.01 | 100%(453/453) | Non-Toxin | nsp3 |
| 159 | ORF1ab | DQA1*05:01/DQB1*02:01 | 15 | Consensus (comb.lib./smm/nn) | SQMEIDFLELAMDEF | 0.05 | 0.05 | 99.11%(449/453) | Non-Toxin | nsp15 |
| 160 | ORF1ab | DPA1*01:03/DPB1*02:01 | 15 | Consensus (comb.lib./smm/nn) | STQWSLFFFLYENAF | 0.05 | 0.05 | 92.27%(418/453) | Non-Toxin | nsp6 |
| 161 | ORF1ab | DQA1*01:01/DQB1*05:01 | 15 | Consensus (comb.lib./smm/nn) | STQWSLFFFLYENAF | 0.1 | 0.1 | 92.27%(418/453) | Non-Toxin | nsp6 |
| 162 | ORF1ab | DPA1*03:01/DPB1*04:02 | 15 | Consensus (comb.lib./smm/nn) | TKFLTENLLLYIDIN | 0.05 | 0.05 | 99.77%(452/453) | Non-Toxin | nsp3 |
| 163 | ORF1ab | DPA1*03:01/DPB1*04:02 | 15 | Consensus (comb.lib./smm/nn) | TLEETKFLTENLLLY | 0.05 | 0.05 | 99.77%(452/453) | Non-Toxin | nsp3 |
| 164 | ORF1ab | DPA1*01/DPB1*04:01 | 15 | Consensus (comb.lib./smm) | TLVFLFVAAIFYLIT | 0.11 | 0.11 | 99.77%(452/453) | Non-Toxin | nsp4 |
| 165 | ORF1ab | DRB1*07:01 | 15 | Consensus (comb.lib./smm/nn) | TPLVPFWITIAYIIC | 0.03 | 0.03 | 100%(453/453) | Non-Toxin | nsp4 |
| 166 | ORF1ab | DPA1*02:01/DPB1*14:01 | 15 | NetMHCIIpan | TQMNLKYAISAKNRA | 0.11 | 0.11 | 99.77%(452/453) | Non-Toxin | nsp12 |
| 167 | ORF1ab | DPA1*01:03/DPB1*02:01 | 15 | Consensus (comb.lib./smm/nn) | TQWSLFFFLYENAFL | 0.04 | 0.04 | 92.27%(418/453) | Non-Toxin | nsp6 |
| 168 | ORF1ab | DQA1*01:01/DQB1*05:01 | 15 | Consensus (comb.lib./smm/nn) | TQWSLFFFLYENAFL | 0.09 | 0.09 | 92.27%(418/453) | Non-Toxin | nsp6 |
| 169 | ORF1ab | DPA1*01/DPB1*04:01 | 15 | Consensus (comb.lib./smm) | VFLFVAAIFYLITPV | 0.11 | 0.11 | 99.77%(452/453) | Non-Toxin | nsp4 |
| 170 | ORF1ab | DRB1*07:01 | 15 | Consensus (comb.lib./smm/nn) | VGNICYTPSKLIEYT | 0.13 | 0.13 | 99.77%(452/453) | Non-Toxin | nsp4 |
| 171 | ORF1ab | DRB1*11:01 | 15 | Consensus (smm/nn/sturniolo) | VNEFYAYLRKHFSMM | 0.05 | 0.05 | 100%(453/453) | Non-Toxin | nsp12 |
| 172 | ORF1ab | DRB1*07:01 | 15 | Consensus (comb.lib./smm/nn) | VPFWITIAYIICIST | 0.03 | 0.03 | 100%(453/453) | Non-Toxin | nsp4 |
| 173 | ORF1ab | DRB1*01:01 | 15 | Consensus (comb.lib./smm/nn) | VQQESPFVMMSAPPA | 0.1 | 0.1 | 99.77%(452/453) | Non-Toxin | nsp3 |
| 174 | ORF1ab | DQA1*01:01/DQB1*05:01 | 15 | Consensus (comb.lib./smm/nn) | VQSTQWSLFFFLYEN | 0.11 | 0.11 | 92.27%(418/453) | Non-Toxin | nsp6 |
| 175 | ORF1ab | DPA1*01:03/DPB1*02:01 | 15 | Consensus (comb.lib./smm/nn) | VRMYIFFASFYYVWK | 0.03 | 0.03 | 98.45%(446/453) | Non-Toxin | nsp3 |
| 176 | ORF1ab | DRB1*15:01 | 15 | Consensus (smm/nn/sturniolo) | VRMYIFFASFYYVWK | 0.1 | 0.1 | 98.45%(446/453) | Non-Toxin | nsp3 |
| 177 | ORF1ab | DPA1*01/DPB1*04:01 | 15 | Consensus (comb.lib./smm) | VTLVFLFVAAIFYLI | 0.11 | 0.11 | 99.77%(452/453) | Non-Toxin | nsp4 |
| 178 | ORF1ab | DPA1*01:03/DPB1*02:01 | 15 | Consensus (comb.lib./smm/nn) | WFLAYILFTRFFYVL | 0.01 | 0.01 | 100%(453/453) | Non-Toxin | nsp3 |
| 179 | ORF1ab | DPA1*02:01/DPB1*01:01 | 15 | Consensus (comb.lib./smm/nn) | WFLAYILFTRFFYVL | 0.06 | 0.06 | 100%(453/453) | Non-Toxin | nsp3 |
| 180 | ORF1ab | DPA1*01/DPB1*04:01 | 15 | Consensus (comb.lib./smm) | WFLAYILFTRFFYVL | 0.08 | 0.08 | 100%(453/453) | Non-Toxin | nsp3 |
| 181 | ORF1ab | DPA1*02:01/DPB1*05:01 | 15 | Consensus (comb.lib./smm/nn) | WLKQLIKVTLVFLFV | 0.13 | 0.13 | 99.77%(452/453) | Non-Toxin | nsp4 |
| 182 | ORF1ab | DQA1*01:02/DQB1*06:02 | 15 | Consensus (comb.lib./smm/nn) | YAFASEAARVVRSIF | 0.08 | 0.08 | 99.77%(452/453) | Non-Toxin | nsp2 |
| 183 | ORF1ab | DPA1*01:03/DPB1*02:01 | 15 | Consensus (comb.lib./smm/nn) | YIFFASFYYVWKSYV | 0.05 | 0.05 | 99.55%(451/453) | Non-Toxin | nsp3 |
| 184 | ORF1ab | DRB3*02:02 | 15 | NetMHCIIpan | YIFWRNTNPIQLSSY | 0.05 | 0.05 | 98.89%(448/453) | Non-Toxin | nsp16 |
| 185 | ORF1ab | DPA1*01:03/DPB1*02:01 | 15 | Consensus (comb.lib./smm/nn) | YILFTRFFYVLGLAA | 0.08 | 0.08 | 100%(453/453) | Non-Toxin | nsp3 |
| S.No. | ORF | Allele | Length | Method used | Peptide | Percentile Rank | Adjusted rank | Conservancy | Toxicity | |
| 186 | ORF3a | DPA1*01/DPB1*04:01 | 15 | Consensus (comb.lib./smm) | APFLYLYALVYFLQS | 0.12 | 0.12 | 96.88%(466/481) | Non-Toxin | |
| 187 | ORF3a | DPA1*02:01/DPB1*14:01 | 15 | NetMHCIIpan | DFVRATATIPIQASL | 0.12 | 0.12 | 99.37%(478/481) | Non-Toxin | |
| 188 | ORF3a | DPA1*01:03/DPB1*02:01 | 15 | Consensus (comb.lib./smm/nn) | DTGVEHVTFFIYNKI | 1.1 | 1.1 | 99.37%(478/481) | Non-Toxin | |
| 189 | ORF3a | DPA1*02:01/DPB1*05:01 | 15 | Consensus (comb.lib./smm/nn) | DTGVEHVTFFIYNKI | 1.6 | 1.6 | 99.37%(478/481) | Non-Toxin | |
| 190 | ORF3a | DRB1*04:05 | 15 | Consensus (smm/nn/sturniolo) | FFIYNKIVDEPEEHV | 0.94 | 0.94 | 99.58%(479/481) | Non-Toxin | |
| 191 | ORF3a | DPA1*01/DPB1*04:01 | 15 | Consensus (comb.lib./smm) | FLYLYALVYFLQSIN | 0.12 | 0.12 | 96.67%(465/481) | Non-Toxin | |

| | | | | | | | | | |
|---|---|---|---|---|---|---|---|---|---|
| 192 | ORF3a | DPA1*02:01/DPB1*14:01 | 15 | NetMHCIIpan | FVRATATIPIQASLP | 0.12 | 0.12 | 99.37%(478/481) | Non-Toxin |
| 193 | ORF3a | DPA1*01:03/DPB1*02:01 | 15 | Consensus (comb.lib./smm/nn) | GVEHVTFFIYNKIVD | 1.2 | 1.2 | 99.37%(478/481) | Non-Toxin |
| 194 | ORF3a | DRB1*04:05 | 15 | Consensus (smm/nn/sturniolo) | HVTFFIYNKIVDEPE | 0.84 | 0.84 | 99.58%(479/481) | Non-Toxin |
| 195 | ORF3a | DRB1*01:01 | 15 | Consensus (comb.lib./smm/nn) | LLFVTVYSHLLLVAA | 0.1 | 0.1 | 97.08%(467/481) | Non-Toxin |
| 196 | ORF3a | DPA1*01/DPB1*04:01 | 15 | Consensus (comb.lib./smm) | LYLYALVYFLQSINF | 0.12 | 0.12 | 96.46%(464/481) | Non-Toxin |
| 197 | ORF3a | DPA1*01/DPB1*04:01 | 15 | Consensus (comb.lib./smm) | PFLYLYALVYFLQSI | 0.12 | 0.12 | 96.67%(465/481) | Non-Toxin |
| 198 | ORF3a | DPA1*02:01/DPB1*14:01 | 15 | NetMHCIIpan | SDFVRATATIPIQAS | 0.12 | 0.12 | 99.37%(478/481) | Non-Toxin |
| 199 | ORF3a | DRB1*04:05 | 15 | Consensus (smm/nn/sturniolo) | TFFIYNKIVDEPEEH | 0.81 | 0.81 | 99.58%(479/481) | Non-Toxin |
| 200 | ORF3a | DPA1*01:03/DPB1*02:01 | 15 | Consensus (comb.lib./smm/nn) | TGVEHVTFFIYNKIV | 0.93 | 0.93 | 99.37%(478/481) | Non-Toxin |
| 201 | ORF3a | DPA1*02:01/DPB1*05:01 | 15 | Consensus (comb.lib./smm/nn) | TGVEHVTFFIYNKIV | 1.4 | 1.4 | 99.37%(478/481) | Non-Toxin |
| 202 | ORF3a | DRB1*04:05 | 15 | Consensus (smm/nn/sturniolo) | VTFFIYNKIVDEPEE | 0.81 | 0.81 | 99.58%(479/481) | Non-Toxin |
| 203 | ORF3a | DPA1*01/DPB1*04:01 | 15 | Consensus (comb.lib./smm) | YLYALVYFLQSINFV | 0.14 | 0.14 | 96.46%(464/481) | Non-Toxin |
| S.No. | ORF | Allele | Length | Method used | Peptide | Percentile Rank | Adjusted rank | Conservancy | Toxicity |
| 204 | ORF6 | DRB1*15:01 | 15 | Consensus (smm/nn/sturniolo) | EILLIIMRTFKVSIW | 0.1 | 0.1 | 99.58% (479/481) | Non-Toxin |
| 205 | ORF6 | DQA1*01:01/DQB1*05:01 | 15 | Consensus (comb.lib./smm/nn) | FKVSIWNLDYIINLI | 0.02 | 0.02 | 99.38% (478/481) | Non-Toxin |
| 206 | ORF6 | DRB1*15:01 | 15 | Consensus (smm/nn/sturniolo) | ILLIIMRTFKVSIWN | 0.1 | 0.1 | 99.38% (478/481) | Non-Toxin |
| 207 | ORF6 | DQA1*01:01/DQB1*05:01 | 15 | Consensus (comb.lib./smm/nn) | KVSIWNLDYIINLII | 0.02 | 0.02 | 99.38% (478/481) | Non-Toxin |
| 208 | ORF6 | DQA1*01:01/DQB1*05:01 | 15 | Consensus (comb.lib./smm/nn) | TFKVSIWNLDYIINL | 0.02 | 0.02 | 99.38% (478/481) | Non-Toxin |
| 209 | ORF6 | DQA1*01:01/DQB1*05:01 | 15 | Consensus (comb.lib./smm/nn) | VSIWNLDYIINLIIK | 0.05 | 0.05 | 99.38% (478/481) | Non-Toxin |
| | ORF | Allele | Length | Method used | Peptide | Percentile Rank | Adjusted rank | Conservancy | Toxicity |
| 210 | ORF7a | DRB1*01:01 | 15 | Consensus (comb.lib./smm/nn) | IILFLALITLATCEL | 0.16 | 0.16 | 99.79% (479/480) | Non-Toxin |
| 211 | ORF7a | DRB1*01:01 | 15 | Consensus (comb.lib./smm/nn) | ILFLALITLATCELY | 0.16 | 0.16 | 99.79% (479/480) | Non-Toxin |
| 212 | ORF7a | DRB1*01:01 | 15 | Consensus (comb.lib./smm/nn) | KIILFLALITLATCE | 0.16 | 0.16 | 99.58% (478/480) | Non-Toxin |
| 213 | ORF7a | DRB1*01:01 | 15 | Consensus (comb.lib./smm/nn) | MKIILFLALITLATC | 0.16 | 0.16 | 99.58% (478/480) | Non-Toxin |
| 214 | ORF7a | DPA1*03:01/DPB1*04:02 | 15 | Consensus (comb.lib./smm/nn) | MKIILFLALITLATC | 0.39 | 0.39 | 99.58% (478/480) | Non-Toxin |
| S.No. | ORF | Allele | Length | Method used | Peptide | Percentile Rank | Adjusted rank | Conservancy | Toxicity |
| 215 | ORF7b | DRB4*01:01 | 15 | Consensus (comb.lib./smm/nn) | AFLLFLVLIMLIIFW | 0.14 | 0.14 | 99.58% (235/236) | Non-Toxin |
| 216 | ORF7b | DPA1*03:01/DPB1*04:02 | 15 | Consensus (comb.lib./smm/nn) | CFLAFLLFLVLIMLI | 0.03 | 0.03 | 97.88% (231/236) | Non-Toxin |
| 217 | ORF7b | DPA1*01:03/DPB1*02:01 | 15 | Consensus (comb.lib./smm/nn) | CFLAFLLFLVLIMLI | 0.08 | 0.08 | 97.88% (231/236) | Non-Toxin |
| 218 | ORF7b | DPA1*01/DPB1*04:01 | 15 | Consensus (comb.lib./smm) | CFLAFLLFLVLIMLI | 0.11 | 0.11 | 97.88% (231/236) | Non-Toxin |
| 219 | ORF7b | DPA1*03:01/DPB1*04:02 | 15 | Consensus (comb.lib./smm/nn) | DFYLCFLAFLLFLVL | 0.03 | 0.03 | 98.31% (232/236) | Non-Toxin |
| 220 | ORF7b | DPA1*01:03/DPB1*02:01 | 15 | Consensus (comb.lib./smm/nn) | DFYLCFLAFLLFLVL | 0.08 | 0.08 | 98.31% (232/236) | Non-Toxin |
| 221 | ORF7b | DPA1*01/DPB1*04:01 | 15 | Consensus (comb.lib./smm) | DFYLCFLAFLLFLVL | 0.09 | 0.09 | 98.31% (232/236) | Non-Toxin |
| 222 | ORF7b | DPA1*03:01/DPB1*04:02 | 15 | Consensus (comb.lib./smm/nn) | FLAFLLFLVLIMLII | 0.06 | 0.06 | 97.88% (231/236) | Non-Toxin |
| 223 | ORF7b | DRB4*01:01 | 15 | Consensus (comb.lib./smm/nn) | FLAFLLFLVLIMLII | 0.14 | 0.14 | 97.88% (231/236) | Non-Toxin |
| 224 | ORF7b | DRB4*01:01 | 15 | Consensus (comb.lib./smm/nn) | FLLFLVLIMLIIFWF | 0.19 | 0.19 | 99.58% (235/236) | Non-Toxin |
| 225 | ORF7b | DPA1*03:01/DPB1*04:02 | 15 | Consensus (comb.lib./smm/nn) | FYLCFLAFLLFLVLI | 0.03 | 0.03 | 97.88% (231/236) | Non-Toxin |
| 226 | ORF7b | DPA1*01:03/DPB1*02:01 | 15 | Consensus (comb.lib./smm/nn) | FYLCFLAFLLFLVLI | 0.06 | 0.06 | 97.88% (231/236) | Non-Toxin |
| 227 | ORF7b | DPA1*01/DPB1*04:01 | 15 | Consensus (comb.lib./smm) | FYLCFLAFLLFLVLI | 0.09 | 0.09 | 97.88% (231/236) | Non-Toxin |
| 228 | ORF7b | DPA1*01:03/DPB1*02:01 | 15 | Consensus (comb.lib./smm/nn) | IDFYLCFLAFLLFLV | 0.06 | 0.06 | 98.31% (232/236) | Non-Toxin |
| 229 | ORF7b | DPA1*03:01/DPB1*04:02 | 15 | Consensus (comb.lib./smm/nn) | IDFYLCFLAFLLFLV | 0.16 | 0.16 | 98.31% (232/236) | Non-Toxin |
| 230 | ORF7b | DPA1*03:01/DPB1*04:02 | 15 | Consensus (comb.lib./smm/nn) | LAFLLFLVLIMLIIF | 0.08 | 0.08 | 99.58% (235/236) | Non-Toxin |
| 231 | ORF7b | DRB4*01:01 | 15 | Consensus (comb.lib./smm/nn) | LAFLLFLVLIMLIIF | 0.12 | 0.12 | 99.58% (235/236) | Non-Toxin |
| 232 | ORF7b | DPA1*03:01/DPB1*04:02 | 15 | Consensus (comb.lib./smm/nn) | LCFLAFLLFLVLIML | 0.02 | 0.02 | 97.88% (231/236) | Non-Toxin |
| 233 | ORF7b | DPA1*01:03/DPB1*02:01 | 15 | Consensus (comb.lib./smm/nn) | LCFLAFLLFLVLIML | 0.06 | 0.06 | 97.88% (231/236) | Non-Toxin |
| 234 | ORF7b | DPA1*01/DPB1*04:01 | 15 | Consensus (comb.lib./smm) | LCFLAFLLFLVLIML | 0.09 | 0.09 | 97.88% (231/236) | Non-Toxin |
| 235 | ORF7b | DPA1*01:03/DPB1*02:01 | 15 | Consensus (comb.lib./smm/nn) | LIDFYLCFLAFLLFL | 0.05 | 0.05 | 98.31% (232/236) | Non-Toxin |
| 236 | ORF7b | DPA1*01:03/DPB1*02:01 | 15 | Consensus (comb.lib./smm/nn) | LSLIDFYLCFLAFLL | 0.15 | 0.15 | 98.31% (232/236) | Non-Toxin |
| 237 | ORF7b | DQA1*01:01/DQB1*05:01 | 15 | Consensus (comb.lib./smm/nn) | LVLIMLIIFWFSLEL | 0.14 | 0.14 | 99.58% (235/236) | Non-Toxin |
| 238 | ORF7b | DPA1*01:03/DPB1*02:01 | 15 | Consensus (comb.lib./smm/nn) | SLIDFYLCFLAFLLF | 0.07 | 0.07 | 98.31% (232/236) | Non-Toxin |
| 239 | ORF7b | DPA1*03:01/DPB1*04:02 | 15 | Consensus (comb.lib./smm/nn) | YLCFLAFLLFLVLIM | 0.02 | 0.02 | 97.88% (231/236) | Non-Toxin |

| | | | | | | | | | |
|---|---|---|---|---|---|---|---|---|---|
| 240 | ORF7b | DPA1*01:03/DPB1*02:01 | 15 | Consensus (comb.lib./smm/nn) | YLCFLAFLLFLVLIM | 0.06 | 0.06 | 97.88% (231/236) | Non-Toxin |
| 241 | ORF7b | DPA1*01/DPB1*04:01 | 15 | Consensus (comb.lib./smm) | YLCFLAFLLFLVLIM | 0.09 | 0.09 | 97.88% (231/236) | Non-Toxin |
| S.No. | ORF | Allele | Length | Method used | Peptide | Percentile Rank | Adjusted rank | Conservancy | Toxicity |
| 242 | ORF8 | DRB3*01:01 | 15 | Consensus (comb.lib./smm/nn) | CTQHQPYVVDDPCPI | 0.08 | 0.08 | 99.17% (476/480) | Non-Toxin |
| 243 | ORF8 | DRB3*01:01 | 15 | Consensus (comb.lib./smm/nn) | HQPYVVDDPCPIHFY | 0.08 | 0.08 | 99.17% (476/480) | Non-Toxin |
| 244 | ORF8 | DQA1*01:01/DQB1*05:01 | 15 | Consensus (comb.lib./smm/nn) | LVVRCSFYEDFLEYH | 0.45 | 0.45 | 99.58% (478/480) | Non-Toxin |
| 245 | ORF8 | DRB3*01:01 | 15 | Consensus (comb.lib./smm/nn) | QHQPYVVDDPCPIHF | 0.08 | 0.08 | 99.17% (476/480) | Non-Toxin |
| 246 | ORF8 | DRB3*01:01 | 15 | Consensus (comb.lib./smm/nn) | QPYVVDDPCPIHFYS | 0.07 | 0.07 | 99.17% (476/480) | Non-Toxin |
| 247 | ORF8 | DRB3*01:01 | 15 | Consensus (comb.lib./smm/nn) | TQHQPYVVDDPCPIH | 0.08 | 0.08 | 99.17% (476/480) | Non-Toxin |
| S.No. | ORF | Allele | Length | Method used | Peptide | Percentile Rank | Adjusted rank | Conservancy | Toxicity |
| 248 | S protein | DRB3*01:01 | 15 | Consensus (comb.lib./smm/nn) | ADSFVIRGDEVRQIA | 0.49 | 0.49 | 93.63% (470/502) | Non-Toxin |
| 249 | S protein | DRB3*02:02 | 15 | NetMHCIIpan | ADYSVLYNSASFSTF | 0.85 | 0.85 | 93.82% (471/502) | Non-Toxin |
| 250 | S protein | DQA1*01:02/DQB1*06:02 | 15 | Consensus (comb.lib./smm/nn) | AGLIAIVMVTIMLCC | 1.7 | 1.7 | 93.82% (471/502) | Non-Toxin |
| 251 | S protein | DRB1*07:01 | 15 | Consensus (comb.lib./smm/nn) | AIPTNFTISVTTEIL | 0.4 | 0.4 | 93.82% (471/502) | Non-Toxin |
| 252 | S protein | DRB1*15:01 | 15 | Consensus (smm/nn/sturniolo) | CSNLLLQYGSFCTQL | 0.58 | 0.58 | 93.82% (471/502) | Non-Toxin |
| 253 | S protein | DRB3*01:01 | 15 | Consensus (comb.lib./smm/nn) | DSFVIRGDEVRQIAP | 0.51 | 0.51 | 93.63% (470/502) | Non-Toxin |
| 254 | S protein | DRB3*02:02 | 15 | NetMHCIIpan | DYSVLYNSASFSTFK | 0.68 | 0.68 | 93.82% (471/502) | Non-Toxin |
| 255 | S protein | DRB1*15:01 | 15 | Consensus (smm/nn/sturniolo) | ECSNLLLQYGSFCTQ | 0.72 | 0.72 | 93.82% (471/502) | Non-Toxin |
| 256 | S protein | DRB5*01:01 | 15 | Consensus (smm/nn/sturniolo) | EFVFKNIDGYFKIYS | 0.17 | 0.17 | 92.03% (462/502) | Non-Toxin |
| 257 | S protein | DRB3*02:02 | 15 | NetMHCIIpan | EGVFVSNGTHWFVTQ | 0.21 | 0.21 | 93.23% (468/502) | Non-Toxin |
| 258 | S protein | DRB1*04:05 | 15 | Consensus (smm/nn/sturniolo) | ESIVRFPNITNLCPF | 0.77 | 0.77 | 93.43% (469/502) | Non-Toxin |
| 259 | S protein | DRB1*15:01 | 15 | Consensus (smm/nn/sturniolo) | FQTLLALHRSYLTPG | 1.1 | 1.1 | 93.23% (468/502) | Non-Toxin |
| 260 | S protein | DRB5*01:01 | 15 | Consensus (smm/nn/sturniolo) | FVFKNIDGYFKIYSK | 0.17 | 0.17 | 92.03% (462/502) | Non-Toxin |
| 261 | S protein | DRB3*01:01 | 15 | Consensus (comb.lib./smm/nn) | FVIRGDEVRQIAPGQ | 0.54 | 0.54 | 93.63% (470/502) | Non-Toxin |
| 262 | S protein | DQA1*05:01/DQB1*03:01 | 15 | Consensus (comb.lib./smm/nn) | GFIAGLIAIVMVTIM | 1.6 | 1.6 | 93.82% (471/502) | Non-Toxin |
| 263 | S protein | DRB5*01:01 | 15 | Consensus (smm/nn/sturniolo) | GINITRFQTLLALHR | 0.52 | 0.52 | 93.43% (469/502) | Non-Toxin |
| 264 | S protein | DRB1*11:01 | 15 | Consensus (smm/nn/sturniolo) | GNYNYLYRLFRKSNL | 0.22 | 0.22 | 92.63% (465/502) | Non-Toxin |
| 265 | S protein | DQA1*01:02/DQB1*06:02 | 15 | Consensus (comb.lib./smm/nn) | IAGLIAIVMVTIMLC | 1.6 | 1.6 | 93.82% (471/502) | Non-Toxin |
| 266 | S protein | DRB1*07:01 | 15 | Consensus (comb.lib./smm/nn) | IAIPTNFTISVTTEI | 0.47 | 0.47 | 93.82% (471/502) | Non-Toxin |
| 267 | S protein | DRB5*01:01 | 15 | Consensus (smm/nn/sturniolo) | INITRFQTLLALHRS | 0.32 | 0.32 | 93.23% (468/502) | Non-Toxin |
| 268 | S protein | DRB1*07:01 | 15 | Consensus (comb.lib./smm/nn) | IPTNFTISVTTEILP | 0.52 | 0.52 | 93.82% (471/502) | Non-Toxin |
| 269 | S protein | DRB5*01:01 | 15 | Consensus (smm/nn/sturniolo) | ITRFQTLLALHRSYL | 0.26 | 0.26 | 93.23% (468/502) | Non-Toxin |
| 270 | S protein | DPA1*02:01/DPB1*14:01 | 15 | NetMHCIIpan | ITRFQTLLALHRSYL | 0.43 | 0.43 | 93.23% (468/502) | Non-Toxin |
| 271 | S protein | DQA1*05:01/DQB1*03:01 | 15 | Consensus (comb.lib./smm/nn) | IWLGFIAGLIAIVMV | 0.51 | 0.51 | 93.43% (469/502) | Non-Toxin |
| 272 | S protein | DRB3*02:02 | 15 | NetMHCIIpan | KTQSLLIVNNATNVV | 0.17 | 0.17 | 93.82% (471/502) | Non-Toxin |
| 273 | | DRB1*13:02 | 15 | Consensus (smm/nn/sturniolo) | KTQSLLIVNNATNVV | 0.01 | 0.01 | 93.82% (471/502) | Non-Toxin |
| 274 | S protein | DQA1*05:01/DQB1*03:01 | 15 | Consensus (comb.lib./smm/nn) | LGFIAGLIAIVMVTI | 1.6 | 1.6 | 93.82% (471/502) | Non-Toxin |
| 275 | S protein | DRB1*13:02 | 15 | Consensus (smm/nn/sturniolo) | LIVNNATNVVIKVCE | 0.03 | 0.03 | 93.23% (468/502) | Non-Toxin |
| 276 | S protein | DRB1*13:02 | 15 | Consensus (smm/nn/sturniolo) | LLIVNNATNVVIKVC | 0.01 | 0.01 | 93.23% (468/502) | Non-Toxin |
| 277 | S protein | DRB3*02:02 | 15 | NetMHCIIpan | LLIVNNATNVVIKVC | 0.09 | 0.09 | 93.23% (468/502) | Non-Toxin |
| 278 | S protein | DRB1*01:01 | 15 | Consensus (comb.lib./smm/nn) | LSFELLHAPATVCGP | 0.03 | 0.03 | 92.23% (463/502) | Non-Toxin |
| 279 | S protein | DRB5*01:01 | 15 | Consensus (smm/nn/sturniolo) | NITRFQTLLALHRSY | 0.32 | 0.32 | 93.23% (468/502) | Non-Toxin |
| 280 | S protein | DPA1*02:01/DPB1*14:01 | 15 | NetMHCIIpan | NITRFQTLLALHRSY | 0.45 | 0.45 | 93.23% (468/502) | Non-Toxin |
| 281 | S protein | DRB1*15:01 | 15 | Consensus (smm/nn/sturniolo) | NLLLQYGSFCTQLNR | 0.75 | 0.75 | 93.82% (471/502) | Non-Toxin |
| 282 | S protein | DRB1*15:01 | 15 | Consensus (smm/nn/sturniolo) | PTESIVRFPNITNLC | 0.64 | 0.64 | 93.43% (469/502) | Non-Toxin |
| 283 | S protein | DRB1*07:01 | 15 | Consensus (comb.lib./smm/nn) | PTNFTISVTTEILPV | 0.51 | 0.51 | 93.82% (471/502) | Non-Toxin |
| 284 | S protein | DQA1*01:01/DQB1*05:01 | 15 | Consensus (comb.lib./smm/nn) | PWYIWLGFIAGLIAI | 1.9 | 1.9 | 93.43% (469/502) | Non-Toxin |
| 285 | S protein | DPA1*02:01/DPB1*14:01 | 15 | NetMHCIIpan | QLIRAAEIRASANLA | 0.31 | 0.31 | 93.23% (468/502) | Non-Toxin |
| 286 | S protein | DRB1*15:01 | 15 | Consensus (smm/nn/sturniolo) | QPTESIVRFPNITNL | 0.69 | 0.69 | 93.43% (469/502) | Non-Toxin |
| 287 | S protein | DPA1*02:01/DPB1*14:01 | 15 | NetMHCIIpan | QQLIRAAEIRASANL | 0.2 | 0.2 | 93.23% (468/502) | Non-Toxin |
| 288 | S protein | DRB1*13:02 | 15 | Consensus (smm/nn/sturniolo) | QSLLIVNNATNVVIK | 0.01 | 0.01 | 93.63% (470/502) | Non-Toxin |

| | | | | | | | | | |
|---|---|---|---|---|---|---|---|---|---|
| 289 | S protein | DRB3*02:02 | 15 | NetMHCIIpan | QSLLIVNNATNVVIK | 0.02 | 0.02 | 93.63% (470/502) | Non-Toxin |
| 290 | S protein | DRB5*01:01 | 15 | Consensus (smm/nn/sturniolo) | REFVFKNIDGYFKIY | 0.17 | 0.17 | 91.83% (461/502) | Non-Toxin |
| 291 | S protein | DRB3*02:02 | 15 | NetMHCIIpan | REGVFVSNGTHWFVT | 0.2 | 0.2 | 93.43% (469/502) | Non-Toxin |
| 292 | S protein | DRB5*01:01 | 15 | Consensus (smm/nn/sturniolo) | RFQTLLALHRSYLTP | 0.58 | 0.58 | 93.23% (468/502) | Non-Toxin |
| 293 | S protein | DRB1*01:01 | 15 | Consensus (comb.lib./smm/nn) | SFELLHAPATVCGPK | 0.09 | 0.09 | 92.23% (463/502) | Non-Toxin |
| 294 | S protein | DRB3*01:01 | 15 | Consensus (comb.lib./smm/nn) | SFVIRGDEVRQIAPG | 0.51 | 0.51 | 93.63% (470/502) | Non-Toxin |
| 295 | S protein | DRB1*04:05 | 15 | Consensus (smm/nn/sturniolo) | SIVRFPNITNLCPFG | 0.98 | 0.98 | 93.43% (469/502) | Non-Toxin |
| 296 | S protein | DRB1*13:02 | 15 | Consensus (smm/nn/sturniolo) | SKTQSLLIVNNATNV | 0.03 | 0.03 | 93.82% (471/502) | Non-Toxin |
| 297 | S protein | DRB1*13:02 | 15 | Consensus (smm/nn/sturniolo) | SLLIVNNATNVVIKV | 0.01 | 0.01 | 93.63% (470/502) | Non-Toxin |
| 298 | S protein | DRB3*02:02 | 15 | NetMHCIIpan | SLLIVNNATNVVIKV | 0.03 | 0.03 | 93.63% (470/502) | Non-Toxin |
| 299 | S protein | DRB1*15:01 | 15 | Consensus (smm/nn/sturniolo) | SNLLLQYGSFCTQLN | 0.6 | 0.6 | 93.82% (471/502) | Non-Toxin |
| 300 | S protein | DQA1*05:01/DQB1*03:01 | 15 | Consensus (comb.lib./smm/nn) | SSGWTAGAAAYYVGY | 0.94 | 0.94 | 93.43% (469/502) | Non-Toxin |
| 301 | S protein | DQA1*05:01/DQB1*03:01 | 15 | Consensus (comb.lib./smm/nn) | SSSGWTAGAAAYYVG | 1.1 | 1.1 | | Non-Toxin |
| 302 | S protein | DRB1*15:01 | 15 | Consensus (smm/nn/sturniolo) | TESIVRFPNITNLCP | 0.69 | 0.69 | 93.43% (469/502) | Non-Toxin |
| 303 | S protein | DRB1*07:01 | 15 | Consensus (comb.lib./smm/nn) | TNFTISVTTEILPVS | 0.52 | 0.52 | 93.82% (471/502) | Non-Toxin |
| 304 | S protein | DRB1*13:02 | 15 | Consensus (smm/nn/sturniolo) | TQSLLIVNNATNVVI | 0.01 | 0.01 | 93.63% (470/502) | Non-Toxin |
| 305 | S protein | DRB3*02:02 | 15 | NetMHCIIpan | TQSLLIVNNATNVVI | 0.06 | 0.06 | 93.63% (470/502) | Non-Toxin |
| 306 | S protein | DRB5*01:01 | 15 | Consensus (smm/nn/sturniolo) | TRFQTLLALHRSYLT | 0.35 | 0.35 | 93.23% (468/502) | Non-Toxin |
| 307 | S protein | DRB1*01:01 | 15 | Consensus (comb.lib./smm/nn) | VLSFELLHAPATVCG | 0.03 | 0.03 | 92.23% (463/502) | Non-Toxin |
| 308 | S protein | DRB1*01:01 | 15 | Consensus (comb.lib./smm/nn) | VVLSFELLHAPATVC | 0.03 | 0.03 | 92.23% (463/502) | Non-Toxin |
| 309 | S protein | DRB1*01:01 | 15 | Consensus (comb.lib./smm/nn) | VVVLSFELLHAPATV | 0.09 | 0.09 | 92.23% (463/502) | Non-Toxin |
| 310 | S protein | DQA1*05:01/DQB1*03:01 | 15 | Consensus (comb.lib./smm/nn) | WLGFIAGLIAIVMVT | 1.6 | 1.6 | 93.63% (470/502) | Non-Toxin |
| 311 | S protein | DQA1*05:01/DQB1*03:01 | 15 | Consensus (comb.lib./smm/nn) | WYIWLGFIAGLIAIV | 0.58 | 0.58 | 93.43% (469/502) | Non-Toxin |
| 312 | S protein | DRB3*01:01 | 15 | Consensus (comb.lib./smm/nn) | YADSFVIRGDEVRQI | 0.49 | 0.49 | 93.63% (470/502) | Non-Toxin |
| 313 | S protein | DQA1*05:01/DQB1*03:01 | 15 | Consensus (comb.lib./smm/nn) | YIWLGFIAGLIAIVM | 0.51 | 0.51 | 93.43% (469/502) | Non-Toxin |
| 314 | S protein | DRB3*02:02 | 15 | NetMHCIIpan | YSVLYNSASFSTFKC | 0.66 | 0.66 | 93.82% (471/502) | Non-Toxin |

**Supplementary table S4:** Population coverage by all the overlapping CTL and HTL epitopes forming epitope clusters.

| population/area | Class combined | | |
|---|---|---|---|
| | coverage[a] | average_hit[b] | pc90[c] |
| Algeria | 79.67% | 20.09 | 1.97 |
| American Samoa | 98.75% | 65.28 | 38.26 |
| Argentina | 99.74% | 95.44 | 52.01 |
| Australia | 98.16% | 78.28 | 31.37 |
| Austria | 99.99% | 113.11 | 66.15 |
| Belarus | 43.81% | 3.23 | 0.71 |
| Belgium | 99.76% | 87.08 | 49.51 |
| Bolivia | 38.38% | 5.02 | 0.97 |
| Borneo | 38.38% | 6.32 | 0.97 |
| Brazil | 99.99% | 140.15 | 88.36 |
| Bulgaria | 99.68% | 95.65 | 53.93 |
| Burkina Faso | 67.18% | 32.67 | 3.96 |
| Cameroon | 99.98% | 136.25 | 72.31 |

| | | | |
|---|---|---|---|
| Canada | 89.48% | 20.21 | 3.8 |
| Cape Verde | 99.76% | 117.59 | 58.01 |
| Central Africa | 99.96% | 129.22 | 68.79 |
| Central African Republic | 66.32% | 20.16 | 1.78 |
| Central America | 80.55% | 20.65 | 3.09 |
| Chile | 99.48% | 89.01 | 43.08 |
| China | 99.83% | 121.35 | 63.51 |
| Colombia | 78.87% | 21.53 | 1.89 |
| Congo | 93.93% | 22.36 | 9.86 |
| Cook Islands | 100.00% | 76.33 | 46.33 |
| Costa Rica | 76.74% | 16.09 | 2.58 |
| Croatia | 99.99% | 126.99 | 77.32 |
| Cuba | 99.77% | 111.02 | 58.37 |
| Czech Republic | 99.95% | 117.22 | 65.85 |
| Denmark | 95.41% | 25.93 | 10.65 |
| East Africa | 99.98% | 135.39 | 74.29 |
| East Asia | 99.81% | 115.86 | 57.72 |
| Ecuador | 99.71% | 95.91 | 46.78 |
| England | 100.00% | 131.07 | 82.91 |
| Equatorial Guinea | 72.14% | 14.25 | 3.23 |
| Ethiopia | 96.17% | 29.61 | 11.44 |
| Europe | 100.00% | 170.31 | 117.87 |
| Fiji | 96.74% | 25.75 | 11.2 |
| Finland | 100.00% | 120.96 | 68.64 |
| France | 100.00% | 176.9 | 125.21 |
| Gabon | 99.95% | 73.91 | 44.53 |
| Gambia | 99.90% | 65.49 | 41.85 |
| Georgia | 99.85% | 109.46 | 57.53 |
| Germany | 100.00% | 137.81 | 86.57 |
| Greece | 87.50% | 22.14 | 3.2 |
| Guatemala | 20.40% | 4.99 | 0.75 |
| Guinea-Bissau | 98.74% | 96.3 | 36.3 |
| Hong Kong | 96.05% | 58.93 | 25.23 |
| India | 99.94% | 123.02 | 71.53 |
| Indonesia | 96.59% | 63.06 | 23.28 |
| Iran | 99.72% | 100.42 | 51.81 |
| Ireland Northern | 99.99% | 117.02 | 72.08 |
| Ireland South | 99.99% | 115.21 | 70.53 |
| Israel | 98.99% | 82.96 | 34.51 |
| Italy | 99.82% | 110.18 | 59.32 |
| Ivory Coast | 67.75% | 25.43 | 3.41 |

| | | | |
|---|---|---|---|
| Jamaica | 82.25% | 18.28 | 2.25 |
| Japan | 99.95% | 140.06 | 78.99 |
| Jordan | 98.07% | 67.52 | 25.05 |
| Kenya | 99.96% | 125.27 | 66.5 |
| Kiribati | 56.35% | 9.61 | 2.06 |
| Korea; South | 99.74% | 106.67 | 52.49 |
| Lebanon | 61.98% | 8.81 | 1.05 |
| Liberia | 91.53% | 35.2 | 26.27 |
| Macedonia | 97.86% | 37.84 | 12.81 |
| Malaysia | 90.59% | 50.93 | 6.9 |
| Mali | 96.02% | 83.84 | 22.38 |
| Martinique | 71.03% | 25.57 | 1.38 |
| Mexico | 100.00% | 149.99 | 98.41 |
| Mongolia | 99.46% | 88.15 | 46.32 |
| Morocco | 99.86% | 109.88 | 59.86 |
| Nauru | 82.88% | 15.4 | 2.34 |
| Netherlands | 87.50% | 21.51 | 3.2 |
| New Caledonia | 99.97% | 88.65 | 59.66 |
| New Zealand | 90.84% | 43.25 | 5.16 |
| Nigeria | 90.03% | 39.47 | 26.01 |
| Niue | 91.02% | 38.7 | 10.12 |
| North Africa | 99.56% | 100.23 | 45.59 |
| North America | 100.00% | 166.28 | 113.61 |
| Northeast Asia | 99.84% | 121.14 | 63.63 |
| Norway | 97.19% | 31.38 | 12.9 |
| Oceania | 99.86% | 113.14 | 62.67 |
| Oman | 99.69% | 89.6 | 48.6 |
| Pakistan | 97.13% | 56.16 | 26.19 |
| Papua New Guinea | 100.00% | 122.52 | 89.36 |
| Paraguay | 25.70% | 4.63 | 2.42 |
| Peru | 100.00% | 106.29 | 54.99 |
| Philippines | 96.26% | 64.15 | 37.87 |
| Poland | 99.99% | 127.2 | 78.58 |
| Portugal | 99.86% | 112.89 | 61.91 |
| Romania | 99.67% | 93.68 | 50.23 |
| Russia | 100.00% | 157.35 | 107.43 |
| Rwanda | 64.44% | 20.73 | 1.12 |
| Samoa | 97.31% | 52.21 | 17.85 |
| Sao Tome and Principe | 98.33% | 85.65 | 31.47 |
| Saudi Arabia | 99.51% | 90.79 | 46.85 |
| Scotland | 98.48% | 48.85 | 18.32 |

| | | | |
|---|---|---|---|
| Senegal | 96.67% | 90.69 | 29.48 |
| Serbia | 73.37% | 24.84 | 4.13 |
| Singapore | 96.73% | 65.18 | 25.2 |
| Slovenia | 99.99% | 80.78 | 59.26 |
| South Africa | 93.56% | 63.39 | 16.23 |
| South America | 99.87% | 119.99 | 66.69 |
| South Asia | 99.97% | 131.94 | 80.9 |
| Southeast Asia | 99.26% | 79.48 | 36.82 |
| Southwest Asia | 98.54% | 80.38 | 33.12 |
| Spain | 100.00% | 130.17 | 77.24 |
| Sri Lanka | 52.39% | 22.7 | 5.25 |
| Sudan | 96.25% | 72.21 | 24.29 |
| Sweden | 100.00% | 184.71 | 128.28 |
| Taiwan | 99.78% | 90.33 | 47.35 |
| Thailand | 99.04% | 71.41 | 30.17 |
| Tokelau | 94.65% | 43.18 | 44.11 |
| Tonga | 87.20% | 36.78 | 4.69 |
| Tunisia | 99.63% | 101.6 | 52.14 |
| Turkey | 97.28% | 52.55 | 17.49 |
| Uganda | 99.98% | 136.66 | 76.57 |
| Ukraine | 50.64% | 3.87 | 0.81 |
| United Kingdom | 31.77% | 8.9 | 4.1 |
| United States | 100.00% | 167.87 | 115.74 |
| Venezuela | 95.65% | 71.89 | 16.7 |
| Vietnam | 95.32% | 60.75 | 24.19 |
| West Africa | 99.99% | 154.32 | 89.11 |
| West Indies | 99.90% | 118.21 | 64.3 |
| Zambia | 98.10% | 89.47 | 33.81 |
| Zimbabwe | 99.78% | 97.75 | 38.46 |
| **World** | **99.98%** | **150.46** | **92.94** |
| **Average** | **91.11** | **79.27** | **41.42** |
| **Standard deviation** | **16.97** | **46.69** | **32.37** |

Note: Following allele(s) were not available at IEDB "Population coverage" tool database, and therefore not included in the calculation, please note that allele names are case sensitive: DPA1*01, DRB5*01:01, DRB3*02:02, DRB3*01:01, DRB4*01:01.

**a** projected population coverage
**b** average number of epitope hits / HLA combinations recognized by the population
**c** minimum number of epitope hits / HLA combinations recognized by 90% of the population

**Supplementary table S5:** Construct of CTL-MPV-1, CTL-MPV-2, CTL-MPV-3, HTL-MPV-1, and HTL-MPV-2. Physicochemical property analysis based on the amino acid sequences of all the designed three CTL and two HTL Multi-Patch Vaccines.

| S.No. | Vaccine | Vaccine constructs |
|---|---|---|
| 1 | CTL-MPV-1 construct<br><br>(Comprising of identified CTL Ag-PATCHES from Membrane and Spike protein of SARS-CoV-2): | GIGDPVTCLKSGAICHPVFCPRRYKQIGTCGLPGTKCCKKPEAAAKGTITVEELKKLL<br>EQWNLVIGFLFLTWICLLQFAYANRNRFLYIIKLIFLWLLWPVTLACFVLAAVGGGG<br>SRINWITGGIAIAMACLVGLMWLSYFIASFRLFARTRSMWSFNGGGGSKEITVATSR<br>TLSYYKLGASQRVAGDSGFAAYSRYRIGNYKLGGGGSTTRTQLPPAYTNSFTRGVYY<br>PDKVFRSSVGGGGSSTQDLFLPFFSNVTWFGGGGSVLPFNDGVYFASTEKSNIIRGW<br>IFGGGGSGVYYHKNNKSWMESEFRVYSSANNCTFEYVSQPFLGGGGSKQGNFKNLR<br>EFVFKNIDGYFKIYSKHTPIGGGGSEPLVDLPIGINITRFQTLLAGGGGSTPGDSSSGW<br>TAGAAAYYVGYLQPRTFLLKGGGGSSETKCTLKSFTVEKGIYQTSNFRGGGGSFPNIT<br>NLCPFGEVFNATRFASVGGGGSRISNCVADYSVLYNSASFSTFKCYGGGGSNVYADSF<br>VIRGDEVRGGGGSSKVGGNYNYLYRLFRKSNLKPFERGGGGSNTSNQVAVLYQDVN<br>CTEVGGGGSRVYSTGSNVFGGGGSSPRRARSVASQSIIAYGGGGSFTISVTTEILPVSM<br>TKTSVDCTMYGGGGSNTQEVFAQVKQIYKTPPIKGGGGSKRSFIEDLLFNKVTLAGG<br>GGSPLLTDEMIAQYGGGGSLQIPFAMQMAYRFNGIGGGGSFPQSAPHGVVFGGGGSF<br>VSNGTHWFVTQRGGGGSYEQYIKWPWYIWLGFIAGLIAIVEAAAKGIINTLQKYYCR<br>VRGGRCAVLSCLPKEEQIGKCSTRGRKCCRRKKHHHHHH |
| 2 | CTL-MPV-2 construct<br>(Comprising of identified CTL Ag-PATCHES from Nucleocapsid protein and ORF1ab proteins of SARS-CoV-2): | GIGDPVTCLKSGAICHPVFCPRRYKQIGTCGLPGTKCCKKPEAAAKNQRNAPRITFG<br>GPSGGGGSRSKQRRPQGLPNNTASWFTALTQHGKGGGGSNSSPDDQIGYYRRATRR<br>IRGGDGKMKDLSPRWYFYYLGGGGSLPYGANKDGIIWGGGGSKSAAEASKKPRQKR<br>TATKAYNVTQAFGGGGSQELIRQGTDYGGGGSIAQFAPSASAFFGMSRIGMEVTPSG<br>TWLTYGGGGSLLNKHIDAYKTFPPTEPKKGGGGSDEWSMATYYLFGGGGSETISLA<br>GSYKGGGGSTQVVDMSMTYGGGGSAVMYMGTLSYEQFGGGGSLVAEWFLAYILFT<br>RFFYVGGGGSRMYIFFASFYYVWKSYGGGGSAYVNTFSSTFNVPMEKGGGGSCLAYY<br>FMRFRRAFGGGGSMVMFTPLVPFWITIAYGGGGSFYWFFSNYLKRRVVFGGGGSLP<br>SLATVAYFNMVYGGGGSLILMTARTVYGGGGSTACTDDNALAYYGGGGSYTMADLV<br>YALGGGGSKLFDRYFKYWDQTYGGGGSARLYYDSMSYGGGGSVDTDFVNEFYAYLR<br>KHFSMGGGGSFPLCANGQVFGLYGGGGSIPLMYKGLPWNVVRGGGGSYVMHANYIF<br>WEAAAKGIINTLQKYYCRVRGGRCAVLSCLPKEEQIGKCSTRGRKCCRRKKHHHHH<br>H |
| 3 | CTL-MPV-3 construct<br>(Comprising of | GIGDPVTCLKSGAICHPVFCPRRYKQIGTCGLPGTKCCKKPEAAAK*SEETGTLIVNSVL*<br>*LFLAFVVFLLVTLAILTALRLCAYGGGGSVSLVKPSFYVYSRVKNLNSSRGGGGS*FMRIF<br>TIGTVTLKGGGGSTIPIQASLPFGWLIVGVALLAVFQSASKIITLKKRWGGGGSHFVC<br>NLLLLFVTVGGGGSLLVAAGLEAPFLYLYALVYFLQSINFVRIIMRLWLCWKCRGGG<br>GSYFLCWHTNCYDYGGGGSTTSPISEHDYQIGGYGGGGSYYQLYSTQLSTDTGVEHV |

| | | |
|---|---|---|
| | identified CTL Ag-PATCHES from Envelope protein, ORF3a protein, ORF6 protein, ORF7a protein, ORF7b protein, ORF8 protein, and ORF10 protein of SARS-CoV-2): | TFFIYNKIGGGGSFHLVDFQVTIAEILLIIMRTFKVSIWNLDYIIGGGGSMKIILFLALIT<br>LATCELYHYGGGGSLLKEPCSSGTYEGNSPFHPLADNKFALTCFSTQFAFACPDGVK<br>HVYQLRARSVSPKLFIRGGGGSEVQELYSPIFLIVAAIVFITLCFTLKRKGGGGSMIELS<br>LIDFYLCFLAFLLFLVLIMLIIFWFSLELGGGGSMKFLVFLGIITTVAAFGGGGSYVVD<br>DPCPIHFYSKWYIRVGARKSAPLIELCGGGGSIQYIDIGNYTVSCLPFTINCQEPKLGS<br>LVVRCSFYEDFLEYHDVRVVLGGGGSMGYINVFAFPFTIYSLLLCREAAAKGIINTLQ<br>KYYCRVRGGRCAVLSCLPKEEQIGKCSTRGRKCCRRKKHHHHHH |
| <u>4</u> | <u>HTL-MPV-1 construct</u> (Comprising of identified CTL Ag-PATCHES from Envelope protein, Membrane protein, Spike protein, Nucleocapsid protein, ORF3a protein of SARS-CoV-2): | GIGDPVTCLKSGAICHPVFCPRRYKQIGTCGLPGTKCCKKPEAAAKVNSVLLFLAFVV<br>FLLVTLAILTGGGGSIKLIFLWLLWPVTLACGGGGSVGLMWLSYFIASFRLFARGGGG<br>SSRTLSYYKLGASQRVAGDSGGGGSSKTQSLLIVNNATNVVIKVCEGGGGSREFVFKN<br>IDGYFKIYSKGGGGSGINITRFQTLLALHRSYLTPGDSSSGWTAGAAAYYVGYGGGGS<br>QPTESIVRFPNITNLCPFGGGGGSADYSVLYNSASFSTFKCGGGGSYADSFVIRGDEV<br>RQIAPGQGGGGSVVVLSFELLHAPATVCGPKGGGGSIAIPTNFTISVTTEILPVSGGGG<br>SQQLIRAAEIRASANLAGGGGSREGVFVSNGTHWFVTQGGGGSPWYIWLGFIAGLIA<br>IVMVTIMLCCGGGGSDDQIGYYRRATRRIRGGDGGGGGSGTRNPANNAAIVLQLPQG<br>TTLGGGGSWPQIAQFAPSASAFFGMSRGGGGSTPSGTWLTYTGAIKLDDKDGGGGS<br>SDFVRATATIPIQASLPGGGGSAPFLYLYALVYFLQSINFVGGGGSDTGVEHVTFFIYN<br>KIVDEPEEHVEAAAKGIINTLQKYYCRVRGGRCAVLSCLPKEEQIGKCSTRGRKCCRR<br>KKHHHHHH |
| <u>5</u> | <u>HTL-MPV-2 construct</u> (Comprising of identified CTL Ag-PATCHES from ORF1ab protein, ORF6 protein, ORF7a protein, ORF7b protein, ORF8 protein, ORF10 protein) of SARS-CoV-2): | GIGDPVTCLKSGAICHPVFCPRRYKQIGTCGLPGTKCCKKPEAAAKAIILASFSASTSA<br>FVETGGGGSYAFASEAARVVRSIFSGGGGSTLEETKFLTENLLLYIDINGGGGSVQQE<br>SPFVMMSAPPAQYELGGGGSLLQLCTFTRSTNSRIKASMGGGGSSKLINIIIWFLLLS<br>VCLGGGGSEWFLAYILFTRFFYVLGLAAIMQLFFSYFAVHFISNSWLMWLIINLVQM<br>AGGGGSVRMYIFFASFYYVWKSYVGGGGSWLKQLIKVTLVFLFVAAIFYLITPVHGG<br>GGSFSAVGNICYTPSKLIEYTGGGGSDISASIVAGGIVAIVVTCLGGGGSFTPLVPFWIT<br>IAYIICISTGGGGSVQSTQWSLFFFLYENAFLPGGGGSQAIASEFSSLPSYAAFATGGGG<br>STQMNLKYAISAKNRARTGGGGSRAMPNMLRIMASLVLARKHGGGGSFVNEFYAY<br>LRKHFSMMILGGGGSPEVKILNNLGVDIAANGGGGSSQMEIDFLELAMDEFIERYGG<br>GGSFAWWTAFVTNVNASSSGGGGSHANYIFWRNTNPIQLSSYGGGGSEILLIIMRTF<br>KVSIWNLDYIINLIIKGGGGSMKIILFLALITLATCELYGGGGSLSLIDFYLCFLAFLLFL<br>VLIMLIIFWFSLELGGGGSCTQHQPYVVDDPCPIHFYSGGGGSYINVFAFPFTIYSLLL<br>CRMEAAAKGIINTLQKYYCRVRGGRCAVLSCLPKEEQIGKCSTRGRKCCRRKKHHH<br>HHH |

**Supplementary table S6:** INF-γ inducing POSITIVE epitopes with a score of 1 or more than 1, screened from the CTL MPVs.

| S.No. | CTL MPV | Start-END | Sequence | Method | Result | Score |
|---|---|---|---|---|---|---|
| 1 | CTL-MPV-1 | 21-36 | RRYKQIGTCGLPGTK | MERCI | POSITIVE | 1 |
| 2 | CTL-MPV-1 | 22-37 | RYKQIGTCGLPGTKC | MERCI | POSITIVE | 1 |
| 3 | CTL-MPV-1 | 23-38 | YKQIGTCGLPGTKCC | MERCI | POSITIVE | 1 |
| 4 | CTL-MPV-1 | 24-39 | KQIGTCGLPGTKCCK | MERCI | POSITIVE | 1 |
| 5 | CTL-MPV-1 | 25-40 | QIGTCGLPGTKCCKK | MERCI | POSITIVE | 1 |
| 6 | CTL-MPV-1 | 41-56 | EAAAKGTITVEELKK | MERCI | POSITIVE | 1 |
| 7 | CTL-MPV-1 | 42-57 | AAAKGTITVEELKKL | MERCI | POSITIVE | 1 |
| 8 | CTL-MPV-1 | 43-58 | AAKGTITVEELKKLL | MERCI | POSITIVE | 1 |
| 9 | CTL-MPV-1 | 44-59 | AKGTITVEELKKLLE | MERCI | POSITIVE | 1 |
| 10 | CTL-MPV-1 | 150-165 | TRSMWSFNGGGGSKE | MERCI | POSITIVE | 1 |
| 11 | CTL-MPV-1 | 151-166 | RSMWSFNGGGGSKEI | MERCI | POSITIVE | 1 |
| 12 | CTL-MPV-1 | 152-167 | SMWSFNGGGGSKEIT | MERCI | POSITIVE | 1 |
| 13 | CTL-MPV-1 | 153-168 | MWSFNGGGGSKEITV | MERCI | POSITIVE | 1 |
| 14 | CTL-MPV-1 | 154-169 | WSFNGGGGSKEITVA | MERCI | POSITIVE | 1 |
| 15 | CTL-MPV-1 | 155-170 | SFNGGGGSKEITVAT | MERCI | POSITIVE | 1 |
| 16 | CTL-MPV-1 | 156-171 | FNGGGGSKEITVATS | MERCI | POSITIVE | 1 |
| 17 | CTL-MPV-1 | 157-172 | NGGGGSKEITVATSR | MERCI | POSITIVE | 1 |
| 18 | CTL-MPV-1 | 158-173 | GGGGSKEITVATSRT | MERCI | POSITIVE | 1 |
| 19 | CTL-MPV-1 | 159-174 | GGGSKEITVATSRTL | MERCI | POSITIVE | 1 |
| 20 | CTL-MPV-1 | 163-178 | KEITVATSRTLSYYK | MERCI | POSITIVE | 1 |
| 21 | CTL-MPV-1 | 164-179 | EITVATSRTLSYYKL | MERCI | POSITIVE | 1 |
| 22 | CTL-MPV-1 | 165-180 | ITVATSRTLSYYKLG | MERCI | POSITIVE | 1 |
| 23 | CTL-MPV-1 | 166-181 | TVATSRTLSYYKLGA | MERCI | POSITIVE | 1 |
| 24 | CTL-MPV-1 | 167-182 | VATSRTLSYYKLGAS | MERCI | POSITIVE | 1 |
| 25 | CTL-MPV-1 | 168-183 | ATSRTLSYYKLGASQ | MERCI | POSITIVE | 1 |
| 26 | CTL-MPV-1 | 169-184 | TSRTLSYYKLGASQR | MERCI | POSITIVE | 1 |
| 27 | CTL-MPV-1 | 170-185 | SRTLSYYKLGASQRV | MERCI | POSITIVE | 1 |
| 28 | CTL-MPV-1 | 219-234 | TNSFTRGVYYPDKVF | MERCI | POSITIVE | 1 |
| 29 | CTL-MPV-1 | 220-235 | NSFTRGVYYPDKVFR | MERCI | POSITIVE | 1 |
| 30 | CTL-MPV-1 | 221-236 | SFTRGVYYPDKVFRS | MERCI | POSITIVE | 1 |
| 31 | CTL-MPV-1 | 222-237 | FTRGVYYPDKVFRSS | MERCI | POSITIVE | 1 |
| 32 | CTL-MPV-1 | 262-277 | GSVLPFNDGVYFAST | MERCI | POSITIVE | 1 |
| 33 | CTL-MPV-1 | 263-278 | SVLPFNDGVYFASTE | MERCI | POSITIVE | 1 |
| 34 | CTL-MPV-1 | 264-279 | VLPFNDGVYFASTEK | MERCI | POSITIVE | 1 |
| 35 | CTL-MPV-1 | 265-280 | LPFNDGVYFASTEKS | MERCI | POSITIVE | 1 |
| 36 | CTL-MPV-1 | 266-281 | PFNDGVYFASTEKSN | MERCI | POSITIVE | 1 |
| 37 | CTL-MPV-1 | 267-282 | FNDGVYFASTEKSNI | MERCI | POSITIVE | 1 |
| 38 | CTL-MPV-1 | 268-283 | NDGVYFASTEKSNII | MERCI | POSITIVE | 1 |

| | | | | | | |
|---|---|---|---|---|---|---|
| 39 | CTL-MPV-1 | 269-284 | DGVYFASTEKSNIIR | MERCI | POSITIVE | 1 |
| 40 | CTL-MPV-1 | 270-285 | GVYFASTEKSNIIRG | MERCI | POSITIVE | 1 |
| 41 | CTL-MPV-1 | 271-286 | VYFASTEKSNIIRGW | MERCI | POSITIVE | 1 |
| 42 | CTL-MPV-1 | 285-300 | WIFGGGGSGVYYHKN | MERCI | POSITIVE | 3 |
| 43 | CTL-MPV-1 | 286-301 | IFGGGGSGVYYHKNN | MERCI | POSITIVE | 3 |
| 44 | CTL-MPV-1 | 287-302 | FGGGGSGVYYHKNNK | MERCI | POSITIVE | 3 |
| 45 | CTL-MPV-1 | 288-303 | GGGGSGVYYHKNNKS | MERCI | POSITIVE | 4 |
| 46 | CTL-MPV-1 | 289-304 | GGGSGVYYHKNNKSW | MERCI | POSITIVE | 4 |
| 47 | CTL-MPV-1 | 290-305 | GGSGVYYHKNNKSWM | MERCI | POSITIVE | 4 |
| 48 | CTL-MPV-1 | 291-306 | GSGVYYHKNNKSWME | MERCI | POSITIVE | 4 |
| 49 | CTL-MPV-1 | 292-307 | SGVYYHKNNKSWMES | MERCI | POSITIVE | 4 |
| 50 | CTL-MPV-1 | 293-308 | GVYYHKNNKSWMESE | MERCI | POSITIVE | 4 |
| 51 | CTL-MPV-1 | 294-309 | VYYHKNNKSWMESEF | MERCI | POSITIVE | 2 |
| 52 | CTL-MPV-1 | 343-358 | FVFKNIDGYFKIYSK | MERCI | POSITIVE | 1 |
| 53 | CTL-MPV-1 | 344-359 | VFKNIDGYFKIYSKH | MERCI | POSITIVE | 1 |
| 54 | CTL-MPV-1 | 345-360 | FKNIDGYFKIYSKHT | MERCI | POSITIVE | 1 |
| 55 | CTL-MPV-1 | 346-361 | KNIDGYFKIYSKHTP | MERCI | POSITIVE | 1 |
| 56 | CTL-MPV-1 | 347-362 | NIDGYFKIYSKHTPI | MERCI | POSITIVE | 1 |
| 57 | CTL-MPV-1 | 348-363 | IDGYFKIYSKHTPIG | MERCI | POSITIVE | 1 |
| 58 | CTL-MPV-1 | 349-364 | DGYFKIYSKHTPIGG | MERCI | POSITIVE | 1 |
| 59 | CTL-MPV-1 | 350-365 | GYFKIYSKHTPIGGG | MERCI | POSITIVE | 1 |
| 60 | CTL-MPV-1 | 351-366 | YFKIYSKHTPIGGGG | MERCI | POSITIVE | 1 |
| 61 | CTL-MPV-1 | 409-424 | YVGYLQPRTFLLKGG | MERCI | POSITIVE | 1 |
| 62 | CTL-MPV-1 | 410-425 | VGYLQPRTFLLKGGG | MERCI | POSITIVE | 1 |
| 63 | CTL-MPV-1 | 411-426 | GYLQPRTFLLKGGGG | MERCI | POSITIVE | 1 |
| 64 | CTL-MPV-1 | 412-427 | YLQPRTFLLKGGGGS | MERCI | POSITIVE | 1 |
| 65 | CTL-MPV-1 | 413-428 | LQPRTFLLKGGGGSS | MERCI | POSITIVE | 1 |
| 66 | CTL-MPV-1 | 414-429 | QPRTFLLKGGGGSSE | MERCI | POSITIVE | 2 |
| 67 | CTL-MPV-1 | 415-430 | PRTFLLKGGGGSSET | MERCI | POSITIVE | 2 |
| 68 | CTL-MPV-1 | 416-431 | RTFLLKGGGGSSETK | MERCI | POSITIVE | 2 |
| 69 | CTL-MPV-1 | 417-432 | TFLLKGGGGSSETKC | MERCI | POSITIVE | 2 |
| 70 | CTL-MPV-1 | 418-433 | FLLKGGGGSSETKCT | MERCI | POSITIVE | 2 |
| 71 | CTL-MPV-1 | 419-434 | LLKGGGGSSETKCTL | MERCI | POSITIVE | 1 |
| 72 | CTL-MPV-1 | 420-435 | LKGGGGSSETKCTLK | MERCI | POSITIVE | 1 |
| 73 | CTL-MPV-1 | 421-436 | KGGGGSSETKCTLKS | MERCI | POSITIVE | 1 |
| 74 | CTL-MPV-1 | 422-437 | GGGGSSETKCTLKSF | MERCI | POSITIVE | 1 |
| 75 | CTL-MPV-1 | 423-438 | GGGSSETKCTLKSFT | MERCI | POSITIVE | 1 |
| 76 | CTL-MPV-1 | 508-523 | GGSNVYADSFVIRGD | MERCI | POSITIVE | 2 |
| 77 | CTL-MPV-1 | 509-524 | GSNVYADSFVIRGDE | MERCI | POSITIVE | 2 |
| 78 | CTL-MPV-1 | 510-525 | SNVYADSFVIRGDEV | MERCI | POSITIVE | 2 |
| 79 | CTL-MPV-1 | 511-526 | NVYADSFVIRGDEVR | MERCI | POSITIVE | 2 |

| | | | | | | |
|---|---|---|---|---|---|---|
| 80 | CTL-MPV-1 | 512-527 | VYADSFVIRGDEVRG | MERCI | POSITIVE | 2 |
| 81 | CTL-MPV-1 | 513-528 | YADSFVIRGDEVRGG | MERCI | POSITIVE | 2 |
| 82 | CTL-MPV-1 | 514-529 | ADSFVIRGDEVRGGG | MERCI | POSITIVE | 2 |
| 83 | CTL-MPV-1 | 515-530 | DSFVIRGDEVRGGGG | MERCI | POSITIVE | 2 |
| 84 | CTL-MPV-1 | 516-531 | SFVIRGDEVRGGGGS | MERCI | POSITIVE | 2 |
| 85 | CTL-MPV-1 | 591-606 | VFGGGGSSPRRARSV | MERCI | POSITIVE | 1 |
| 86 | CTL-MPV-1 | 592-607 | FGGGGSSPRRARSVA | MERCI | POSITIVE | 1 |
| 87 | CTL-MPV-1 | 593-608 | GGGGSSPRRARSVAS | MERCI | POSITIVE | 1 |
| 88 | CTL-MPV-1 | 594-609 | GGGSSPRRARSVASQ | MERCI | POSITIVE | 1 |
| 89 | CTL-MPV-1 | 751-766 | THWFVTQRGGGGSYE | MERCI | POSITIVE | 1 |
| 90 | CTL-MPV-1 | 752-767 | HWFVTQRGGGGSYEQ | MERCI | POSITIVE | 1 |
| 91 | CTL-MPV-1 | 753768 | WFVTQRGGGGSYEQY | MERCI | POSITIVE | 1 |
| 92 | CTL-MPV-1 | 754-769 | FVTQRGGGGSYEQYI | MERCI | POSITIVE | 1 |
| 93 | CTL-MPV-1 | 755-770 | VTQRGGGGSYEQYIK | MERCI | POSITIVE | 1 |
| 94 | CTL-MPV-1 | 756-771 | TQRGGGGSYEQYIKW | MERCI | POSITIVE | 1 |
| 95 | CTL-MPV-1 | 757-772 | QRGGGGSYEQYIKWP | MERCI | POSITIVE | 1 |
| 96 | CTL-MPV-1 | 758-773 | RGGGGSYEQYIKWPW | MERCI | POSITIVE | 1 |
| 97 | CTL-MPV-1 | 759-774 | GGGGSYEQYIKWPWY | MERCI | POSITIVE | 1 |
| 98 | CTL-MPV-1 | 760-775 | GGGSYEQYIKWPWYI | MERCI | POSITIVE | 1 |
| 99 | CTL-MPV-1 | 825-840 | STRGRKCCRRKKHHH | MERCI | POSITIVE | 1 |
| 100 | CTL-MPV-1 | 826-841 | TRGRKCCRRKKHHHH | MERCI | POSITIVE | 1 |
| 101 | CTL-MPV-1 | 827-842 | RGRKCCRRKKHHHHH | MERCI | POSITIVE | 1 |
| 102 | CTL-MPV-1 | 828-843 | GRKCCRRKKHHHHHH | MERCI | POSITIVE | 2 |
| 103 | CTL-MPV-1 | 829-844 | RKCCRRKKHHHHHH | MERCI | POSITIVE | 2 |
| 104 | CTL-MPV-1 | 830-845 | KCCRRKKHHHHHH | MERCI | POSITIVE | 1 |
| 105 | CTL-MPV-1 | 831-846 | CCRRKKHHHHHH | MERCI | POSITIVE | 1 |
| 106 | CTL-MPV-1 | 832-847 | CRRKKHHHHHH | MERCI | POSITIVE | 1 |
| 107 | CTL-MPV-1 | 833-848 | RRKKHHHHHH | MERCI | POSITIVE | 1 |
| 108 | CTL-MPV-1 | 834849 | RKKHHHHHH | MERCI | POSITIVE | 1 |

| S.No. | CTL MPV | Start-END | Sequence | Method | Result | Score |
|---|---|---|---|---|---|---|
| 109 | CTL-MPV-2 | 21-36 | RRYKQIGTCGLPGTK | MERCI | POSITIVE | 1 |
| 110 | CTL-MPV-2 | 22-37 | RYKQIGTCGLPGTKC | MERCI | POSITIVE | 1 |
| 111 | CTL-MPV-2 | 23-38 | YKQIGTCGLPGTKCC | MERCI | POSITIVE | 1 |
| 112 | CTL-MPV-2 | 24-39 | KQIGTCGLPGTKCCK | MERCI | POSITIVE | 1 |
| 113 | CTL-MPV-2 | 25-40 | QIGTCGLPGTKCCKK | MERCI | POSITIVE | 1 |
| 114 | CTL-MPV-2 | 37-52 | CKKPEAAAKNQRNAP | MERCI | POSITIVE | 1 |
| 115 | CTL-MPV-2 | 66-81 | SKQRRPQGLPNNTAS | MERCI | POSITIVE | 2 |
| 116 | CTL-MPV-2 | 67-82 | KQRRPQGLPNNTASW | MERCI | POSITIVE | 2 |
| 117 | CTL-MPV-2 | 68-83 | QRRPQGLPNNTASWF | MERCI | POSITIVE | 2 |
| 118 | CTL-MPV-2 | 69-84 | RRPQGLPNNTASWFT | MERCI | POSITIVE | 2 |
| 119 | CTL-MPV-2 | 70-85 | RPQGLPNNTASWFTA | MERCI | POSITIVE | 2 |

| | | | | | | |
|---|---|---|---|---|---|---|
| 120 | CTL-MPV-2 | 71-86 | PQGLPNNTASWFTAL | MERCI | POSITIVE | 3 |
| 121 | CTL-MPV-2 | 72-87 | QGLPNNTASWFTALT | MERCI | POSITIVE | 1 |
| 122 | CTL-MPV-2 | 73-88 | GLPNNTASWFTALTQ | MERCI | POSITIVE | 1 |
| 123 | CTL-MPV-2 | 74-89 | LPNNTASWFTALTQH | MERCI | POSITIVE | 1 |
| 124 | CTL-MPV-2 | 75-90 | PNNTASWFTALTQHG | MERCI | POSITIVE | 1 |
| 125 | CTL-MPV-2 | 76-91 | NNTASWFTALTQHGK | MERCI | POSITIVE | 1 |
| 126 | CTL-MPV-2 | 77-92 | NTASWFTALTQHGKG | MERCI | POSITIVE | 1 |
| 127 | CTL-MPV-2 | 78-93 | TASWFTALTQHGKGG | MERCI | POSITIVE | 1 |
| 128 | CTL-MPV-2 | 173-188 | AYNVTQAFGGGGSQE | MERCI | POSITIVE | 1 |
| 129 | CTL-MPV-2 | 174-189 | YNVTQAFGGGGSQEL | MERCI | POSITIVE | 1 |
| 130 | CTL-MPV-2 | 175-190 | NVTQAFGGGGSQELI | MERCI | POSITIVE | 1 |
| 131 | CTL-MPV-2 | 176-191 | VTQAFGGGGSQELIR | MERCI | POSITIVE | 1 |
| 132 | CTL-MPV-2 | 177-192 | TQAFGGGGSQELIRQ | MERCI | POSITIVE | 1 |
| 133 | CTL-MPV-2 | 178-193 | QAFGGGGSQELIRQG | MERCI | POSITIVE | 1 |
| 134 | CTL-MPV-2 | 179-194 | AFGGGGSQELIRQGT | MERCI | POSITIVE | 1 |
| 135 | CTL-MPV-2 | 180-195 | FGGGGSQELIRQGTD | MERCI | POSITIVE | 1 |
| 136 | CTL-MPV-2 | 181-196 | GGGGSQELIRQGTDY | MERCI | POSITIVE | 1 |
| 137 | CTL-MPV-2 | 182-197 | GGGSQELIRQGTDYG | MERCI | POSITIVE | 1 |
| 138 | CTL-MPV-2 | 252-267 | PKKGGGGSDEWSMAT | MERCI | POSITIVE | 19 |
| 139 | CTL-MPV-2 | 253-268 | KKGGGGSDEWSMATY | MERCI | POSITIVE | 19 |
| 140 | CTL-MPV-2 | 254-269 | KGGGGSDEWSMATYY | MERCI | POSITIVE | 19 |
| 141 | CTL-MPV-2 | 255-270 | GGGGSDEWSMATYYL | MERCI | POSITIVE | 19 |
| 142 | CTL-MPV-2 | 256-271 | GGGSDEWSMATYYLF | MERCI | POSITIVE | 19 |
| 143 | CTL-MPV-2 | 257-272 | GGSDEWSMATYYLFG | MERCI | POSITIVE | 14 |
| 144 | CTL-MPV-2 | 314-329 | SYEQFGGGGSLVAEW | MERCI | POSITIVE | 2 |
| 145 | CTL-MPV-2 | 315-330 | YEQFGGGGSLVAEWF | MERCI | POSITIVE | 2 |
| 146 | CTL-MPV-2 | 316-331 | EQFGGGGSLVAEWFL | MERCI | POSITIVE | 2 |
| 147 | CTL-MPV-2 | 317-332 | QFGGGGSLVAEWFLA | MERCI | POSITIVE | 2 |
| 148 | CTL-MPV-2 | 318-333 | FGGGGSLVAEWFLAY | MERCI | POSITIVE | 2 |
| 149 | CTL-MPV-2 | 319-334 | GGGGSLVAEWFLAYI | MERCI | POSITIVE | 2 |
| 150 | CTL-MPV-2 | 320-335 | GGGSLVAEWFLAYIL | MERCI | POSITIVE | 2 |
| 151 | CTL-MPV-2 | 321-336 | GGSLVAEWFLAYILF | MERCI | POSITIVE | 2 |
| 152 | CTL-MPV-2 | 322-337 | GSLVAEWFLAYILFT | MERCI | POSITIVE | 2 |
| 153 | CTL-MPV-2 | 323-338 | SLVAEWFLAYILFTR | MERCI | POSITIVE | 1 |
| 154 | CTL-MPV-2 | 481-496 | GSTACTDDNALAYYG | MERCI | POSITIVE | 5 |
| 155 | CTL-MPV-2 | 482-497 | STACTDDNALAYYGG | MERCI | POSITIVE | 5 |
| 156 | CTL-MPV-2 | 483-498 | TACTDDNALAYYGGG | MERCI | POSITIVE | 4 |
| 157 | CTL-MPV-2 | 484-499 | ACTDDNALAYYGGGG | MERCI | POSITIVE | 1 |
| 158 | CTL-MPV-2 | 485-500 | CTDDNALAYYGGGGS | MERCI | POSITIVE | 1 |
| 159 | CTL-MPV-2 | 486-501 | TDDNALAYYGGGGSY | MERCI | POSITIVE | 1 |
| 160 | CTL-MPV-2 | 487-502 | DDNALAYYGGGGSYT | MERCI | POSITIVE | 1 |

| | | | | | | |
|---|---|---|---|---|---|---|
| 161 | CTL-MPV-2 | 544-559 | GGGGSVDTDFVNEFY | MERCI | POSITIVE | 1 |
| 162 | CTL-MPV-2 | 545-560 | GGGSVDTDFVNEFYA | MERCI | POSITIVE | 1 |
| 163 | CTL-MPV-2 | 546-561 | GGSVDTDFVNEFYAY | MERCI | POSITIVE | 1 |
| 164 | CTL-MPV-2 | 547-562 | GSVDTDFVNEFYAYL | MERCI | POSITIVE | 1 |
| 165 | CTL-MPV-2 | 658-673 | STRGRKCCRRKKHHH | MERCI | POSITIVE | 1 |
| 166 | CTL-MPV-2 | 659-674 | TRGRKCCRRKKHHHH | MERCI | POSITIVE | 1 |
| 167 | CTL-MPV-2 | 660-675 | RGRKCCRRKKHHHHH | MERCI | POSITIVE | 1 |
| 168 | CTL-MPV-2 | 661-676 | GRKCCRRKKHHHHHH | MERCI | POSITIVE | 2 |
| 169 | CTL-MPV-2 | 662-677 | RKCCRRKKHHHHHH | MERCI | POSITIVE | 2 |
| 170 | CTL-MPV-2 | 663-678 | KCCRRKKHHHHHH | MERCI | POSITIVE | 1 |
| 171 | CTL-MPV-2 | 664-679 | CCRRKKHHHHHH | MERCI | POSITIVE | 1 |
| 172 | CTL-MPV-2 | 665-680 | CRRKKHHHHHH | MERCI | POSITIVE | 1 |
| 173 | CTL-MPV-2 | 666-681 | RRKKHHHHHH | MERCI | POSITIVE | 1 |
| 174 | CTL-MPV-2 | 667-682 | RKKHHHHHH | MERCI | POSITIVE | 1 |
| **S.No.** | **CTL MPV** | **Start-END** | **Sequence** | **Method** | **Result** | **Score** |
| 175 | CTL-MPV-3 | 21-36 | RRYKQIGTCGLPGTK | MERCI | POSITIVE | 1 |
| 176 | CTL-MPV-3 | 22-37 | RYKQIGTCGLPGTKC | MERCI | POSITIVE | 1 |
| 177 | CTL-MPV-3 | 23-38 | YKQIGTCGLPGTKCC | MERCI | POSITIVE | 1 |
| 178 | CTL-MPV-3 | 24-39 | KQIGTCGLPGTKCCK | MERCI | POSITIVE | 1 |
| 179 | CTL-MPV-3 | 25-40 | QIGTCGLPGTKCCKK | MERCI | POSITIVE | 1 |
| 180 | CTL-MPV-3 | 37-52 | CKKPEAAAKSEETGT | MERCI | POSITIVE | 2 |
| 181 | CTL-MPV-3 | 38-53 | KKPEAAAKSEETGTL | MERCI | POSITIVE | 1 |
| 182 | CTL-MPV-3 | 39-54 | KPEAAAKSEETGTLI | MERCI | POSITIVE | 1 |
| 183 | CTL-MPV-3 | 40-55 | PEAAAKSEETGTLIV | MERCI | POSITIVE | 1 |
| 184 | CTL-MPV-3 | 41-56 | EAAAKSEETGTLIVN | MERCI | POSITIVE | 1 |
| 185 | CTL-MPV-3 | 42-57 | AAAKSEETGTLIVNS | MERCI | POSITIVE | 1 |
| 186 | CTL-MPV-3 | 43-58 | AAKSEETGTLIVNSV | MERCI | POSITIVE | 1 |
| 187 | CTL-MPV-3 | 118-133 | FTIGTVTLKGGGGST | MERCI | POSITIVE | 2 |
| 188 | CTL-MPV-3 | 119-134 | TIGTVTLKGGGGSTI | MERCI | POSITIVE | 2 |
| 189 | CTL-MPV-3 | 120-135 | IGTVTLKGGGGSTIP | MERCI | POSITIVE | 1 |
| 190 | CTL-MPV-3 | 121-136 | GTVTLKGGGGSTIPI | MERCI | POSITIVE | 1 |
| 191 | CTL-MPV-3 | 122-137 | TVTLKGGGGSTIPIQ | MERCI | POSITIVE | 1 |
| 192 | CTL-MPV-3 | 158-173 | SKIITLKKRWGGGGS | MERCI | POSITIVE | 1 |
| 193 | CTL-MPV-3 | 159-174 | KIITLKKRWGGGGSH | MERCI | POSITIVE | 1 |
| 194 | CTL-MPV-3 | 160-175 | IITLKKRWGGGGSHF | MERCI | POSITIVE | 1 |
| 195 | CTL-MPV-3 | 161-176 | ITLKKRWGGGGSHFV | MERCI | POSITIVE | 1 |
| 196 | CTL-MPV-3 | 162-177 | TLKKRWGGGGSHFVC | MERCI | POSITIVE | 1 |
| 197 | CTL-MPV-3 | 199-214 | APFLYLYALVYFLQS | MERCI | POSITIVE | 1 |
| 198 | CTL-MPV-3 | 200-215 | PFLYLYALVYFLQSI | MERCI | POSITIVE | 2 |
| 199 | CTL-MPV-3 | 201-216 | FLYLYALVYFLQSIN | MERCI | POSITIVE | 2 |
| 200 | CTL-MPV-3 | 202-217 | LYLYALVYFLQSINF | MERCI | POSITIVE | 2 |

| | | | | | | |
|---|---|---|---|---|---|---|
| 201 | CTL-MPV-3 | 203-218 | YLYALVYFLQSINFV | MERCI | POSITIVE | 2 |
| 202 | CTL-MPV-3 | 204-219 | LYALVYFLQSINFVR | MERCI | POSITIVE | 2 |
| 203 | CTL-MPV-3 | 205-220 | YALVYFLQSINFVRI | MERCI | POSITIVE | 2 |
| 204 | CTL-MPV-3 | 206-221 | ALVYFLQSINFVRII | MERCI | POSITIVE | 2 |
| 205 | CTL-MPV-3 | 220-235 | IMRLWLCWKCRGGGG | MERCI | POSITIVE | 1 |
| 206 | CTL-MPV-3 | 221-236 | MRLWLCWKCRGGGGS | MERCI | POSITIVE | 1 |
| 207 | CTL-MPV-3 | 222-237 | RLWLCWKCRGGGGSY | MERCI | POSITIVE | 1 |
| 208 | CTL-MPV-3 | 223-238 | LWLCWKCRGGGGSYF | MERCI | POSITIVE | 1 |
| 209 | CTL-MPV-3 | 224-239 | WLCWKCRGGGGSYFL | MERCI | POSITIVE | 1 |
| 210 | CTL-MPV-3 | 233-248 | GGSYFLCWHTNCYDY | MERCI | POSITIVE | 1 |
| 211 | CTL-MPV-3 | 270-285 | GGSYYQLYSTQLSTD | MERCI | POSITIVE | 2 |
| 212 | CTL-MPV-3 | 271-286 | GSYYQLYSTQLSTDT | MERCI | POSITIVE | 2 |
| 213 | CTL-MPV-3 | 272-287 | SYYQLYSTQLSTDTG | MERCI | POSITIVE | 2 |
| 214 | CTL-MPV-3 | 273-288 | YYQLYSTQLSTDTGV | MERCI | POSITIVE | 3 |
| 215 | CTL-MPV-3 | 274-289 | YQLYSTQLSTDTGVE | MERCI | POSITIVE | 2 |
| 216 | CTL-MPV-3 | 275-290 | QLYSTQLSTDTGVEH | MERCI | POSITIVE | 3 |
| 217 | CTL-MPV-3 | 276-291 | LYSTQLSTDTGVEHV | MERCI | POSITIVE | 2 |
| 218 | CTL-MPV-3 | 277-292 | YSTQLSTDTGVEHVT | MERCI | POSITIVE | 2 |
| 219 | CTL-MPV-3 | 278-293 | STQLSTDTGVEHVTF | MERCI | POSITIVE | 2 |
| 220 | CTL-MPV-3 | 279-294 | TQLSTDTGVEHVTFF | MERCI | POSITIVE | 2 |
| 221 | CTL-MPV-3 | 280-295 | QLSTDTGVEHVTFFI | MERCI | POSITIVE | 2 |
| 222 | CTL-MPV-3 | 281-296 | LSTDTGVEHVTFFIY | MERCI | POSITIVE | 1 |
| 223 | CTL-MPV-3 | 362-377 | GGGGSLLKEPCSSGT | MERCI | POSITIVE | 1 |
| 224 | CTL-MPV-3 | 363-378 | GGGSLLKEPCSSGTY | MERCI | POSITIVE | 1 |
| 225 | CTL-MPV-3 | 364-379 | GGSLLKEPCSSGTYE | MERCI | POSITIVE | 1 |
| 226 | CTL-MPV-3 | 365-380 | GSLLKEPCSSGTYEG | MERCI | POSITIVE | 1 |
| 227 | CTL-MPV-3 | 366-381 | SLLKEPCSSGTYEGN | MERCI | POSITIVE | 1 |
| 228 | CTL-MPV-3 | 515-530 | TVAAFGGGGSYVVDD | MERCI | POSITIVE | 1 |
| 229 | CTL-MPV-3 | 516-531 | VAAFGGGGSYVVDDP | MERCI | POSITIVE | 1 |
| 230 | CTL-MPV-3 | 517-532 | AAFGGGGSYVVDDPC | MERCI | POSITIVE | 1 |
| 231 | CTL-MPV-3 | 518-533 | AFGGGGSYVVDDPCP | MERCI | POSITIVE | 1 |
| 232 | CTL-MPV-3 | 519-534 | FGGGGSYVVDDPCPI | MERCI | POSITIVE | 1 |
| 233 | CTL-MPV-3 | 520-535 | GGGGSYVVDDPCPIH | MERCI | POSITIVE | 1 |
| 234 | CTL-MPV-3 | 521-536 | GGGSYVVDDPCPIHF | MERCI | POSITIVE | 1 |
| 235 | CTL-MPV-3 | 522-537 | GGSYVVDDPCPIHFY | MERCI | POSITIVE | 1 |
| 236 | CTL-MPV-3 | 523-538 | GSYVVDDPCPIHFYS | MERCI | POSITIVE | 1 |
| 237 | CTL-MPV-3 | 524-539 | SYVVDDPCPIHFYSK | MERCI | POSITIVE | 1 |
| 238 | CTL-MPV-3 | 525-540 | YVVDDPCPIHFYSKW | MERCI | POSITIVE | 1 |
| 239 | CTL-MPV-3 | 526-541 | VVDDPCPIHFYSKWY | MERCI | POSITIVE | 1 |
| 240 | CTL-MPV-3 | 527-542 | VDDPCPIHFYSKWYI | MERCI | POSITIVE | 1 |
| 241 | CTL-MPV-3 | 528-543 | DDPCPIHFYSKWYIR | MERCI | POSITIVE | 1 |

| | | | | | | |
|---|---|---|---|---|---|---|
| 242 | CTL-MPV-3 | 529-544 | DPCPIHFYSKWYIRV | MERCI | POSITIVE | 1 |
| 243 | CTL-MPV-3 | 530-545 | PCPIHFYSKWYIRVG | MERCI | POSITIVE | 1 |
| 244 | CTL-MPV-3 | 531-546 | CPIHFYSKWYIRVGA | MERCI | POSITIVE | 1 |
| 245 | CTL-MPV-3 | 532-547 | PIHFYSKWYIRVGAR | MERCI | POSITIVE | 1 |
| 246 | CTL-MPV-3 | 533-548 | IHFYSKWYIRVGARK | MERCI | POSITIVE | 1 |
| 247 | CTL-MPV-3 | 534-549 | HFYSKWYIRVGARKS | MERCI | POSITIVE | 1 |
| 248 | CTL-MPV-3 | 672-687 | STRGRKCCRRKKHHH | MERCI | POSITIVE | 1 |
| 249 | CTL-MPV-3 | 673-688 | TRGRKCCRRKKHHHH | MERCI | POSITIVE | 1 |
| 250 | CTL-MPV-3 | 674-689 | RGRKCCRRKKHHHHH | MERCI | POSITIVE | 1 |
| 251 | CTL-MPV-3 | 675-690 | GRKCCRRKKHHHHHH | MERCI | POSITIVE | 2 |
| 252 | CTL-MPV-3 | 676-691 | RKCCRRKKHHHHHH | MERCI | POSITIVE | 2 |
| 253 | CTL-MPV-3 | 677-692 | KCCRRKKHHHHHH | MERCI | POSITIVE | 1 |
| 254 | CTL-MPV-3 | 678-693 | CCRRKKHHHHHH | MERCI | POSITIVE | 1 |
| 255 | CTL-MPV-3 | 679-694 | CRRKKHHHHHH | MERCI | POSITIVE | 1 |
| 256 | CTL-MPV-3 | 680-695 | RRKKHHHHHH | MERCI | POSITIVE | 1 |
| 257 | CTL-MPV-3 | 681-696 | RKKHHHHHH | MERCI | POSITIVE | 1 |

**Supplementary table S7:** INF-γ inducing POSITIVE epitopes with a score of 1 or more than 1, screened from the HTL MPVs.

| S.No. | HTL MPV | Start-END | Sequence | Method | Result | Score |
|---|---|---|---|---|---|---|
| 1 | HTL-MPV-1 | 21-36 | RRYKQIGTCGLPGTK | MERCI | POSITIVE | 1 |
| 2 | HTL-MPV-1 | 22-37 | RYKQIGTCGLPGTKC | MERCI | POSITIVE | 1 |
| 3 | HTL-MPV-1 | 23-38 | YKQIGTCGLPGTKCC | MERCI | POSITIVE | 1 |
| 4 | HTL-MPV-1 | 24-39 | KQIGTCGLPGTKCCK | MERCI | POSITIVE | 1 |
| 5 | HTL-MPV-1 | 25-40 | QIGTCGLPGTKCCKK | MERCI | POSITIVE | 1 |
| 6 | HTL-MPV-1 | 37-52 | CKKPEAAAKVNSVLL | MERCI | POSITIVE | 1 |
| 7 | HTL-MPV-1 | 110-125 | ARGGGGSSRTLSYYK | MERCI | POSITIVE | 1 |
| 8 | HTL-MPV-1 | 111-126 | RGGGGSSRTLSYYKL | MERCI | POSITIVE | 1 |
| 9 | HTL-MPV-1 | 112-127 | GGGGSSRTLSYYKLG | MERCI | POSITIVE | 1 |
| 10 | HTL-MPV-1 | 113-128 | GGGSSRTLSYYKLGA | MERCI | POSITIVE | 1 |
| 11 | HTL-MPV-1 | 114-129 | GGSSRTLSYYKLGAS | MERCI | POSITIVE | 1 |
| 12 | HTL-MPV-1 | 115-130 | GSSRTLSYYKLGASQ | MERCI | POSITIVE | 1 |
| 13 | HTL-MPV-1 | 116-131 | SSRTLSYYKLGASQR | MERCI | POSITIVE | 1 |
| 14 | HTL-MPV-1 | 117-132 | SRTLSYYKLGASQRV | MERCI | POSITIVE | 1 |
| 15 | HTL-MPV-1 | 155-170 | VVIKVCEGGGGSREF | MERCI | POSITIVE | 2 |
| 16 | HTL-MPV-1 | 156-171 | VIKVCEGGGGSREFV | MERCI | POSITIVE | 1 |
| 17 | HTL-MPV-1 | 157-172 | IKVCEGGGGSREFVF | MERCI | POSITIVE | 1 |
| 18 | HTL-MPV-1 | 158-173 | KVCEGGGGSREFVFK | MERCI | POSITIVE | 1 |
| 19 | HTL-MPV-1 | 159-174 | VCEGGGGSREFVFKN | MERCI | POSITIVE | 1 |

| | | | | | | |
|---|---|---|---|---|---|---|
| 20 | HTL-MPV-1 | 160-175 | CEGGGGSREFVFKNI | MERCI | POSITIVE | 1 |
| 21 | HTL-MPV-1 | 161-176 | EGGGGSREFVFKNID | MERCI | POSITIVE | 1 |
| 22 | HTL-MPV-1 | 162-177 | GGGGSREFVFKNIDG | MERCI | POSITIVE | 1 |
| 23 | HTL-MPV-1 | 163-178 | GGGSREFVFKNIDGY | MERCI | POSITIVE | 1 |
| 24 | HTL-MPV-1 | 169-184 | FVFKNIDGYFKIYSK | MERCI | POSITIVE | 1 |
| 25 | HTL-MPV-1 | 170-185 | VFKNIDGYFKIYSKG | MERCI | POSITIVE | 1 |
| 26 | HTL-MPV-1 | 171-186 | FKNIDGYFKIYSKGG | MERCI | POSITIVE | 1 |
| 27 | HTL-MPV-1 | 172-187 | KNIDGYFKIYSKGGG | MERCI | POSITIVE | 1 |
| 28 | HTL-MPV-1 | 173-188 | NIDGYFKIYSKGGGG | MERCI | POSITIVE | 1 |
| 29 | HTL-MPV-1 | 174-189 | IDGYFKIYSKGGGGS | MERCI | POSITIVE | 1 |
| 30 | HTL-MPV-1 | 227-242 | GGGGSQPTESIVRFP | MERCI | POSITIVE | 1 |
| 31 | HTL-MPV-1 | 228-243 | GGGSQPTESIVRFPN | MERCI | POSITIVE | 5 |
| 32 | HTL-MPV-1 | 229-244 | GGSQPTESIVRFPNI | MERCI | POSITIVE | 5 |
| 33 | HTL-MPV-1 | 230-245 | GSQPTESIVRFPNIT | MERCI | POSITIVE | 8 |
| 34 | HTL-MPV-1 | 231-246 | SQPTESIVRFPNITN | MERCI | POSITIVE | 8 |
| 35 | HTL-MPV-1 | 232-247 | QPTESIVRFPNITNL | MERCI | POSITIVE | 7 |
| 36 | HTL-MPV-1 | 279-294 | YADSFVIRGDEVRQI | MERCI | POSITIVE | 2 |
| 37 | HTL-MPV-1 | 280-295 | ADSFVIRGDEVRQIA | MERCI | POSITIVE | 2 |
| 38 | HTL-MPV-1 | 281-296 | DSFVIRGDEVRQIAP | MERCI | POSITIVE | 2 |
| 39 | HTL-MPV-1 | 282-297 | SFVIRGDEVRQIAPG | MERCI | POSITIVE | 2 |
| 40 | HTL-MPV-1 | 283-298 | FVIRGDEVRQIAPGQ | MERCI | POSITIVE | 2 |
| 41 | HTL-MPV-1 | 363-378 | ANLAGGGGSREGVFV | MERCI | POSITIVE | 1 |
| 42 | HTL-MPV-1 | 364-379 | NLAGGGGSREGVFVS | MERCI | POSITIVE | 1 |
| 43 | HTL-MPV-1 | 365-380 | LAGGGGSREGVFVSN | MERCI | POSITIVE | 1 |
| 44 | HTL-MPV-1 | 366-381 | AGGGGSREGVFVSNG | MERCI | POSITIVE | 1 |
| 45 | HTL-MPV-1 | 367-382 | GGGGSREGVFVSNGT | MERCI | POSITIVE | 1 |
| 46 | HTL-MPV-1 | 368-383 | GGGSREGVFVSNGTH | MERCI | POSITIVE | 1 |
| 47 | HTL-MPV-1 | 469-484 | GGSWPQIAQFAPSAS | MERCI | POSITIVE | 1 |
| 48 | HTL-MPV-1 | 470-485 | GSWPQIAQFAPSASA | MERCI | POSITIVE | 1 |
| 49 | HTL-MPV-1 | 471-486 | SWPQIAQFAPSASAF | MERCI | POSITIVE | 1 |
| 50 | HTL-MPV-1 | 472-487 | WPQIAQFAPSASAFF | MERCI | POSITIVE | 1 |
| 51 | HTL-MPV-1 | 473-488 | PQIAQFAPSASAFFG | MERCI | POSITIVE | 1 |
| 52 | HTL-MPV-1 | 474-489 | QIAQFAPSASAFFGM | MERCI | POSITIVE | 1 |
| 53 | HTL-MPV-1 | 542-557 | APFLYLYALVYFLQS | MERCI | POSITIVE | 1 |
| 54 | HTL-MPV-1 | 543-558 | PFLYLYALVYFLQSI | MERCI | POSITIVE | 2 |
| 55 | HTL-MPV-1 | 544-559 | FLYLYALVYFLQSIN | MERCI | POSITIVE | 2 |
| 56 | HTL-MPV-1 | 545-560 | LYLYALVYFLQSINF | MERCI | POSITIVE | 2 |
| 57 | HTL-MPV-1 | 546-561 | YLYALVYFLQSINFV | MERCI | POSITIVE | 2 |
| 58 | HTL-MPV-1 | 547-562 | LYALVYFLQSINFVG | MERCI | POSITIVE | 2 |
| 59 | HTL-MPV-1 | 548-563 | YALVYFLQSINFVGG | MERCI | POSITIVE | 2 |
| 60 | HTL-MPV-1 | 549-564 | ALVYFLQSINFVGGG | MERCI | POSITIVE | 2 |

| | | | | | | |
|---|---|---|---|---|---|---|
| 61 | HTL-MPV-1 | 550-565 | LVYFLQSINFVGGGG | MERCI | POSITIVE | 2 |
| 62 | HTL-MPV-1 | 551-566 | VYFLQSINFVGGGGS | MERCI | POSITIVE | 2 |
| 63 | HTL-MPV-1 | 568-583 | GVEHVTFFIYNKIVD | MERCI | POSITIVE | 2 |
| 64 | HTL-MPV-1 | 569-584 | VEHVTFFIYNKIVDE | MERCI | POSITIVE | 4 |
| 65 | HTL-MPV-1 | 570-585 | EHVTFFIYNKIVDEP | MERCI | POSITIVE | 4 |
| 66 | HTL-MPV-1 | 571-586 | HVTFFIYNKIVDEPE | MERCI | POSITIVE | 4 |
| 67 | HTL-MPV-1 | 572-587 | VTFFIYNKIVDEPEE | MERCI | POSITIVE | 4 |
| 68 | HTL-MPV-1 | 573-588 | TFFIYNKIVDEPEEH | MERCI | POSITIVE | 4 |
| 69 | HTL-MPV-1 | 574-589 | FFIYNKIVDEPEEHV | MERCI | POSITIVE | 2 |
| 70 | HTL-MPV-1 | 575-590 | FIYNKIVDEPEEHVE | MERCI | POSITIVE | 1 |
| 71 | HTL-MPV-1 | 579-594 | KIVDEPEEHVEAAAK | MERCI | POSITIVE | 1 |
| 72 | HTL-MPV-1 | 580-595 | IVDEPEEHVEAAAKG | MERCI | POSITIVE | 1 |
| 73 | HTL-MPV-1 | 581-596 | VDEPEEHVEAAAKGI | MERCI | POSITIVE | 1 |
| 74 | HTL-MPV-1 | 582-597 | STRGRKCCRRKKHHH | MERCI | POSITIVE | 1 |
| 75 | HTL-MPV-1 | 583-598 | TRGRKCCRRKKHHHH | MERCI | POSITIVE | 1 |
| 76 | HTL-MPV-1 | 584-599 | RGRKCCRRKKHHHHH | MERCI | POSITIVE | 1 |
| 77 | HTL-MPV-1 | 585-600 | GRKCCRRKKHHHHHH | MERCI | POSITIVE | 2 |
| 78 | HTL-MPV-1 | 5860-601 | RKCCRRKKHHHHHH | MERCI | POSITIVE | 2 |
| 79 | HTL-MPV-1 | 587-602 | KCCRRKKHHHHHH | MERCI | POSITIVE | 1 |
| 80 | HTL-MPV-1 | 588-603 | CCRRKKHHHHHH | MERCI | POSITIVE | 1 |
| 81 | HTL-MPV-1 | 589-604 | CRRKKHHHHHH | MERCI | POSITIVE | 1 |
| 82 | HTL-MPV-1 | 590-605 | RRKKHHHHHH | MERCI | POSITIVE | 1 |
| 83 | HTL-MPV-1 | 591-606 | RKKHHHHHH | MERCI | POSITIVE | 1 |
| **S.No.** | **CTL MPV** | **Start-END** | **Sequence** | **Method** | **Result** | **Score** |
| 84 | HTL-MPV-2 | 21-36 | RRYKQIGTCGLPGTK | MERCI | POSITIVE | 1 |
| 85 | HTL-MPV-2 | 22-37 | RYKQIGTCGLPGTKC | MERCI | POSITIVE | 1 |
| 86 | HTL-MPV-2 | 23-38 | YKQIGTCGLPGTKCC | MERCI | POSITIVE | 1 |
| 87 | HTL-MPV-2 | 24-39 | KQIGTCGLPGTKCCK | MERCI | POSITIVE | 1 |
| 88 | HTL-MPV-2 | 25-40 | QIGTCGLPGTKCCKK | MERCI | POSITIVE | 1 |
| 89 | HTL-MPV-2 | 114-129 | QQESPFVMMSAPPAQ | MERCI | POSITIVE | 2 |
| 90 | HTL-MPV-2 | 137-152 | LLQLCTFTRSTNSRI | MERCI | POSITIVE | 1 |
| 91 | HTL-MPV-2 | 138-153 | LQLCTFTRSTNSRIK | MERCI | POSITIVE | 1 |
| 92 | HTL-MPV-2 | 139-154 | QLCTFTRSTNSRIKA | MERCI | POSITIVE | 1 |
| 93 | HTL-MPV-2 | 140-155 | LCTFTRSTNSRIKAS | MERCI | POSITIVE | 1 |
| 94 | HTL-MPV-2 | 170-185 | FLLLSVCLGGGGSEW | MERCI | POSITIVE | 1 |
| 95 | HTL-MPV-2 | 171-186 | LLLSVCLGGGGSEWF | MERCI | POSITIVE | 1 |
| 96 | HTL-MPV-2 | 172-187 | LLSVCLGGGGSEWFL | MERCI | POSITIVE | 1 |
| 97 | HTL-MPV-2 | 173-188 | LSVCLGGGGSEWFLA | MERCI | POSITIVE | 1 |
| 98 | HTL-MPV-2 | 174-189 | SVCLGGGGSEWFLAY | MERCI | POSITIVE | 1 |
| 99 | HTL-MPV-2 | 175-190 | VCLGGGGSEWFLAYI | MERCI | POSITIVE | 1 |
| 100 | HTL-MPV-2 | 176-191 | CLGGGGSEWFLAYIL | MERCI | POSITIVE | 1 |

| | | | | | | |
|---|---|---|---|---|---|---|
| 101 | HTL-MPV-2 | 177-192 | LGGGGSEWFLAYILF | MERCI | POSITIVE | 1 |
| 102 | HTL-MPV-2 | 178-193 | GGGGSEWFLAYILFT | MERCI | POSITIVE | 1 |
| 103 | HTL-MPV-2 | 179-194 | GGGSEWFLAYILFTR | MERCI | POSITIVE | 1 |
| 104 | HTL-MPV-2 | 185-200 | FLAYILFTRFFYVLG | MERCI | POSITIVE | 1 |
| 105 | HTL-MPV-2 | 186-201 | LAYILFTRFFYVLGL | MERCI | POSITIVE | 1 |
| 106 | HTL-MPV-2 | 187-202 | AYILFTRFFYVLGLA | MERCI | POSITIVE | 1 |
| 107 | HTL-MPV-2 | 188-203 | YILFTRFFYVLGLAA | MERCI | POSITIVE | 1 |
| 108 | HTL-MPV-2 | 198-213 | LGLAAIMQLFFSYFA | MERCI | POSITIVE | 1 |
| 109 | HTL-MPV-2 | 199-214 | GLAAIMQLFFSYFAV | MERCI | POSITIVE | 1 |
| 110 | HTL-MPV-2 | 200-215 | LAAIMQLFFSYFAVH | MERCI | POSITIVE | 1 |
| 111 | HTL-MPV-2 | 201-216 | AAIMQLFFSYFAVHF | MERCI | POSITIVE | 1 |
| 112 | HTL-MPV-2 | 202-217 | AIMQLFFSYFAVHFI | MERCI | POSITIVE | 1 |
| 113 | HTL-MPV-2 | 203-218 | IMQLFFSYFAVHFIS | MERCI | POSITIVE | 1 |
| 114 | HTL-MPV-2 | 204-219 | MQLFFSYFAVHFISN | MERCI | POSITIVE | 2 |
| 115 | HTL-MPV-2 | 205-220 | QLFFSYFAVHFISNS | MERCI | POSITIVE | 2 |
| 116 | HTL-MPV-2 | 206-221 | LFFSYFAVHFISNSW | MERCI | POSITIVE | 2 |
| 117 | HTL-MPV-2 | 207-222 | FFSYFAVHFISNSWL | MERCI | POSITIVE | 1 |
| 118 | HTL-MPV-2 | 364-379 | VQSTQWSLFFFLYEN | MERCI | POSITIVE | 1 |
| 119 | HTL-MPV-2 | 365-380 | QSTQWSLFFFLYENA | MERCI | POSITIVE | 1 |
| 120 | HTL-MPV-2 | 366-381 | STQWSLFFFLYENAF | MERCI | POSITIVE | 1 |
| 121 | HTL-MPV-2 | 367-382 | TQWSLFFFLYENAFL | MERCI | POSITIVE | 1 |
| 122 | HTL-MPV-2 | 368-383 | QWSLFFFLYENAFLP | MERCI | POSITIVE | 1 |
| 123 | HTL-MPV-2 | 369-384 | WSLFFFLYENAFLPG | MERCI | POSITIVE | 1 |
| 124 | HTL-MPV-2 | 370-385 | SLFFFLYENAFLPGG | MERCI | POSITIVE | 1 |
| 125 | HTL-MPV-2 | 371-386 | LFFFLYENAFLPGGG | MERCI | POSITIVE | 1 |
| 126 | HTL-MPV-2 | 372-387 | FFFLYENAFLPGGGG | MERCI | POSITIVE | 1 |
| 127 | HTL-MPV-2 | 381-396 | LPGGGGSQAIASEFS | MERCI | POSITIVE | 2 |
| 128 | HTL-MPV-2 | 382-397 | PGGGGSQAIASEFSS | MERCI | POSITIVE | 2 |
| 129 | HTL-MPV-2 | 383-398 | GGGGSQAIASEFSSL | MERCI | POSITIVE | 2 |
| 130 | HTL-MPV-2 | 384-399 | GGGSQAIASEFSSLP | MERCI | POSITIVE | 2 |
| 131 | HTL-MPV-2 | 385-400 | GGSQAIASEFSSLPS | MERCI | POSITIVE | 2 |
| 132 | HTL-MPV-2 | 386-401 | GSQAIASEFSSLPSY | MERCI | POSITIVE | 2 |
| 133 | HTL-MPV-2 | 387-402 | SQAIASEFSSLPSYA | MERCI | POSITIVE | 2 |
| 134 | HTL-MPV-2 | 388-403 | QAIASEFSSLPSYAA | MERCI | POSITIVE | 2 |
| 135 | HTL-MPV-2 | 389-404 | AIASEFSSLPSYAAF | MERCI | POSITIVE | 2 |
| 136 | HTL-MPV-2 | 409-424 | GSTQMNLKYAISAKN | MERCI | POSITIVE | 1 |
| 137 | HTL-MPV-2 | 410-425 | STQMNLKYAISAKNR | MERCI | POSITIVE | 1 |
| 138 | HTL-MPV-2 | 411-426 | TQMNLKYAISAKNRA | MERCI | POSITIVE | 1 |
| 139 | HTL-MPV-2 | 412-427 | QMNLKYAISAKNRAR | MERCI | POSITIVE | 1 |
| 140 | HTL-MPV-2 | 413-428 | MNLKYAISAKNRART | MERCI | POSITIVE | 1 |
| 141 | HTL-MPV-2 | 414-429 | NLKYAISAKNRARTG | MERCI | POSITIVE | 1 |

| | | | | | | |
|---|---|---|---|---|---|---|
| 142 | HTL-MPV-2 | 415-430 | LKYAISAKNRARTGG | MERCI | POSITIVE | 1 |
| 143 | HTL-MPV-2 | 448-463 | ARKHGGGGSFVNEFY | MERCI | POSITIVE | 2 |
| 144 | HTL-MPV-2 | 449-464 | RKHGGGGSFVNEFYA | MERCI | POSITIVE | 2 |
| 145 | HTL-MPV-2 | 450-465 | KHGGGGSFVNEFYAY | MERCI | POSITIVE | 2 |
| 146 | HTL-MPV-2 | 451-466 | HGGGGSFVNEFYAYL | MERCI | POSITIVE | 2 |
| 147 | HTL-MPV-2 | 452-467 | GGGGSFVNEFYAYLR | MERCI | POSITIVE | 2 |
| 148 | HTL-MPV-2 | 453-468 | GGGSFVNEFYAYLRK | MERCI | POSITIVE | 2 |
| 149 | HTL-MPV-2 | 454-469 | GGSFVNEFYAYLRKH | MERCI | POSITIVE | 2 |
| 150 | HTL-MPV-2 | 455-470 | GSFVNEFYAYLRKHF | MERCI | POSITIVE | 2 |
| 151 | HTL-MPV-2 | 456-471 | SFVNEFYAYLRKHFS | MERCI | POSITIVE | 2 |
| 152 | HTL-MPV-2 | 518-533 | RYGGGGSFAWWTAFV | MERCI | POSITIVE | 1 |
| 153 | HTL-MPV-2 | 519-534 | YGGGGSFAWWTAFVT | MERCI | POSITIVE | 1 |
| 154 | HTL-MPV-2 | 520-535 | GGGGSFAWWTAFVTN | MERCI | POSITIVE | 1 |
| 155 | HTL-MPV-2 | 521-536 | GGGSFAWWTAFVTNV | MERCI | POSITIVE | 1 |
| 156 | HTL-MPV-2 | 522-537 | GGSFAWWTAFVTNVN | MERCI | POSITIVE | 1 |
| 157 | HTL-MPV-2 | 523-538 | GSFAWWTAFVTNVNA | MERCI | POSITIVE | 1 |
| 158 | HTL-MPV-2 | 524-539 | SFAWWTAFVTNVNAS | MERCI | POSITIVE | 1 |
| 159 | HTL-MPV-2 | 525-540 | FAWWTAFVTNVNASS | MERCI | POSITIVE | 1 |
| 160 | HTL-MPV-2 | 526-541 | AWWTAFVTNVNASSS | MERCI | POSITIVE | 1 |
| 161 | HTL-MPV-2 | 655-670 | GGGSCTQHQPYVVDD | MERCI | POSITIVE | 1 |
| 162 | HTL-MPV-2 | 656-671 | GGSCTQHQPYVVDDP | MERCI | POSITIVE | 1 |
| 163 | HTL-MPV-2 | 657-672 | GSCTQHQPYVVDDPC | MERCI | POSITIVE | 1 |
| 164 | HTL-MPV-2 | 658-673 | SCTQHQPYVVDDPCP | MERCI | POSITIVE | 1 |
| 165 | HTL-MPV-2 | 659-674 | CTQHQPYVVDDPCPI | MERCI | POSITIVE | 1 |
| 166 | HTL-MPV-2 | 660-675 | TQHQPYVVDDPCPIH | MERCI | POSITIVE | 1 |
| 167 | HTL-MPV-2 | 661-676 | QHQPYVVDDPCPIHF | MERCI | POSITIVE | 1 |
| 168 | HTL-MPV-2 | 662-677 | HQPYVVDDPCPIHFY | MERCI | POSITIVE | 1 |
| 169 | HTL-MPV-2 | 663-678 | QPYVVDDPCPIHFYS | MERCI | POSITIVE | 1 |
| 170 | HTL-MPV-2 | 664-679 | PYVVDDPCPIHFYSG | MERCI | POSITIVE | 1 |
| 171 | HTL-MPV-2 | 665-680 | YVVDDPCPIHFYSGG | MERCI | POSITIVE | 1 |
| 172 | HTL-MPV-2 | 740-755 | STRGRKCCRRKKHHH | MERCI | POSITIVE | 1 |
| 173 | HTL-MPV-2 | 741-756 | TRGRKCCRRKKHHHH | MERCI | POSITIVE | 1 |
| 174 | HTL-MPV-2 | 742-757 | RGRKCCRRKKHHHHH | MERCI | POSITIVE | 1 |
| 175 | HTL-MPV-2 | 743-758 | GRKCCRRKKHHHHHH | MERCI | POSITIVE | 2 |
| 176 | HTL-MPV-2 | 744-759 | RKCCRRKKHHHHHH | MERCI | POSITIVE | 2 |
| 177 | HTL-MPV-2 | 745-760 | KCCRRKKHHHHHH | MERCI | POSITIVE | 1 |
| 178 | HTL-MPV-2 | 746-761 | CCRRKKHHHHHH | MERCI | POSITIVE | 1 |
| 179 | HTL-MPV-2 | 747-762 | CRRKKHHHHHH | MERCI | POSITIVE | 1 |
| 180 | HTL-MPV-2 | 748-763 | RRKKHHHHHH | MERCI | POSITIVE | 1 |
| 181 | HTL-MPV-2 | 749-764 | RKKHHHHHH | MERCI | POSITIVE | 1 |

**Supplementary table S8:** Parameters for the tertiary structure homology modeling of all the CTL and HTL MPVs by the I-TASSER tool.

| S.No. | MPVs | PDB hit | C-Score | TM-Score | RMSD (Å) |
|---|---|---|---|---|---|
| 1 | CTL-MPV-1 | 6cv0A | -2.39 | 0.43 ± 0.14 | 14.5 ± 3.7 |
| 2 | CTL-MPV-2 | 5n8pA | -0.64 | 0.63 ± 0.13 | 9.5 ± 4.6 |
| 3 | CTL-MPV-3 | 6s7tA | -1.23 | 0.56 ± 0.15 | 11.0 ± 4.6 |
| 4 | HTL-MPV-1 | 6p2mA | -1.67 | 0.51 ± 0.15 | 11.9 ±4.4 |
| 5 | HTL-MPV-2 | 5kdvA | -0.82 | 0.61 ± 0.14 | 10.2 ± 4.6 |

**Supplementary table S9:** Refinement parameter values for CTL and HTL MPV models after refinement by GalaxyRefine tool.

| S.No. | MPV Models | GDT-HA | RMSD | MolProbity | Clash score | Poor rotamers | Rama favored |
|---|---|---|---|---|---|---|---|
| 1 | CTL-MPV-1 | 0.8652 | 0.632 | 2.594 | 18.6 | 1.7 | 85.2 |
| 2 | CTL-MPV-2 | 0.9452 | 0.427 | 2.55 | 22.3 | 1.4 | 88.3 |
| 3 | CTL-MPV-3 | 0.9311 | 0.474 | 2.398 | 20.4 | 0.7 | 88.2 |
| 4 | HTL-MPV-1 | 0.9239 | 0.477 | 2.577 | 26.6 | 1.1 | 85.7 |
| 5 | HTL-MPV-2 | 0.9488 | 0.425 | 2.494 | 22.1 | 1.4 | 89.8 |

**MolProbity** score indicates the log-weighted combination score of the clash score, the percentage of Ramachandran not favored residues and the percentage of bad side-chain rotamers
**Clash score:** number of atomic clashes per 1000 atoms.
**Poor rotamers:** the percentages of rotamer outliers
**RMSD** value in Å indicated deviation from initial model.
**GDT-HA** (global distance test-High Accuracy): backbone structure accuracy measured by GDT-HA
**Rama favored** is the percentage of residues which come in the favored region of the Ramachandran plot

**Supplementary table S10:** B cell linear epitopes screened from CTL MPVs.

| CTL MPV | No. | Start | End | B Cell Linear Epitopes | Number of residues | Score |
|---|---|---|---|---|---|---|
| CTL-MPV-1 | 1 | 309 | 372 | FRVYSSANNCTFEYVSQPFLGGGGSKQGNFKNLREFVFKNIDGYFKIYSKHTPIGGGGSEPLVD | 64 | 0.773 |
| CTL-MPV-1 | 2 | 380 | 440 | TRFQTLLAGGGGSTPGDSSSGWTAGAAAYYVGYLQPRTFLLKGGGGSSETKCTLKSFTVEK | 61 | 0.725 |
| CTL-MPV-1 | 3 | 38 | 94 | CKKPEAAAKGTITVEELKKLLEQWNLVIGFLFLTWICLLQFAYANRNRFLYIIKLIF | 57 | 0.787 |
| CTL-MPV-1 | 4 | 116 | 165 | SRINWITGGIAIAMACLVGLMWLSYFIASFRLFARTRSMWSFNGGGGSKE | 50 | 0.675 |
| CTL-MPV-1 | 5 | 701 | 743 | AQYGGGGSLQIPFAMQMAYRFNGIGGGGSFPQSAPHGVVFGGG | 43 | 0.785 |
| CTL-MPV-1 | 6 | 800 | 842 | YYCRVRGGRCAVLSCLPKEEQIGKCSTRGRKCCRRKKHHHHHH | 43 | 0.776 |
| CTL-MPV-1 | 7 | 587 | 624 | TGSNVFGGGGSSPRRARSVASQSIIAYGGGGSFTISVT | 38 | 0.805 |
| CTL-MPV-1 | 8 | 455 | 482 | FPNITNLCPFGEVFNATRFASVGGGGSR | 28 | 0.653 |

| | | | | | | |
|---|---|---|---|---|---|---|
| CTL-MPV-1 | 9 | 654 | 676 | AQVKQIYKTPPIKGGGGSKRSFI | 23 | 0.557 |
| CTL-MPV-1 | 10 | 501 | 513 | TFKCYGGGGSNVY | 13 | 0.71 |
| CTL-MPV-1 | 11 | 182 | 190 | SQRVAGDSG | 9 | 0.528 |
| CTL-MPV-1 | 12 | 644 | 651 | GGGSNTQE | 8 | 0.584 |
| **CTL MPV** | **No.** | **Start** | **End** | **B Cell Linear Epitopes** | **Number of residues** | **Score** |
| CTL-MPV-2 | 1 | 577 | 675 | ANGQVFGLYGGGGSIPLMYKGLPWNVVRGGGGSYVMHANYIFWEAAAKGIINTLQKYYCRVRGGRCAVLSCLPKEEQIGKCSTRGRKCCRRKKHHHHHH | 99 | 0.801 |
| CTL-MPV-2 | 2 | 1 | 105 | GIGDPVTCLKSGAICHPVFCPRRYKQIGTCGLPGTKCCKKPEAAAKNQRNAPRITFGGPSGGGGSRSKQRRPQGLPNNTASWFTALTQHGKGGGGSNSSPDDQIG | 105 | 0.789 |
| CTL-MPV-2 | 3 | 351 | 367 | IFFASFYYVWKSYGGGG | 17 | 0.732 |
| CTL-MPV-2 | 4 | 332 | 343 | AYILFTRFFYVG | 12 | 0.707 |
| CTL-MPV-2 | 5 | 374 | 389 | FSSTFNVPMEKGGGGS | 16 | 0.677 |
| CTL-MPV-2 | 6 | 248 | 259 | FPPTEPKKGGGG | 12 | 0.65 |
| CTL-MPV-2 | 7 | 188 | 198 | ELIRQGTDYGG | 11 | 0.634 |
| CTL-MPV-2 | 8 | 310 | 321 | YMGTLSYEQFGG | 12 | 0.613 |
| CTL-MPV-2 | 9 | 442 | 451 | VFGGGGSLPS | 10 | 0.6 |
| CTL-MPV-2 | 10 | 548 | 566 | SVDTDFVNEFYAYLRKHFS | 19 | 0.597 |
| CTL-MPV-2 | 11 | 272 | 278 | GGGGSET | 7 | 0.588 |
| CTL-MPV-2 | 12 | 396 | 406 | MRFRRAFGGGG | 11 | 0.566 |
| CTL-MPV-2 | 13 | 292 | 301 | TQVVDMSMTY | 10 | 0.535 |
| CTL-MPV-2 | 14 | 527 | 534 | TYGGGGSA | 8 | 0.533 |
| **CTL MPV** | **No.** | **Start** | **End** | **B Cell Linear Epitopes** | **Number of residues** | **Score** |
| CTL-MPV-3 | 1 | 627 | 689 | YSLLLCREAAAKGIINTLQKYYCRVRGGRCAVLSCLPKEEQIGKCSTRGRKCCRRKKHHHHHH | 63 | 0.843 |
| CTL-MPV-3 | 2 | 316 | 365 | EILLIIMRTFKVSIWNLDYIIGGGGSMKIILFLALITLATCELYHYGGGG | 50 | 0.814 |
| CTL-MPV-3 | 3 | 18 | 61 | VFCPRRYKQIGTCGLPGTKCCKKPEAAAKSEETGTLIVNSVLLF | 44 | 0.742 |
| CTL-MPV-3 | 4 | 481 | 515 | LLFLVLIMLIIFWFSLELGGGGSMKFLVFLGIITT | 35 | 0.78 |
| CTL-MPV-3 | 5 | 590 | 608 | VRCSFYEDFLEYHDVRVVL | 19 | 0.726 |
| CTL-MPV-3 | 6 | 106 | 122 | NSSRGGGGSFMRIFTIG | 17 | 0.649 |
| CTL-MPV-3 | 7 | 537 | 551 | SKWYIRVGARKSAPL | 15 | 0.645 |
| CTL-MPV-3 | 8 | 152 | 163 | LAVFQSASKIIT | 12 | 0.734 |
| CTL-MPV-3 | 9 | 367 | 378 | LLKEPCSSGTYE | 12 | 0.703 |
| CTL-MPV-3 | 10 | 75 | 86 | LTALRLCAYGGG | 12 | 0.627 |
| CTL-MPV-3 | 11 | 277 | 288 | LYSTQLSTDTGV | 12 | 0.605 |
| CTL-MPV-3 | 12 | 126 | 134 | LKGGGGSTI | 9 | 0.642 |
| CTL-MPV-3 | 13 | 231 | 239 | RGGGGSYFL | 9 | 0.522 |
| CTL-MPV-3 | 14 | 431 | 438 | SEVQELYS | 8 | 0.59 |
| CTL-MPV-3 | 15 | 522 | 529 | GGSYVVDD | 8 | 0.532 |

**Supplementary table S11:** B cell Discontinuous epitopes screened from CTL MPVs.

| CTL MPV | No. | B Cell Discontinuous Epitopes residues | Number of residues | Score |
|---|---|---|---|---|
| CTL-MPV-1 | 1 | H838, H839, H840, H841 | 4 | 0.901 |
| CTL-MPV-1 | 2 | T587, G588, S589, N590, V591, F592, G593, G594, G595, G596, S597, S598, P599, R600, R601, A602, R603, S604, V605, A606, S607, Q608, S609, I610, I611, A612, Y613, G614, G615, G616, G617, S618, F619, T620, I621, S622, V623, T624, E626, A714, M715, Q716, M717, A718, Y719, R720, F721, N722, G723, I724, G725, G726, G727, G728, S729, F730, P731, Q732, S733, A734, P735, H736, G737, V738, V739, F740, G741, G742, G743, L797, Y800, Y801, C802, R803, V804, R805, G806, G807, R808, C809, A810, V811, L812, S813, C814, L815, P816, K817, E818, E819, Q820, I821, G822, K823, C824, S825, T826, R827, G828, R829, K830, C831, C832, R833, K835, K836, H837 | 107 | 0.789 |
| CTL-MPV-1 | 3 | F309, R310, V311, Y312, S313, S314, A315, N316, N317, C318, T319, F320, E321, Y322, V323, S324, Q325, P326, F327, L328, G329, G330, G331, G332, S333, K334, Q335, G336, N337, F338, K339, N340, L341, R342, E343, V345, F346, K347, N348, I349, D350, G351, Y352, F353, K354, I355, Y356, S357, K358, H359, T360, P361, I362, G363, G364, G365, G366, S367, E368, P369, L370, V371, D372, L373, T380, R381, F382, Q383, T384, L385, L386, A387, G388, G389, G390, G391, S392, T393, P394, G395, D396, | 169 | 0.723 |

| | | | | |
|---|---|---|---|---|
| | | S397, S398, S399, G400, W401, T402, A403, G404, A405, A406, A407, Y408, Y409, V410, G411, Y412, L413, Q414, P415, R416, T417, F418, L419, L420, K421, G422, G423, G424, G425, S426, S427, E428, T429, K430, C431, T432, L433, K434, S435, F436, T437, V438, E439, K440, N447, F448, F455, P456, N457, I458, T459, N460, L461, C462, P463, F464, G465, E466, V467, F468, N469, A470, T471, R472, F473, A474, S475, V476, G477, G478, G479, G480, S481, R482, I483, T501, F502, K503, C504, Y505, G506, G507, G508, G509, S510, N511, V512, Y513 | | |
| CTL-MPV-1 | 4 | K39, K40, P41, E42, A43, A44, A45, K46, G47, T48, I49, T50, V51, E52, E53, L54, K55, K56, L57, L58, E59, Q60, W61, N62, L63, V64, I65, G66, F67, L68, F69, L70, T71, W72, I73, C74, L75, L76, Q77, F78, A79, Y80, A81, N82, R83, N84, R85, F86, L87, Y88, I89, I90, K91, L92, I93, F94, L95, G113, G114, G115, R117, I118, N119, W120, I121, T122, G123, G124, I125, A126, I127, A128, M129, A130, C131, L132, V133, G134, L135, M136, W137, L138, S139, Y140, F141, I142, A143, S144, F145, R146, V185, A186, G187, D188, S189, G190, F191, Y194, D231, V233, R235, T245, Q246, D247, L248 | 105 | 0.721 |
| CTL-MPV-1 | 5 | K666, G667, G668, G669, G670, S671, K672, R673, S674, F675, A701, Q702, G704, G705, G706, G707, S708, L709, Q710, I711, P712, F713 | 22 | 0.636 |
| CTL-MPV-1 | 6 | R152, S153, M154, W155, S156, F157, N158, G159, G160, G161, G162, S163, K164, E165, S209, T210, T211, R212 | 18 | 0.57 |
| CTL-MPV-1 | 7 | S636, V637, D638, T640, G643, G644, G645, G646, S647, N648, Q650, E651, A654, Q655, K657, Q658, I659, Y660 | 18 | 0.554 |
| CTL MPV | **No.** | **B Cell Discontinuous Epitopes residues** | **Number of residues** | **Score** |
| CTL-MPV-2 | 1 | R667,K669,H670 | 3 | 0.875 |
| CTL-MPV-2 | 2 | G1,I2,G3,D4,P5,V6,T7,C8,L9,K10,S11,G12,A13,I14,C15,H16,P17,V18,F19,C20,P21,R22,R23,Y24,K25,Q26,I27,G28,T29,C30,G31,L32,P33,G34,T35,K36,C37,C38,K39,K40,P41,E42,A43,A44,A45,K46,N47,Q48,R49,N50,A51,P52,R53,I54,T55,F56,G57,G58,P59,S60,G61,G62,G63,G64,S65,R66,S67,K68,Q69,R70,R71,P72,Q73,G74,L75,P76,N77,N78,T79,A80,S81,W82,F83,T84,A85,L86,T87,H89,K91,G92,G93,G94,G95,S96,N97,S98,S99,P100,D101,D102,Q103,I104,G105,Y106,Y107,I114,D118,G119 | 108 | 0.777 |
| CTL-MPV-2 | 3 | Q526,T527,Y528,G529,G530,G531,G532,S533,A534,G547,S548,V549,D550,T551,D552,F553,V554,N555,E556,F557,Y558,A559,Y560,L561,H564,F565,S566,M567,G569,A577,N578,G579,Q580,V581,F582,G583,L584,Y585,G586,G587,G588,G589,S590,I591,P592,L593,M594,Y595,K596,G597,L598,P599,W600,N601,V602,V603,R604,G605,G606,G607,G608,S609,Y610,V611,M612,H613,A614,N615,Y616,I617,F618,W619,E620,A621,A622,A623,K624,G625,I626,I627,N628,T629,L630,Q631,K632,Y633,Y634,C635,R636,V637,R638,G639,G640,R641,C642,A643,V644,L645,S646,C647,L648,P649,K650,E651,E652,Q653,I654,G655,K656,C657,S658,T659,R660,G661,R662,K663,C664,C665,R666,H671,H672,H673,H674,H675 | 124 | 0.737 |
| CTL-MPV-2 | 4 | E188,L189,I190,R191,Q192,G193,T194,D195 | 8 | 0.665 |
| CTL-MPV-2 | 5 | T247,F248,P249,P250,T251,E252,P253,K254,K255,G256,G257,G258,G259,G272,G273,G274,G275,S276,E277,T278,T292,Q293,V294,V295,D296,M297,M299,Y301,M311,G312,T313,L314,S315,Y316,E317,Q318,F319,G320,G321,L325,F330,L331,A332,Y333,I334,L335,F336,T337,R338,F339,F340,Y341,G343,I351,F352,F353,A354,S355,F356,Y357,Y358,W360,K361,S362,Y363,G364,G365,G366,G367,S375,S376,T377,F378,N379,V380,P381,M382,E383,G385,G386,G387,G388,S389,C390,R397,F398,R399,R400,A401,F402,G403,G404,G405,G406,L414,V415,P416,I419,I421,A422,Y436,V442,G444,G445,G446,G447,S448,L449,P450,S451,G465 | 111 | 0.626 |
| CTL MPV | **No.** | **B Cell Discontinuous Epitopes residues** | **Number of residues** | **Score** |
| CTL-MPV-3 | 1 | L9, K10, S11, I14, V18, F19, C20, P21, R22, R23, Y24, K25, Q26, I27, G28, T29, C30, G31, L32, P33, G34, T35, K36, C37, C38, K39, K40, P41, E42, A43, A44, A45, K46, S47, E48, E49, T50, G51, T52, L53, I54, V55, N56, S57, V58, L60, F61, F64, R231, G232, G233, G234, G235, S236, Y237, F238, C240, W241, Y274, L277, Y278, S279, T280, Q281, L282, S283, T284, D285, G287, V288, V291, E316, I317, L319, I320, I321, M322, R323, T324, F325, K326, V327, S328, I329, W330, N331, L332, D333, Y334, I335, I336, G337, G338, G339, G340, S341, M342, K343, I344, I345, L346, F347, L348, A349, L350, I351, T352, L353, A354, T355, C356, E357, L358, Y359, H360 | 115 | 0.743 |
| CTL-MPV-3 | 2 | I118, F119, T120, I121, G122, L126, K127, G128, G129, G130, G131, S132, T133, I134, P135, I136, R542, V543, G544, A545, R546, K547, S548, A549, P550, L551, I552, I564, D565, I566, G567, N568, Y569, T570, V571, N579, Q581, E582, P583, G586, S587, V590, R591, C592, S593, F594, Y595, E596, D597, F598, L599, E600, Y601, D603, V604, R605, V606, V607, L608, G612, S613, I617, F620, F624, Y627, S628, L629, L630, L631, C632, R633, E634, A635, A636, A637, K638, G639, I640, I641, N642, T643, L644, Q645, K646, Y647, Y648, C649, R650, V651, R652, G653, G654, R655, C656, A657, V658, L659, S660, C661, L662, P663, K664, E665, E666, I668, G669, K670, C671, S672, T673, G675, R676, K677, C678, C679, R680, K682, K683, H684, H685, H686, H687, H688, H689 | 124 | 0.724 |
| CTL-MPV-3 | 3 | G188, G189, G190, S420, P421, F424, I425, R426, G427, G428, S431, E432, Q434, E435, L436, Y437, S438, P439, L481, L482, F483, L484, V485, L486, I487, M488, L489, I490, I491, F492, W493, F494, S495, L496, E497, L498, G499, G500, G501, G502, S503, M504, K505, F506, L507, V508, L510, G511, I512, T514, T515 | 51 | 0.716 |
| CTL-MPV-3 | 4 | L152, A153, V154, F155, Q156, S157, A158, S159, K160, I161, I162, T163, R167 | 13 | 0.713 |
| CTL-MPV-3 | 5 | N106, S107, S108, R109, G110, G111, G112, G113, S114, M116, R117, Q137 | 12 | 0.689 |
| CTL-MPV-3 | 6 | S304, L307, V308, D309, G362, G363, G364, G365, S366, L368, K369, E370, P371, C372, S373, S374, G375, T376, Y377, E378, G379, A387, D388, N389, K390, F391, A392 | 27 | 0.596 |

| | | | | |
|---|---|---|---|---|
| CTL-MPV-3 | 7 | T71, L75, T76, L78, R79, L80, C81, A82, Y83, G84, G85, G86, G522, G523, S524, Y525, V526, V527, D528, S537, K538 | 21 | 0.586 |

**Supplementary table S12:** B cell linear epitopes screened from HTL MPVs.

| HTL MPV | No. | Chain | Start | End | B Cell Linear Epitopes | Number of residues | Score |
|---|---|---|---|---|---|---|---|
| HTL-MPV-1 | 1 | _ | 273 | 312 | CGGGGSYADSFVIRGDEVRQIAPGQGGGGSVVVLSFELLH | 40 | 0.736 |
| HTL-MPV-1 | 2 | _ | 504 | 532 | YTGAIKLDDKDGGGGSSDFVRATATIPIQ | 29 | 0.783 |
| HTL-MPV-1 | 3 | _ | 184 | 211 | KGGGGSGINITRFQTLLALHRSYLTPGD | 28 | 0.749 |
| HTL-MPV-1 | 4 | _ | 129 | 155 | SQRVAGDSGGGGSSKTQSLLIVNNATN | 27 | 0.71 |
| HTL-MPV-1 | 5 | _ | 549 | 571 | ALVYFLQSINFVGGGGSDTGVEH | 23 | 0.783 |
| HTL-MPV-1 | 6 | _ | 234 | 256 | PTESIVRFPNITNLCPFGGGGGS | 23 | 0.672 |
| HTL-MPV-1 | 7 | _ | 461 | 482 | PQGTTLGGGGSWPQIAQFAPSA | 22 | 0.775 |
| HTL-MPV-1 | 8 | _ | 419 | 440 | GGSDDQIGYYRRATRRIRGGDG | 22 | 0.697 |
| HTL-MPV-1 | 9 | _ | 162 | 179 | EGGGGSREFVFKNIDGYF | 18 | 0.813 |
| HTL-MPV-1 | 10 | _ | 109 | 124 | LFARGGGGSSRTLSYY | 16 | 0.635 |
| HTL-MPV-1 | 11 | _ | 488 | 500 | MSRGGGGSTPSGT | 13 | 0.632 |
| HTL-MPV-1 | 12 | _ | 634 | 644 | CRRKKHHHHHH | 11 | 0.764 |
| HTL-MPV-1 | 13 | _ | 618 | 628 | PKEEQIGKCST | 11 | 0.614 |
| HTL-MPV-1 | 14 | _ | 579 | 587 | KIVDEPEEH | 9 | 0.809 |
| HTL-MPV-1 | 15 | _ | 449 | 457 | NPANNAAIV | 9 | 0.653 |
| HTL-MPV-1 | 16 | _ | 54 | 60 | LAFVVFL | 7 | 0.594 |
| HTL MPV | No. | Chain | Start | End | B Cell Linear Epitopes | Number of residues | Score |
| HTL-MPV-2 | 1 | _ | 567 | 591 | GSEILLIIMRTFKVSIWNLDYIINL | 25 | 0.793 |
| HTL-MPV-2 | 2 | _ | 646 | 663 | IFWFSLELGGGGSCTQHQ | 18 | 0.786 |
| HTL-MPV-2 | 3 | _ | 184 | 198 | EWFLAYILFTRFFYV | 15 | 0.782 |
| HTL-MPV-2 | 4 | _ | 690 | 757 | PFTIYSLLLCRMEAAAKGIINTLQKYYCRVRGGRCAVLSCLPKEEQIGKCSTRGRKCCRRKKHHHHHH | 68 | 0.761 |
| HTL-MPV-2 | 5 | _ | 22 | 44 | RRYKQIGTCGLPGTKCCKKPEAA | 23 | 0.752 |
| HTL-MPV-2 | 6 | _ | 1 | 15 | GIGDPVTCLKSGAIC | 15 | 0.749 |
| HTL-MPV-2 | 7 | _ | 210 | 219 | SYFAVHFISN | 10 | 0.728 |
| HTL-MPV-2 | 8 | _ | 248 | 265 | FYYVWKSYVGGGGSWLKQ | 18 | 0.712 |
| HTL-MPV-2 | 9 | _ | 385 | 402 | GGSQAIASEFSSLPSYAA | 18 | 0.698 |
| HTL-MPV-2 | 10 | _ | 456 | 475 | SFVNEFYAYLRKHFSMMILG | 20 | 0.674 |
| HTL-MPV-2 | 11 | _ | 421 | 434 | AKNRARTGGGGSRA | 14 | 0.667 |
| HTL-MPV-2 | 12 | _ | 294 | 306 | SAVGNICYTPSKL | 13 | 0.633 |
| HTL-MPV-2 | 13 | _ | 615 | 628 | ELYGGGGSLSLIDF | 14 | 0.579 |
| HTL-MPV-2 | 14 | _ | 556 | 562 | NPIQLSS | 7 | 0.555 |

**Supplementary table S13:** B cell Discontinuous epitopes screened from HTL MPVs.

| HTL MPV | No. | B Cell Discontinuous Epitope residues | Number of residues | Score |
|---|---|---|---|---|
| HTL-MPV-1 | 1 | M413, G419, G420, S421, D422, D423, Q424, I425, G426, Y427, Y428, R430, A431, T432, R433, R434, I435, R436, G437, G438, D439, G440, N449, P450, A451, N452, N453, A454, A455, I456, V457, L460, P461, Q462, G463, T464, T465, L466, G467, G468, G469, G470, S471, W472, P473, Q474, I475, A476, Q477, F478, A479, P480, S481, A482, S483, M488, S489, R490, G491, G492, G493, G494, S495, T496, P497, S498, | 121 | 0.727 |

| | | | | |
|---|---|---|---|---|
| | | G499, T500, Y504, T505, G506, A507, I508, K509, L510, D511, D512, K513, D514, G515, G516, G517, G518, S519, S520, D521, F522, V523, R524, A525, T526, A527, T528, I529, P530, I531, Q532, L547, A549, L550, V551, Y552, F553, L554, Q555, S556, I557, N558, F559, V560, G561, G562, G563, G564, S565, D566, T567, G568, V569, E570, V572 | | |
| HTL-MPV-1 | 2 | V261, L262, Y263, N264, S265, A266, S267, F268, C273, G274, G275, G276, G277, S278, Y279, A280, D281, S282, F283, V284, I285, R286, G287, D288, E289, V290, R291, Q292, I293, A294, P295, G296, Q297, G298, G299, G300, G301, S302, V303, V304, V305, L306, S307, F308, T331, N332 | 46 | 0.716 |
| HTL-MPV-1 | 3 | I596, N597, T598, L599 | 4 | 0.703 |
| HTL-MPV-1 | 4 | L54, A55, F56, V57, V58, F59, L60, K75, F78, L79, L82, W83, V85, T86, L87, L109, F110, A111, R112, G113, G114, G115, G116, S117, S118, R119, T120, L121, S122, Y124, A128, S129, Q130, R131, V132, A133, G134, D135, S136, G137, G138, G139, G140, S141, S142, K143, T144, Q145, S146, L147, L148, I149, V150, N151, N152, A153, T154, N155, E162, G163, G164, G165, G166, S167, R168, E169, F170, V171, F172, N174, I175, D176, G177, Y178, F179, K184, G185, G186, G187, G188, S189, G190, I191, N192, I193, T194, R195, F196, Q197, T198, L199, L200, A201, L202, H203, R204, S205, Y206, L207, T208, P209, G210, D211, A218, G219, A220, A221, Y227, G228, P234, T235, E236, I238, V239, R240, F241, P242, N243, I244, T245, N246, L247, C248, P249, F250, G251, G252, G253, G254, G255, S256 | 131 | 0.682 |
| HTL-MPV-1 | 5 | Y603, C604, V606, C634, R635, R636, H639, H640, H643, H644 | 10 | 0.675 |
| HTL-MPV-1 | 6 | G1, I2, G368, G369, G370, S371, R372, K579, I580, V581, D582, E583, P584, E585, E586, H587, P618, K619, E620, E621, Q622, G624, K625, C626, S627, T628, R629, G630, R631, C633, K637, K638, H641 | 33 | 0.652 |
| HTL-MPV-1 | 7 | F19, C20, P21, R22 | 4 | 0.52 |
| HTL-MPV-1 | 8 | E309, L310, L311, H312, P343, V344, S345, G346, G347, G348, G349, S350, Q351 | 13 | 0.517 |
| **HTL MPV** | **No.** | **B Cell Discontinuous Epitope residues** | **Number of residues** | **Score** |
| HTL-MPV-2 | 1 | F19,R23,K25,Q26,I27,G28,T29,C30,G31,L32,P33,G34,T35,K36,C37,C38,K39,K40,P41,E42,A43,A44,K46,V61,E62,T63,G64,G65,G66,G67,S68,E74,A76,V78,V79,R80,F83,S84,G85,G86,G87,G88,S89,T90,L91,E92,E93,T94,K95,F96,L97,T98,E99,N100,L101,L102,L103,Y104,I105,D106,I107,N108,G109,G110,G111,G112,S113,V114,Q115,Q116,E117,S118,P119,F120,V121,M122,M123,S124,A125,P126,P127,A128,Q129,Y130,E131,L132,G133,G134,G135,G136,S137,L138,L139,Q140,L141,C142,T143,F144,T145,S147,T148,N149,S150,R151,A154,S155,G157,G158,G159,G160,S161,S162,K163,L164,S183,E184,W185,F186,L187,A188,Y189,I190,L191,F192,T193,R194,F195,F196,Y197,V198 | 130 | 0.755 |
| HTL-MPV-2 | 2 | G453,S456,F457,V458,N459,E460,Y462,A463,Y464,L465,R466,K467,H468,F469,S470,M471,M472,I473,L474,G475,G476,G564,G565,G566,G567,S568,E569,I570,L571,L572,I573,I574,M575,R576,T577,F578,K579,V580,S581,I582,W583,N584,L585,D586,Y587,I588,I589,N590,L591,K594,Y617,G618,G619,G620,G621,S622,L623,S624,L625,I626,D627,F628,I646,F647,W648,F649,S650,L651,E652,L653,G654,G655,G656,G657,S658,C659,T660,Q661,H662,Q663,P664,Y683,V686,F687,P690,F691,T692,I693,Y694,S695,L696,L697,L698,C699,R700,E702,A703,A704,A705,K706,G707,I708,I709,N710,T711,L712,Q713,K714,Y715,C717,R718,V719,R720,G721,G722,R723,C724,A725,V726,L727,S728,C729,L730,P731,K732,E733,E734,Q735,I736,G737,K738,C739,S740,T741,R742,G743,R744,K745,C746,C747,R748,K750,K751,H752,H753,H755,H756,H757 | 148 | 0.724 |

| | | | | |
|---|---|---|---|---|
| HTL-MPV-2 | 3 | S294,A295,V296,G297,I299,C300,Y301,G384,G385,G386,S387,Q388,A389,I390,A391,S392,E393,F394,S395,S396,L397,P398,S399,Y400,A401,A402,K416,Y417,A418,A421,K422,R424,A425,R426,T427,G428,G429,G430,G431,S432,R433,A434 | 42 | 0.666 |
| HTL-MPV-2 | 4 | G1,I2,G3,D4,P5,V6,T7,C8,L9,K10,S11,G12,A13,I14,C15,V18,R22,M205,L207,S210,Y211,F212,A213,V214,H215,F216,I217,S218,N219,F248,Y249,V251,W252,K253,S254,Y255,V256,G257,G258,G259,G260,S261,W262,L263,K264,Q265,K268,L282,I283,T284,P285,V286,H287,G288,G289,G290,G291,S292,T302,P303,S304,K305,L306,T332,L334,G335,G336,G337,G338,S339,F340,T341,Q365,T367 | 74 | 0.638 |
| HTL-MPV-2 | 5 | N556,P557,I558,Q559,L560,S561,S562 | 7 | 0.555 |

**Supplementary table S14:** Analysis of codon-optimized cDNA of all the MPVs.

| S.No. | MPVs | GC content | CAI (Codon Adaptation Index) score | Tandem rare codons |
|---|---|---|---|---|
| 1 | CTL-MPV-1 | 67.84% | 1 | 0% |
| 2 | CTL-MPV-2 | 69.72% | 1 | 0% |
| 3 | CTL-MPV-3 | 66.22% | 1 | 0% |
| 4 | HTL-MPV-1 | 71.10% | 1 | 0% |
| 5 | HTL-MPV-2 | 66.58% | 1 | 0% |
| | Ideal values | 30-70% | 0.8-1.0 | <30% |
